# A phenotypic screen identifies a compound series that induces differentiation of acute myeloid leukemia cells *in vitro* and shows anti-tumour effects *in vivo*

**DOI:** 10.1101/2020.12.18.423389

**Authors:** Laia Josa-Culleré, Katrina S. Madden, Thomas J. Cogswell, Thomas R. Jackson, Tom S. Carter, Douzi Zhang, Graham Trevitt, Stephen G. Davies, Paresh Vyas, Graham M. Wynne, Thomas A. Milne, Angela J. Russell

## Abstract

Induction of differentiation is a promising therapeutic strategy against acute myeloid leukemia. However, current differentiation therapies are effective only to specific patient populations. To identify novel differentiation agents with wider efficacy, we developed a phenotypic high-throughput screen with a range of genetically diverse cell lines. From the resulting hits, one chemical scaffold was optimised in terms of activity and physicochemical properties to yield OXS007417, which was also able to decrease tumour volume in a murine in vivo xenograft model.

## Introduction

Acute myeloid leukemia (AML) is the most aggressive type of blood cancer, and the second most common leukemia in adults. AML arises from genetic mutation(s) which cause an arrest of differentiation at the progenitor or precursor stage, blocking production of downstream blood lineages. This leads to the accumulation of immature myeloid cells – blasts – in the bone marrow and peripheral blood, which disrupts normal haematopoiesis.^1^ AML can occur at all ages, but predominantly affects elderly people, with 80% of patients being over 60 years of age.^2,3^

AML can originate from accumulation of different combinations of genetic abnormalities, which underlies the highly heterogeneous nature of the disease. Different AML subtypes have been described, with the FAB^4^ and the WHO^5^ classifications being the most common ones. The standard of care for many years has been intensive chemotherapy with cytarabine in combination with anthracycline-derived antibiotics such as daunorubicin to induce apoptosis of AML blasts, which often cannot be tolerated by elderly patients. Other cytotoxic targeted therapies have had little success, as most select for specific clones, leading to selection pressure and resistance or relapse.

An alternative to cytotoxic agents is differentiation therapy. While initial attempts failed to induce clinical remission,^6,7^ a breakthrough in clinical oncology was achieved when all-transretinoic acid (ATRA) was shown to release myeloid blasts from differentiation arrest in the subset of AML patients with acute promyelocytic leukemia (APL). Previously APL had been the worst prognostic subsets of AML with complete remission (CR) rates of 40% after chemotherapy,^8^ however treatment with a combination of ATRA and arsenic trioxide induced CR in 90% of patients.^9^ Although this represents only 10% of the AML patient population, these encouraging results have resulted in growing interest in differentiation therapy research in AML.^10^ In the recent years, new therapeutic agents capable of overcoming the differentiation blockade have been described, such as DHODH,^11,12^ IDH1 and IDH2,^13,14^ FLT3^15^ and LSD1^16,17^ inhibitors, some of which target specific mutations. These examples suggest that inducing differentiation in AML could be more effective and less toxic than current therapies, however novel agents are needed which are effective for wider or more refractive patient populations.

To identify new agents capable of inducing differentiation to AML blasts, which do not target a specific mutation or patient subpopulation, a phenotypic medium throughput assay was developed and used to screen a library of drug-like small molecules. From the resulting hits, a medicinal chemistry optimisation program was initiated, ultimately to provide OXS007417. This small molecule was capable of inducing differentiation to a range of AML cells *in vitro*, had favourable pharmacokinetic properties and reduced tumour volume in a murine AML xenograft *in vivo* model.

## Materials and methods

Experimental procedures can be found in the Supplementary Information file.

## Results

### A phenotypic assay identifies novel small molecules that can differentiate AML cell lines

To find new small molecules that can differentiate AML cells, the cell-surface marker CD11b was chosen as it is upregulated upon myeloid differentiation regardless of the lineage that the cells specialise into. Three AML cell lines representing different population subtypes (HL-60, THP-1 and OCI-AML3) were treated with a compound library of 1,000 small molecules. Cells were stained with CD11b-PE and DAPI, and upregulation of CD11b in the live cells was detected via flow cytometry. For the initial screen, compounds were dosed at 10 μM for 4 days. The compounds that upregulated CD11b in all three cell lines by more than 10% were selected for further investigation. Firstly, to detect autofluorescent compounds, which could show an increase in fluorescence independent of CD11b upregulation, cells treated with the compounds were stained with isotype-PE. This would also identify compounds showing a non-specific increase of the antibody. Differentiation ability was then further investigated through morphology analysis and assessment of cell number and viability following compound treatment. These were performed in six AML cell lines with high genetic diversity (HL-60, THP-1, OCI-AML3, Kasumi-1, KG-1 and ME-1).

Those compounds causing a specific CD11b increase, changes to morphology indicative of differentiation, and decrease of cell numbers in all six cell lines were resynthesised. The compounds where the biological activity was confirmed with the authentic solid sample were considered to be confirmed hits.

One of the confirmed hits of the screen was imidazopyridine OXS000675 (Figure 2A), which induced differentiation in all six AML cell lines (Figure 2B-F, Figures S1-S8). After a four-day treatment, OXS000675 increased CD11b by up to 92%, with EC_50_ values of 2.3 – 7.6 μM. The morphology of the cells treated with OXS000675 showed clear changes indicative of differentiation as compared to negative controls, such as a pronounced increase in size and granularity. Also, the number of viable cells stopped increasing after 24 h, consistent with a block in proliferation.

**Figure 1.**
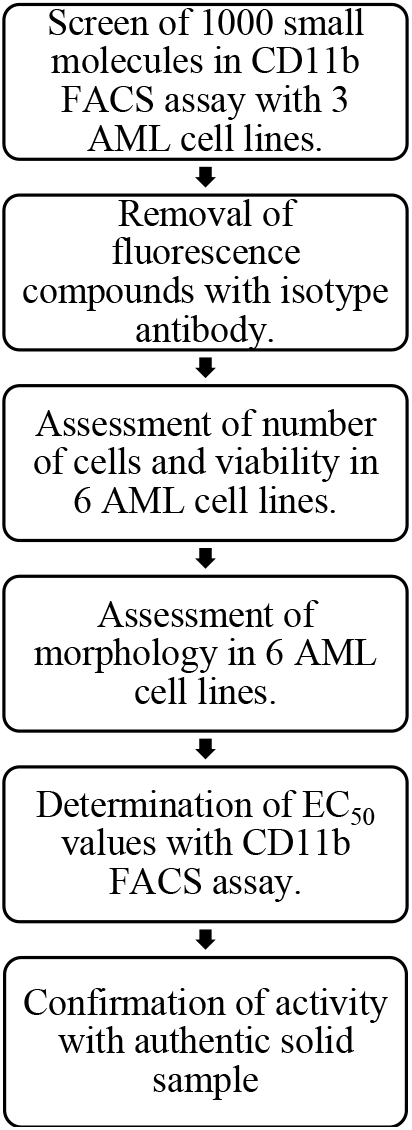
Screen workflow from primary assay to confirmed hits.

**Figure 2.**
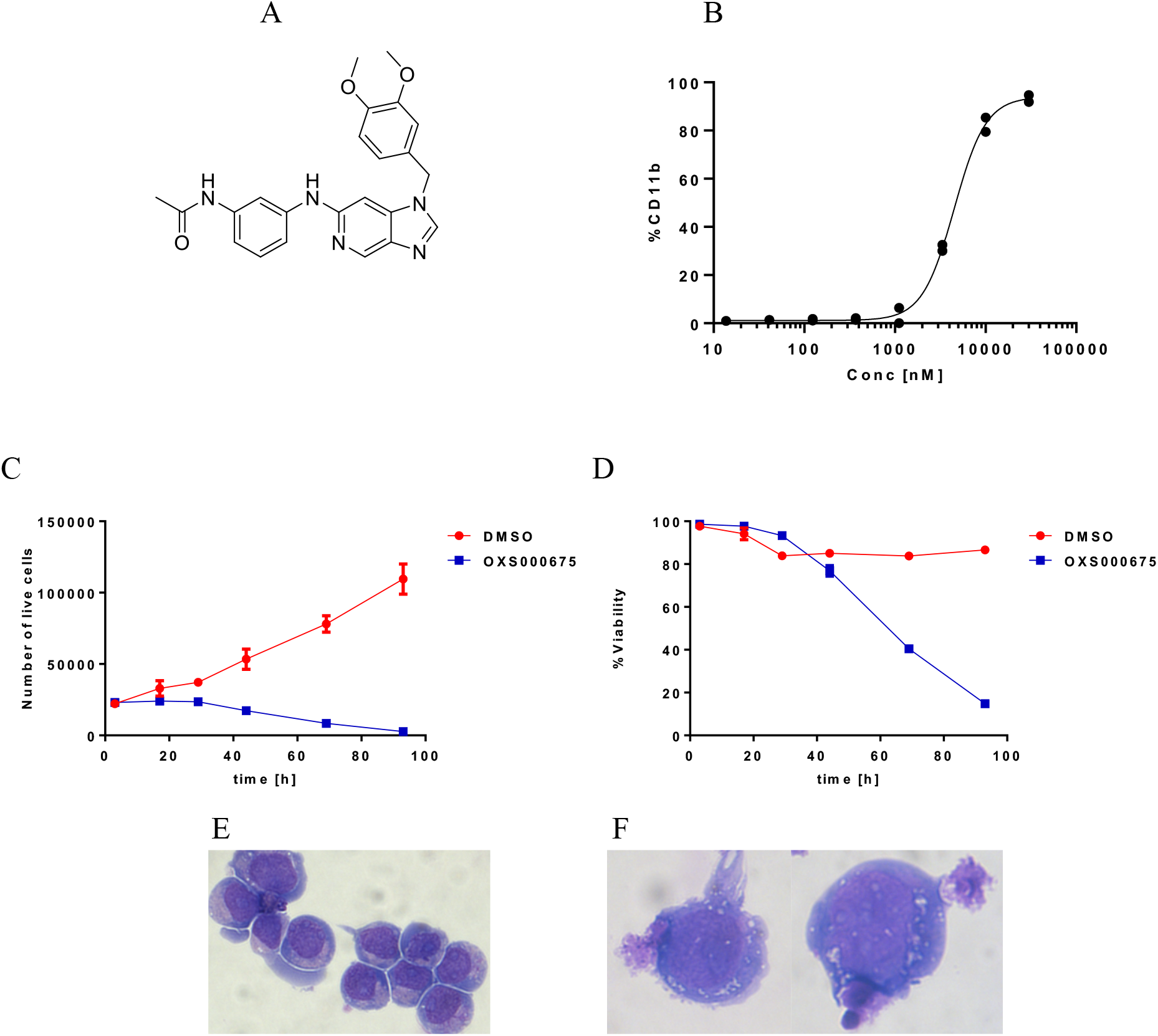
*In vitro* profile of the confirmed hit OXS000675 in HL-60 cells. A) Structure of OXS000675; B) Concentration-response curve of %CD11b determined by flow cytometry, HL-60 cells treated with OXS000675; C) Number of live cells per well and D) %Viability of HL-60 cells treated with either DMSO control or 10 μM OXS000675 over four days, determined with acridine orange and DAPI; E) Morphology of HL-60 cells treated with DMSO control, and F) OXS000675 10 μM.

### Optimisation of the activity and physicochemical properties of OXS000675 identifies OXS007417

OXS000675 was weakly active in the CD11b assay (4.8 μM in HL-60 cells), had good aqueous solubility (>200 μM) and had reasonable metabolic stability in mouse S9 (mS9) liver fraction (t_1/2_ = 42 min). Its overall profile was deemed to be a good starting point for a medicinal chemistry programme. The initial optimisation was focussed on biological activity of the analogues evaluated in HL-60 cells only, and an initial assessment of physicochemical properties was based on metabolic stability and aqueous solubility. The metabolic stability of the analogues was assessed in mouse liver S9 fraction, and is indicated as extraction ratio (ER).^18^.

OXS000675 and analogues were easily accessed in 4 steps from 2,4-dichloro-5-nitropyridine (Scheme 1).^19^ The imidazopyridine N-1 substituent was first introduced by regioselective nucleophilic aromatic substitution of an amine on the 4-chloro position of the nitropyridine. Subsequent reduction of the nitro group and cyclisation with triethylorthoformate provided imidazopyridine **4**. This could then be derivatised at the C-6 position via either Buchwald-Hartwig or Suzuki-Miyaura coupling to give **5** and **6** respectively.

**Scheme 1.**
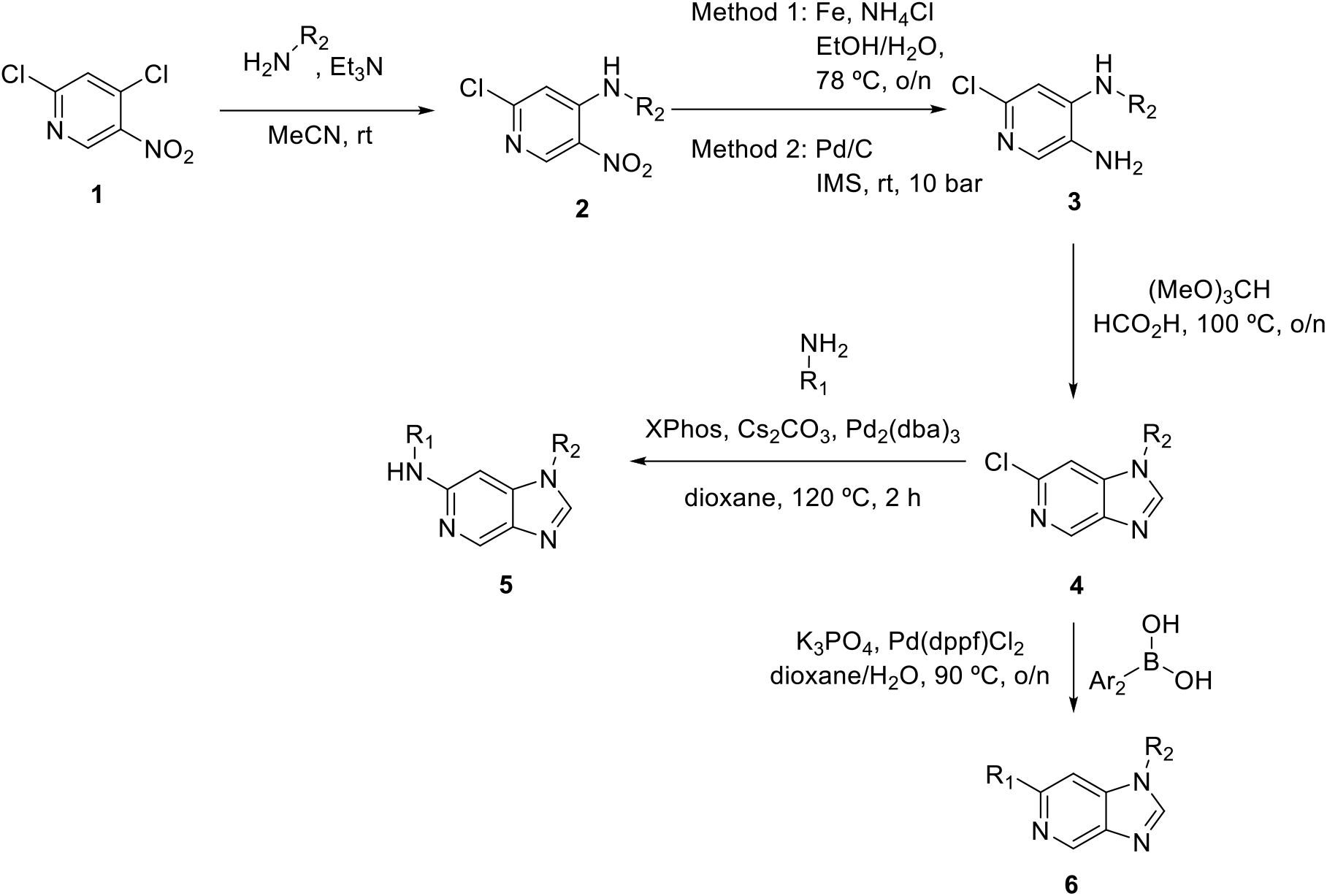
Synthetic route to analogues of OXS000675

Initial SAR studies on the N-1 position of the imidazopyridine core established that removal of the methylene linker was highly beneficial for activity, providing OXS007014 with a 35-fold increase in potency, but at the expense of metabolic stability. Pleasingly, a favourable level of solubility was retained (Table 1).

**Table 1.**
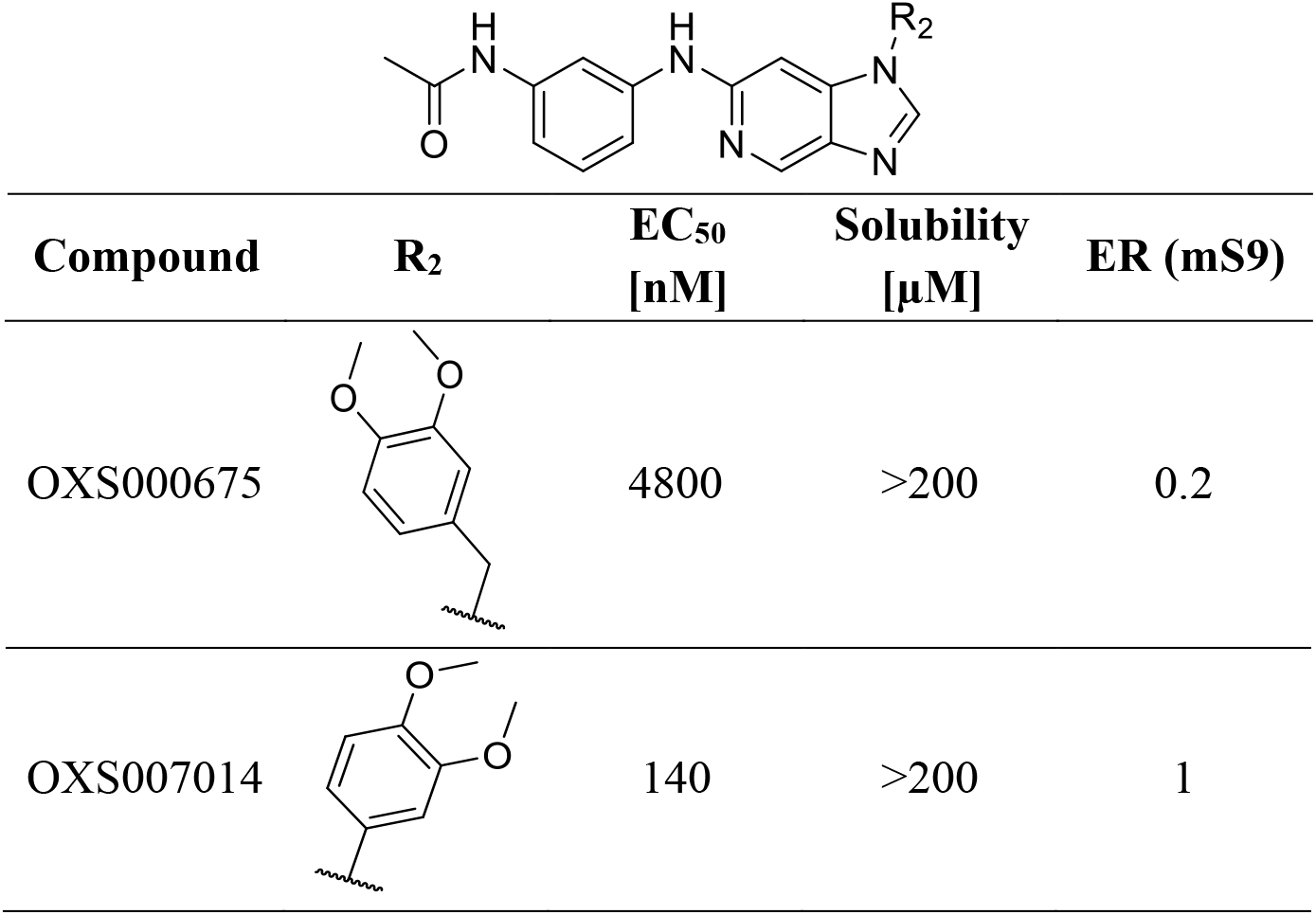
Structure-Activity Relationships of N-1 analogues

|  |  |  |  |  |
| --- | --- | --- | --- | --- |
| Compound | $\text{R}_2$ | $\text{EC}_{50}$<br>[nM] | Solubility<br>[$\mu\text{M}$] | ER (mS9) |
| OXS000675 |  | 4800 | >200 | 0.2 |
| OXS007014 |  | 140 | >200 | 1 |

**Table 2.**
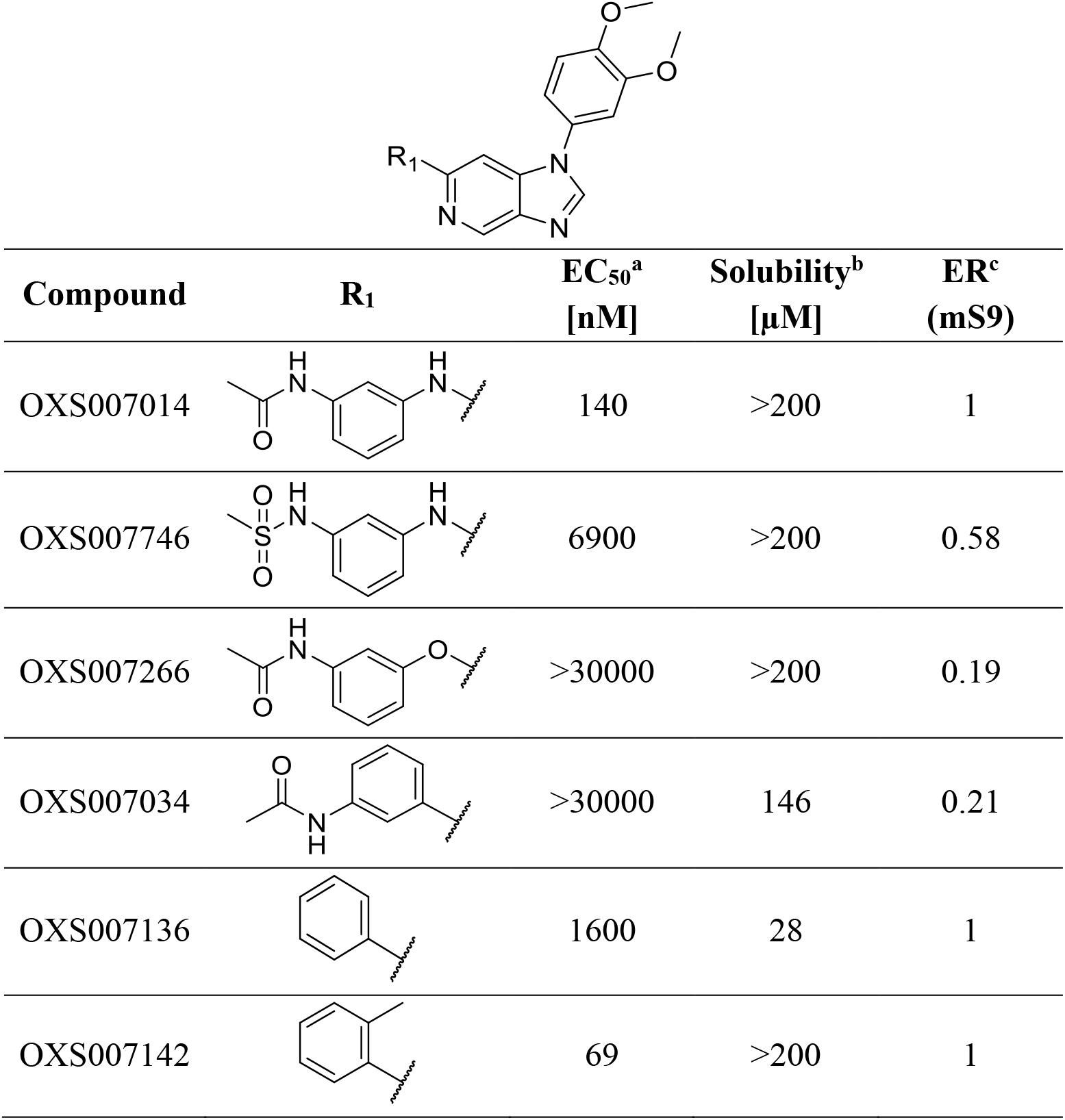
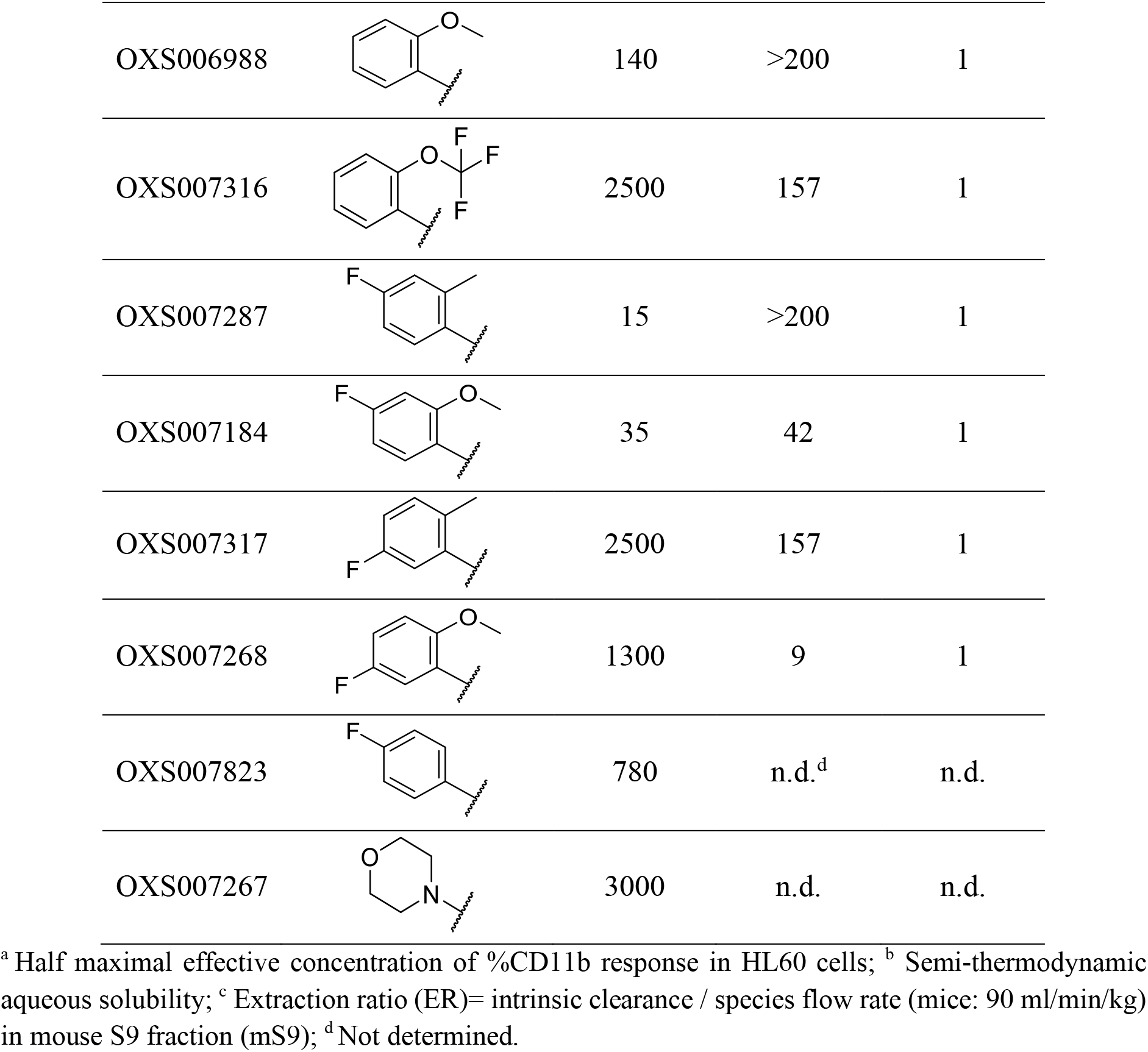
Structure-Activity Relationships of C-6 analogues

|  |  |  |  |  |
| --- | --- | --- | --- | --- |
| Compound | R<sub>1</sub> | EC<sub>50</sub><sup>a</sup><br>[nM] | Solubility<sup>b</sup><br>[μM] | ER<sup>c</sup><br>(mS9) |
| OXS007014 |  | 140 | >200 | 1 |
| OXS007746 |  | 6900 | >200 | 0.58 |
| OXS007266 |  | >30000 | >200 | 0.19 |
| OXS007034 |  | >30000 | 146 | 0.21 |
| OXS007136 |  | 1600 | 28 | 1 |
| OXS007142 |  | 69 | >200 | 1 |
| OXS006988 |  | 140 | >200 | 1 |
| OXS007316 |  | 2500 | 157 | 1 |
| OXS007287 |  | 15 | >200 | 1 |
| OXS007184 |  | 35 | 42 | 1 |
| OXS007317 |  | 2500 | 157 | 1 |
| OXS007268 |  | 1300 | 9 | 1 |
| OXS007823 |  | 780 | n.d.<sup>d</sup> | n.d. |
| OXS007267 |  | 3000 | n.d. | n.d. |

<sup>a</sup> Half maximal effective concentration of %CD11b response in HL60 cells; <sup>b</sup> Semi-thermodynamic aqueous solubility; <sup>c</sup> Extraction ratio (ER)= intrinsic clearance / species flow rate (mice: 90 ml/min/kg) in mouse S9 fraction (mS9); <sup>d</sup> Not determined.

At the C-6 position, another significant gain in potency was achieved when the amine linker was removed, and the aryl was substituted in the *ortho* position. Both *o*-methyl and methoxy groups were well tolerated. An additional fluorine atom in the *para* position led to another significant enhancement in activity, taking compounds OXS007287 and OXS007184 into the low nM range (EC_50_ = 15 and 35 nM). This is in striking contrast to OXS007317 and OXS07268, which have the fluorine in the *meta* position instead and are only weakly active.

Despite the promising activity of these compounds, all the examples had high clearance in mS9 (ER 1). The most potent group on C-6 (*o*-Me-*p*-F, OXS007287) was chosen for the next generation of analogues, where attention was focussed on improving metabolic stability whilst maintaining activity (Table 3). It was hypothesised that this could be achieved through modification of electronic properties on the aromatic ring attached to N-1, aiming to reduce the overall electron density and substitute one or both of the putatively metabolically labile methoxy groups.

**Table 3.**
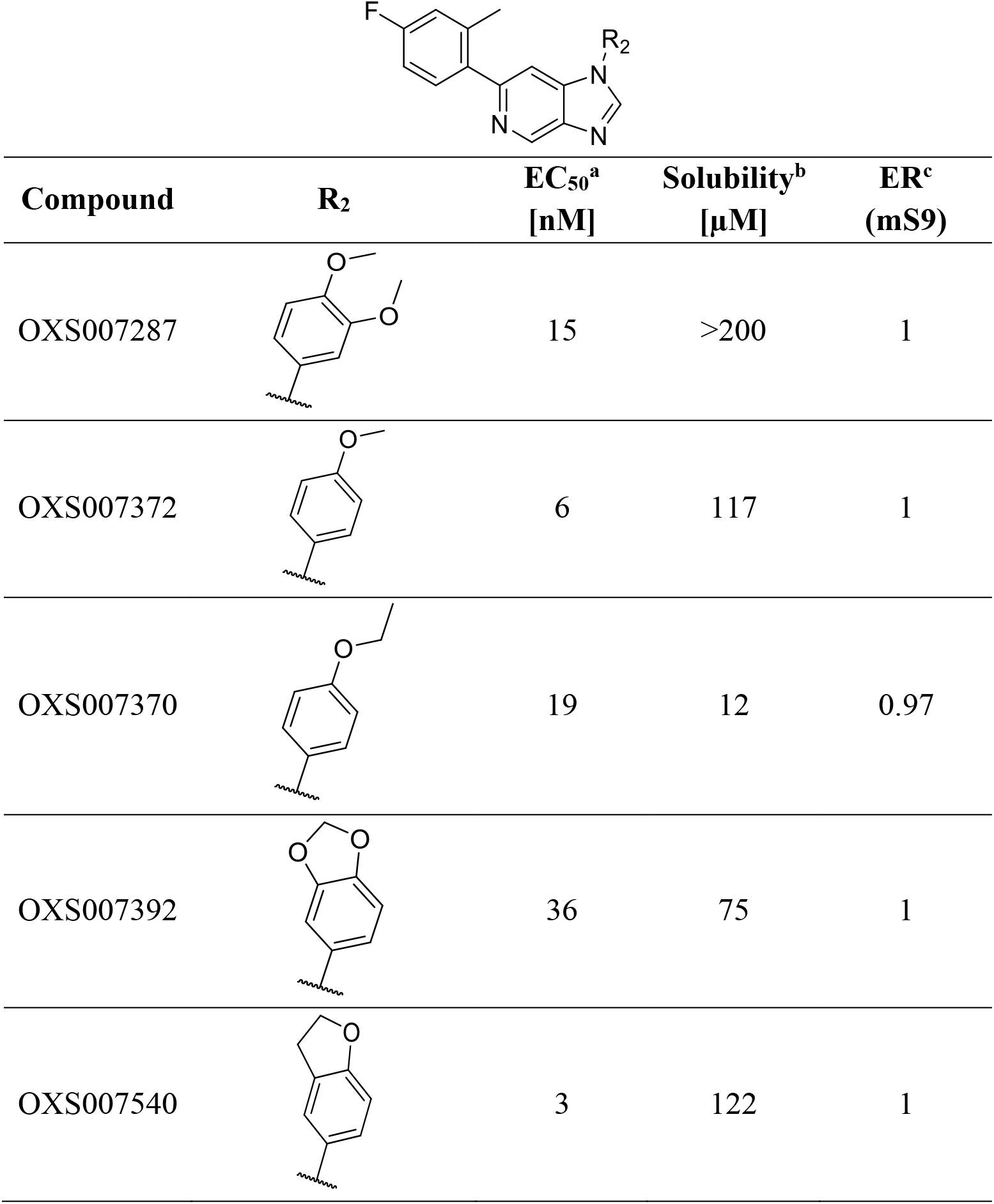
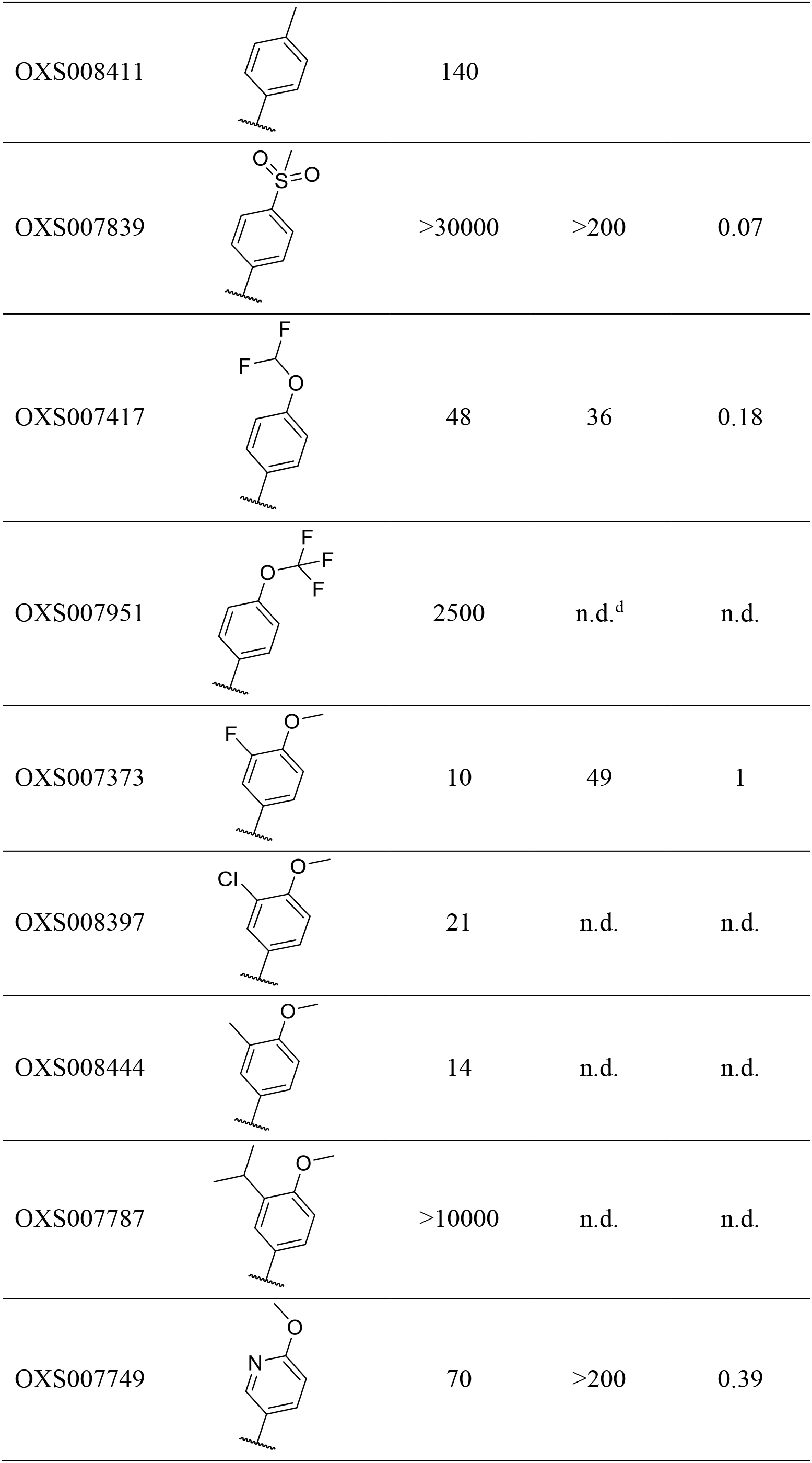
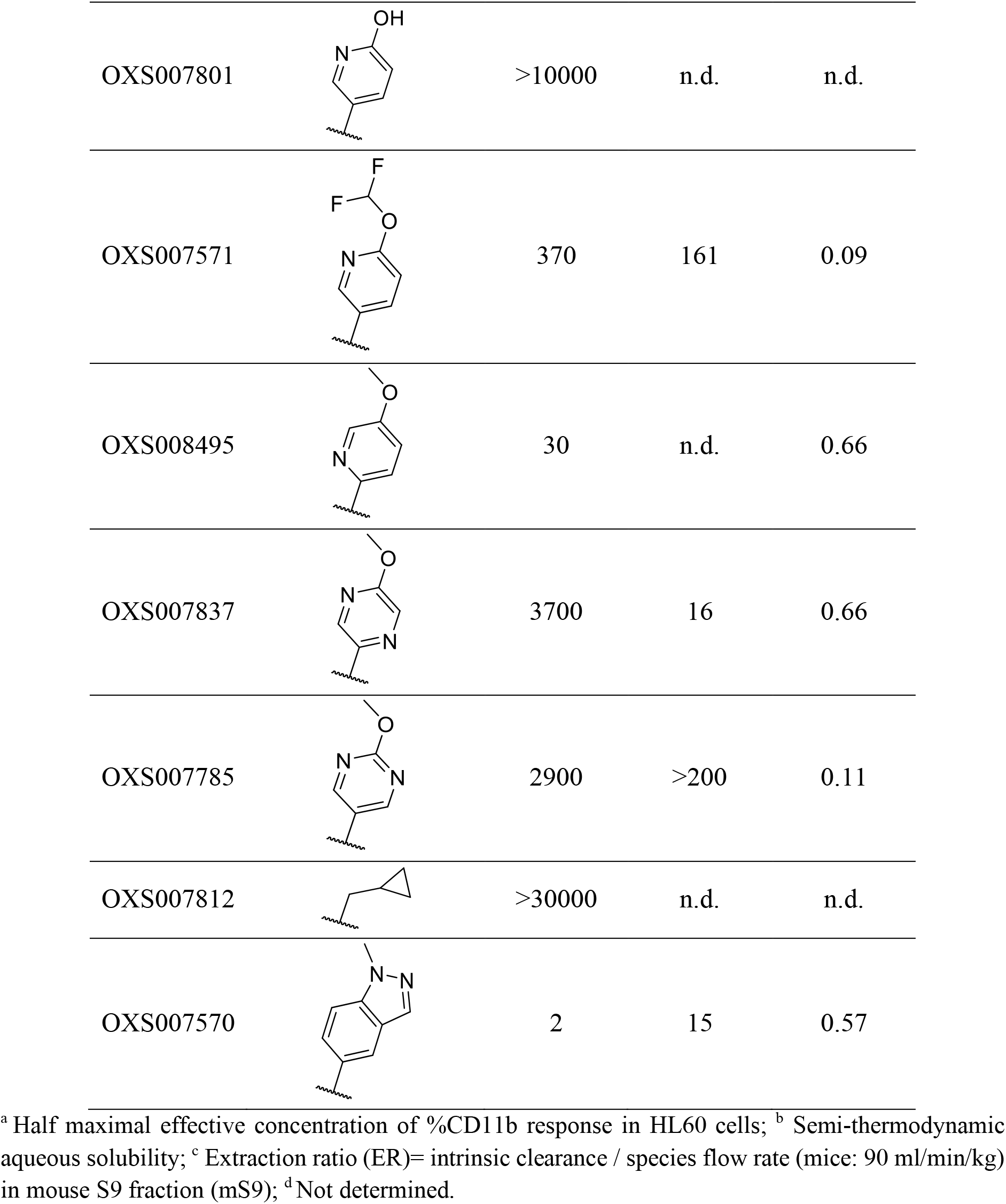
Structure-Activity Relationships of N-1 analogues

|  |  |  |  |  |
| --- | --- | --- | --- | --- |
| Compound | R<sub>2</sub> | EC<sub>50</sub><sup>a</sup><br>[nM] | Solubility<sup>b</sup><br>[μM] | ER<sup>c</sup><br>(mS9) |
| OXS007287 |  | 15 | >200 | 1 |
| OXS007372 |  | 6 | 117 | 1 |
| OXS007370 |  | 19 | 12 | 0.97 |
| OXS007392 |  | 36 | 75 | 1 |
| OXS007540 |  | 3 | 122 | 1 |
| OXS008411 |  | 140 |  |  |
| OXS007839 |  | >30000 | >200 | 0.07 |
| OXS007417 |  | 48 | 36 | 0.18 |
| OXS007951 |  | 2500 | n.d.<sup>d</sup> | n.d. |
| OXS007373 |  | 10 | 49 | 1 |
| OXS008397 |  | 21 | n.d. | n.d. |
| OXS008444 |  | 14 | n.d. | n.d. |
| OXS007787 |  | >10000 | n.d. | n.d. |
| OXS007749 |  | 70 | >200 | 0.39 |
| OXS007801 |  | >10000 | n.d. | n.d. |
| OXS007571 |  | 370 | 161 | 0.09 |
| OXS008495 |  | 30 | n.d. | 0.66 |
| OXS007837 |  | 3700 | 16 | 0.66 |
| OXS007785 |  | 2900 | >200 | 0.11 |
| OXS007812 |  | >30000 | n.d. | n.d. |
| OXS007570 |  | 2 | 15 | 0.57 |

<sup>a</sup> Half maximal effective concentration of %CD11b response in HL60 cells; <sup>b</sup> Semi-thermodynamic aqueous solubility; <sup>c</sup> Extraction ratio (ER)= intrinsic clearance / species flow rate (mice: 90 ml/min/kg) in mouse S9 fraction (mS9); <sup>d</sup> Not determined.

The *m*-methoxy group was found to not be required for biological activity; its removal led to a slight increase in potency, but unfortunately not in metabolic stability (OXS007372). Substitution of the *m*-position with either electron withdrawing (OXS007373, OXS008397) or sterically bulky (OXS007787) groups was not beneficial either. Adding nitrogens to the N-1 aromatic ring (OXS007749, OXS007837, OXS007785) improved metabolic stability, however this was at the cost of reduced biological activity. Encouragingly though, it was found that replacing the *p*-methoxy with a *p*-OCHF_2_ (OXS007417) gave a large improvement in metabolic stability, whilst causing only a modest drop in activity (EC_50_ = 48 nM). It was found that further reducing the electron density at this key group by incorporating an extra fluorine atom (OCF_3_, OXS007951) greatly reduced the activity. It was also found that replacement of the OCHF_2_ group with a structurally dissimilar electron withdrawing group such as a sulfone (OXS007839) eliminated biological activity, despite having an improved stability and solubility profile. OXS007417 seemed to give the best balance between biological activity and metabolic stability. The OCHF_2_ group appears to be a useful middle ground reducing the analogue’s susceptibility to metabolic oxidation, whilst preserving some accessibility of the oxygen lone pairs for bonding interactions.

The 1-methylindazole group was found to be an excellent bioisostere of the methoxy group, providing the highly potent OXS007570 with EC_50_ = 2 nM. However, it displayed poorer solubility and higher metabolic clearance than OXS007417.

With the more metabolically stable analogues in hand, substituent tolerability on the C-6 position was further analysed (Table 4).

**Table 4.**
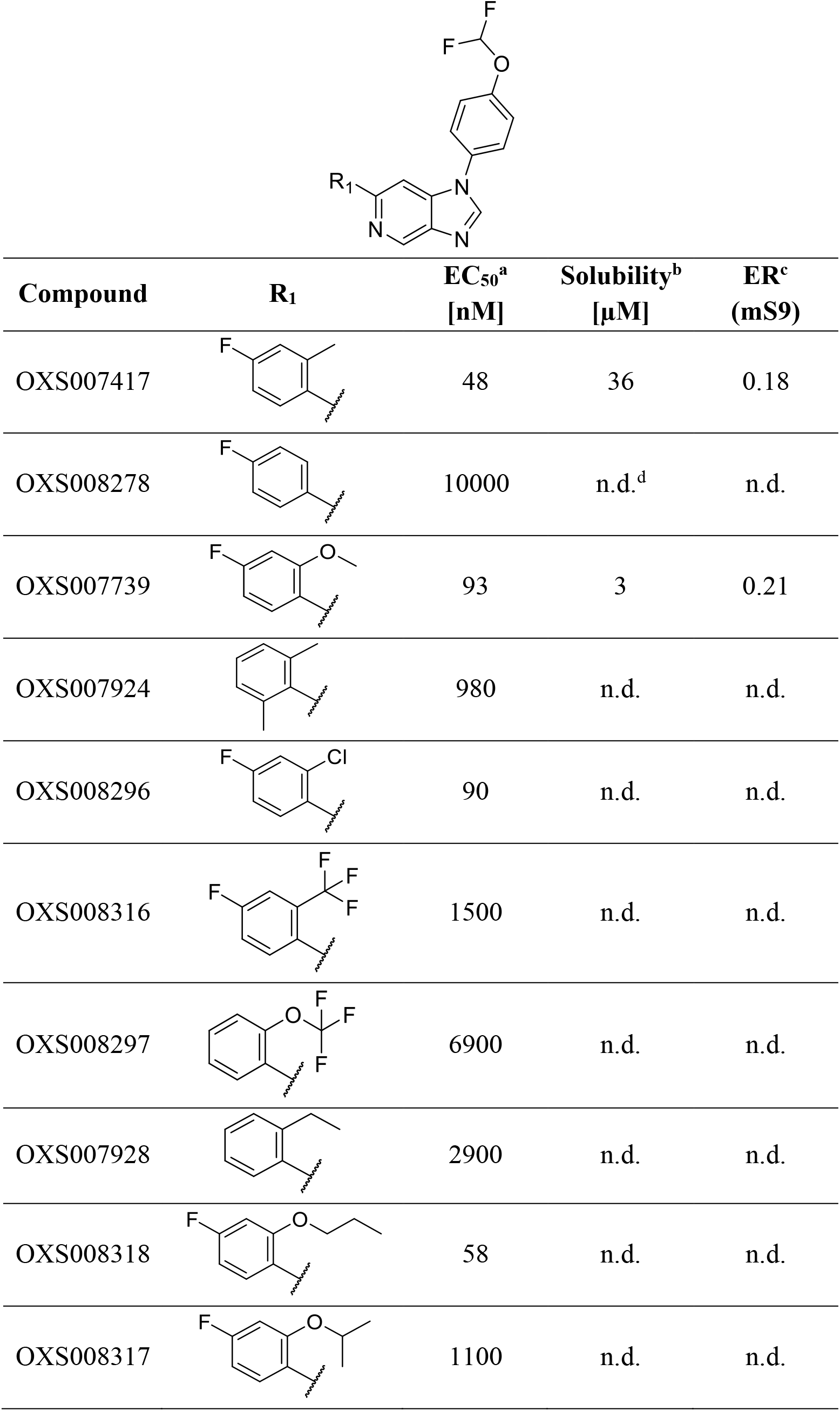
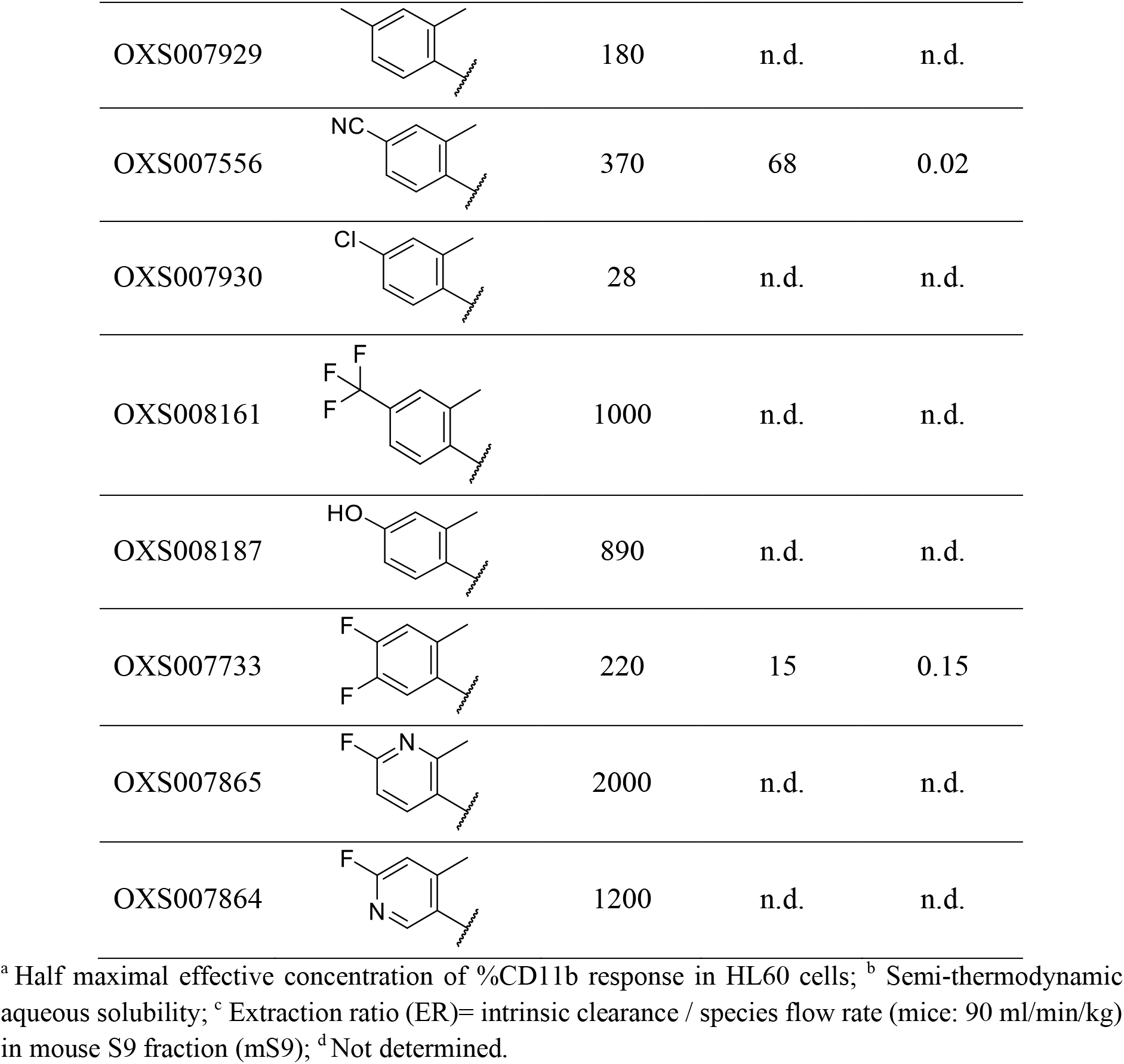
Structure-Activity Relationships of second-generation C-6 analogues

|  |  |  |  |  |
| --- | --- | --- | --- | --- |
| Compound | R<sub>1</sub> | EC<sub>50</sub><sup>a</sup><br>[nM] | Solubility<sup>b</sup><br>[μM] | ER<sup>c</sup><br>(mS9) |
| OXS007417 |  | 48 | 36 | 0.18 |
| OXS008278 |  | 10000 | n.d.<sup>d</sup> | n.d. |
| OXS007739 |  | 93 | 3 | 0.21 |
| OXS007924 |  | 980 | n.d. | n.d. |
| OXS008296 |  | 90 | n.d. | n.d. |
| OXS008316 |  | 1500 | n.d. | n.d. |
| OXS008297 |  | 6900 | n.d. | n.d. |
| OXS007928 |  | 2900 | n.d. | n.d. |
| OXS008318 |  | 58 | n.d. | n.d. |
| OXS008317 |  | 1100 | n.d. | n.d. |
| OXS007929 |  | 180 | n.d. | n.d. |
| OXS007556 |  | 370 | 68 | 0.02 |
| OXS007930 |  | 28 | n.d. | n.d. |
| OXS008161 |  | 1000 | n.d. | n.d. |
| OXS008187 |  | 890 | n.d. | n.d. |
| OXS007733 |  | 220 | 15 | 0.15 |
| OXS007865 |  | 2000 | n.d. | n.d. |
| OXS007864 |  | 1200 | n.d. | n.d. |

<sup>a</sup> Half maximal effective concentration of %CD11b response in HL60 cells; <sup>b</sup> Semi-thermodynamic aqueous solubility; <sup>c</sup> Extraction ratio (ER)= intrinsic clearance / species flow rate (mice: 90 ml/min/kg) in mouse S9 fraction (mS9); <sup>d</sup> Not determined.

At the *ortho*-position, the methyl group gave the best activity. Analogues containing a chloro OXS008296, methoxy OXS007739 and propoxy OXS008318 groups were also tolerated, while electron withdrawing groups (CF_3_, OCF_3_) led to a significant decrease in potency. At the *para* position, chloro substituted OXS007930 was equivalent to fluoro in terms of activity, but the latter was preferred as it was less lipophilic. Cyano containing OXS007556 was also well tolerated, and provided a further increase in metabolic stability and solubility. Pyridine derivatives OXS007865 and OXS007864 were prepared in an effort to reduce clogP, but the additional nitrogens were not tolerated.

The importance of the *ortho* group on the C-6 aryl unit is remarkable. It is clearly evidenced by comparing the *non-ortho* substituted OXS007136 and OXS008278, with the corresponding *o*-methyl-substituted OXS007142 and OXS007417, which are 23- and 21-fold more active respectively. It was hypothesised that this jump in potency could be due to steric interference between the methyl group and the imidazopyridine core, inducing a molecular twist which would reduce planarity in the molecules. This conformational change caused an advantage not only in activity, but also a gain in solubility, consistent with a less productive crystal packing of the less planar molecules. To provide further evidence, DFT optimisations were performed on OXS007417 and OXS008278. As shown on Figure 3, the methyl group of OXS007417 is predicted to cause rotation of the aryl group, increasing the dihedral angle between the core and the LHS, thus conveying greater three dimensional character onto the molecule. In an attempt to increase the dihedral angle, molecules with 2,6-substitution (OXS007924) or bulkier *ortho* groups (OXS008317) were prepared, but these led to a decrease in activity, suggesting that there was a size limitation within the target binding site.

**Figure 3.**
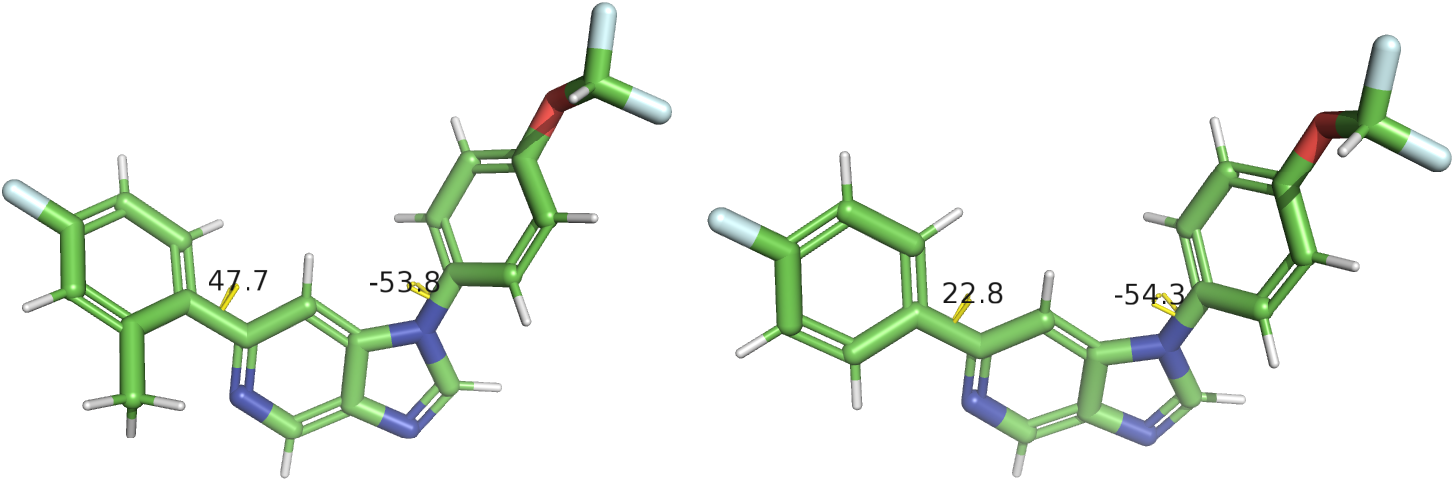
Predicted conformations of OXS007417 and OXS008278 from DFT energy minimisation. The dihedral angles between the bicyclic core and the N-1 and C-6 aryl groups are shown.

Substitution of the C-2 position of the imidazole was not well tolerated (Table 5).

**Table 5.**
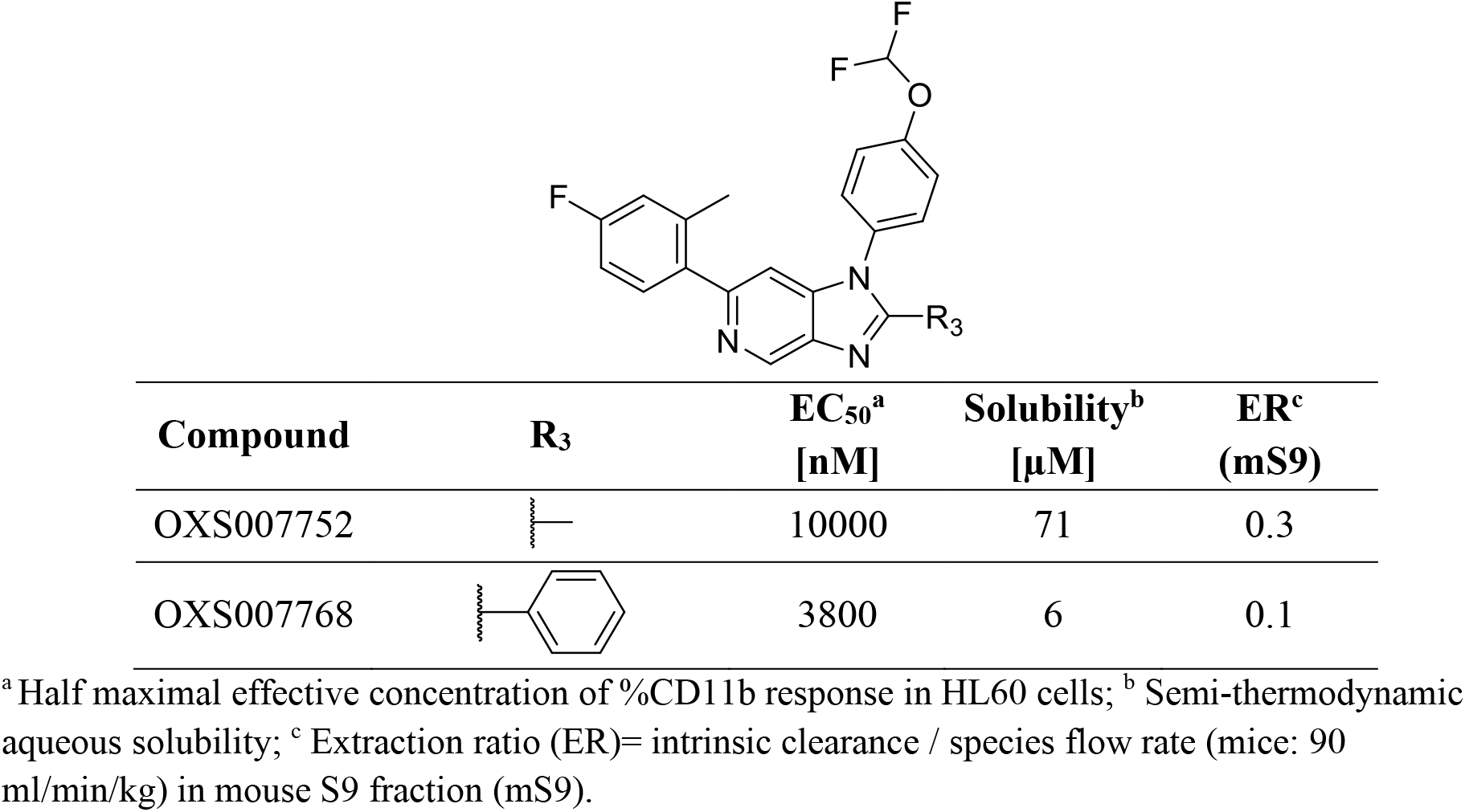
Structure-Activity Relationships of C-2 analogues

| Compound | R<sub>3</sub> | EC<sub>50</sub><sup>a</sup><br>[nM] | Solubility<sup>b</sup><br>[μM] | ER<sup>c</sup><br>(mS9) |
| --- | --- | --- | --- | --- |
| OXS007752 |  | 10000 | 71 | 0.3 |
| OXS007768 |  | 3800 | 6 | 0.1 |

<sup>a</sup> Half maximal effective concentration of %CD11b response in HL60 cells; <sup>b</sup> Semi-thermodynamic aqueous solubility; <sup>c</sup> Extraction ratio (ER)= intrinsic clearance / species flow rate (mice: 90 ml/min/kg) in mouse S9 fraction (mS9).

**Table 5.** Pharmacokinetic parameters of OXS007417 determined in the single dose 3 mg/kg, and repeat dose 60 mg/kg studies

| Parameter | 3 mg/kg<br>single dose | 60 mg/kg<br>repeat dose |  |
| --- | --- | --- | --- |
|  |  | Day 1 | Day 4 |
| $T_{1/2}^a$ (h) | 1.5 | | 8.1 |
| $AUC_{0-8h}^b$<br>(ng/mL.h) | 363 | 9860 | 9225 |
| $AUC_{0-8h}/dose$<br>(ng/mL.h<br>/(mg/kg)) | 121 | 164 | 154 |
| $C_{max}^c$ (ng/ml) | 128 | 2212 | 2675 |
| $C_{max}/dose$<br>(ng/mL/(mg/kg)) | 43 | 37 | 45 |
| $Cl_{obs}^d/F^e$<br>(ml/min/kg) | 129 | | |
| F (%) | 48 |  |  |

<sup>a</sup> Elimination half-life; <sup>b</sup> Area under the curve; <sup>c</sup> Maximum concentration; <sup>d</sup> Clearance; <sup>e</sup> Bioavailability.

Overall, OXS007417 gives the best balance of high potency, with reasonable metabolic stability and solubility, and was therefore chosen as a lead molecule for progression into further *in vitro* and *in vivo* studies (Figure 4). In spite of the moderate solubility at neutral pH, there was an improvement in acidic buffer (pH 1), probably due to the basic nature of the imidazopyridine core.

A more detailed *in vitro* ADME assessment of OXS007417 was undertaken. It showed reasonable stability in mouse hepatocytes (ER = 0.31), high cellular permeability measured in a Caco-2 assay (P_app_ = 14 x10^−6^ cm/s) and high plasma protein binding (PPB, 99% bound).

**Figure 4.**
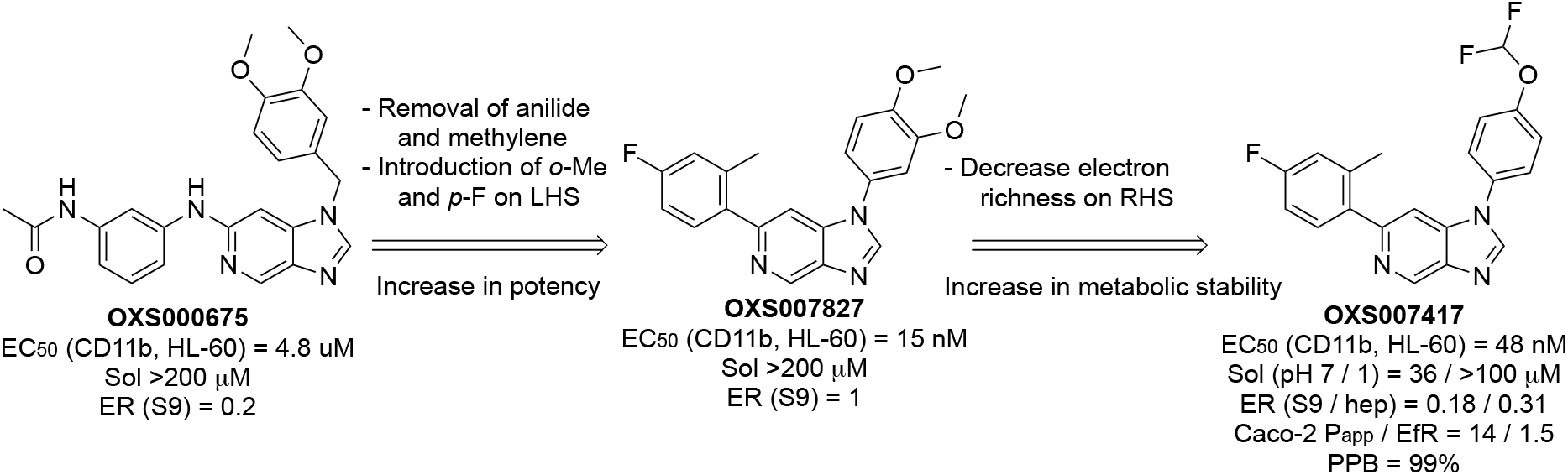
Overview of evolution from the hit OXS000675 to the *in vivo* lead molecule OXS007417.

### OXS007417 differentiates AML cell lines *in vitro*

Following the optimisation of the original hit molecule OXS000675 into OXS007417, it was important to verify that the lead molecule continued to be effective across multiple AML cell lines, and displayed a mechanistic profile consistent with differentiation.

Gratifyingly, CD11b flow cytometry analysis of THP-1 and OCI-AML3 cells treated with OXS007417 showed similar values to HL-60 (Figure 5A,B). Also, OXS007417 caused a clear decrease in cell numbers in the panel of six AML cell lines (Figure 5C). In some of the cell lines, and in particular ME-1, the decrease in viability was small, consistent with induction of differentiation and block in proliferation, rather than direct apoptosis. OXS007417 also caused morphological changes consistent with differentiation, such as a lighter cytoplasm, higher cytoplasm to nucleus ratio, increase in granulation and change on cell shape and size (Figure 5E-H and Figures S9-S14).

**Figure 5.**
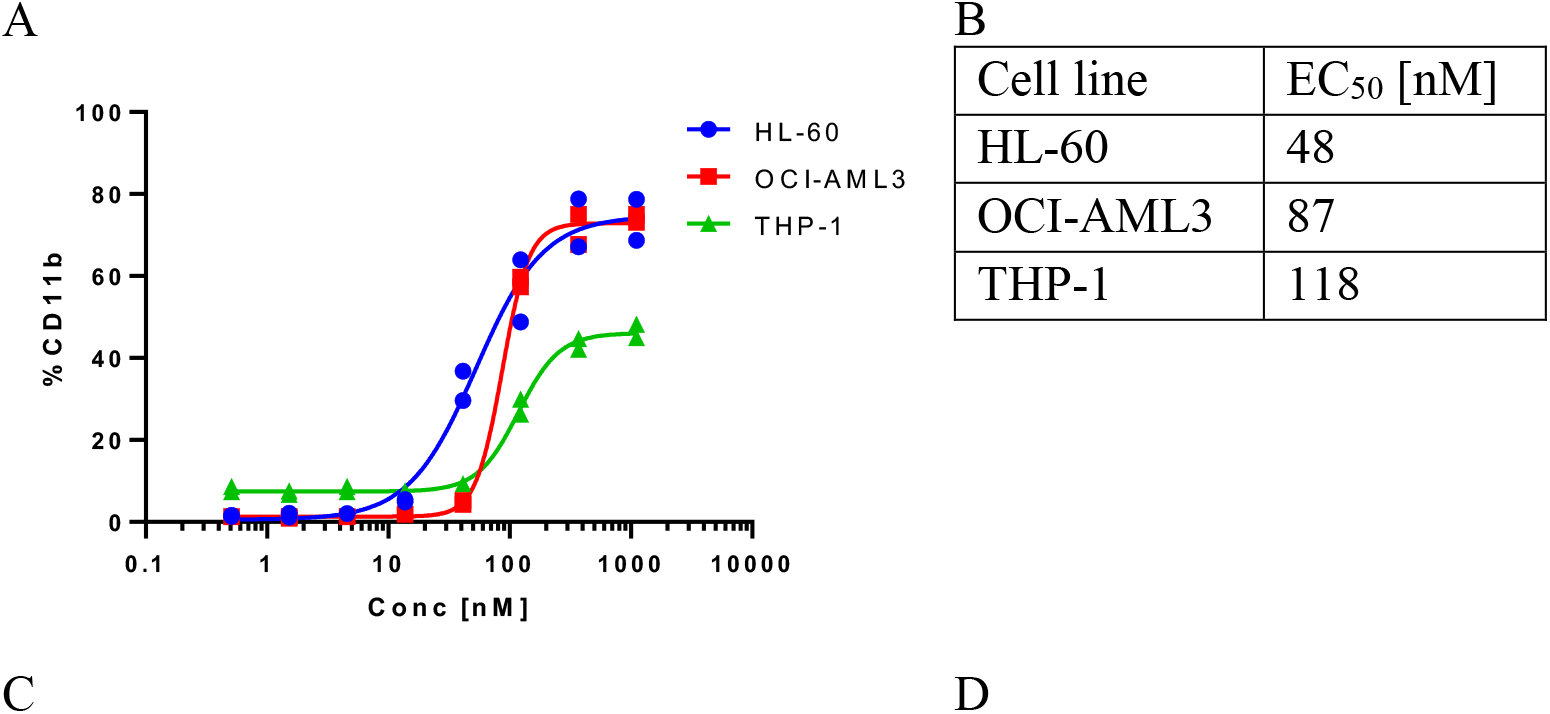
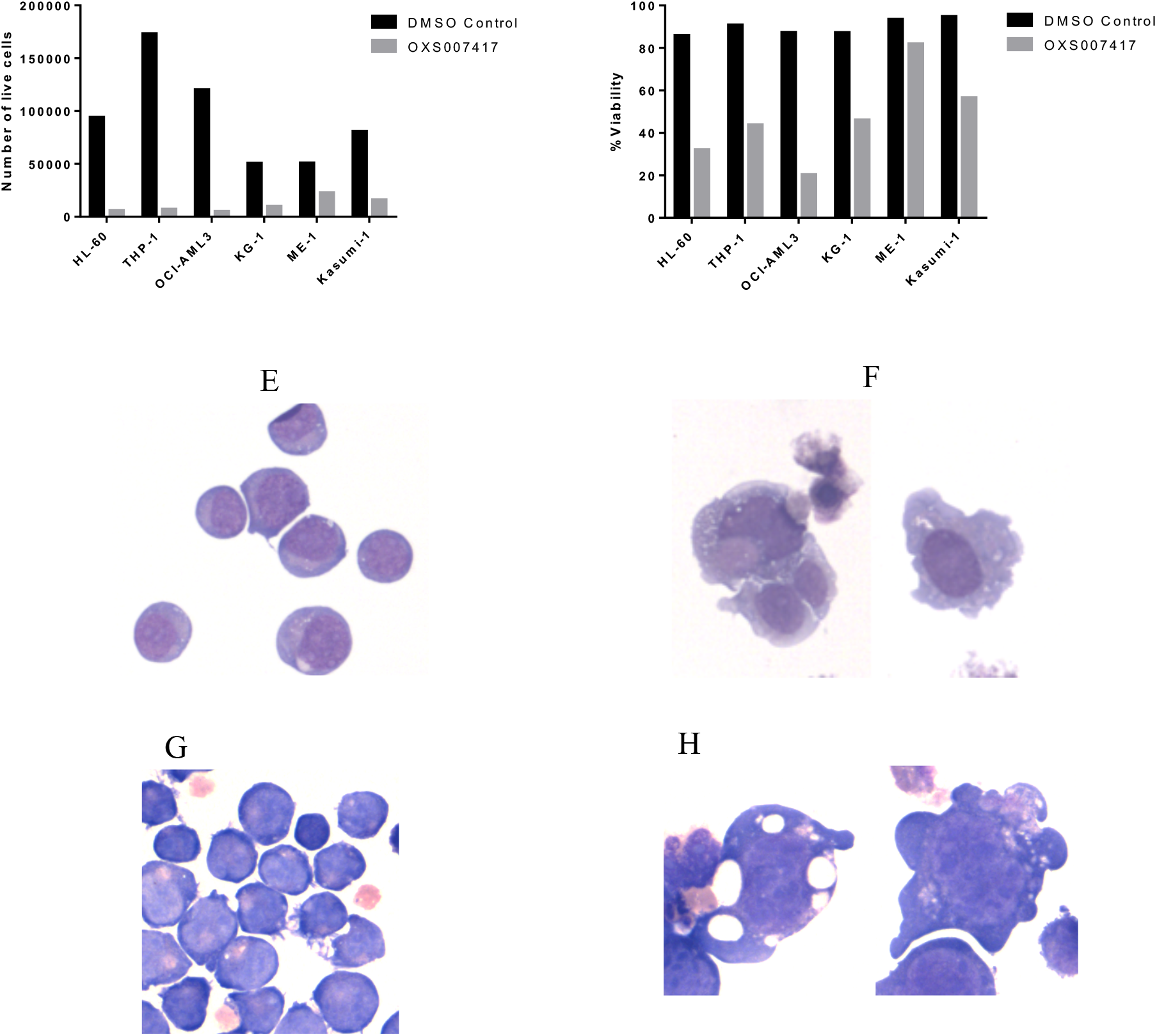
*In vitro* activity of OXS007417. A) Concentration-response curves of %CD11b determined by flow cytometry, HL-60, OCI-AML3 and THP-1 cells treated with OXS0007417, and B); C) Number of live cells per well and D) %viability of HL-60 cells treated with either DMSO control or 370 nM OXS007417 over 4 days, determined with acridine orange and PI; Morphology of HL-60 cells treated with E) DMSO control, F) OXS0007417 370 nM; Morphology of THP-1 cells treated with E) DMSO control, F) OXS0007417 370 nM

### OXS007417 has favourable pharmacokinetic properties and is well tolerated at a therapeutic dose

Next, the pharmacokinetic profile of OXS007417 was evaluated in male CD-1 mice. In the first study, OXS007417 was dosed at 3 mg/kg PO, and 1 mg/kg IV (Figure 6A, Table 5). The compound had moderate clearance (64 ml/min/kg) and volume of distribution (4.6 L/kg), leading to an elimination half-life of 1.5 h, and reasonable bioavailability (48%). At 3 mg/kg, the C_max_ was 128 ng/mL, which is 7-fold higher than the EC_50_ *in vitro*, and compound concentration above the EC_50_ was maintained for 5 h. It should be noted though that the high protein binding determined for OXS007417 (99%) means that the free drug available is likely to be significantly less than this.

**Figure 6.**
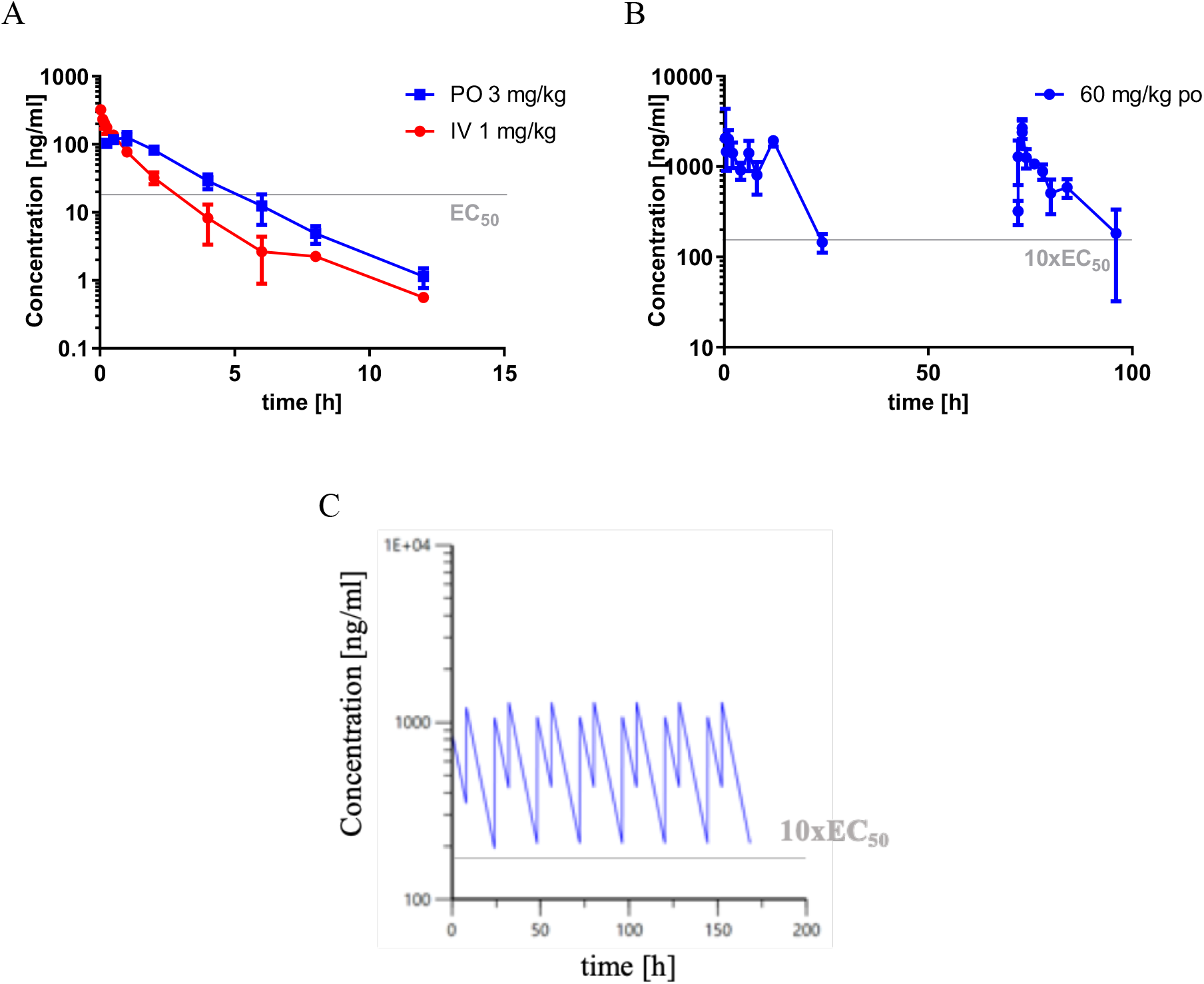
Pharmacokinetic characterisation of OXS007417. A) Observed blood-concentration profile in mice following single 3 mg/kg PO and 1 mg/kg IV dosing of OXS007417; B) Observed blood-concentration profile in mice following 60 mg/kg PO dosing of OXS007417 at days 1 and 4; C) Simulated exposure of OXS007417 at 30 mg/kg BID.

**Figure 7.**
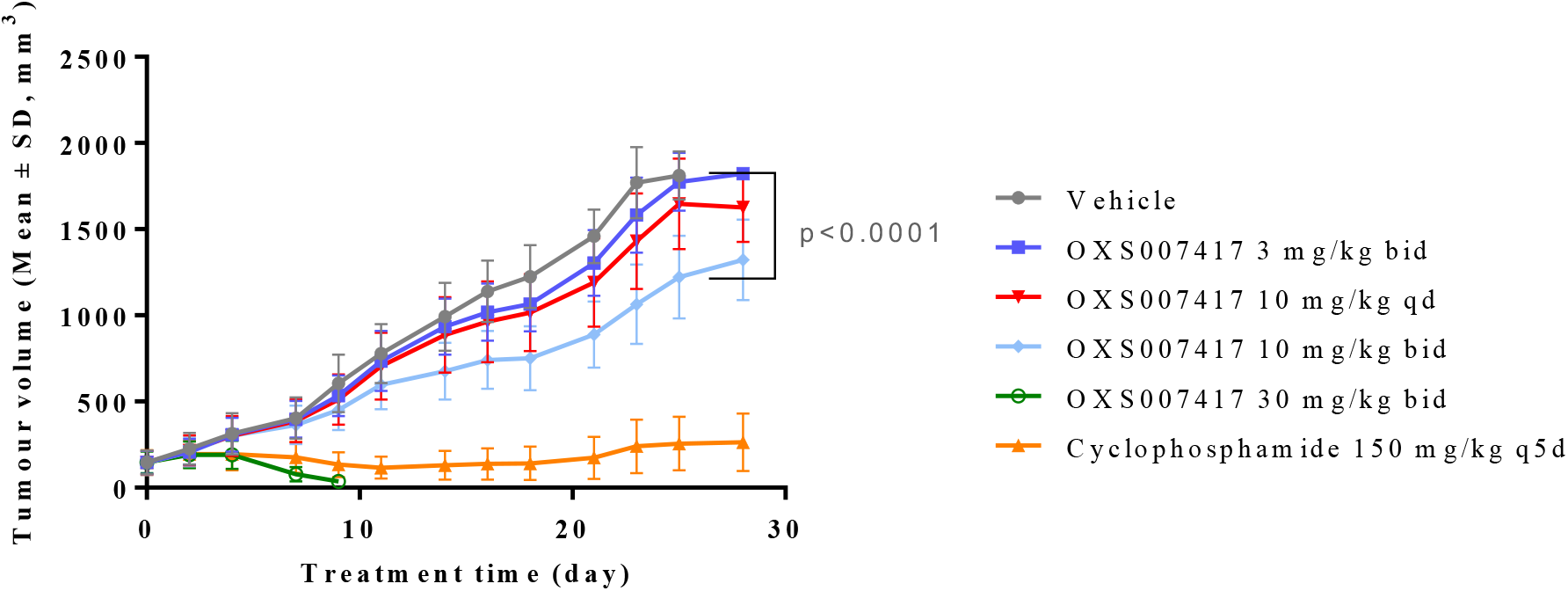
Evolution of tumour volume in the HL-60 subcutaneous model (female NOD SCID mice). Values shown are mean ±SD; n=10 at day 0 for all vehicle and OXS007417 groups, and n=5 for the cyclophosphamide group.

To maintain constant exposure levels above EC_50_ (48 nM = 18 ng/mL), repeat dosing at a higher compound dose was contemplated. A repeat dose study at 60 mg/kg was undertaken in CD-1 mice, where compound levels were measured on days 1 and 4 (Figure 6B). The study confirmed linear PK between the 3 and 60 mg/kg dose levels (Table 5), as indicated by both AUC and C_max_. The half-life, however, was significantly increased at 60 mg/kg, which could be indicative of saturation of clearance, or could reflect slower absorption of drug at the higher dose level. The similar exposure levels at days 1 and 4 indicated no accumulation of compound over time.

To estimate an appropriate dose level, the observed blood concentration-time data were fit to a single compartment PK model (Figure S15, see supplementary information for details) and then used to simulate the exposure over 1 week at 30 mg/kg twice daily (Figure 6C). The predicted values showed that at 30 mg/kg BID the compound exposure levels would be constantly maintained above 10xEC_50_. For subsequent efficacy studies, a dose of 10 mg/kg twice daily was considered to be appropriate to sustain sufficient exposure levels, whilst mitigating effects of protein binding.

In subsequent tolerability studies, dosing of compound for 7 days at 10 mg/kg BID was well tolerated with no apparent adverse effects.

Overall, PK and tolerance studies identified 10 mg/kg BID as a suitable efficacy testing dose for OXS007417, and demonstrated viability of maintaining exposure levels above EC_50_.

### OXS007417 reduces tumour volume in an HL-60 subcutaneous xenograft model

To study the ability of OXS007417 to inhibit tumour growth *in vivo*, a subcutaneous xenograft model was used. HL-60 cells were implanted onto the flank of female NOD SCID mice, and tumours were allowed to reach a volume of 150 mm^3^ before start of treatment. OXS007417 was administered PO at 3 mg/kg BID, 10 mg/kg QD and BID, and 30 mg/kg BID for 4 weeks. Cyclophosphamide (IP, 150 mg/kg Q5D), a clinical cytotoxic drug, was used as an internal control.

Treatment with OXS007417 was well tolerated at 3 and 10 mg/kg and did not lead to significant body weight loss. At 10 mg/kg twice daily dosing, OXS007417 significantly delayed the growth of HL-60 tumours, with T/C (ratio of tumour volume in treated mice versus control) of 57%, when compared to vehicle group (p<0.0001). A clear dose-dependent response was observed: at 10 mg/kg once daily dosing, the T/C was 79%, thus with a significant slowing of the growth of the tumours when compared to vehicle (p=0.0155), but to a lesser extent than at twice daily dosing. Tumours in 3 mg/kg BID treated animals followed a similar pattern to those treated with vehicle (p=0.3637).

Treatment at 30 mg/kg had a striking effect on tumour growth, almost diminishing tumour volume entirely on day 7. However, this dose was not well tolerated, and this arm of the study was discontinued at Day 9.

At the end of the study, blood samples were taken for bioanalysis at 1, 4 and 8 h following the final dose treatment. Exposure of OXS007417 was detected in all treatment groups; it was dose-dependent and decreased over time following compound administration. The exposure levels were also in agreement with those measured in the earlier PK studies.

## Discussion

Despite recent advances in the treatment of AML, resistance and relapse following treatment is still a major problem, and novel therapeutic approaches are imperative. Differentiation therapy is a promising alternative to the traditional cytotoxic chemotherapy; however, clinical application has so far only been successful in a small subset of patients. The high heterogeneity of the patients with AML challenges the search for a “one fits all” drug, but a differentiation agent effective in all AML patients regardless of their mutation status would be a major breakthrough.

In this work, a medium throughput phenotypic screen was developed to identify novel small molecules capable of differentiating AML cells. Phenotypic assays are particularly attractive in the search for drugs without a predefined target. To ensure that the effect was not limited to a specific patient population, a range of cell lines representing a variety of AML subtypes were utilised. CD11b was chosen as a marker of differentiation, and its upregulation was detected by flow cytometry. The hits were confirmed by assessing their effect on cell proliferation and morphology. One of the confirmed hits was OXS000675, which was effective in differentiating the panel of 6 cell lines, albeit with relatively modest potency (EC_50_ 2.2 – 7.8 μM).

A medicinal chemistry programme was undertaken to improve the potency and physicochemical properties of OXS000675. Removal of the amine and methylene linkers at C-6 and N-1 respectively led to a significant increase in potency. At the C-6 position, *ortho*-substitution was also critical for activity, and a *para-fluorine* atom was also beneficial. The importance of the *ortho* substituent was attributed to a twist of the molecule, which was supported by DFT calculations.

Despite the promising potency of these initial compounds, they lacked sufficient metabolic stability for further progression. Detailed investigation of the N-1 aromatic ring revealed that the OCHF_2_ group was capable of reducing electron density, thus increasing stability, while maintaining high activity.

The most promising compound OXS007417 was confirmed to induce differentiation *in vitro* in 6 AML cell lines, and it was progressed to *in vivo* studies. Pharmacokinetic studies at 3 mg/kg and 60 mg/kg, as well as tolerability studies, identified that dosing at 10 mg/kg twice daily would be a suitable dose.

In a subcutaneous study with HL-60 cells, treatment with OXS007417 at 10 mg/kg BID resulted in significant delay of tumour growth. A lower dose (3 mg/kg BID) was not effective, and a higher dose (30 mg/kg QD) was not well tolerated. Overall, this study identified an effective dose where OXS007417 was capable of inducing significant antitumour activity without adverse effects.

## Conclusion

In summary, a phenotypic screen was developed to identify novel small molecules capable of differentiating AML cells. A medicinal chemistry programme to optimise one of the hit molecules identified OXS007417, which induced differentiation of a range of AML cell lines *in vitro*. OXS007417 had favourable pharmacokinetic properties and showed anti-tumour effects *in vivo* in an AML subcutaneous model.

## Acknowledgements

We gratefully acknowledge Professor Malcolm MacCoss, Dr Alan Naylor and Dr Robert Westwood for helpful discussions. The xenograft study was performed by AxisBio Discovery Services, UK. We acknowledge Key Organics Ltd, UK for support with syntheses. This work was supported by OxStem Oncology (LJC, KSM, TJC, TRJ, TSC, DZ, GMW).

## References

(1) Khwaja, A.; Bjorkholm, M.; Gale, R. E.; Levine, R. L.; Jordan, C. T.; Ehninger, G.; Bloomfield, C. D.; Estey, E.; Burnett, A.; Cornelissen, J. J.; Scheinberg, D. A.; Bouscary, D.; Linch, D. C. Acute Myeloid Leukaemia. Nat. Rev. Dis. Prim. 2016, 2, 16010.

(2) Juliusson, G.; Antunovic, P.; Derolf, A.; Lehmann, S.; Mollgard, L.; Stockelberg, D.; Tidefelt, U.; Wahlin, A.; Hoglund, M. Age and Acute Myeloid Leukemia: Real World Data on Decision to Treat and Outcomes from the Swedish Acute Leukemia Registry. Blood 2009, 113 (18), 4179–4187.

(3) Ferrara, F.; Schiffer, C. A. Acute Myeloid Leukaemia in Adults. Lancet 2013, 381 (9865), 484–495.

(4) Bennett, J. M.; Catovsky, D.; Daniel, M.; Flandrin, G.; Galton, D. A. G.; Gralnick, H. R.; Sultan, C. Proposals for the Classification of the Acute Leukaemias French-American-British (FAB) Co-operative Group. Br. J. Haematol. 1976, 33 (4), 451–458.

(5) Ley, T. J.; Miller, C.; Ding, L.; Raphael, B. J.; Mungall, A. J.; Robertson, G.; Hoadley, K.; Triche, T. J.; Laird, P. W.; Baty, J. D.; Fulton, L. L.; Fulton, R.; Heath, S. E.; Kalicki-Veizer, J.; Kandoth, C.; Klco, J. M.; Koboldt, D. C.; Kanchi, K. L.; Kulkarni, S.; Lamprecht, T. L.; Larson, D. E.; Lin, G.; Lu, C.; McLellan, M. D.; McMichael, J. F.; Payton, J.; Schmidt, H.; Spencer, D. H.; Tomasson, M. H.; Wallis, J. W.; Wartman, L. D.; Watson, M. A.; Welch, J.; Wendl, M. C.; Ally, A.; Balasundaram, M.; Birol, I.; Butterfield, Y.; Chiu, R.; Chu, A.; Chuah, E.; Chun, H. J.; Corbett, R.; Dhalla, N.; Guin, R.; He, A.; Hirst, C.; Hirst, M.; Holt, R. A.; Jones, S.; Karsan, A.; Lee, D.; Li, H. I.; Marra, M. A.; Mayo, M.; Moore, R. A.; Mungall, K.; Parker, J.; Pleasance, E.; Plettner, P.; Schein, J.; Stoll, D.; Swanson, L.; Tam, A.; Thiessen, N.; Varhol, R.; Wye, N.; Zhao, Y.; Gabriel, S.; Getz, G.; Sougnez, C.; Zou, L.; Leiserson, M. D. M.; Vandin, F.; Wu, H. T.; Applebaum, F.; Baylin, S. B.; Akbani, R.; Broom, B. M.; Chen, K.; Motter, T. C.; Nguyen, K.; Weinstein, J. N.; Zhang, N.; Ferguson, M. L.; Adams, C.; Black, A.; Bowen, J.; Gastier-Foster, J.; Grossman, T.; Lichtenberg, T.; Wise, L.; Davidsen, T.; Demchok, J. A.; Mills Shaw, K. R.; Sheth, M.; Sofia, H. J.; Yang, L.; Downing, J. R.; Eley, G.; Alonso, S.; Ayala, B.; Baboud, J.; Backus, M.; Barletta, S. P.; Berton, D. L.; Chu, A. L.; Girshik, S.; Jensen, M. A.; Kahn, A.; Kothiyal, P.; Nicholls, M. C.; Pihl, T. D.; Pot, D. A.; Raman, R.; Sanbhadti, R. N.; Snyder, E. E.; Srinivasan, D.; Walton, J.; Wan, Y.; Wang, Z.; Issa, J. P. J.; Beau, M. Le; Carroll, M.; Kantarjian, H.; Kornblau, S.; Bootwalla, M. S.; Lai, P. H.; Shen, H.; Van Den Berg, D. J.; Weisenberger, D. J.; Link, D. C.; Walter, M. J.; Ozenberger, B. A.; Mardis, E. R.; Westervelt, P.; Graubert, T. A.; DiPersio, J. F.; Wilson, R. K. Genomic and Epigenomic Landscapes of Adult de Novo Acute Myeloid Leukemia. N. Engl. J. Med. 2013, 368 (22), 2059–2074.

(6) Nilsson, B. Probable in Vivo Induction of Differentiation by Retinoic Acid of Promyelocytes in Acute Promyelocytic Ieukaemia. Br. J. Haematol. 1984, 57, 365–371.

(7) Sampi, K.; Honma, Y.; Hozumi, M.; Sakurai, M. Discrepancy between In-Vitro and in-Vivo Inductions of Differentiation by Retinoids of Human Acute Promyelocytic Leukemia Cells in Relapse. Leuk. Res. 1985, 9 (12), 1475–1478.

(8) Wang, Z. Y.; Chen, Z. Acute Promyelocytic Leukemia: From Highly Fatal to Highly Curable. Blood. 2008, pp 2505–2515.

(9) Lo-Coco, F.; Avvisati, G.; Vignetti, M.; Thiede, C.; Orlando, S. M.; Iacobelli, S.; Ferrara, F.; Fazi, P.; Cicconi, L.; Di Bona, E.; Specchia, G.; Sica, S.; Divona, M.; Levis, A.; Fiedler, W.; Cerqui, E.; Breccia, M.; Fioritoni, G.; Salih, H. R.; Cazzola, M.; Melillo, L.; Carella, A. M.; Brandts, C. H.; Morra, E.; Von Lilienfeld-Toal, M.; Hertenstein, B.; Wattad, M.; Lübbert, M.; Hänel, M.; Schmitz, N.; Link, H.; Kropp, M. G.; Rambaldi, A.; La Nasa, G.; Luppi, M.; Ciceri, F.; Finizio, O.; Venditti, A.; Fabbiano, F.; Döhner, K.; Sauer, M.; Ganser, A.; Amadori, S.; Mandelli, F.; Döhner, H.; Ehninger, G.; Schlenk, R. F.; Platzbecker, U. Retinoic Acid and Arsenic Trioxide for Acute Promyelocytic Leukemia. N. Engl. J. Med. 2013, 369 (2), 111–121.

(10) De Thé, H. Differentiation Therapy Revisited. Nat. Rev. Cancer 2018, 18 (2), 117–127.

(11) Sykes, D. B.; Kfoury, Y. S.; Mercier, F. E.; Wawer, M. J.; Law, J. M.; Haynes, M. K.; Lewis, T. A.; Schajnovitz, A.; Jain, E.; Lee, D.; Meyer, H.; Pierce, K. A.; Tolliday, N. J.; Waller, A.; Ferrara, S. J.; Eheim, A. L.; Stoeckigt, D.; Maxcy, K. L.; Cobert, J. M.; Bachand, J.; Szekely, B. A.; Mukherjee, S.; Sklar, L. A.; Kotz, J. D.; Clish, C. B.; Sadreyev, R. I.; Clemons, P. A.; Janzer, A.; Schreiber, S. L.; Scadden, D. T. Inhibition of Dihydroorotate Dehydrogenase Overcomes Differentiation Blockade in Acute Myeloid Leukemia. Cell 2016, 167 (1), 171–186.e15.

(12) Christian, S.; Merz, C.; Evans, L.; Gradl, S.; Seidel, H.; Friberg, A.; Eheim, A.; Lejeune, P.; Brzezinka, K.; Zimmermann, K.; Ferrara, S.; Meyer, H.; Lesche, R.; Stoeckigt, D.; Bauser, M.; Haegebarth, A.; Sykes, D. B.; Scadden, D. T.; Losman, J. A.; Janzer, A. The Novel Dihydroorotate Dehydrogenase (DHODH) Inhibitor BAY 2402234 Triggers Differentiation and Is Effective in the Treatment of Myeloid Malignancies. Leukemia 2019.

(13) Lu, C.; Ward, P. S.; Kapoor, G. S.; Rohle, D.; Turcan, S.; Abdel-Wahab, O.; Edwards, C. R.; Khanin, R.; Figueroa, M. E.; Melnick, A.; Wellen, K. E.; Oĝrourke, D. M.; Berger, S. L.; Chan, T. A.; Levine, R. L.; Mellinghoff, I. K.; Thompson, C. B. IDH Mutation Impairs Histone Demethylation and Results in a Block to Cell Differentiation. Nature 2012, 483 (7390), 474–478.

(14) Wang, F.; Travins, J.; Delabarre, B.; Penard-lacronique, V.; Schalm, S.; Hansen, E.; Straley, K.; Kernytsky, A.; Liu, W.; Gliser, C.; Yang, H.; Gross, S.; Artin, E.; Saada, V.; Mylonas, E.; Quivoron, C.; Popovici-muller, J.; Saunders, J. O.; Salituro, F. G.; Yan, S.; Murray, S.; Wei, W.; Gao, Y.; Dang, L. Targeted Inhibition of Mutant IDH2 in Leukemia Cells Induces Cellular Differentiation. Science 2013, 340 (6132), 622–626.

(15) Sexauer, A.; Perl, A.; Yang, X.; Borowitz, M.; Gocke, C.; Rajkhowa, T.; Thiede, C.; Frattini, M.; Nybakken, G. E.; Pratz, K.; Karp, J.; Smith, B. D.; Levis, M. Terminal Myeloid Differentiation in Vivo Is Induced by FLT3 Inhibition in FLT3/ITD AML. Blood 2012, 120 (20), 4205–4214.

(16) Schenk, T.; Chen, W. C.; Göllner, S.; Howell, L.; Jin, L.; Hebestreit, K.; Klein, H. U.; Popescu, A. C.; Burnett, A.; Mills, K.; Casero, R. A.; Marton, L.; Woster, P.; Minden, M. D.; Dugas, M.; Wang, J. C. Y.; Dick, J. E.; Müller-Tidow, C.; Petrie, K.; Zelent, A. Inhibition of the LSD1 (KDM1A) Demethylase Reactivates the All-Trans-Retinoic Acid Differentiation Pathway in Acute Myeloid Leukemia. Nat. Med. 2012, 18 (4), 605–611.

(17) Hitchin, J. R.; Blagg, J.; Burke, R.; Burns, S.; Cockerill, M. J.; Fairweather, E. E.; Hutton, C.; Jordan, A. M.; McAndrew, C.; Mirza, A.; Mould, D.; Thomson, G. J.; Waddell, I.; Ogilvie, D. J. Development and Evaluation of Selective, Reversible LSD1 Inhibitors Derived from Fragments. Medchemcomm 2013, 4 (11), 1513–1522.

(18) Extraction Ratio (ER) Is Calculated as the Measured Intrinsic Clearance over the Hepatic Blood Flow of Each Species (90 Ml/Min/Kg for Mice). It Has the Advantage over Other Parameters (Half-Life, Clearance) That It Is Independent of the Species Used.

(19) Bryan, M. C.; Chan, B.; Hanan, E.; Heffron, T.; Purkey, H.; Elliott, R. L.; Heald, R.; Knight, J.; Lainchbury, M.; Seward, E. M. Aminopyrimidine Compounds as Inhibitors of T790M Containing EGFR Mutants. WO2014081718, 2014.

