## Supplementary Figures for "A phenotypic screen identifies a compound series that induces differentiation of acute myeloid leukemia cells *in vitro* and shows anti-tumour effects *in vivo*"

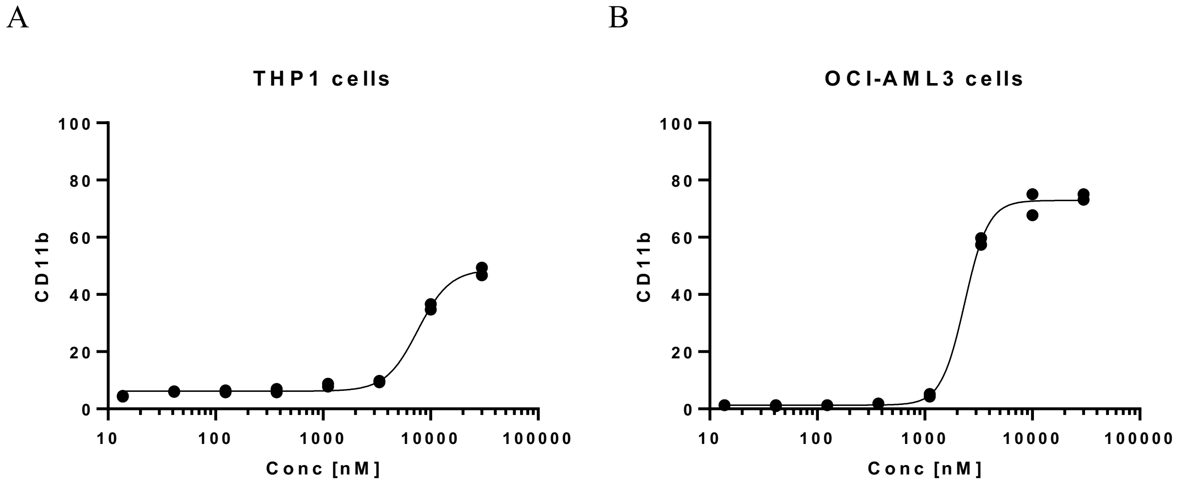


Figure S1. Concentration-response curves of OXS000675. %CD11b determined by flow cytometry. (A) THP1 cells ($EC_{50}$ 7.6 μM) and (B) OCI-AML3 cells ($EC_{50}$ 7.6 μM).

A

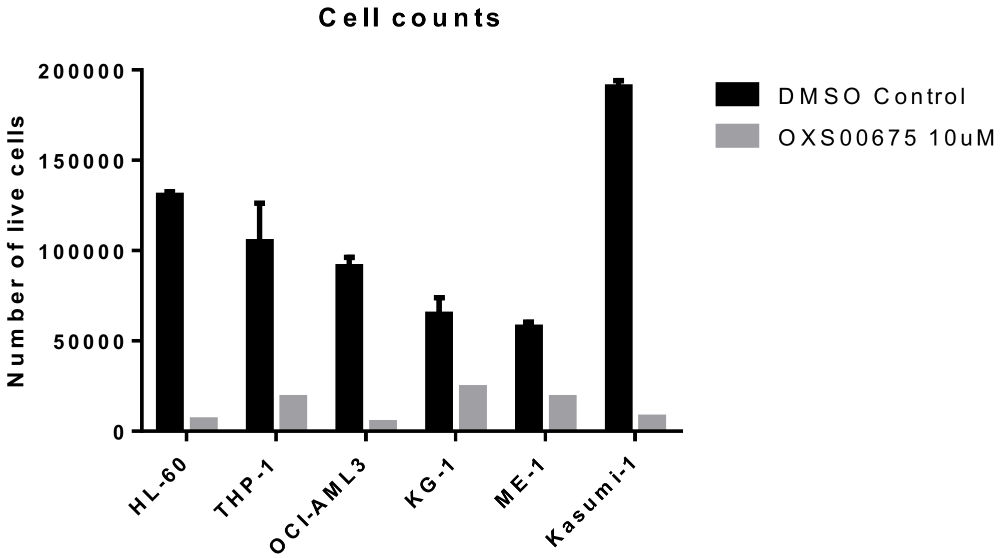


B

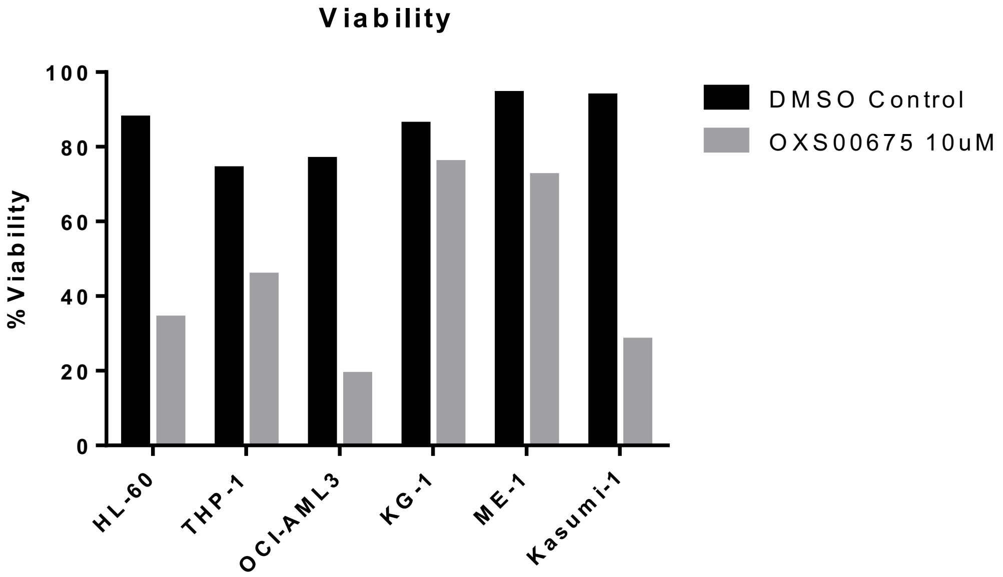


Figure S2. A) Number of live cells per well and B) %viability of cell lines treated with either DMSO control or 10 µM OXS000675 over 4 days, determined with acridine orange and DAPI.

A B

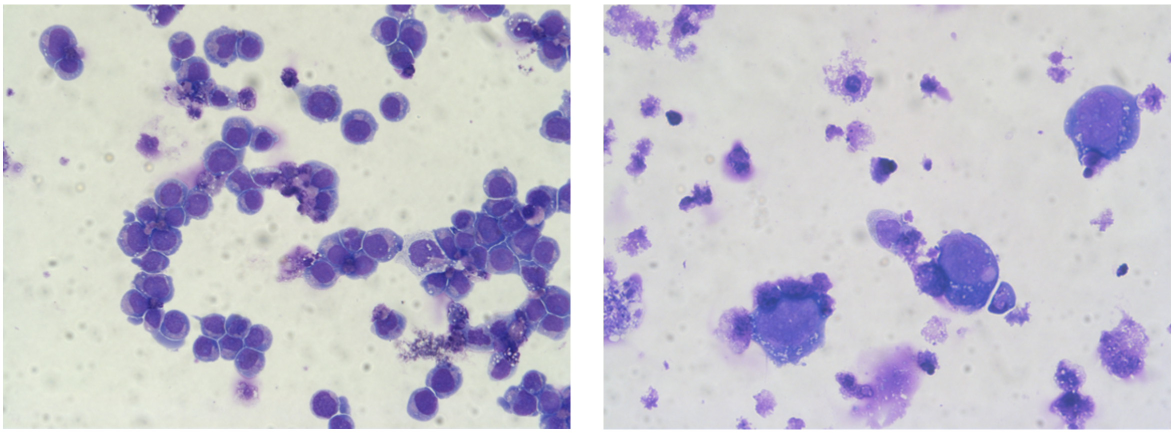

Figure S3. Cytospins of HL-60 cells treated with (A) DMSO control or (B) OXS000675 10 μM stained with Modified Wright stain.

A B

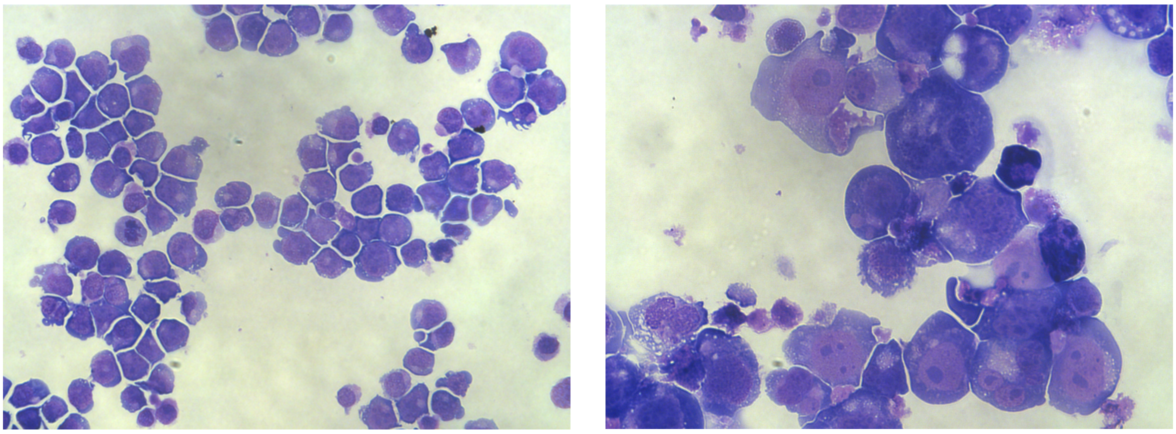

Figure S4. Cytospins of THP-1 cells treated with (A) DMSO control or (B) OXS000675 10 μM stained with Modified Wright stain.

A B

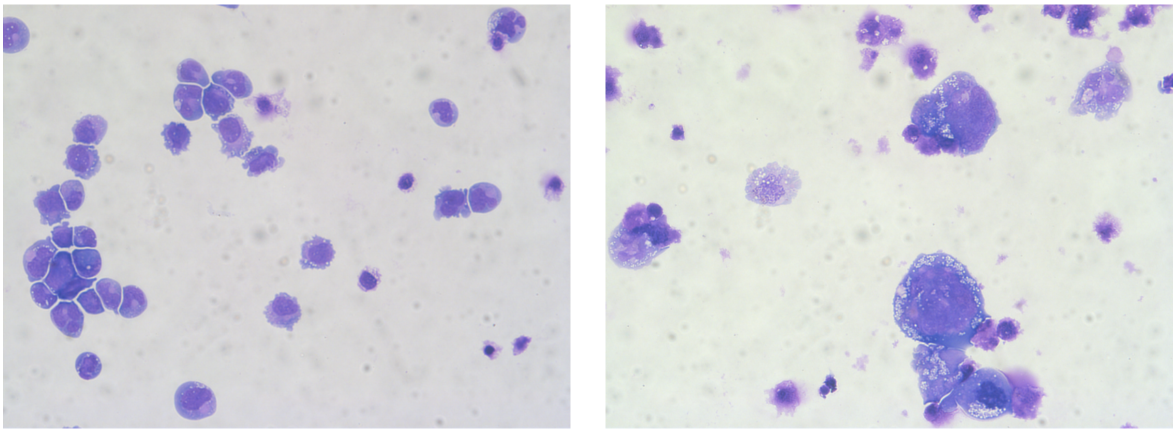

Figure S5. Cytospins of OCI-AML3 cells treated with (A) DMSO control or (B) OXS000675 10 μM stained with Modified Wright stain.

A B

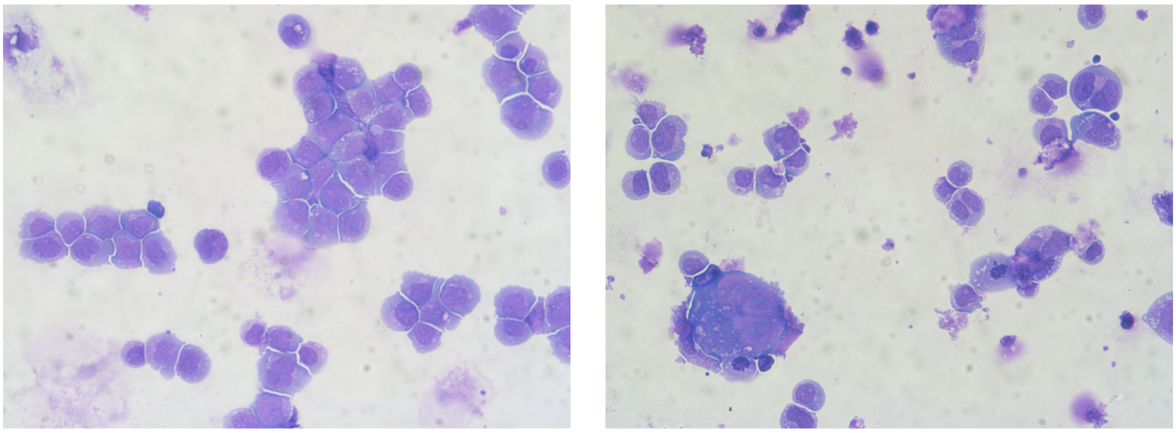

Figure S6. Cytospins of KG-1 cells treated with (A) DMSO control or (B) OXS000675 10 µM stained with Modified Wright stain.

A B

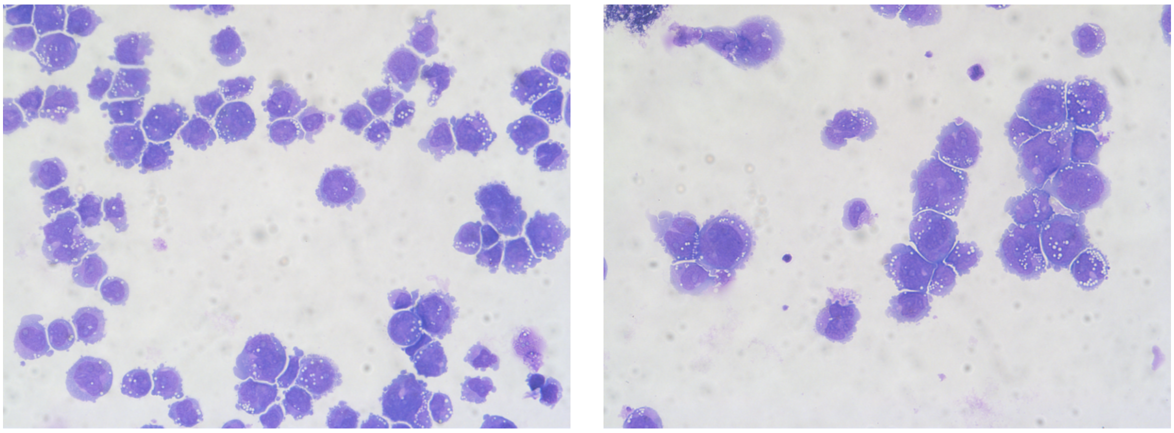

Figure S7. Cytospins of ME-1 cells treated with (A) DMSO control or (B) OXS000675 10 µM stained with Modified Wright stain.

A B

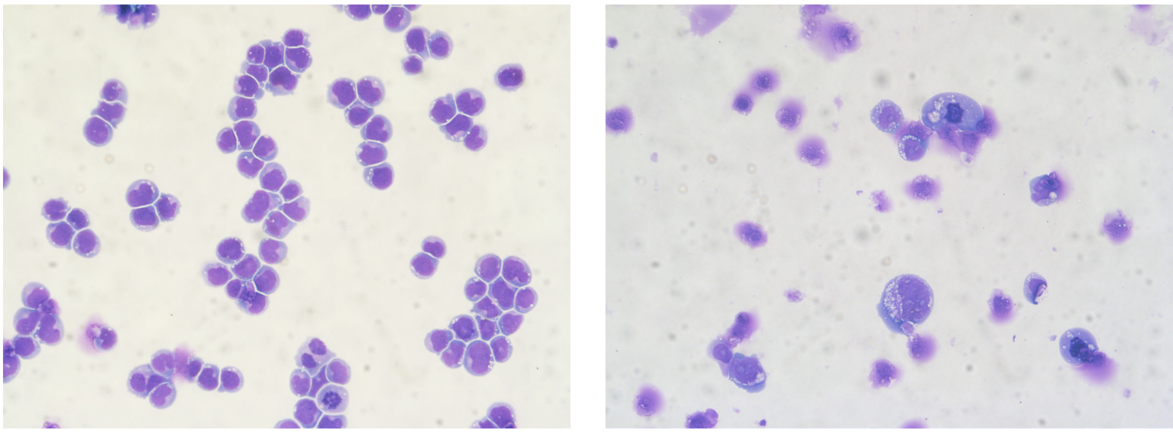

Figure S8. Cytospins of Kasumi-1 cells treated with (A) DMSO control or (B) OXS000675 10 µM stained with Modified Wright stain.

A B

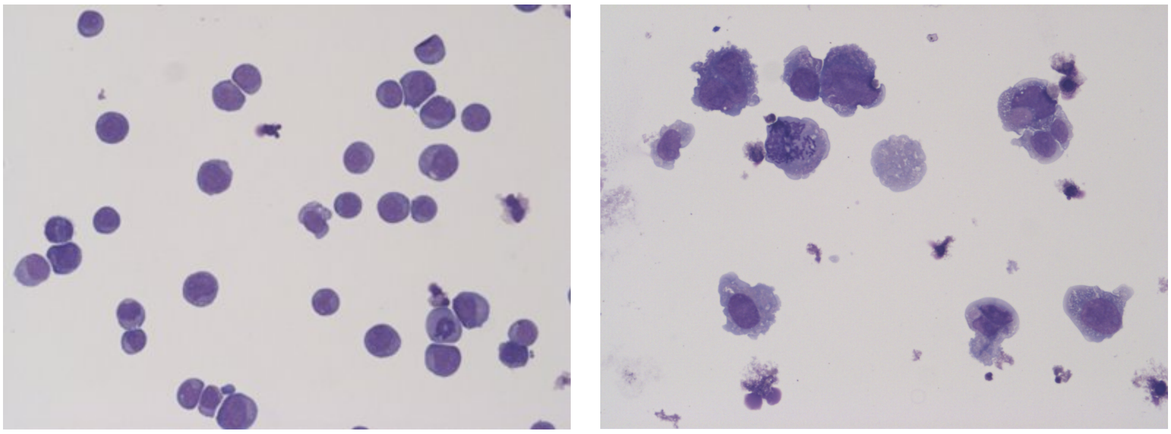

Figure S9. Cytospins of HL-60 cells treated with (A) DMSO control or (B) OXS007417 370 nM stained with Modified Wright stain.

A B

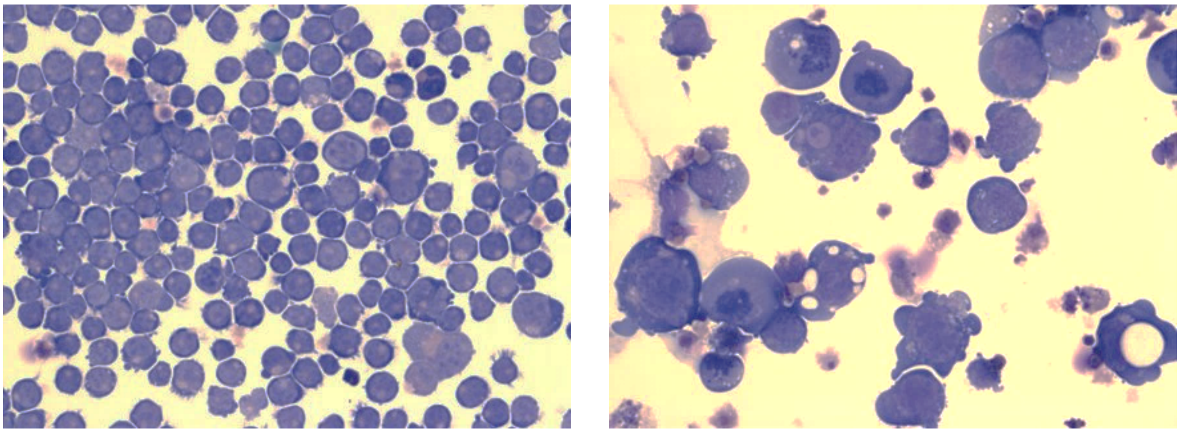

Figure S10. Cytospins of THP-1 cells treated with (A) DMSO control or (B) OXS007417 370 nM stained with Modified Wright stain.

A B

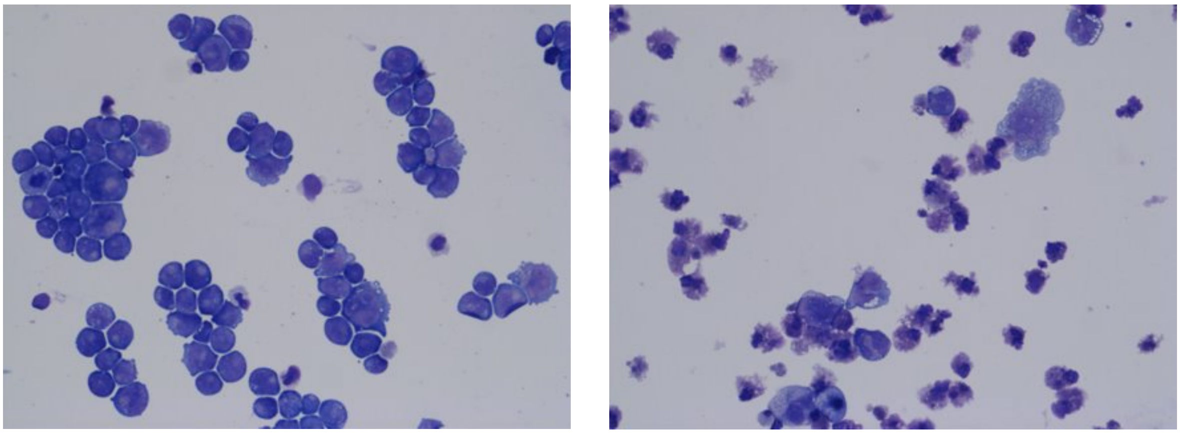

Figure S11. Cytospins of OCI-AML3 cells treated with (A) DMSO control or (B) OXS007417 370 nM stained with Modified Wright stain.

A B

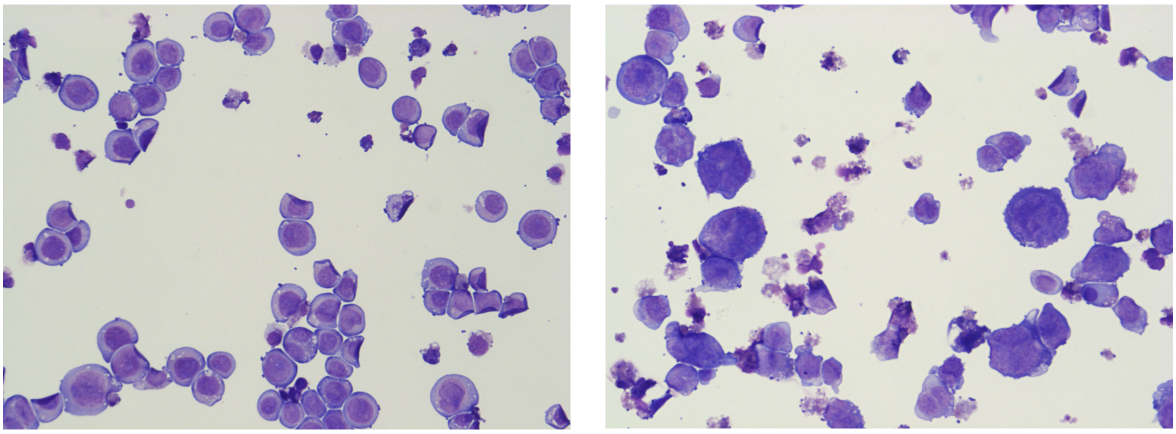

Figure S12. Cytospins of KG-1 cells treated with (A) DMSO control or (B) OXS007417 41 nM stained with Modified Wright stain.

A B

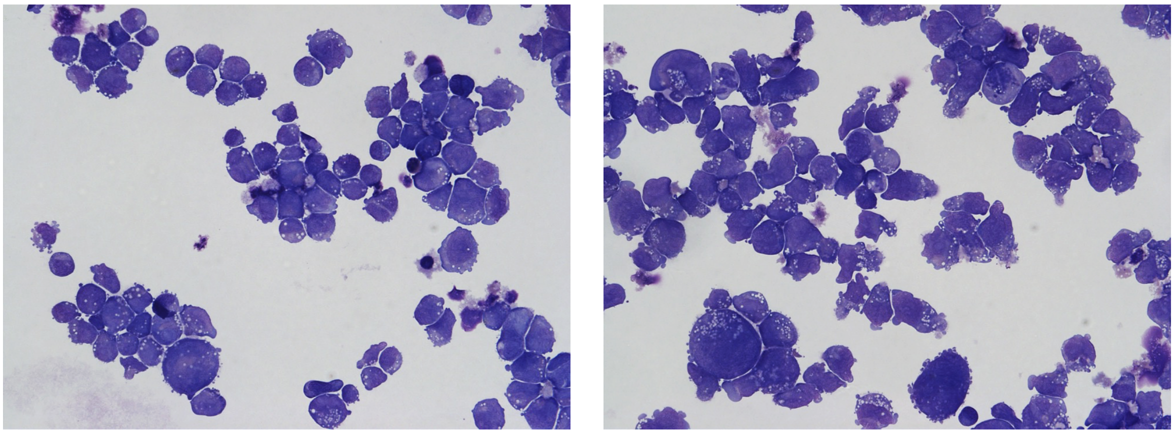

Figure S13. Cytospins of ME-1 cells treated with (A) DMSO control or (B) OXS007417 370 nM stained with Modified Wright stain.

A B

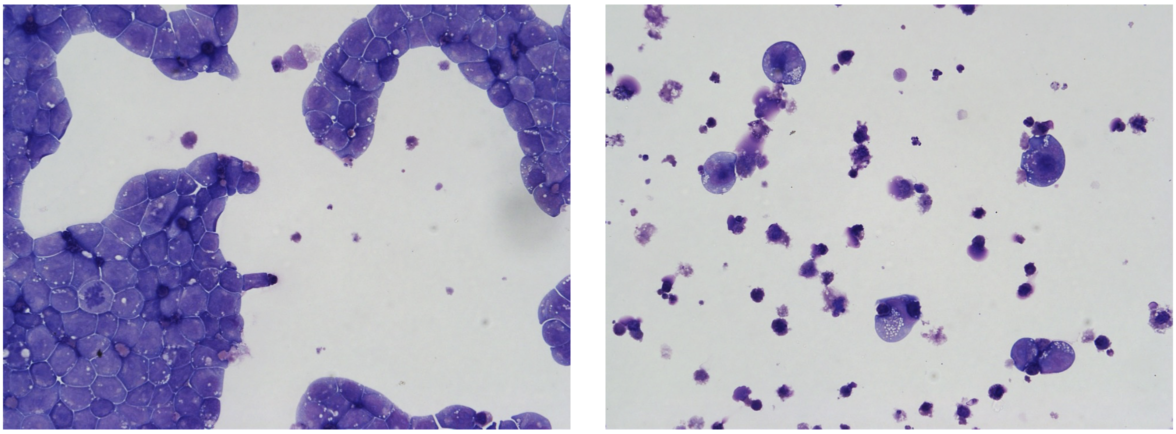

Figure S14. Cytospins of Kasumi-1 cells treated with (A) DMSO control or (B) OXS007417 370 nM stained with Modified Wright stain.

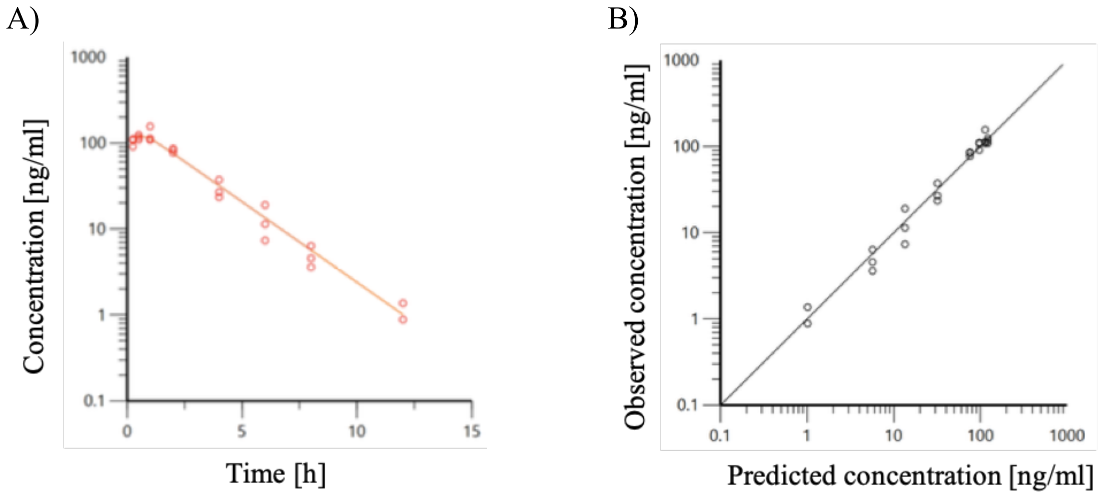


C)

| Parameter | Estimate | CV% |
|---|---|---|
| $K_a$ [$h^{-1}$] | 4.13 | 12.7 |
| V [mL] | 18700 | 5.32 |
| CL [mL/h] | 8060 | 4.37 |

Figure S15. PK model parameters and observed vs predicted values for OXS007417. (A) Predicted (line) versus observed (circles) blood concentration-time profile. (B) Observed versus predicted blood concentrations (solid line represents predicted=observed). (C) Fitted model parameters and their CV%.
