## Supplementary Information for "A phenotypic screen identifies a compound series that induces differentiation of acute myeloid leukemia cells *in vitro* and shows anti-tumour effects *in vivo*"

### SUPPORTING INFORMATION

#### Biological testing experimental procedures

##### Cell Culture

AML cell lines HL-60, THP-1, OCI-AML3, Kasumi-1, KG-1 and ME-1 were purchased from the American Type Culture Collection (ATCC). The cells were maintained in RPMI supplemented with 10% fetal bovine serum and 1% L-glutamine.

##### Flow Cytometry

PE mouse-anti-human-CD11b/Mac-1 (ICRF44, Cat 555388) and PE-mouse-IgG1-κ-isotype control (MOPC-21, Cat 555749) were purchased from BD Bioscience. Cells were suspended in 100 µL growth media (RPMI + 10% FBS + 1% L-glutamine) at a density of $2x10^4$ cells/well of a V-bottom 96 well plate (Corning) and grown for 96 hours in the presence of compound at the required concentration. After 96 hours, the plate was centrifuged, media was removed, and cells were resuspended in 50 µL of blocking buffer (IMDM, no phenol red + 10% FBS) containing 2.5 µL of either CD11b or isotype control antibody and incubated on ice in the dark for 20 min. Cells were washed twice in 200 µL of staining buffer (IMDM, no phenol red + 1% FBS) before being resuspended in 200 µL of staining buffer containing 1µg/ml of DAPI to help identify dead cells. Flow cytometry data was collected on an Attune NxT and analysed using the Attune NxT software (Life Technologies).

##### Cell counts and viability assessment

Solution 13 containing acridin orange and DAPI was purchased from ChemoMetec (Cat 910-3013). After the appropriate cell treatment, one volume of solution 13 was added into 19 volumes of the pre-mixed cell suspension, and analysed using the NucleoCounter® NC-300™ (ChemoMetec).

##### Morphology assessment with Wright staining

Cells were prepared in staining buffer (IMDM, no phenol red + 2% FBS) at a concentration of approximately $1x10^5$ cells/mL. Cytospins were made (1,000 rpm, 10 min), and the cells were allowed to air dry. Cells were stained with Modified Wright stain using a Hematek®. Stained cells were allowed to air dry and coverslips were affixed with DPX mount prior to microscopy.

**PK modelling**

A single compartment PK model with first order rate of absorption (Ka) parameterised by clearance (CL) and volume of distribution (V) was fit to the observed data with multiplicative weighting in Phoneix64®. Plots of observed vs predicted concentration and predicted and observed concentration vs time (Figure S15) reasonably capture the observed data.

***In Vivo* Leukemia Analysis: Subcutaneous Model**

Female NOD SCID mice aged 5-7 weeks of age were used for the HL-60 subcutaneous model. Cells ($5x10^6$, 1:1 in Matrigel) were implanted subcutaneously under the flank of each mouse. When tumours reached approximately 150 $mm^3$, mice were grouped randomly into treatment groups based on their bodyweight to ensure equal distribution. Mice were treated by indicated treatment regimes. Drug solutions were formulated immediately prior to dosing.

| Group | n | Treatment | Dose Level | Dosing Route | Dosing Regimen |
|---|---|---|---|---|---|
| 1 | 10 | Vehicle | -- | PO | BID |
| 2 | 5 | Cyclophosphamide | 150 mg/kg | IP | Q5D |
| 3 | 10 | OXS007417 | 10 mg/kg | PO | BID |
| 4 | 10 | OXS007417 | 10 mg/kg | PO | QD |
| 5 | 10 | OXS007417 | 3 mg/kg | PO | BID |
| 6 | 10 | OXS007417 | 30 mg/kg | PO | BID |

Cyclophosphamide was formulated in sterile saline. Dosing volume was 10 ml/kg for IP dosing.

OXS007417 was formulated in DMSO (final concentration of 5%) and PBS + 0.1% Tween 20. Dosing volume was 10 ml/kg for PO dosing.

The length and width of the tumours were measured 3 times per week using digital callipers. The study was terminated at the end of the 28 day treatment period.

All protocols used in this study were approved by the Axis Bioservices Animal Welfare and Ethical Review Committee, and all procedures are carried out under the guidelines of the Animal (Scientific Procedures) Act 1986.

### Chemistry synthesis experimental procedures

#### General information

All reactions involving moisture sensitive reagents were carried out under a nitrogen or an argon atmosphere. Anhydrous solvents were dried by passing over an activated alumina column, under an inert atmosphere, using a solvent purification system. All other solvents and reagents were used as supplied (analytical or HPLC grade) without prior purification. *Flash* column chromatography was performed on Kieselgel 60 silica gel (230-400 mesh particle size) on a glass column or on a Biotage SP4 automated flash column chromatography platform. NMR spectra were recorded on Bruker Advance spectrometers at 400 or 500 MHz in the deuterated solvent stated at room temperature. The field was locked by external referencing to the relevant deuteron resonance. Chemical shifts (δ) are reported in parts per million (ppm) and coupling constants (*J*) are quoted in Hz. Data are reported as follows: chemical shift, multiplicity (s = singlet, d = doublet, t = triplet, q = quartet, sext = sextuplet, hept = heptet, and m = multiplet), coupling constant and integration. Low-resolution mass spectra (*m/z*) were recorded on an Agilent 1260 Infinity II with Diode Array and Single Quadrupole Detectors in solutions of MeOH. A selected peak is reported in Daltons and its intensity given as percentage of the base peak. High resolution mass spectra (HRMS) were run on a Bruker microTOF (ESI and APCI) or on a Waters GCT (EI).

#### General synthetic route

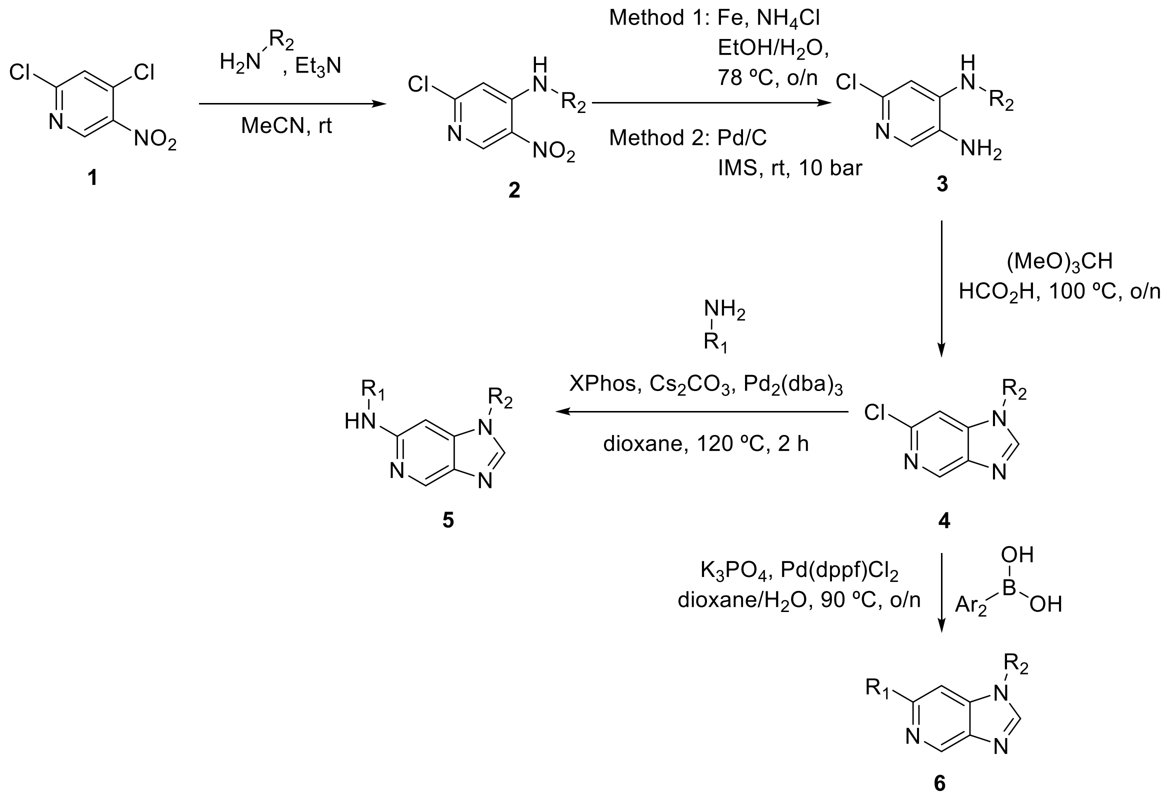

#### General procedure A: Nucleophilic aromatic substitution

To a solution of 2,4-dichloro-5-nitro-pyridine (1.0 equiv) in acetonitrile (*c* 0.5 M) were added the desired amine (1.0 equiv) and triethylamine (2.0 equiv). The mixture was stirred at room temperature until full conversion, and concentrated under reduced pressure. The resulting residue was redissolved in EtOAc, washed with 1x$H_2O$ and 1xbrine, dried over anhydrous $Na_2SO_4$ and concentrated to give the amine **2**, which was used in the following step without further purification.

#### General procedure B.1: Nitro reduction with Fe/$NH_4Cl$

To a suspension of nitro **2** (1.0 equiv) in ethanol/water (1:2, *c* 0.2) were added ammonium chloride (4 equiv) and iron powder (5 equiv). The reaction mixture was heated at reflux for 14 h, then cooled to room temperature, diluted with DCM and filtered through Celite. The filtrate was basified with saturated aqueous $NaHCO_3$, extracted with 3xDCM, washed with brine, dried over anhydrous $Na_2SO_4$ and concentrated to give amine **3**, which was used in the following step without further purification.

**General procedure B.2: Nitro reduction via hydrogenation**

A solution of nitro **2** (1.0 equiv) in IMS (*c* 0.01 M) was placed in an autoclave. Pd/C (10 wt.%, 0.05 equiv) wetted with water was added and the reaction was heated to 20 °C under 10 bar $H_2$ for 2 h. The mixture was then filtered through Celite® and evaporated to dryness. The crude residue was purified by *flash* column chromatography to give the amine.

**General procedure C: Imidazole formation**

Formic acid (1 equiv) was added to a solution of amine **3** (1.0 equiv) in triethylorthoformate (*c* 0.3 M), and the mixture was stirred at 100 °C overnight. The reaction mixture was concentrated, and to the resulting residue was added saturated aqueous $NaHCO_3$, extracted with 3xEtOAc, dried over anhydrous $Na_2SO_4$ and concentrated to give imidazopyridine **4**, which was used in the following step without further purification.

**General procedure D: Buchwald**

To a solution of **4** (1.0 equiv) in 1,4-dioxane (*c* 0.2 M) under argon were added the desired amine (1.0 equiv), caesium carbonate (2 equiv), X-Phos (0.2 equiv) and $Pd_2(dba)_3$ (0.05 equiv). The reaction mixture was heated at 120 °C for 2 h, then diluted with EtOAc, filtered through Celite and concentrated. The crude product was purified by *flash* column chromatography to give **5**.

**General procedure E.1: Suzuki reaction**

To a solution of **5** (1 equiv) in 1,4-dioxane/water (4:1, *c* 0.2 M) under argon were added potassium phosphate (3 equiv), the desired boronic acid (1.2 equiv) and $Pd(dppf)Cl_2$ (0.1 equiv). The mixture was heated at 100 °C overnight, then diluted with EtOAc, filtered through Celite and concentrated. The crude product was purified by *flash* column chromatography to give the desired product **6**.

**General procedure E.2: Suzuki reaction**

To a solution of chloride **5** (1 equiv) in degassed DME (*c* 0.5 M) under argon were added potassium carbonate (2 equiv), the required boronic acid (1.2 equiv) and Pd(dppf)$Cl_2$ (0.1 equiv). The mixture was heated to 80 °C overnight, then diluted with EtOAc, filtered through Celite and concentrated. The crude product was purified by *flash* column chromatography to give the desired product **6**.

**2-Chloro-*N*-[(3,4-dimethoxyphenyl)methyl]-5-nitro-pyridin-4-amine (7)**

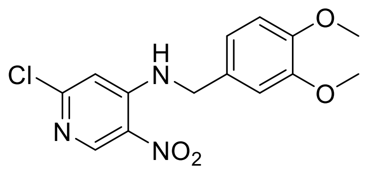

General procedure A; yield 95% (800 mg, 2.47 mmol); yellow solid. $^1$H NMR (500 MHz, $CDCl_3$) δ 9.01 (s, 1H), 8.40 (t, *J* = 5.3 Hz, 1H), 6.87 (d, *J* = 1.1 Hz, 2H), 6.81 (d, *J* = 1.2 Hz, 1H), 6.77 (s, 1H), 4.44 (d, *J* = 5.3 Hz, 2H), 3.88 (s, 3H), 3.87 (s, 3H); $^{13}$C NMR (125 MHz, $CDCl_3$) δ 156.6, 149.7, 149.6, 149.2, 149.2, 129.6, 127.6, 119.8, 111.7, 110.5, 107.2, 56.1, 56.1, 47.1; *m/z* ($ESI^+$) 324.0 ($[M+H]^+$, 53%); HRMS ($ESI^+$) $C_{14}H_{15}N_3O_4Cl^+$ ($[M+H]^+$) requires 324.0746; found 324.0747.

**6-Chloro-*N*$^4$-(3,4-dimethoxybenzyl)pyridine-3,4-diamine (8)**

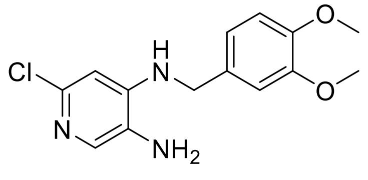

General procedure B.1 from **7** (603 mg, 1.86 mmol); yield quant. (550 mg, 1.86 mmol); dark purple solid. $^1$H NMR (400 MHz, $CDCl_3$) δ 7.61 (s, 1H), 6.90 – 6.79 (m, 3H), 6.46 (s, 1H), 4.69 (t, *J* =

5.3 Hz, 1H), 4.24 (d, $J$ = 5.2 Hz, 2H), 3.86 (s, 3H), 3.85 (s, 3H); $^{13}$C NMR (100 MHz, $CDCl_3$) δ 149.3, 148.7, 147.2, 144.7, 136.6, 129.8, 128.1, 120.1, 111.4, 111.0, 104.3, 56.0, 56.0, 47.3; *m/z* (ESI$^+$) 294.1 ([M+H]$^+$, 100%); HRMS (ESI$^+$) $C_{14}H_{17}N_3O_2Cl^+$ ([M+H]$^+$) requires 294.10038; found 294.10036.

**6-Chloro-1-(3,4-dimethoxybenzyl)-1*H*-imidazo[4,5-*c*]pyridine (9)**

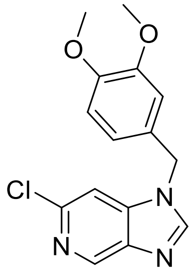

General procedure C from **8** (129 mg, 0.439 mmol); yield quant. (133 mg, 0.438 mmol); dark orange oil. $^1$H NMR (400 MHz, $CDCl_3$) δ 8.85 (t, $J$ = 1.0 Hz, 1H), 7.97 (s, 1H), 7.26 (s, 1H), 6.85 (d, $J$ = 8.2 Hz, 1H), 6.75 (dd, $J$ = 8.2, 2.1 Hz, 1H), 6.67 (d, $J$ = 2.1 Hz, 1H), 5.24 (s, 2H), 3.87 (s, 3H), 3.81 (s, 5H); $^{13}$C NMR (100 MHz, $CDCl_3$) δ 149.8, 149.6, 145.7, 144.0, 142.3, 141.0, 140.7, 126.2, 120.2, 111.6, 110.4, 105.6, 56.1, 56.1, 49.3; *m/z* (ESI$^+$) 304.1 ([M+H]$^+$, 33%); HRMS (ESI$^+$) $C_{15}H_{15}N_3O_2Cl^+$ ([M+H]$^+$) requires 304.08473; found 304.08464.

***N*-(3-((1-(3,4-Dimethoxybenzyl)-1*H*-imidazo[4,5-*c*]pyridin-6-yl)amino)phenyl)acetamide (OXS000675)**

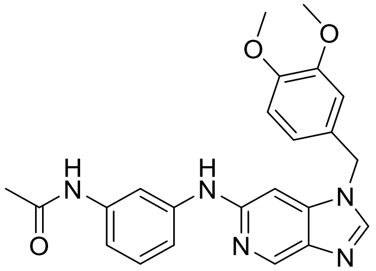

General procedure D from **9** (118 mg, 0.389 mmol); yield 36% (59 mg, 0.14 mmol); brown solid. $^{1}$H NMR (400 MHz, DMSO-$d_6$) δ 9.82 (s, 1H), 8.80 (s, 1H), 8.56 (s, 1H), 8.30 (s, 1H), 7.77 (t, *J* = 2.1 Hz, 1H), 7.23 (d, *J* = 8.1 Hz, 1H), 7.10 (t, *J* = 8.0 Hz, 1H), 7.07 – 6.98 (m, 2H), 6.94 – 6.86 (m, 2H), 6.77 (dd, *J* = 8.2, 2.0 Hz, 1H), 5.30 (s, 2H), 3.71 (s, 3H), 3.71 (s, 3H), 2.03 (s, 3H); $^{13}$C NMR (100 MHz, DMSO-$d_6$) δ 168.2, 150.9, 148.8, 148.4, 145.1, 142.8, 140.8, 139.6, 138.9, 136.1, 128.6, 128.6, 119.8, 112.4, 111.9, 111.5, 111.0, 108.4, 89.8, 55.5, 55.5, 47.5, 24.0; *m/z* (ESI$^+$) 418.1 ([M+H]$^+$, 45%); HRMS (ESI$^+$) $C_{23}H_{24}N_5O_3^+$ ([M+H]$^+$) requires 418.18737; found 418.18692.

**2-Chloro-*N*-(3,4-dimethoxyphenyl)-5-nitropyridin-4-amine (10)**

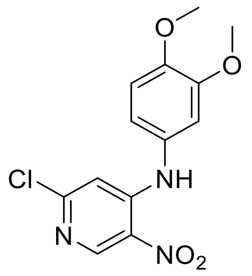

General procedure A; yield 97% (781 mg, 2.52 mmol); purple solid. $^{1}$H NMR (400 MHz, $CDCl_3$) δ 9.49 (s, 1H), 9.00 (s, 1H), 6.93 (d, *J* = 8.5 Hz, 1H), 6.83 (ddd, *J* = 8.4, 2.4, 0.7 Hz, 1H), 6.77 (s, 1H), 6.74 (d, *J* = 2.4 Hz, 1H), 3.90 (s, 3H), 3.86 (s, 3H); $^{13}$C NMR (100 MHz, $CDCl_3$) δ 156.4, 150.2, 149.6, 149.2, 148.8, 129.4, 128.6, 118.4, 111.9, 109.7, 108.3, 56.2, 56.2; *m/z* (ESI$^+$) 310.0 ([M+H]$^+$, 90%); HRMS (ESI$^+$) $C_{13}H_{13}N_3O_4Cl^+$ ([M+H]$^+$) requires 310.05891; found 310.05881.

**6-Chloro-N$^4$-(3,4-dimethoxyphenyl)pyridine-3,4-diamine (11)**

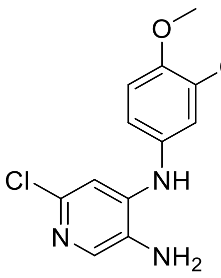

General procedure B.1 from **10** (686 mg, 2.22 mmol); yield 99% (614 mg, 2.20 mmol); purple solid. $^{1}$H NMR (400 MHz, $CDCl_3$) δ 7.71 (s, 1H), 6.86 (d, *J* = 8.4 Hz, 1H), 6.77 – 6.61 (m, 3H), 6.09 (s, 1H), 3.88 (s, 3H), 3.83 (s, 3H); $^{13}$C NMR (100 MHz, $CDCl_3$) δ 149.9, 146.7, 145.6, 144.4, 137.6, 132.5, 128.9, 115.4, 112.1, 107.7, 106.3, 56.3, 56.1; *m/z* ($ESI^+$) 280.1 ($[M+H]^+$, 100%); HRMS ($ESI^+$) $C_{13}H_{15}N_3O_2Cl^+$ ($[M+H]^+$) requires 280.08473; found 280.08443.

**6-Chloro-1-(3,4-dimethoxyphenyl)-1*H*-imidazo[4,5-*c*]pyridine (12)**

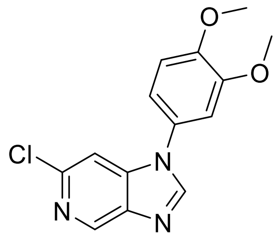

General procedure C from **11** (576 mg, 2.06 mmol); yield 94% (558 mg, 1.93 mmol); white solid. $^{1}$H NMR (400 MHz, $CDCl_3$) δ 8.89 (d, *J* = 0.9 Hz, 1H), 8.10 (s, 1H), 7.39 (d, *J* = 1.0 Hz, 1H), 7.03 (d, *J* = 8.5 Hz, 1H), 7.00 (dd, *J* = 8.4, 2.2 Hz, 1H), 6.91 (d, *J* = 2.2 Hz, 1H), 3.96 (s, 3H), 3.93 (s, 3H); $^{13}$C NMR (100 MHz, $CDCl_3$) δ 150.3, 149.8, 145.2, 144.6, 142.5, 141.2, 140.5, 127.6, 116.9, 111.9, 108.0, 105.8, 56.4, 56.4; *m/z* ($ESI^+$) 290.1 ($[M+H]^+$, 95%); HRMS ($ESI^+$) $C_{14}H_{13}N_3O_2Cl^+$ ($[M+H]^+$) requires 290.06908; found 290.06894.

***N*-(3-((1-(3,4-Dimethoxyphenyl)-1*H*-imidazo[4,5-*c*]pyridin-6-yl)amino)phenyl)acetamide (OXS007014)**

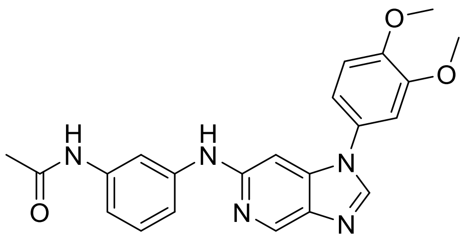

General procedure D from **12** (50 mg, 0.17 mmol); yield 85% (59 mg, 0.15 mmol); brown solid. $^{1}$H NMR (500 MHz, DMSO-$d_6$) δ 9.83 (s, 1H), 8.87 (s, 1H), 8.67 (d, $J$ = 0.9 Hz, 1H), 8.41 (s, 1H), 7.85 (t, $J$ = 2.1 Hz, 1H), 7.38 – 7.31 (m, 1H), 7.26 (d, $J$ = 2.1 Hz, 1H), 7.18 – 7.16 (m, 2H), 7.11 (d, $J$ = 8.0 Hz, 1H), 7.06 (dt, $J$ = 8.3, 1.3 Hz, 1H), 7.01 (d, $J$ = 1.0 Hz, 1H), 3.86 (s, 3H), 3.84 (s, 3H), 2.03 (s, 3H); $^{13}$C NMR (125 MHz, DMSO-$d_6$) δ 168.1, 151.9, 149.6, 148.3, 144.1, 142.7, 140.6, 139.6, 139.2, 135.8, 128.7, 128.4, 115.6, 112.5, 112.3, 111.0, 108.2, 108.0, 89.6, 55.9, 55.9, 24.1; *m/z* (ESI$^+$) 404.1 ([M+H]$^+$, 100%); HRMS (ESI$^+$) $C_{22}H_{22}N_5O_3^+$ ([M+H]$^+$) requires 404.17172; found 404.17161.

***N*-[3-[[1-(3,4-Dimethoxyphenyl)imidazo[4,5-c]pyridin-6-yl]amino]phenyl]methanesulfonamide (OXS007746)**

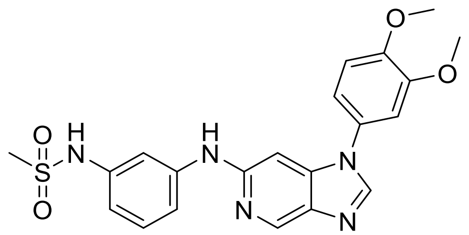

General procedure D from **12** (81 mg, 0.27 mmol); yield 37% (46 mg, 0.10 mmol); brown solid.$^{1}$H NMR (500 MHz, $CDCl_3$) δ 8.83 (s, 1H), 8.01 (s, 1H), 7.41 (t, $J$ = 2.2 Hz, 1H), 7.26 (d, $J$ = 8.0 Hz, 1H), 7.10 – 6.99 (m, 5H), 6.96 (d, $J$ = 2.0 Hz, 1H), 6.88 (s, 1H), 6.79 (dd, $J$ = 7.9, 1.3 Hz, 1H), 3.98 (s, 3H), 3.95 (s, 3H), 3.01 (s, 3H); $^{13}$C NMR (126 MHz, $CDCl_3$) δ 151.0, 150.1, 149.3, 143.6, 142.9, 141.6, 141.4, 137.9, 136.7, 130.5, 128.3, 116.7, 115.3, 113.4, 111.9, 109.9, 107.9, 89.2, 56.4, 56.3, 39.4; *m/z*(ESI$^+$) 440.1 ([M+H]$^+$, 100%); HRMS (ESI$^+$) $C_{21}H_{22}N_5O_4S^+$ ([M+H]$^+$) requires 440.1387; found 440.1386.

**$N$-(3-((1-(3,4-Dimethoxyphenyl)-1$H$-imidazo[4,5-$c$]pyridin-6-yl)oxy)phenyl)acetamide (OXS007266)**

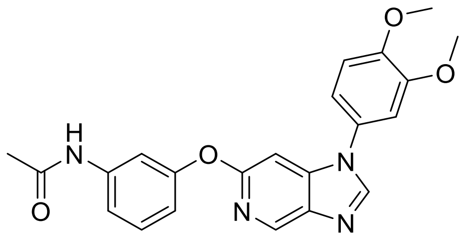

To a solution of **12** (100 mg, 0.345 mmol) in degassed 1,4-dioxane (2 mL) under argon were added 3-acetamidophenol (52 mg, 0.34 mmol) caesium carbonate (225 mg, 0.690 mmol), XantPhos (40 mg, 0.069 mmol) and $Pd(OAc)_2$ (3.9 mg, 0.017 mmol). The reaction mixture was heated at 120 °C for 3 days. The reaction mixture was allowed to cool to room temperature, concentrated onto Celite® and then purified by silica gel chromatography (0 – 10% MeOH in DCM, then 0 – 10% MeOH in EtOAc) to give **OXS007266** (28 mg, 0.069 mmol, 20%) as a pale brown solid.$^{1}$H NMR (500 MHz, $CDCl_3$) δ 8.78 (s, 1H), 8.06 (s, 1H), 7.40 (s, 1H), 7.31 – 7.28 (m, 2H), 7.20 (d, $J$ = 8.3 Hz, 1H), 7.02 (s, 2H), 6.96 (d, $J$ = 16.1 Hz, 2H), 6.86 (d, $J$ = 8.1 Hz, 1H), 3.97 (s, 3H), 3.93 (s, 3H), 2.13 (s, 3H); $^{13}$C NMR (126 MHz, $CDCl_3$) δ 168.3, 159.6, 156.2, 150.3, 149.6, 144.7, 142.3, 140.6, 139.3, 138.7, 130.1, 128.3, 116.6, 116.1, 115.4, 112.0, 107.9, 93.0, 56.4, 24.4; $m/z$ (ESI$^+$) 405.1 ([M+H]$^+$, 100%);HRMS (ESI$^+$) $C_{22}H_{21}O_4N_4$ ([M+H]$^+$) requires 405.1557; found 405.1555.

**$N$-(3-(1-(3,4-dimethoxyphenyl)-1$H$-imidazo[4,5-$c$]pyridin-6-yl)phenyl)acetamide (OXS007034)**

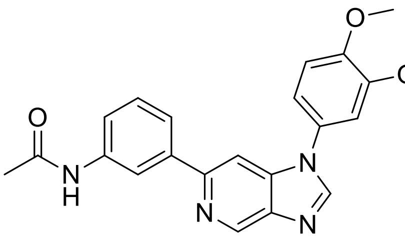

General procedure E.2 from **12** (39 mg, 0.13 mmol); yield 88% (46 mg, 0.12 mmol); brown solid. $^1$H NMR (500 MHz, $CDCl_3$) δ 9.23 (s, 1H), 8.15 (s, 1H), 8.11 (t, *J* = 2.0 Hz, 1H), 7.80 (d, *J* = 1.1 Hz, 1H), 7.77 – 7.63 (m, 2H), 7.54 (s, 1H), 7.42 (t, *J* = 7.9 Hz, 1H), 7.34 – 7.20 (m, 1H), 7.10-7.08 (m, 2H), 7.01 (d, *J* = 2.2 Hz, 1H), 4.01 (s, 3H), 3.97 (s, 3H), 2.20 (s, 3H); $^{13}$C NMR (126 MHz, CDCl3) δ 168.4, 151.2, 150.2, 149.6, 144.5, 142.9, 140.7, 140.3, 140.2, 138.5, 129.5, 128.1, 122.8, 120.0, 118.5, 116.9, 111.9, 108.2, 102.5, 56.4, 56.3, 24.7; *m/z* ($ESI^+$) 389.1 ($[M+H]^+$, 100%); HRMS ($ESI^+$) $C_{22}H_{21}N_4O_3^+$ ($[M+H]^+$) requires 389.16082; found 389.16136.

**1-(3,4-Dimethoxyphenyl)-6-phenyl-1*H*-imidazo[4,5-*c*]pyridine (OXS007136)**

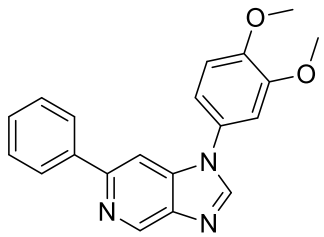

General procedure E.2 from **12** (100 mg, 0.345mmol); yield 75% (77 mg, 0.23 mmol); beige solid. $^1$H NMR (500 MHz, $CDCl_3$) δ 9.25 (d, *J* = 1.1 Hz, 1H), 8.12 (s, 1H), 8.03 – 7.98 (m, 2H), 7.77 (d, *J* = 1.0 Hz, 1 H),7.46 (dd, *J* = 8.3, 6.9 Hz, 2H), 7.41 – 7.35 (m, 1H), 7.12 – 7.04 (m, 2H), 6.99 (d, *J* = 2.1 Hz, 1H), 3.99 (s, 3H), 3.95 (s, 3H); $^{13}$C NMR (126 MHz, $CDCl_3$) δ 152.1, 150.3, 149.7, 144.4, 143.1, 140.3, 140.1, 128.9, 128.6, 128.3, 127.2, 117.0, 112.0, 108.3, 102.4, 56.5; *m/z* ($ESI^+$) 332.1 ($[M+H]^+$, 100%);HRMS ($ESI^+$) $C_{21}H_{18}O_2N_3$ ($[M+H]^+$) requires 332.13935; found 332.13922.

**1-(3,4-Dimethoxyphenyl)-6-(*o*-tolyl)-1*H*-imidazo[4,5-*c*]pyridine (OXS007142)**

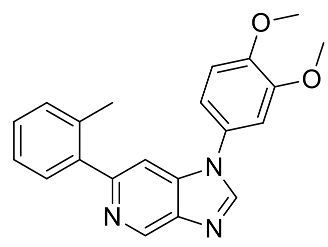

General procedure E.2 from **12** (90 mg, 0.31 mmol); yield 67% (72 mg, 0.21mmol); beige solid. $^1$H NMR (500 MHz, $CDCl_3$) δ 9.25 (d, *J* = 1.1 Hz, 1H), 8.15 (s, 1H), 7.49 (d, *J* = 1.1 Hz, 1H), 7.40 (dt, *J* = 7.2, 1.2 Hz, 1H), 7.29 (d, *J* = 1.1 Hz, 1H), 7.07 – 7.00 (m, 2H), 6.98 (d, *J* = 2.4 Hz, 1H), 3.96 (s, 3H), 3.93 (s, 3H), 2.37 (s, 3H); $^{13}$C NMR (126 MHz, $CDCl_3$) δ 154.1, 150.2, 149.6, 144.2, 142.6, 141.0, 139.6, 136.1, 130.9, 130.1, 128.2, 126.0, 116.8, 111.9, 108.1, 105.9, 56.4, 20.6; *m/z* ($ESI^+$) 346.2 ($[M+H]^+$, 100%); HRMS ($ESI^+$) $C_{21}H_{20}O_2N_3$ ($[M+H]^+$) requires 346.15500; found 346.15555.

**1-(3,4-Dimethoxyphenyl)-6-(2-methoxyphenyl)-1*H*-imidazo[4,5-*c*]pyridine (OXS006988)**

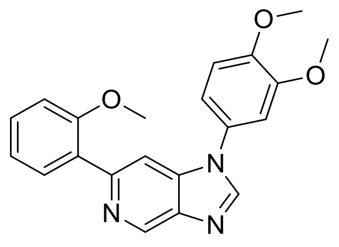

General procedure E.1 from **12** (50 mg, 0.17 mmol); yield 80% (50 mg, 0.14 mmol); brown solid. $^1$H NMR (400 MHz, $CDCl_3$) δ 9.25 (d, *J* = 1.0 Hz, 1H), 8.13 (s, 1H), 7.92 (d, *J* = 1.1 Hz, 1H), 7.82 (dd, *J* = 7.6, 1.8 Hz, 1H), 7.35 (ddd, *J* = 8.3, 7.4, 1.8 Hz, 1H), 7.09 (td, *J* = 7.5, 1.2 Hz, 1H), 7.06 – 6.98 (m, 4H), 3.97 (s, 3H), 3.94 (s, 3H), 3.82 (s, 3H); $^{13}$C NMR (100 MHz, $CDCl_3$) δ 156.8, 150.1, 149.8, 149.4, 144.2, 142.9, 140.0, 139.3, 131.7, 129.7, 129.6, 128.6, 121.2, 116.7, 111.9, 111. 6, 108.0, 106.9, 56.4, 56.4 55.9; *m/z* ($ESI^+$) 362.1 ($[M+H]^+$, 100%); HRMS ($ESI^+$) $C_{21}H_{20}N_3O_3^+$ ($[M+H]^+$) requires 362.14992; found 362.14938.

**1-(3,4-Dimethoxyphenyl)-6-(2-(trifluoromethoxy)phenyl)-1*H*-imidazo[4,5-*c*]pyridine (OXS007316)**

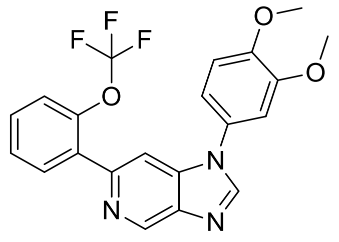

General procedure E.2 from **12** (90 mg, 0.31 mmol); yield 32% (41 mg, 0.099 mmol); pale brown foam. $^{1}$H NMR (500 MHz, $CDCl_3$) δ 9.28 (d, *J* = 1.1 Hz, 1H), 8.17 (s, 1H), 7.99 – 7.87 (m, 1H), 7.82 (d, *J* = 1.0 Hz, 1H), 7.42 – 7.37 (m, 1H), 7.37 – 7.24 (m, 2H), 7.04 (t, *J* = 1.5 Hz, 2H), 7.00 (d, *J* = 1.9 Hz, 1H), 3.97 (s, 3H), 3.94 (s, 3H); $^{13}$C NMR (126 MHz, $CDCl_3$) δ 150.3, 149.6, 147.9, 146.3, 144.7, 143.4, 140.4, 139.4, 134.3, 132.4, 129.6, 128.2, 127.5, 121.5, 116.7, 111.9, 107.9, 106.8, 56.4, 56.3; *m/z* ($ESI^+$) 416.1 ($[M+H]^+$, 100%); HRMS ($ESI^+$) $C_{21}H_{17}O_3N_3F_3$ ($[M+H]^+$) requires 416.1217; found 416.1207.

**1-(3,4-Dimethoxyphenyl)-6-(4-fluoro-2-methylphenyl)-1*H*-imidazo[4,5-*c*]pyridine(OXS007287)**

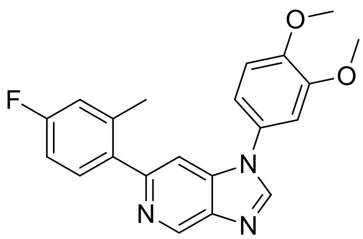

General procedure E.2 from **12** (90 mg, 0.31 mmol); yield 81% (92 mg, 0.25 mmol); beige solid. $^{1}$H NMR (500 MHz, $CDCl_3$) δ 9.32 – 9.16 (m, 1H), 8.16 (s, 1H), 7.45 – 7.42 (m, 1H), 7.37 (dd, *J* = 8.5, 5.9 Hz, 1H), 7.08 – 7.01 (m, 2H), 7.01 – 6.90 (m, 3H), 3.97 (s, 3H), 3.93 (s, 3H), 2.36 (s, 3H); $^{13}$C NMR (126 MHz, $CDCl_3$) δ 162.6 (d, *J* = 246 Hz), 153.2, 150.3, 149.7, 144.4, 142.7,

139.8, 138.8 (d, $J$ = 8 Hz), 131.7 (d, $J$ = 8 Hz), 128.3, 117.4 (d, $J$ = 21 Hz), 116.9, 112.8 (d, $J$ = 21 Hz), 112.0, 108.2, 105.9, 56.4 (d, $J$ = 4 Hz), 29.9, 20.7; *m/z* ($ESI^+$) 364.1 ($[M+H]^+$, 100%); HRMS ($ESI^+$) $C_{21}H_{19}O_2N_3F$($[M+H]^+$) requires 364.1456; found 364.1451.

**1-(3,4-Dimethoxyphenyl)-6-(4-fluoro-2-methoxyphenyl)-1*H*-imidazo[4,5-*c*]pyridine (OXS007184)**

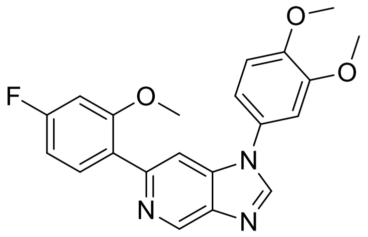

General procedure E.2 from **12** (90 mg, 0.31 mmol); yield 49% (58 mg, 0.15mmol); pale brown solid. $^1$H NMR (500 MHz, $CDCl_3$) δ 9.24 (d, $J$ = 1.1 Hz, 1H), 8.13 (s, 1H), 7.87 (d, $J$ = 1.1 Hz, 1H), 7.82 (dd, $J$ = 8.6, 7.0 Hz, 1H), 7.11 – 7.02 (m, 2H), 7.01 (d, $J$ = 2.2 Hz, 1H), 6.79 (td, $J$ = 8.3, 2.4 Hz, 1H), 6.72 (dd, $J$ = 10.9, 2.4 Hz, 1H), 3.98 (s, 3H), 3.94 (s, 3H), 3.81 (s, 3H); $^{13}$C NMR (126 MHz, $CDCl_3$) δ 164.7, 162.7, 157.9, 150.2, 149.5, 149.0, 144.3, 142.9, 140.0, 139.4, 132.8, 128.5, 125.7, 116.7, 111.9, 108.1, 107.7 (d, $J$= 21 Hz), 106.7, 99.6 (d, $J$= 26 Hz), 56.4, 56.1; *m/z* ($ESI^+$) 380.1($[M+H]^+$, 100%); HRMS ($ESI^+$) $C_{21}H_{19}O_3N_3F$($[M+H]^+$) requires 380.1405; found 380.1404.

**1-(3,4-Dimethoxyphenyl)-6-(5-fluoro-2-methylphenyl)-1*H*-imidazo[4,5-*c*]pyridine (OXS007317)**

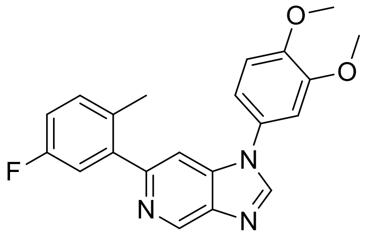

General procedure E.2 from **12** (90 mg, 0.31 mmol); yield 37% (37 mg, 0.10 mmol); beige solid. $^1$H NMR (500 MHz, $CDCl_3$) δ 9.24 (d, *J* = 1.1 Hz, 1H), 8.17 (s, 1H), 7.47 (d, *J* = 1.1 Hz, 1H), 7.23 – 7.18 (m, 1H), 7.14 (dd, *J* = 9.5, 2.8 Hz, 1H), 7.06 – 7.01 (m, 2H), 6.98 (dd, *J* = 8.9, 2.5 Hz, 2H), 3.97 (s, 3H), 3.93 (s, 3H), 2.33 (s, 3H); $^{13}$C NMR (126 MHz, $CDCl_3$) δ 161.1 (d, *J* = 244 Hz), 152.9 (d, *J* = 2 Hz), 150.3, 149.7, 144.5, 142.8, 142.5 (d, *J* = 7 Hz), 140.0, 139.7, 132.2 (d, *J* = 8 Hz), 131.7 (d, *J* = 3 Hz), 128.2, 116.9, 116.8 (d, *J* = 22 Hz), 114.8 (d, *J* = 21 Hz), 112.0, 108.2, 105.9, 56.4 (d, *J* = 4 Hz), 19.8; *m/z* (ESI$^+$) 364.1 ([M+H]$^+$, 100%); HRMS (ESI$^+$) $C_{21}H_{19}O_2N_3F$ ([M+H]$^+$) requires 364.1456; found 364.1451.

**1-(3,4-Dimethoxyphenyl)-6-(5-fluoro-2-methoxyphenyl)-1*H*-imidazo[4,5-*c*]pyridine (OXS007268)**

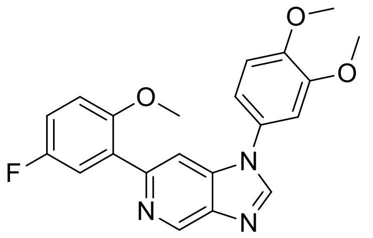

General procedure E from **12** (90 mg, 0.31 mmol); yield 75% (89 mg, 0.24 mmol); beige solid. $^1$H NMR (500 MHz, $CDCl_3$) δ 9.25 (d, *J* = 1.0 Hz, 1H), 8.15 (s, 1H), 8.00 (d, *J* = 1.1 Hz, 1H), 7.65 (dd, *J* = 9.5, 3.2 Hz, 1H), 7.13 – 6.99 (m, 4H), 6.93 (dd, *J* = 9.0, 4.4 Hz, 1H), 3.98 (s, 3H), 3.95 (s, 3H), 3.80 (s, 3H); $^{13}$C NMR (126 MHz, $CDCl_3$) δ 157.5 (d, *J* = 239 Hz), 153.1 (d, *J* = 2 Hz), 150.2, 149.5, 148.5, 144.5, 143.0, 140.2, 139.4, 128.5, 118.2 (d, *J* = 24 Hz), 116.7, 115.6 (d, *J* = 23 Hz), 112.9 (d, *J* = 8 Hz), 111.9, 108.1, 107.0, 56.4, 29.9; *m/z* (ESI$^+$) 380.1 ([M+H]$^+$, 100%); HRMS (ESI$^+$) $C_{21}H_{19}O_3N_3F$([M+H]$^+$) requires 380.1405; found 380.1401.

**1-(3,4-Dimethoxyphenyl)-6-(4-fluorophenyl)-1*H*-imidazo[4,5-*c*]pyridine (OXS007823)**

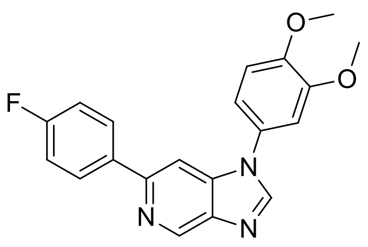

General procedure E.2 from **12** (100 mg, 0.346mmol); yield 75% (91 mg, 0.26 mmol); white solid. $^{1}$H NMR (500 MHz, $CDCl_3$) δ 9.25 (d, *J* = 0.8 Hz, 1H), 8.15 (s, 1H), 8.01 – 7.98 (m, 2H), 7.73 (d, *J* = 0.8 Hz, 1H), 7.18 – 7.14 (m, 2H), 7.10 – 7.08 (m, 2H), 7.00 (d, *J* = 1.43 Hz, 1H), 4.01 (s, 3H), 3.97 (s, 3H); $^{13}$C NMR (126 MHz, $CDCl_3$) δ 163.2 (d, *J* = 248.2 Hz), 150.9, 150.2, 149.6, 145.1, 143.4, 143.0, 142.5, 140.3, 140.1, 136.1 (d, *J* = 3.0 Hz), 128.8 (d, *J* = 8.7 Hz), 128.1, 117.0, 116.8, 115.6 (d, *J* = 21.5 Hz), 111.9, 108.3, 56.4, 56.3; *m/z* ($ESI^+$) 350.1 ($[M+H]^+$, 100%); HRMS ($ESI^+$) $C_{20}H_{17}N_3O_2F^+$ ($[M+H]^+$) requires 350.12993; found 350.12991.

**4-(1-(3,4-Dimethoxyphenyl)-1*H*-imidazo[4,5-*c*]pyridin-6-yl)morpholine (OXS007267)**

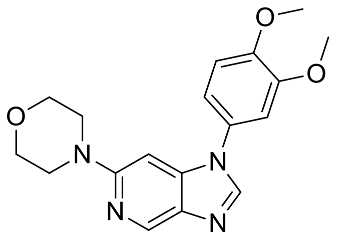

To a solution of **12** (100 mg, 0.346 mmol) in degassed 1,4-dioxane (2 mL) under argon were added morpholine (30 µL, 0.34 mmol) caesium carbonate (225 mg, 0.690 mmol), XantPhos (40 mg, 0.069 mmol) and $Pd(OAc)_2$ (3.9 mg, 0.017 mmol). The reaction mixture was heated at 120 °C for 1 day, then allowed to cool to room temperature, concentrated onto Celite and then purified by silica gel chromatography (0-10% MeOH in DCM) to give the product **OXS007267** (22 mg, 0.065mmol, 19%) as an off-white solid. $^{1}$H NMR (500 MHz, $CDCl_3$) δ 8.81 (d, *J* = 1.0 Hz, 1H), 7.90 (s, 1H), 7.02 (d, *J* = 1.8 Hz, 2H), 6.93 (d, *J* = 2.0 Hz, 1H), 6.55 (d, *J* = 1.0 Hz, 1H), 3.97 (s, 3H), 3.93 (s, 3H), 3.89 – 3.76 (m, 4H), 3.51 – 3.45 (m, 4H); $^{13}$C NMR (126 MHz, $CDCl_3$) δ 157.3, 150.2, 149.4, 143.2, 142.1, 141.0, 135.8, 128.7, 117.0, 111.9, 108.3, 86.8, 67.0, 56.4, 47.2. *m/z*

(ESI$^+$) 341.1 ([M+H]$^+$, 100%); HRMS (ESI$^+$) $C_{18}H_{21}O_3N_4$ ([M+H]$^+$) requires 341.1608; found 341.1608.

**2-Chloro-*N*-(4-methoxyphenyl)-5-nitropyridin-4-amine (13)**

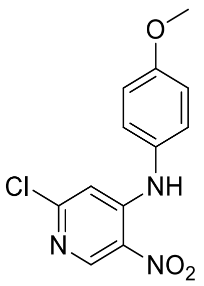

General procedure A; yield quant. (1.50 g, 5.37 mmol); gold solid. $^1$H NMR (400 MHz, $CDCl_3$) δ 9.52 (s, 1H), 9.08 (s, 1H), 7.22 – 7.15 (m, 2H), 7.07 – 6.95 (m, 2H), 6.77 (s, 1H), 3.87 (s, 3H); *m/z* (ESI$^+$) 280.0 ([M+H]$^+$, 100%).

**6-Chloro-*N*$^4$-(4-methoxyphenyl)pyridine-3,4-diamine (14)**

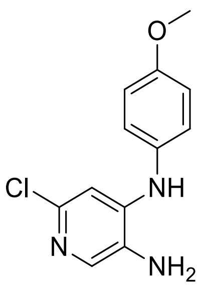

General procedure B.1 from **13** (1.50 g, 5.35 mmol); yield 97% (1.28 g, 5.11 mmol); purple foam. $^1$H NMR (400 MHz, $CDCl_3$) δ 7.74 (s, 1H), 7.13 – 7.05 (m, 2H), 6.98 – 6.90 (m, 2H), 6.69 (s, 1H), 5.94 (s, 1H), 3.83 (s, 3H), 3.22 (s, 2H); *m/z* (ESI$^+$) 250.1 ([M+H]$^+$, 100%).

**6-Chloro-1-(4-methoxyphenyl)-1*H*-imidazo[4,5-*c*]pyridine (15)**

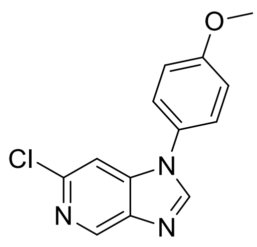

General procedure C from **14** (827 mg, 3.32 mmol); yield 53% (0.456 g, 1.76 mmol); brown solid.$^{1}$H NMR (400 MHz, DMSO-$d_6$) δ 8.88 (d, *J* = 0.9 Hz, 1H), 8.70 (s, 1H), 7.72 – 7.53 (m, 3H), 7.22 – 7.14 (m, 2H), 3.85 (s, 3H); *m/z* (ESI$^+$) 260.2 ([M+H]$^+$, 100%).

**6-(4-Fluoro-2-methylphenyl)-1-(4-methoxyphenyl)-1*H*-imidazo[4,5-*c*]pyridine (OXS007372)**

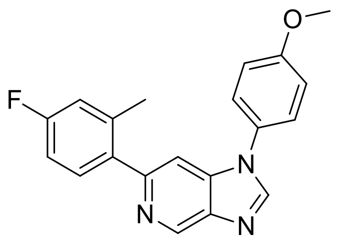

General procedure E.2 from **15** (81 mg, 0.31 mmol); yield 55% (57 mg, 0.17mmol); beige foam. $^{1}$H NMR (500 MHz, $CDCl_3$) δ 9.22 (s, 1H), 8.14 (d, *J* = 1.4 Hz, 1H), 7.42 (d, *J* = 8.9 Hz, 3H), 7.36 (dd, *J* = 8.4, 6.0 Hz, 1H), 7.14 – 7.06 (m, 2H), 7.02 – 6.88 (m, 2H), 3.89 (d, *J* = 1.4 Hz, 3H), 2.35 (s, 3H); $^{13}$C NMR (126 MHz, $CDCl_3$) δ 162.6 (d, *J* = 246 Hz), 160.0, 153.2, 144.3, 142.7, 139.7 (d, *J* = 23 Hz), 137.2 (d, *J* = 3 Hz), 131.6 (d, *J* = 8 Hz), 128.2, 125.7, 117.3 (d, *J* = 21 Hz), 115.5, 112.7 (d, *J* = 21 Hz), 105.9, 55.8, 20.7; *m/z* (ESI$^+$) 334.1 ([M+H]$^+$, 100%); HRMS (ESI$^+$) $C_{20}H_{17}ON_3F$([M+H]$^+$) requires 334.1350; found 334.1349.

**2-Chloro-*N*-(4-ethoxyphenyl)-5-nitropyridin-4-amine (16)**

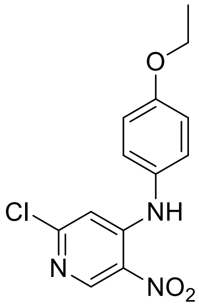

General procedure A; yield 81% (1.23 g, 4.20 mmol); gold solid. $^{1}H$ NMR (400 MHz, $CDCl_3$) δ 9.52 (s, 1H), 9.07 (s, 1H), 7.17 (dd, *J* = 9.0, 0.6 Hz, 2H), 6.99 (d, *J* = 8.9 Hz, 2H), 6.77 (s, 1H), 4.08 (q, *J* = 7.0 Hz, 2H), 1.46 (t, *J* = 7.0 Hz, 3H); *m/z* ($ESI^+$) 294.0 ($[M+H]^+$, 100%).

**6-Chloro-*N*$^{4}$-(4-ethoxyphenyl)pyridine-3,4-diamine (17)**

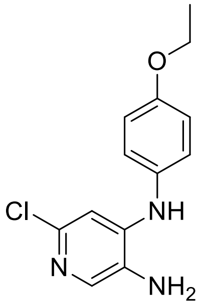

General procedure B.1 from **16** (1.35 g, 4.62 mmol); yield quant. (1.22 g, 4.62 mmol); purple solid.$^{1}H$ NMR (400 MHz, $CDCl_3$) δ 7.74 (s, 1H), 7.08 (d, *J* = 8.9 Hz, 2H), 6.92 (d, *J* = 8.9 Hz, 2H), 6.69 (s, 1H), 5.92 (s, 1H), 4.04 (q, *J* = 7.0 Hz, 2H), 3.21 (s, 2H), 1.43 (t, *J* = 7.0 Hz, 3H); *m/z* ($ESI^+$) 264.1 ($[M+H]^+$, 100%).

**6-Chloro-1-(4-ethoxyphenyl)-1*H*-imidazo[4,5-*c*]pyridine (18)**

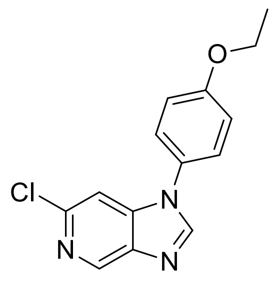

General procedure C from **17** (1.21 g, 4.59 mmol); yield quant. (1.26 g, 4.59 mmol); purple solid.$^{1}$H NMR (400 MHz, $CDCl_3$) δ 8.90 (d, *J* = 0.9 Hz, 1H), 8.08 (s, 1H), 7.47 – 7.29 (m, 2H), 7.07 (d, *J* = 8.9 Hz, 2H), 5.15 (s, 1H), 4.10 (q, *J* = 7.0 Hz, 2H), 1.46 (t, *J* = 7.0 Hz, 3H); *m/z* ($ESI^+$) 274.1 ($[M+H]^+$, 100%).

**1-(4-Ethoxyphenyl)-6-(4-fluoro-2-methylphenyl)-1*H*-imidazo[4,5-*c*]pyridine (OXS007370)**

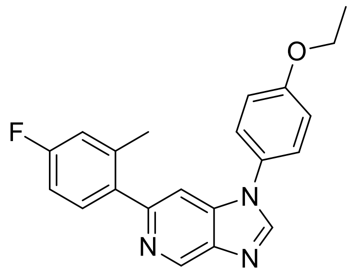

General procedure E.2 from **18** (85 mg, 0.31mmol); yield 76% (82 mg, 0.24mmol); yellow solid. $^{1}$H NMR (500 MHz, $CDCl_3$) δ 9.22 (d, *J* = 1.1 Hz, 1H), 8.14 (s, 1H), 7.44 – 7.38 (m, 3H), 7.38 – 7.34 (m, 1H), 7.07 (d, *J* = 8.8 Hz, 2H), 6.95 (ddd, *J* = 16.8, 9.1, 2.8 Hz, 2H), 4.11 (q, *J* = 7.0 Hz, 2H), 2.35 (s, 3H), 1.47 (t, *J* = 7.0 Hz, 3H); $^{13}$C NMR (126 MHz, $CDCl_3$) δ 162.6 (d, *J* = 246 Hz), 159.4, 153.2, 144.3, 142.7, 139.8, 139.6, 138.8 (d, *J* = 8 Hz), 137.2 (d, *J* = 3 Hz), 131.6 (d, *J* = 8 Hz), 128.0, 125.7, 117.3 (d, *J* = 21 Hz), 116.0, 112.7 (d, *J* = 21 Hz), 105.9, 64.2, 20.7 (d, *J* = 1 Hz), 14.9; *m/z* ($ESI^+$) 348.1 ($[M+H]^+$, 100%); HRMS ($ESI^+$) $C_{21}H_{19}ON_3F$($[M+H]^+$) requires 348.1507; found 348.1511.

***N*-(Benzo[*d*][1,3]dioxol-5-yl)-2-chloro-5-nitropyridin-4-amine (19)**

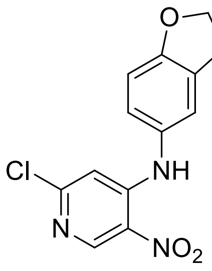

General procedure A; yield quant. (2.05 g, 6.97 mmol); dark gold solid. $^{1}$H NMR (400 MHz, $CDCl_3$) δ 9.47 (s, 1H), 9.07 (s, 1H), 6.90 (d, *J* = 8.7 Hz, 1H), 6.80 (s, 1H), 6.76 – 6.65 (m, 2H), 6.07 (s, 2H); *m/z* ($ESI^+$) 294.0 ($[M+H]^+$, 100%).

**$N^4$-(Benzo[*d*][1,3]dioxol-5-yl)-6-chloropyridine-3,4-diamine (20)**

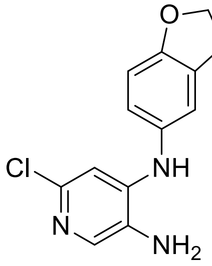

General procedure B.1 from **19** (1.51 g, 5.15 mmol); yield 81% (1.10 g, 4.17 mmol); brown solid.$^{1}$H NMR (400 MHz, $CDCl_3$) δ 7.75 (s, 1H), 6.81 (d, *J* = 8.0 Hz, 1H), 6.73 (s, 1H), 6.69 – 6.54 (m, 2H), 6.00 (s, 2H), 5.94 (s, 1H), 3.23 (s, 2H); *m/z* (ESI+) 264.0 ($[M+H]^+$, 100%).

**1-(Benzo[*d*][1,3]dioxol-5-yl)-6-chloro-1*H*-imidazo[4,5-*c*]pyridine (21)**

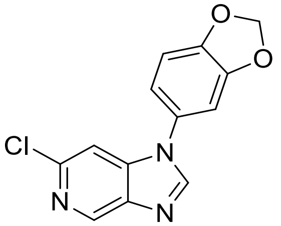

General procedure C from **20** (1.10 g, 4.17 mmol); yield quant. (1.14 g, 4.17mmol); dark brown solid.$^{1}$H NMR (400 MHz, $CDCl_3$) δ 8.92 (d, *J* = 0.9 Hz, 1H), 8.08 (s, 1H), 7.42 (d, *J* = 0.9 Hz, 1H), 6.99 (d, *J* = 8.8 Hz, 1H), 6.93 – 6.89 (m, 2H), 6.13 (s, 2H); *m/z* ($ESI^+$) 274.0 ($[M+H]^+$, 100%).

**1-(Benzo[*d*][1,3]dioxol-5-yl)-6-(4-fluoro-2-methylphenyl)-1*H*-imidazo[4,5-*c*]pyridine (OXS007392)**

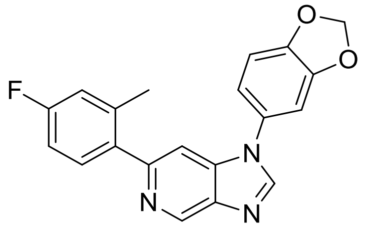

General procedure E.2 from **21** (85 mg, 0.31 mmol); yield 56% (61 mg, 0.18 mmol); beige solid. $^{1}$H NMR (500 MHz, $CDCl_3$) δ 9.22 (d, *J* = 1.0 Hz, 1H), 8.12 (s, 1H), 7.44 (d, *J* = 1.1 Hz, 1H), 7.36 (dd, *J* = 8.5, 5.9 Hz, 1H), 7.03 – 6.89 (m, 5H), 6.11 (s, 2H), 2.35 (s, 3H); $^{13}$C NMR (126 MHz, $CDCl_3$) δ 162.6 (d, *J* = 246 Hz), 153.3, 149.2, 148.2, 144.2, 142.7, 139.8, 139.5, 138.8 (d, *J* = 8 Hz), 137.2 (d, *J* = 3 Hz), 131.6 (d, *J* = 8 Hz), 129.2, 118.0, 117.3 (d, *J* = 21 Hz), 112.7 (d, *J* = 21 Hz), 109.2, 105.9, 105.7, 102.4, 20.7; *m/z* ($ESI^+$) 348.1 ($[M+H]^+$, 100%); HRMS ($ESI^+$) $C_{20}H_{15}O_2N_3F$($[M+H]^+$) requires 348.1143; found 348.1138.

**2-Chloro-*N*-(2,3-dihydrobenzofuran-5-yl)-5-nitropyridin-4-amine (22)**

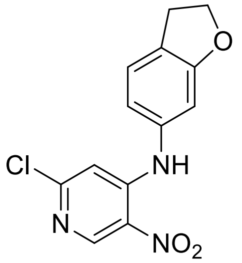

General procedure A; yield 94% (1.42 g, 4.85 mmol); pale brown solid.$^1$H NMR (400 MHz, $CDCl_3$) δ 9.48 (s, 1H), 9.07 (s, 1H), 7.10 – 6.94 (m, 2H), 6.87 (d, *J* = 8.4 Hz, 1H), 6.76 (s, 1H), 4.67 (t, *J* = 8.8 Hz, 2H), 3.28 (t, *J* = 8.8 Hz, 2H); *m/z* (ESI+) 292.0 ($[M+H]^+$, 100%).

**6-Chloro-*N*$^4$-(2,3-dihydrobenzofuran-5-yl)pyridine-3,4-diamine (23)**

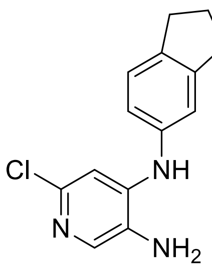

General procedure B.1 from **22** (1.40 g, 4.80 mmol); yield quant. (1.25 g, 4.80 mmol); brown solid.$^1$H NMR (400 MHz, $CDCl_3$) δ 7.74 (s, 1H), 7.02 (dt, *J* = 2.4, 1.2 Hz, 1H), 6.90 (dd, *J* = 8.4, 2.3 Hz, 1H), 6.79 (d, *J* = 8.4 Hz, 1H), 6.64 (s, 1H), 5.88 (s, 1H), 4.62 (t, *J* = 8.7 Hz, 2H), 3.24 (t, *J* = 8.7 Hz, 2H); *m/z* ($ESI^+$) 262.0 ($[M+H]^+$, 100%).

**6-Chloro-1-(2,3-dihydrobenzofuran-5-yl)-1*H*-imidazo[4,5-*c*]pyridine (24)**

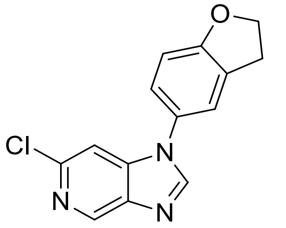

General procedure C from **23** (1.25 g, 4.80 mmol); yield 81% (1.05 g, 3.89 mmol); brown solid.$^1$H NMR (400 MHz, $CDCl_3$) δ 8.92 (d, *J* = 0.9 Hz, 1H), 8.07 (s, 1H), 7.39 (d, *J* = 0.9 Hz, 1H), 7.25 (s, 1H), 7.17 (ddt, *J* = 8.3, 2.3, 0.8 Hz, 1H), 6.95 (d, *J* = 8.4 Hz, 1H), 4.72 (t, *J* = 8.8 Hz, 2H), 3.34 (tt, *J* = 8.6, 1.0 Hz, 2H); *m/z* ($ESI^+$) 272.0 ($[M+H]^+$, 100%).

**1-(2,3-Dihydrobenzofuran-5-yl)-6-(4-fluoro-2-methylphenyl)-1*H*-imidazo[4,5-*c*]pyridine (OXS007540)**

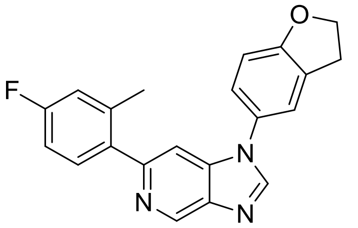

General procedure E.2 from **24** (96 mg, 0.35 mmol); yield 39% (47 mg, 0.14mmol); beige solid. $^{1}$H NMR (500 MHz, $CDCl_3$) δ 9.22 (d, *J* = 1.1 Hz, 1H), 8.12 (s, 1H), 7.41 (d, *J* = 1.1 Hz, 1H), 7.36 (dd, *J* = 8.4, 6.0 Hz, 1H), 7.29 (dt, *J* = 2.5, 1.2 Hz, 1H), 7.22 – 7.17 (m, 1H), 7.02 – 6.83 (m, 3H), 4.70 (t, *J* = 8.8 Hz, 2H), 3.32 (t, *J* = 8.7 Hz, 2H), 2.35 (s, 3H); $^{13}$C NMR (126 MHz, $CDCl_3$) δ 162.6 (d, *J* = 246 Hz), 160.7, 153.1, 144.5, 142.6, 139.8, 138.8 (d, *J* = 8 Hz), 137.2 (d, *J* = 3 Hz), 131.6 (d, *J* = 8 Hz), 129.5, 128.0, 124.7, 121.6, 117.3 (d, *J* = 21 Hz), 112.7 (d, *J* = 21 Hz), 110.5, 105.9, 72.1, 20.7; *m/z* (ESI$^+$) 346.1 ([M+H]$^+$, 100%); HRMS (ESI$^+$) $C_{21}H_{17}ON_3F$([M+H]$^+$) requires 346.13509; found 346.13502.

**2-Chloro-5-nitro-*N*-(p-tolyl)pyridin-4-amine (25)**

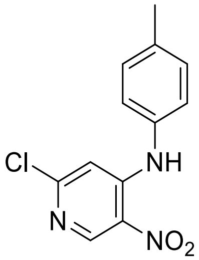

General procedure A; yield 82% (559 mg, 2.12 mmol); yellow solid. $^{1}$H NMR (400 MHz, $CDCl_3$) δ 9.57 (s, 1H), 9.08 (s, 1H), 7.30 (d, *J* = 8 Hz, 2H), 7.16 (d, *J* = 8 Hz, 2H), 6.86 (s,1H), 2.42 (s, 3H); $^{13}$C NMR (100 MHz, $CDCl_3$) δ 156.5, 149.3, 149.2, 138.1, 133.2, 130.9, 129.6, 125.5, 108.1, 21.1; *m/z* (ESI$^+$) 263.9 ([M+H]$^+$, 100%).

**6-Chloro-*N*-(p-tolyl)pyridine-3,4-diamine (26)**

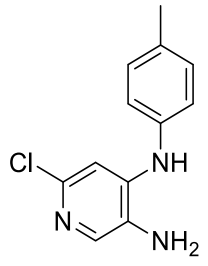

General procedure B.1 from **25** (545 mg, 2.07 mmol); yield 97% (470 mg, 2.01 mmol); yellow solid. $^{1}$H NMR (400 MHz, $CDCl_3$) δ 7.73 (s, 1H), 7.19 – 7.17 (m, 2H), 7.04 – 7.02 (m, 2H), 6.85 (s, 1H), 5.99 (s, 1H), 2.36 (s, 3H); $^{13}$C NMR (100 MHz, $CDCl_3$) δ 144.8, 144.5, 137.9, 136.7, 134.2, 130.3, 129.1, 122.1, 106.6, 20.9; *m/z* (ESI$^+$) 234.0 ([M+H]$^+$).

**6-Chloro-1-(*p*-tolyl)-1*H*-imidazo[4,5-*c*]pyridine (27)**

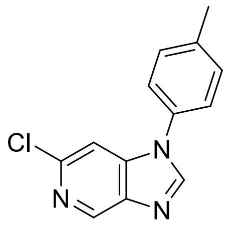

General procedure C from **26** (455 mg, 1.95 mmol); yield 73% (347 mg, 1.42 mmol); yellow solid.$^{1}$H NMR (400 MHz, DMSO-$d_6$) δ 8.88 (s, 1H), 8.72 (s, 1H), 7.64 (s, 1H), 6.59 (d, *J* = 8.4 Hz, 2H), 7.44 (d, *J* = 8 Hz,2H), 2.42 (s, 3H); $^{13}$C NMR (100 MHz, DMSO-$d_6$) δ 147.0, 143.7, 142.0, 141.1, 140.7, 138.7, 132.8, 131.0, 124.2, 106.5, 21.1; *m/z* (ESI$^+$) 244.0 ([M+H]$^+$, 100%).

**6-(4-Fluoro-2-methylphenyl)-1-(4-methylphenyl)-1*H*-imidazo[4,5-*c*]pyridine (OXS008411)**

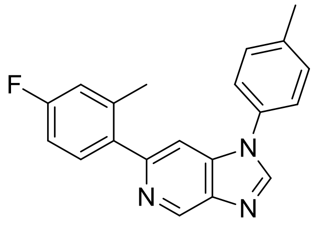

General procedure E.2 from **27** (150 mg, 0.620mmol); yield 29% (57 mg, 0.18 mmol); brown solid. $^1$H NMR (400 MHz, DMSO-$d_6$) δ 9.23 (d, *J* = 1.2 Hz, 1H), 8.17 (s, 1H), 7.47 (d, *J* = 1.2 Hz, 1H), 7.39 – 7.35 (m, 5H), 6.99 (dd, *J* = 2.4, 9.6 Hz, 1H), 6.94 (dt, 2.4, 8.4 Hz, 1H), 2.47 (s, 3H), 2.36 (s, 3H); $^{13}$C NMR (101 MHz, $CDCl_3$) δ 162.6 (d, *J* = 246 Hz), 153.3, 144.1, 142.8, 140.0, 139.4, 138.8 (d, *J* = 8 Hz), 137.3 (d, *J* = 3 Hz), 133.0, 131.7 (d, *J* = 8 Hz), 131.0, 124.0, 117.3 (d, *J* = 21 Hz), 112.7 (d, *J* = 21 Hz), 106.0, 21.3, 20.7; *m/z* (ESI$^+$) 318.2 ([M+H]$^+$, 100%).

**2-Chloro-*N*-(4-(methylsulfonyl)phenyl)-5-nitropyridin-4-amine (28)**

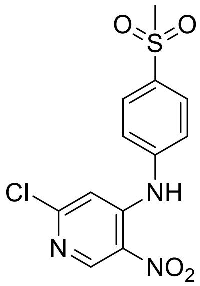

General procedure A; yield 58% (980 mg, 2.99 mmol); brown solid. $^1$H NMR (400 MHz, DMSO-$d_6$) δ 10.04 (br s, 1H), 9.0 (s, 1H), 8.01 – 7.99 (m, 2H), 7.66 – 7.64 (m, 2H), 7.09 (s, 1H), 3.25 (s, 3H); $^{13}$C NMR (100 MHz, DMSO-$d_6$) δ 155.3, 149.3, 147.5, 142.7, 138.4, 132.0, 129.2, 125.2, 109.7, 44.1; *m/z* (ESI$^+$) 328.0 ([M+H]$^+$, 100%).

**6-Chloro-N$^4$-(4-(methylsulfonyl)phenyl)pyridine-3,4-diamine (29)**

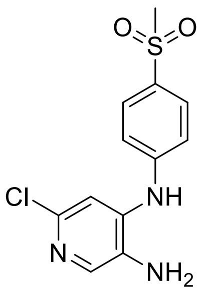

General procedure B.1 from **28** (1.00 g, 3.05 mmol); yield 15% (140 mg, 0.470 mmol); brown solid. $^{1}$H NMR (400 MHz, DMSO-$d_6$) δ 8.30 (s, 1H), 7.80 – 7.78 (m, 3H), 7.20 (dd, *J* = 2.0, 6.8 Hz, 2H), 7.04 (s, 1H), 5.15 (s,2H), 3.12 (s, 3H); $^{13}$C NMR (100 MHz, DMSO-$d_6$) δ 147.1, 138.3, 137.1, 136.3, 136.1, 132.4,129.3, 117.4, 111.9, 44.6;*m/z* (ESI$^+$) 298.1 ([M+H]$^+$, 100%).

**6-Chloro-1-(4-(methylsulfonyl)phenyl)-1*H*-imidazo[4,5-*c*]pyridine (30)**

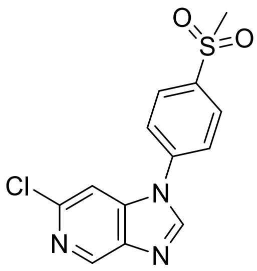

General procedure C from **29** (137 mg, 0.460 mmol); yield 89% (126 mg, 0.409 mmol); light brown solid. $^{1}$H NMR (400 MHz, DMSO-$d_6$) δ 8.92 (d, *J* = 0.4 Hz, 1H), 8.89 (s, 1H), 8.18 (dd, *J* = 2, 6.8 Hz, 2H), 8.05 (dd, *J* = 2, 6.8 Hz, 2H), 7.86 (d, *J* = 0.8 Hz, 1H), 3.12 (s, 3H); $^{13}$C NMR (100 MHz, DMSO-$d_6$) δ 146.8, 144.2, 142.2, 141.3, 140.7, 140.3, 139.4, 129.5, 124.9, 107.0, 44.0; *m/z* (ESI$^+$) 308.0 ([M+H]$^+$, 100%).

**6-(4-Fluoro-2-methylphenyl)-1-(4-(methylsulfonyl)phenyl)-1*H*-imidazo[4,5-*c*]pyridine (OXS007839)**

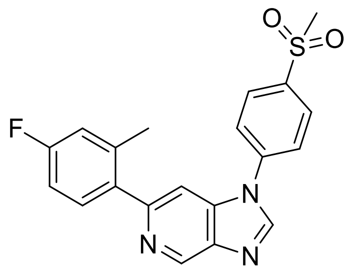

General procedure E from **30** (120 mg, 0.390 mmol); yield 28% (41 mg, 0.11 mmol); off-white solid. $^1$H NMR (400 MHz, DMSO-$d_6$) δ 9.15 (d, *J* = 0.8 Hz, 1H), 8.88 (s, 1H), 8.16 (dd, *J* = 6.8, 8.8 Hz, 2H), 8.08 (dd, *J* =28.8 Hz, 2H), 7.80 (d, *J* = 1.2 Hz, 1H), 7.52 – 7.48 (m, 1H), 7.17 (dd, *J* = 2.8, 10.4 Hz, 1H), 7.09 (td, *J* = 2.8, 8.8Hz, 1H), 3.31 (s, 3H), 2.35 (s, 3H); $^{13}$C NMR (100 MHz, DMSO-$d_6$) δ 161.6 (d, *J* = 244.0 Hz), 152.5, 145.1, 141.5, 139.8 (d, *J* = 11 Hz), 139.2, 138.7 (d, *J* = 8 Hz), 137.9, 137.0 (d, *J* = 3 Hz), 131.8 (d, *J* = 8 Hz), 129.0, 124.0, 116.7 (d, *J* = 21 Hz), 112.2 (d, *J* = 21 Hz), 106.3, 43.4, 20.2; *m/z* (ESI$^+$) 382.1 ([M+H]$^+$, 100%).

**2-Chloro-*N*-(4-(difluoromethoxy)phenyl)-5-nitropyridin-4-amine (31)**

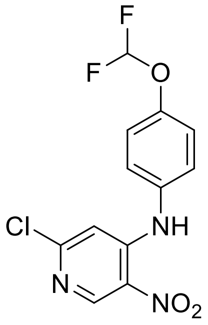

General procedure A; yield 77% (107 g, 338 mmol); gold solid. $^1$H NMR (400 MHz, $CDCl_3$) δ 9.58 (s, 1H), 9.10 (s, 1H), 7.32 – 7.26 (m, 4H), 6.85 (s, 1H), 6.57 (t, *J* = 73.2 Hz, 1H). $^{13}$C NMR (100 MHz, $CDCl_3$) δ 156.8, 150.2, 150.1, 150.0, 149.3, 148.9, 133.1, 129.7, 127.3, 121.6, 118.1, 115.5, 112.9, 108.0; *m/z* (ESI$^+$) 316.0 ([M+H]$^+$, 100%).

**6-Chloro-*N*$^4$-(4-(difluoromethoxy)phenyl)pyridine-3,4-diamine (32)**

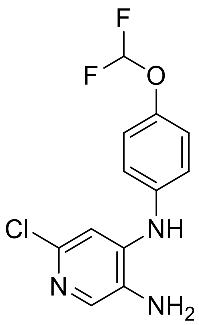

General procedure B.1 from **31** (53.0g, 170 mmol); yield quant. (48.0 g, 170 mmol); pale brown solid. $^1$H NMR (400 MHz, $CDCl_3$) δ 7.79 (s, 1H), 7.17 – 7.11 (m, 4H), 6.86 (s, 1H), 6.51 (t, $J$ = 74 Hz, 1H), 6.06 (s, 1H), 3.29 (s, 2H); $^{13}$C NMR (100 MHz, $CDCl_3$) δ147.3, 147.2, 144.2, 144.0, 138.0, 137.0, 129.6, 123.0, 121.3, 118.5, 115.9, 113.3, 107.0; *m/z* (ESI$^+$) 286.1 ([M+H]$^+$, 100%).

**6-Chloro-1-(4-(difluoromethoxy)phenyl)-1*H*-imidazo[4,5-*c*]pyridine (33)**

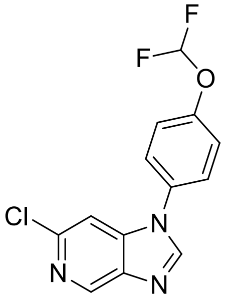

General procedure C from **32** (41.0 g, 150 mmol); yield 86% (39.0 g, 132 mmol); pale brown solid. $^1$H NMR (400 MHz, $CDCl_3$) δ 8.94 (s, 1H), 8.13 (s, 1H), 7.51 – 7.39 (m, 5H), 6.63 (t, $J$ = 72.8 Hz, 1H); $^{13}$C NMR (100 MHz, $CDCl_3$) δ 151.0, 144.9, 144.6, 142.7, 140.7, 140.6, 132.0, 125.6, 121.8, 118.0, 115.4, 112.7, 105.5; *m/z* (ESI$^+$) 296.0 ([M+H]$^+$, 100%).

**1-(4-(Difluoromethoxy)phenyl)-6-(4-fluoro-2-methylphenyl)-1*H*-imidazo[4,5-*c*]pyridine (OXS007417)**

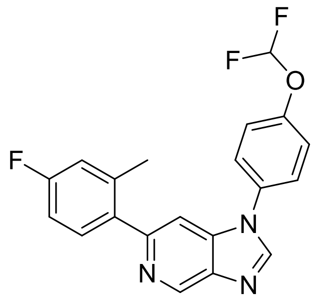

General procedure E from **33** (34 g, 120 mmol); yield 52% (23 g, 62 mmol); beige solid. $^{1}$H NMR (500 MHz, $CDCl_3$) δ 9.25 (d, *J* = 1.1 Hz, 1H), 8.17 (s, 1H), 7.55 – 7.50 (m, 2H), 7.46 (d, *J* = 1.1 Hz, 1H), 7.37 (dd, *J* = 8.5, 5.5 Hz, 3H), 7.04 – 6.90 (m, 2H), 6.60 (t, *J* = 72.8 Hz, 1H), 2.35 (s, 3H); $^{13}$C NMR (126 MHz, $CDCl_3$) δ 162.6 (d, *J* = 247 Hz), 153.6, 150.9 (t, *J* = 3 Hz), 143.9, 142.9, 139.9, 139.3, 138.8 (d, *J* = 8 Hz), 137.0 (d, *J* = 3 Hz), 132.6, 131.6 (d, *J* = 8 Hz), 125.7, 121.9, 117.4 (d, *J* = 21 Hz), 115.5 (t, *J* = 263 Hz), 112.8 (d, *J* = 21 Hz), 105.7, 29.9, 20.7; *m/z* ($ESI^+$) 370.1 ($[M+H]^+$, 100%); HRMS (APCI) $C_{20}H_{15}ON_3F_3$ ($[M+H]^+$) requires 370.11584; found 370.11617.

**2-Chloro-5-nitro-*N*-(4-(trifluoromethoxy)phenyl)pyridin-4-amine (34)**

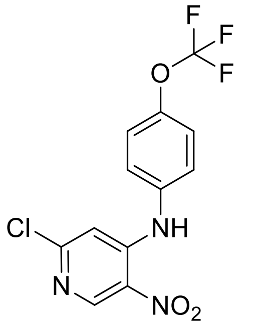

General procedure A; yield 57% (489 mg, 1.47 mmol);yellow solid.$^{1}$H NMR (400 MHz, $CDCl_3$) δ 9.61 (s, 1H), 9.11 (s, 1H), 7.39 – 7.33 (m, 4H), 6.88 (s, 1H); $^{13}$C NMR (100 MHz, $CDCl_3$) δ 156.9, 149.4, 148.6, 148.3, 148.2, 134.5, 129.8, 127.1, 122.8, 121.7, 119.1, 107.9; *m/z* ($ESI^+$) 334.0 ($[M+H]^+$, 100%).

**6-Chloro-N$^4$-(4-(trifluoromethoxy)phenyl)pyridine-3,4-diamine (35)**

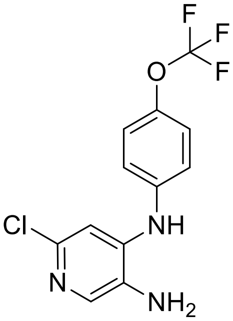

General procedure B.2 from **34** (480 mg, 1.44 mmol); yield 23% (100 mg, 0.329 mmol); brown oil. $^1$H NMR (400 MHz, DMSO-d$_6$) δ 7.86 (br s, 1H), 7.67 (s, 1H), 7.33 (d, *J* = 8.4 Hz, 2H), 7.24 – 7.22 (m, 2H), 6.81 (s, 1H), 5.03 (br s, 2H); $^{13}$C NMR (100 MHz, DMSO-d$_6$) δ 140.7, 139.6, 139.0, 135.3, 134.0, 122.7, 121.5, 107.8; *m/z* (ESI$^+$) 304.0 ([M+H]$^+$, 100%).

**6-Chloro-1-(4-(trifluoromethoxy)phenyl)-1*H*-imidazo[4,5-*c*]pyridine (36)**

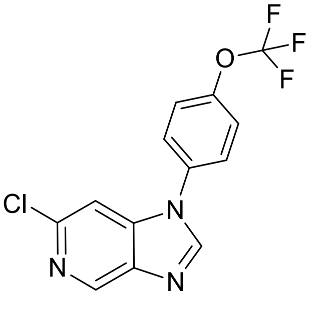

General procedure C from **35** (100 mg, 0.329 mmol), yield 88% (91 mg, 0.29 mmol);orange solid.$^1$H NMR (400 MHz, DMSO-*d*$_6$) δ 8.90 (s, 1H), 8.79 (s, 1H), 7.89 (d, *J* = 8.8 Hz, 2H), 7.76 (s, 1H), 7.64 (d, *J* =8.4 Hz, 2H); $^{13}$C NMR (100 MHz, DMSO-*d*$_6$) δ 148.3, 147.0, 143.9, 142.1, 141.1, 140.6, 134.3, 126.5, 123.2, 106.7; *m/z* (ESI$^+$) 314.0 ([M+H]$^+$, 100%).

**6-(4-Fluoro-2-methylphenyl)-1-(4-(trifluoromethoxy)phenyl)-1*H*-imidazo[4,5-*c*]pyridine (OXS007951)**

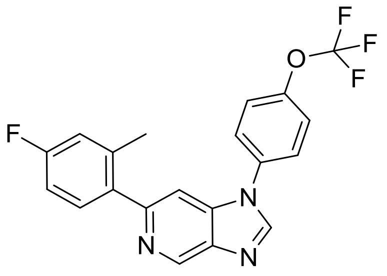

General procedure E.2 from **36** (85 mg, 0.27 mmol), yield 69% (72 mg, 0.19 mmol); yellow solid. $^{1}$H NMR (400 MHz, $CDCl_3$) δ 9.26 (d, *J* = 0.8 Hz, 1H), 8.18 (s, 1H), 7.59 – 7.56 (m, 2H), 7.48 – 7.43 (m, 3H),7.36 (dd, *J* = 6, 8 Hz, 1H), 7.00 (dd, *J* = 2.4, 9.6 Hz, 1H), 6.95 (dt, *J* = 2.4, 8.4 Hz, 1H), 2.36 (s, 3H); $^{13}$C NMR (100 MHz, $CDCl_3$) δ 162.7 (d, *J* = 247 Hz), 153.8, 149.2 – 149.2 (m), 143.7, 143.0, 140.0, 139.2, 138.9 (d, *J* = 8 Hz), 137.0 (d, *J* = 3 Hz), 134.0, 131.6 (d, *J* = 8 Hz), 125.7, 123.1, 117.4 (d, *J* = 21 Hz), 112.8 (d, *J* = 21 Hz), 105.7; *m/z* (ESI$^+$) 388.1 ($[M+H]^+$, 100%).

**2-Chloro-*N*-(3-fluoro-4-methoxyphenyl)-5-nitropyridin-4-amine (37)**

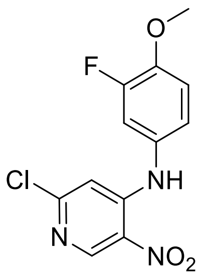

General procedure A; yield quant. (1.10 g, 3.68 mmol); dark yellow solid. $^{1}$H NMR (400 MHz, $CDCl_3$) δ 9.50 (s, 1H), 9.09 (s, 1H), 7.11 – 6.96 (m, 3H), 6.79 (s, 1H), 3.96 (s, 3H); *m/z* (ESI$^+$) 298.0 ($[M+H]^+$, 100%).

**6-Chloro-*N*$^4$-(3-fluoro-4-methoxyphenyl)pyridine-3,4-diamine (38)**

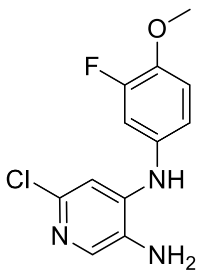

General procedure B.1 from **37**; yield 57% (484 mg, 1.81 mmol); dark purple foam. $^1$H NMR (400 MHz, $CDCl_3$) δ 7.77 (s, 1H), 6.98 (t, *J* = 8.8 Hz, 1H), 6.92 (dd, *J* = 11.9, 2.6 Hz, 1H), 6.88 (ddd, *J* = 8.6, 2.6, 1.3 Hz, 1H), 6.76 (s, 1H), 5.94 (s, 1H), 3.91 (s, 3H), 3.23 (s, 2H);; *m/z* (ESI$^+$) 268.0 ([M+H]$^+$, 100%).

**6-Chloro-1-(3-fluoro-4-methoxyphenyl)-1*H*-imidazo[4,5-*c*]pyridine (39)**

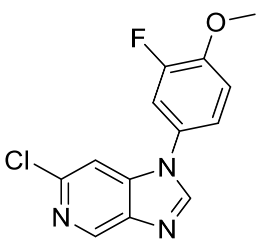

General procedure C from **38**; yield 47% 231 mg, 0.832 mmol); dark yellow solid. $^1$H NMR (400 MHz, DMSO-$d_6$) δ 8.89 (d, *J* = 0.9 Hz, 1H), 8.73 (s, 1H), 7.75 (dd, *J* = 11.9, 2.6 Hz, 1H), 7.71 (d, *J* = 0.9 Hz, 1H), 7.54 (ddd, *J* = 8.8, 2.6, 1.4 Hz, 1H), 7.41 (t, *J* = 9.0 Hz, 1H), 3.94 (s, 3H); *m/z* (ESI$^+$) 278.1 ([M+H]$^+$, 100%).

**6-(4-Fluoro-2-methylphenyl)-1-(3-fluoro-4-methoxyphenyl)-1*H*-imidazo[4,5-*c*]pyridine (OXS007373)**

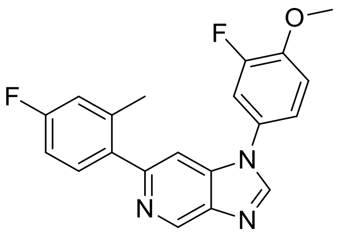

General procedure E.2 from **39** (86 mg, 0.31 mmol; yield 83% (91 mg, 0.26mmol); beige foam. $^{1}$H NMR (500 MHz, $CDCl_3$) δ 9.29 – 9.13 (m, 1H), 8.13 (s, 1H), 7.48 – 7.42 (m, 1H), 7.36 (dd, *J* = 8.4, 5.9 Hz, 1H), 7.29 – 7.19 (m, 2H), 7.15 (t, *J* = 8.6 Hz, 1H), 7.01 – 6.86 (m, 2H), 3.98 (s, 3H), 2.35 (s, 3H); $^{13}$C NMR (126 MHz, $CDCl_3$) δ 162.6 (d, *J* = 246 Hz), 153.7, 153.5, 151.7, 148.4 (d, *J* = 10 Hz), 144.0, 142.8, 139.8, 139.4, 138.8 (d, *J* = 8 Hz), 137.1 (d, *J* = 3 Hz), 131.6 (d, *J* = 8 Hz), 128.0 (d, *J* = 8 Hz), 120.4 (d, *J* = 4 Hz), 117.4 (d, *J* = 21 Hz), 114.4 (d, *J* = 3 Hz), 113.0 (d, *J* = 21 Hz), 112.7 (d, *J* = 21 Hz), 105.7, 56.7, 20.7; *m/z* ($ESI^+$) 352.1 ($[M+H]^+$, 100%); HRMS ($ESI^+$) $C_{20}H_{17}ON_3F$($[M+H]^+$) requires 334.1350; found 334.1349.

**2-Chloro-*N*-(3-chloro-4-methoxyphenyl)-5-nitropyridin-4-amine (40)**

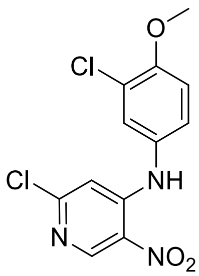

General procedure A; yield 81% (659 mg, 2.10 mmol); yellow solid. $^{1}$H NMR (400 MHz, $CDCl_3$) δ 9.47 (s, 1H), 9.08 (s, 1H), 7.32 (d, *J* = 2.8 Hz, 1H), 7.17 (dd, *J* = 2.4, 8.8 Hz, 1H), 7.04 (d, *J* = 8.4 Hz, 1H), 6.76 (s, 1H), 3.97 (s, 3H); $^{13}$C NMR (100 MHz, $CDCl_3$) δ 156.7, 154.9, 149.3, 149.2, 129.6, 128.8, 128.3, 125.7, 124.0, 113.0, 108.1, 56.5; *m/z* ($ESI^+$) 314.0 ($[M+H]^+$, 100%).

**6-Chloro-$N^4$-(3-chloro-4-methoxyphenyl)pyridine-3,4-diamine (41)**

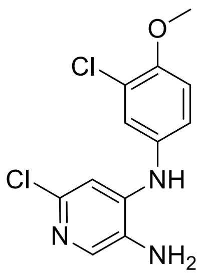

General procedure B.1 from **40** (650 mg, 2.07 mmol); yield quant. (597 mg, 2.10 mmol); off-white solid. $^{1}$H NMR (400 MHz, $CDCl_3$) δ 7.76 (s, 1H), 7.19 (d, *J* = 2.8 Hz, 1H), 7.05 (dd, *J* = 2.8, 8.8 Hz, 1H), 6.95 (d, *J* = 8.8 Hz, 1H), 6.71 (s, 1H), 5.97 (s, 1H), 3.92 (s, 3H); $^{13}$C NMR (100 MHz, $CDCl_3$) δ 152.6, 145.1, 144.7, 138.2, 132.7, 128.8, 125.3, 123.4, 122.4, 113.0, 106.5, 56.5;*m/z* (ESI$^+$) 284.0 ([M+H]$^+$, 100%).

**6-Chloro-1-(3-chloro-4-methoxyphenyl)-1*H*-imidazo[4,5-*c*]pyridine (42)**

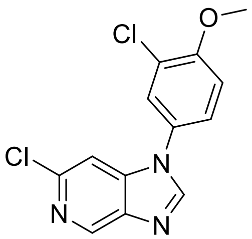

General procedure C from **41** (587 mg, 2.07 mmol); yield 93% (563 mg, 1.91 mmol); off-white solid. $^{1}$H NMR (400 MHz, DMSO-$d_6$) δ 8.86 (s, 1H), 8.68 (s, 1H), 7.86 (d, *J* = 2.8 Hz, 1H), 7.67 (dd, *J* = 2.4, 8.8 Hz, 1H), 7.64 (d, *J* = 0.4 Hz, 1H), 7.38 (d, *J* = 8.8 Hz, 1H), 3.96 (s, 3H); $^{13}$C NMR (100 MHz, DMSO-$d_6$) δ 152.3, 147.2, 143.7, 141.9, 141.0, 140.9, 128.3, 126.6, 125.0, 122.6, 114.2, 106.5, 57.1; *m/z* (ESI$^+$) 294.0 ([M+H]$^+$, 100%).

**1-(3-Chloro-4-methoxyphenyl)-6-(4-fluoro-2-methylphenyl)-1*H*-imidazo[4,5-c]pyridine (OXS008397)**

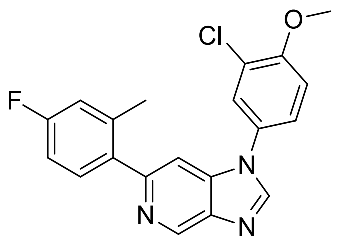

General procedure E.2 from **42** (150 mg, 0.510 mmol); yield 38% (72 mg, 0.20 mmol); off-white solid. $^1$H NMR (400 MHz, DMSO-$d_6$) δ 9.10 (d, $J$ = 1.2 Hz, 1H), 8.67 (s, 1H), 7.88 (d, $J$ = 2.4 Hz, 1H), 7.70 (dd, $J$ = 2.8, 8.8 Hz, 1H), 7.58 (d, $J$ = 1.2 Hz, 1H), 7.49 – 7.45 (m, 1H), 7.37 (d, $J$ = 8.8 Hz, 1H), 7.14 (dd, $J$ = 2.8, 10 Hz, 1H), 7.07 (td, $J$ = 2.8, 8.4 Hz, 1H), 3.95 (s, 3H), 2.34 (s, 3H); $^{13}$C NMR (100 MHz, DMSO-$d_6$) δ 161.5 (d, $J$ = 244 Hz), 154.4, 151.9, 145.3, 141.2, 139.3, 138.6, 138.2 (d, $J$ = 8 Hz), 137.1 (d, $J$ = 3 Hz), 128.2, 125.7, 124.1, 122.0, 116.6 (d, $J$ = 21 Hz), 113.6, 112.2 (d, $J$ = 21.0 Hz), 105.8, 56.5, 20.1; *m/z* (ESI$^+$) 368.0 ([M+H]$^+$, 100%).

**2-Chloro-*N*-(4-methoxy-3-methylphenyl)-5-nitropyridin-4-amine (43)**

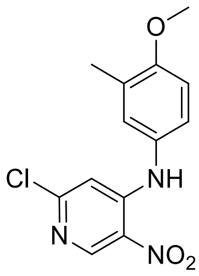

General procedure A; yield 87% (660 mg, 2.24 mmol); yellow solid. $^1$H NMR (400 MHz, $CDCl_3$) δ 9.48 (s, 1H), 9.06 (s, 1H), 7.08 – 7.03 (m, 2H), 6.90 (d, $J$ = 8.4 Hz, 1H), 6.78 (s,1H), 3.89 (s, 3H), 2.26 (s, 3H); $^{13}$C NMR (100 MHz, $CDCl_3$) δ 157.4, 156.3, 149.7, 149.2, 130.0, 129.0, 128.3, 127.9, 124.5, 110.9, 108.2, 55.6,16.3; *m/z* (ESI$^+$) 294.0 ([M+H]$^+$, 100%).

**6-Chloro-*N*-(4-methoxy-3-methylphenyl)pyridine-3,4-diamine (44)**

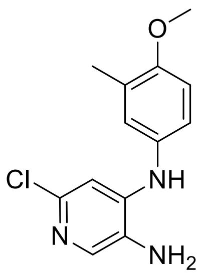

General procedure B.1 from **43** (650mg, 2.20 mmol); yield 18% (106 mg, 0.362mmol); purple solid. $^{1}$H NMR (400 MHz, DMSO-$d_6$) δ 7.54 (s, 1H), 7.43 (s, 1H), 7.01 – 6.94 (m, 3H), 6.45 (s, 1H), 4.87 (s, 2H), 3.79 (s,3H), 2.16 (s, 3H); $^{13}$C NMR (100 MHz, DMSO-$d_6$) δ 154.7, 142.5, 139.6, 134.3, 132.8, 132.1, 127.3, 126.2, 122.1, 117.7, 104.8, 56.0,16.4; *m/z* (ESI$^+$) 264.0 ([M+H]$^+$, 100%).

**6-Chloro-1-(4-methoxy-3-methylphenyl)-1*H*-imidazo[4,5-*c*]pyridine (45)**

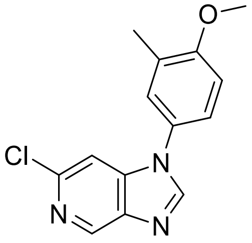

General procedure C from **44** (100 mg, 0.380mmol); yield 94% (98 mg, 0.36 mmol); purple solid. $^{1}$H NMR (400 MHz, DMSO-$d_6$) δ 8.86 (d, $J$ = 0.8 Hz, 1H), 8.65 (s, 1H), 7.60 (d, $J$ = 0.4 Hz, 1H), 7.50 – 7.47 (m, 2H), 7.17 – 7.15 (m, 1H), 3.89 (s, 3H), 2.26 (s, 3H); $^{13}$C NMR (100 MHz, DMSO-$d_6$) δ 157.9, 147.2, 143.6, 142.0, 141.1, 140.9, 128.2, 127.5, 126.6, 123.3, 118.9,106.4, 56.3, 16.3; *m/z* (ESI$^+$) 274.0 ([M+H]$^+$, 100%).

**6-(4-Fluoro-2-methylphenyl)-1-(4-methoxy-3-methylphenyl)-1*H*-imidazo[4,5-*c*]pyridine (OXS008444)**

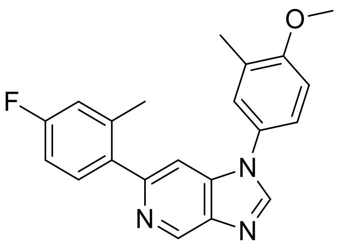

General procedure E.2 from **45** (90 mg, 0.33 mmol); yield 15% (17 mg, 0.049 mmol); off-white solid.$^{1}$H NMR (400 MHz, $CDCl_3$) δ 9.23 (s, 1H), 8.13 (s, 1H), 7.43 (s, 1H), 7.38 – 7.35 (m, 1H), 7.30 – 7.26 (m, 2H), 7.00 – 6.94 (m, 3H), 3.91 (s, 3H), 2.36 (s, 3H), 2.31 (s, 3H); $^{13}$C NMR (101 MHz, $CDCl_3$) δ 162.6 (d, *J* = 246.4 Hz), 158.2, 153.0, 144.5, 142.6, 139.8, 139.7, 138.8 (d, *J* = 7.9 Hz), 137.2 (d, *J* = 2.5 Hz), 131.7 (d, *J* = 8.5 Hz), 129.2, 127.7, 126.7, 122.9, 117.3 (d, *J* = 21.1 Hz), 112.7 (d, *J* = 21.2 Hz), 110.9, 106.0, 55.9, 20.7 (d, *J* = 1.4 Hz), 16.5; *m/z* ($ESI^+$) 348.0 ($[M+H]^+$, 100%).

**2-Chloro-*N*-(3-isopropyl-4-methoxyphenyl)-5-nitropyridin-4-amine (46)**

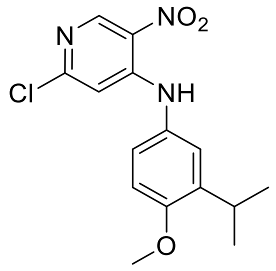

General procedure A; yield 99% (1.70 g, 5.12 mmol); gold solid. $^{1}$H NMR (400 MHz, $CDCl_3$) δ 9.53 (s, 1H), 9.07 (s, 1H), 7.12 – 7.02 (m, 2H), 6.96 – 6.87 (m, 1H), 6.80 (s, 1H), 3.89 (s, 3H), 3.35 (hept, *J* = 6.9 Hz, 1H), 1.22 (d, *J* = 6.9 Hz, 6H); *m/z* ($ESI^+$) 322.0 ($[M+H]^+$, 100%).

**6-Chloro-*N*$^4$-(3-isopropyl-4-methoxyphenyl)pyridine-3,4-diamine (47)**

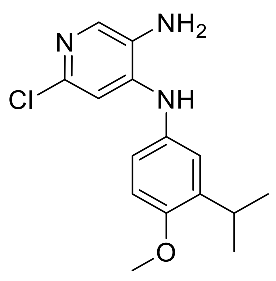

General procedure B.1 from **46** (1.66 g, 5.16 mmol); yield 79% (1.19 mg, 4.10 mmol); grey solid. $^{1}$H NMR (400 MHz, $CDCl_3$) δ 7.75 (s, 1H), 7.01 – 6.94 (m, 2H), 6.89 – 6.83 (m, 1H), 6.71 (s, 1H), 5.90 (s, 1H), 3.85 (s, 3H), 3.33 (hept, *J* = 6.9 Hz, 1H), 3.18 (s, 2H), 1.21 (d, *J* = 6.9 Hz, 6H); *m/z* (ESI$^+$) 292.1 ([M+H]$^+$, 100%).

**6-Chloro-1-(3-isopropyl-4-methoxyphenyl)-1H-imidazo[4,5-c]pyridine (48)**

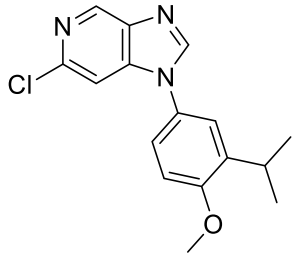

General procedure C from **47** (1.25 g, 4.28 mmol); yield 99% (1.28 g, 4.27 mmol); grey solid. $^{1}$H NMR (400 MHz, $CDCl_3$) δ 8.93 (d, *J* = 0.9 Hz, 1H), 8.10 (s, 1H), 7.38 (d, *J* = 0.9 Hz, 1H), 7.27 – 7.20 (m, 2H), 7.04 – 6.97 (m, 1H), 3.93 (s, 3H), 3.40 (hept, *J* = 6.9 Hz, 1H), 1.26 (d, *J* = 6.9 Hz, 6H); *m/z* (ESI$^+$) 302.1 ([M+H]$^+$, 100%).

**6-(4-Fluoro-2-methylphenyl)-1-(3-isopropyl-4-methoxyphenyl)-1*H*-imidazo[4,5-*c*]pyridine (OXS007787)**

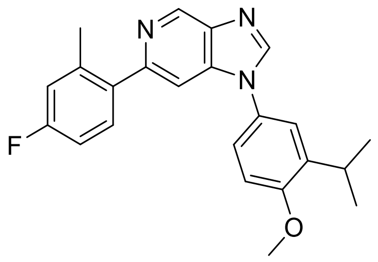

General procedure E.1 from **48** (100 mg, 0.331 mmol); yield 37% (47 mg, 0.13 mmol); white solid. $^{1}$H NMR (400 MHz, $CDCl_3$) δ 9.23 (d, *J* = 1.1 Hz, 1H), 8.15 (s, 1H), 7.42 (d, *J* = 1.1 Hz, 1H), 7.38 (dd, *J* = 8.4, 6.0 Hz, 1H), 7.32 – 7.24 (m, 2H), 7.03 – 6.89 (m, 3H), 3.91 (s, 3H), 3.40 (hept, *J* = 6.8 Hz, 1H), 2.37 (s, 3H), 1.25 (d, *J* = 6.9 Hz, 6H); $^{13}$C NMR (126 MHz, $CDCl_3$) δ 162.5 (d, *J* = 246 Hz), 157.2, 153.0, 144.4, 142.7, 139.8, 139.7, 139.6, 138.7 (d, *J* = 8 Hz), 137.3 (d, *J* = 3 Hz), 131.7 (d, *J* = 8 Hz), 128.1, 122.6, 122.5, 117.3 (d, *J* = 21 Hz), 112.7 (d, *J* = 21 Hz), 111.4, 106.0, 55.9, 27.1, 22.6, 20.7 (d, *J* = 1 Hz); *m/z* (ESI$^{+}$) 376.2 ([M+H]$^{+}$, 100%); HRMS (ESI$^{+}$) $C_{23}H_{23}ON_3F^{+}$ ([M+H]$^{+}$) requires 376.1820; found 376.1819.

**N-(2-Chloro-5-nitropyridin-4-yl)-6-methoxypyridin-3-amine (49)**

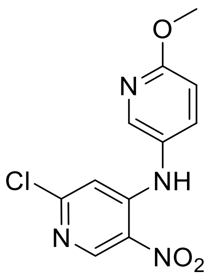

General procedure A; yield 98% (1.43 g, 5.10 mmol); dark brown solid. $^{1}$H NMR (400 MHz, $CDCl_3$) δ 9.43 (s, 1H), 9.09 (s, 1H), 8.13 (dt, *J* = 2.8, 0.7 Hz, 1H), 7.49 (ddd, *J* = 8.7, 2.7, 0.5 Hz, 1H), 6.89 (dd, *J* = 8.8, 0.7 Hz, 1H), 6.69 (s, 1H), 4.00 (s, 3H); *m/z* (ESI$^{+}$) 281.1 ([M+H]$^{+}$, 100%).

**6-Chloro-$N^{4}$-(6-methoxypyridin-3-yl)pyridine-3,4-diamine (50)**

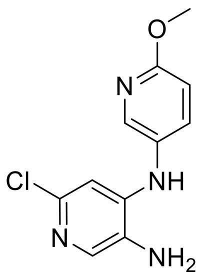

General procedure B.1 from **49** (1.43 g, 5.10 mmol); yield 93% (1.18 g, 4.72 mmol); dark red solid. $^{1}$H NMR (400 MHz, $CDCl_3$) δ 8.03 (d, *J* = 2.7 Hz, 1H), 7.78 (s, 1H), 7.44 (dd, *J* = 8.8, 2.8 Hz, 1H), 6.81 (dd, *J* = 8.8, 0.6 Hz, 1H), 6.58 (s, 1H), 5.90 (s, 1H), 3.96 (s, 3H), 3.22 (s, 2H);*m/z* (ESI+) 251.0 ($[M+H]^+$, 100%).

**6-Chloro-1-(6-methoxypyridin-3-yl)-1*H*-imidazo[4,5-*c*]pyridine (51)**

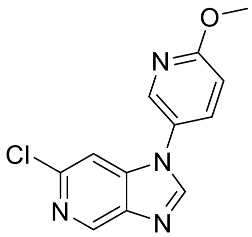

General procedure C from **50** (1.18 g, 4.72 mmol); yield 97% (1.18 g, 4.53 mmol); dark brown solid.$^{1}$H NMR (400 MHz, $CDCl_3$) δ 8.95 (d, *J* = 0.9 Hz, 1H), 8.31 (dd, *J* = 2.8, 0.7 Hz, 1H), 8.07 (s, 1H), 7.67 (dd, *J* = 8.8, 2.8 Hz, 1H), 7.37 (d, *J* = 1.0 Hz, 1H), 6.98 (dd, *J* = 8.8, 0.7 Hz, 1H), 4.03 (s, 3H); *m/z* (ESI+) 261.1 ([M+H]+, 100%).

**6-(4-Fluoro-2-methylphenyl)-1-(6-methoxypyridin-3-yl)-1*H*-imidazo[4,5-*c*]pyridine (OXS007749)**

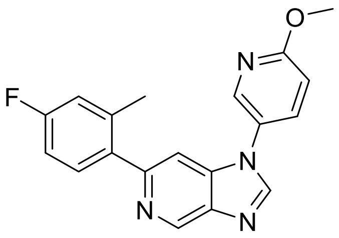

General procedure E.2 from **51** (82 mg, 0.311 mmol); yield 98% (102 mg, 0.305 mmol); pale brown solid. $^1$H NMR (500 MHz, $CDCl_3$) δ 9.25 (d, *J* = 1.0 Hz, 1H), 8.35 (d, *J* = 2.7 Hz, 1H), 8.12 (s, 1H), 7.71 (dd, *J* = 8.8, 2.8 Hz, 1H), 7.39 (d, *J* = 1.0 Hz, 1H), 7.35 (dd, *J* = 8.4, 5.9 Hz, 1H), 7.08 – 6.74 (m, 3H), 4.02 (s, 3H), 2.35 (s, 3H); $^{13}$C NMR (126 MHz, $CDCl_3$) δ 164.3, 162.6 (d, *J* = 246 Hz), 153.6, 144.0, 143.0, 142.8, 139.8 (d, *J* = 3 Hz), 138.8 (d, *J* = 8 Hz), 137.0 (d, *J* = 3 Hz), 135.1, 131.6 (d, *J* = 8 Hz), 126.0, 117.4 (d, *J* = 21 Hz), 112.8 (d, *J* = 21 Hz), 112.6, 105.5, 54.3, 20.7; *m/z* ($ESI^+$) 335.1 ($[M+H]^+$, 100%); HRMS ($ESI^+$) $C_{19}H_{16}ON_4F$($[M+H]^+$) requires 335.1303; found 335.1303.

**5-(6-(4-Fluoro-2-methylphenyl)-1*H*-imidazo[4,5-*c*]pyridin-1-yl)pyridin-2-ol (OXS007801)**

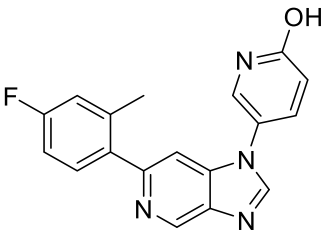

**OXS007749** (60 mg, 0.18 mmol) and pyridine hydrochloride (747 mg, 6.46 mmol) were heated in a vial with a heatgun for 3 minutes. The mixture was cooled, diluted with water and extracted into EtOAc, acidified slightly and extracted further. The combined organics were evaporated to dryness, then, to remove the remaining pyridine, the residue was stirred in water overnight, filtered, washed with further water, triturated with hexane and dried *in vacuo* to give **OXS007801** (15 mg, 0.047 mmol, 26%) as a brown solid.$^1$H NMR (400 MHz, DMSO-$d_6$) δ 11.98 (br s, 1H), 9.08 (d, *J* = 1.2 Hz, 1H), 8.55 (s, 1H), 7.94 (d, *J* = 2.8 Hz, 1H),7.78 (dd, *J* = 3.2, 9.6 Hz, 1H), 7.56 (d, *J* = 0.8 Hz, 1H), 7.48 – 7.44 (m, 1H), 7.17 – 7.13 (m, 1H), 7.10 – 7.06 (m, 1H),6.53 (d, *J* = 9.6 Hz,

1H), 2.34 (s, 3H); $^{13}$C NMR (100 MHz, DMSO-$d_6$) δ 161.6, 161.5 (d, $J$ = 244 Hz), 151.6, 145.9, 141.0, 139.4, 139.2, 138.6, 138.5, 137.1, 131.8 (d, $J$ = 8 Hz), 116.7 (d, $J$ = 21 Hz), 112.2 (d, $J$ = 21.0 Hz), 105.9, 20.2. *m/z* (ESI$^+$) 321.1 ([M+H]$^+$, 100%).

***N*-(2-Chloro-5-nitropyridin-4-yl)-6-(difluoromethoxy)pyridin-3-amine (52)**

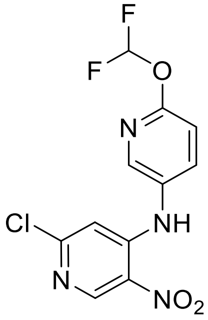

General procedure A at 60 ºC; yield 54% (800 mg, 2.53 mmol); dark brown oil. $^1$H NMR (400 MHz, $CDCl_3$) δ 9.50 (s, 2H), 9.12 (s, 1H), 8.20 (dt, $J$ = 2.8, 0.7 Hz, 1H), 7.71 – 7.68 (m, 1H), 7.48 (t, $J$ = 76.7 Hz, 1H), 6.72 (s, 1H); *m/z* (ESI$^+$) 317.0 ([M+H]$^+$, 100%).

**6-Chloro-*N*$^4$-(6-(difluoromethoxy)pyridin-3-yl)pyridine-3,4-diamine (53)**

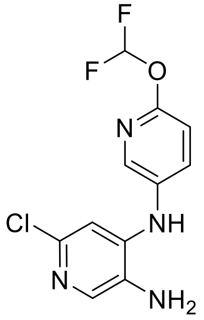

General procedure B.1 from **52**; yield 61% (1.17 g, 2.86 mmol); dark brown gum. $^1$H NMR (400 MHz, $CDCl_3$) δ 8.05 (d, $J$ = 2.7 Hz, 1H), 7.83 (s, 1H), 7.58 (dd, $J$ = 8.7, 2.6 Hz, 1H), 7.42 (d, $J$ =

72.6 Hz, 1H), 6.97 (d, *J* = 8.6 Hz, 1H), 6.71 (s, 1H), 6.02 (s, 1H), 3.28 (s, 2H);*m/z* ($ESI^+$) 287.0 ($[M+H]^+$, 100%).

**6-Chloro-1-(6-(difluoromethoxy)pyridin-3-yl)-1*H*-imidazo[4,5-*c*]pyridine (54)**

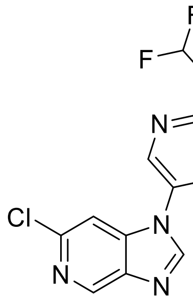

General procedure C from **53**; yield 76% (913 mg, 3.08 mmol); dark brown solid.$^1$H NMR (400 MHz, $CDCl_3$) δ 8.97 (d, *J* = 0.9 Hz, 1H), 8.39 (dd, *J* = 2.8, 0.7 Hz, 1H), 8.10 (s, 1H), 7.88 (dd, *J* = 8.7, 2.8 Hz, 1H), 7.52 (t, *J* = 72.1 Hz, 1H), 7.39 (d, *J* = 1.0 Hz, 1H), 7.19 (dd, *J* = 8.7, 0.7 Hz, 1H); *m/z* (ESI+) 297.0 ($[M+H]^+$, 100%).

**1-(6-(Difluoromethoxy)pyridin-3-yl)-6-(4-fluoro-2-methylphenyl)-1*H*-imidazo[4,5-*c*]pyridine (OXS007571)**

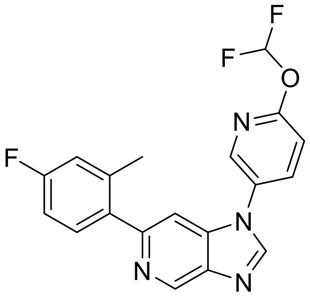

General procedure E.2 from **54** (92 mg, 0.31 mmol); yield 51% (59 mg, 0.16 mmol); off-white foam. $^1$H NMR (500 MHz, $CDCl_3$) δ 9.25 (d, *J* = 1.1 Hz, 1H), 8.17 (s, 1H), 7.56 – 7.50 (m, 2H), 7.46 (d, *J* = 1.1 Hz, 1H), 7.41 – 7.33 (m, 3H), 7.02 – 6.89 (m, 2H), 6.60 (t, *J* = 72.8 Hz, 1H), 2.35

(s, 3H); $^{13}$C NMR (126 MHz, $CDCl_3$) δ 162.7 (d, $J$ = 247 Hz), 158.9 (t, $J$ = 3.9 Hz), 154.0, 143.5, 143.0, 142.8, 139.8, 139.4, 138.8 (d, $J$ = 8 Hz), 136.8 (d, $J$ = 3 Hz), 136.3, 135.3, 131.6 (d, $J$ = 9 Hz), 129.1, 117.4 (d, $J$ = 21 Hz), 114.0 (t, $J$ = 258 Hz), 113.2, 112.8 (d, $J$ = 21 Hz), 105.3, 20.7 (d, $J$ = 1 Hz); *m/z* ($ESI^+$) 370.1 ($[M+H]^+$, 100%); HRMS (ESI) $C_{19}H_{14}ON_4F_3$ ($[M+H]^+$) requires 370.11139; found 370.11142.

***N*-(2-chloro-5-nitropyridin-4-yl)-5-methoxypyridin-2-amine (55)**

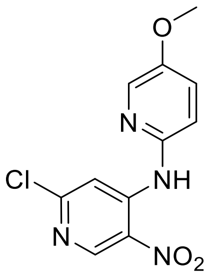

General procedure A; yield 79% (574 mg, 2.05 mmol); red solid. 1H NMR (400 MHz, DMSO-$d_6$) δ 10.14 (s, 1H), 8.96 (s, 1H), 8.26 (s, 1H), 8.15 (d, $J$ = 3.2 Hz, 1H), 7.51 (dd, $J$ =3.2, 8.8 Hz, 1H), 7.33 (d, $J$ = 8.8 Hz, 1H), 3.85 (s, 3H); $^{13}$C NMR (100 MHz, DMSO-$d_6$) δ 155.1, 153.4, 148.8, 1456, 145.5, 134.3, 131.5, 125.2, 117.8, 110.8, 56.5; *m/z* (ESI+) 281.0 ($[M+H]^+$, 100%).

**6-Chloro-*N*⁴-(5-methoxypyridin-2-yl)pyridine-3,4-diamine (56)**

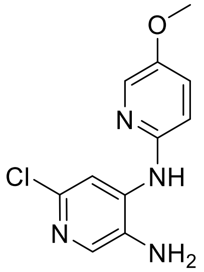

General procedure B.2 from **55** (560 mg, 2.00 mmol); yield 70% (350 mg, 1.40 mmol); brown solid. 1H NMR (400 MHz, DMSO-$d_6$) δ 8.12 – 8.11 (m, 2H), 8.03 (d, $J$ = 2.8 Hz, 1H), 7.65 (s,

1H), 7.40 (dd, $J$ = 3.2, 8.8 Hz,1H), 7.06 (d, $J$ = 8.8 Hz, 1H), 5.07 (s, 2H), 3.80 (s, 3H); $^{13}$C NMR (100 MHz, DMSO-$d_6$) δ 151.0, 149.0, 139.2, 138.0, 135.0, 133.0, 132.8, 125.6, 114.1, 109.3, 56.4; $m/z$ (ESI$^+$) 251.0 ([M+H]$^+$, 100%).

**6-Chloro-1-(5-methoxypyridin-2-yl)-1*H*-imidazo[4,5-*c*]pyridine (57)**

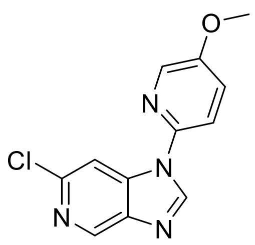

General procedure C from **56** (340 mg, 1.36 mmol); yield 75% (265 mg, 1.02 mmol); beige solid. $_{1}$H NMR (400 MHz, DMSO-$d_6$) δ 9.06 (s, 1H), 8.87 (d, $J$ = 0.8 Hz, 1H), 8.36 (d, $J$ = 3.2 Hz, 1H), 8.18 (d, $J$ = 0.8 Hz,1H), 7.94 (d, $J$ = 8.8 Hz, 1H), 7.71 (dd, $J$ = 3.2, 8.8 Hz, 1H), 3.92 (s, 3H); $^{13}$C NMR (100 MHz, DMSO-$d_6$) δ 155.2, 145.5, 144.0, 142.5, 142.0, 141.3, 139.4, 136.2, 125.0, 116.3, 109.0, 56.7; $m/z$ (ESI$^+$) 261.0 ([M+H]$^+$, 100%).

**6-(4-Fluoro-2-methylphenyl)-1-(5-methoxypyridin-2-yl)-1*H*-imidazo[4,5-*c*]pyridine (OXS008495)**

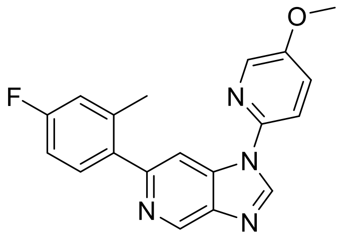

General procedure E.2 from **57** (150 mg, 0.575 mmol); yield 34% (65 mg, 0.19 mmol); yellow solid. $^{1}$H NMR (400 MHz, $CDCl_3$) δ 9.22 (s, 1H), 8.53 (s, 1H), 8.29 (d, $J$ = 2.8 Hz, 1H), 7.95 (s, 1H), 7.51 – 7.39 (m, 3H), 7.02 – 6.94 (m, 2H), 3.95 (s, 3H), 2.37 (s, 3H); $^{13}$C NMR (100 MHz,

$CDCl_3$) δ 162.6 (d, $J$ = 246 Hz), 155.2, 153.7, 142.8, 142.7, 142.5, 140.3, 138.9 (d, $J$ = 8 Hz), 138.0, 137.4 (d, $J$ = 3 Hz), 136.6, 131.7 (d, $J$ = 8 Hz), 124.1, 117.3 (d, $J$ = 21 Hz), 115.4, 112.7 (d, $J$ = 21 Hz), 107.9, 56.3, 20.7 (d, $J$ = 1 Hz); $m/z$ (ESI$^+$) 335.0 ([M+H]$^+$, 100%).

**_N_-(2-Chloro-5-nitropyridin-4-yl)-5-methoxypyrazin-2-amine (58)**

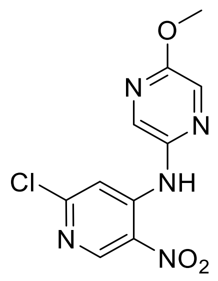

General procedure A; yield 14% (200 mg, 0.71 mmol); yellow solid.$^1$H NMR (400 MHz, DMSO-$d_6$) δ 10.26 (s, 1H), 8.99 (s, 1H), 8.31 (d, $J$ = 1.6 Hz, 1H), 8.21 (d, $J$ = 1.6 Hz, 1H), 7.83(s, 1H), 3.94 (s, 3H); $m/z$ (ESI$^+$) 282.0 ([M+H]$^+$, 100%).

**6-Chloro-_N_$^4$-(5-methoxypyrazin-2-yl)pyridine-3,4-diamine (59)**

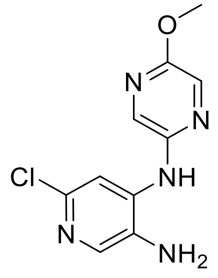

General procedure B.2 from **58** (300 mg, 1.07 mmol);yield 56% (150 mg, 0.596 mmol); brown solid. $^1$H NMR (400 MHz, $CDCl_3$) δ 8.01 (d,$J$ = 1.2 Hz, 1H), 7.92 (d,$J$ = 1.6 Hz, 1H), 7.87 (s, 1H), 7.64 (s, 1H), 6.72 (brs, 1H), 3.97 (s, 3H), 3.25 (brs, 2H);$m/z$ (ESI$^+$) 252.1 ([M+H]$^+$, 100%).

**6-Chloro-1-(5-methoxypyrazin-2-yl)-1H-imidazo[4,5-c]pyridine (60)**

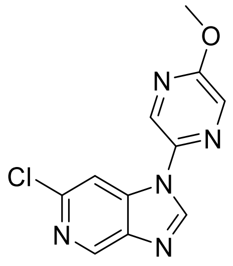

General procedure C from **59** (200 mg, 0.790 mmol); yield 87%; light brown solid (180 mg, 0.688 mmol). $^1$H NMR (400 MHz, DMSO-*d*$_6$) δ 9.06 (s, 1H), 8.90 (d,*J* = 0.8 Hz, 1H), 8.86 (d,*J* = 1.2 Hz, 1H), 8.41 (d,*J* = 1.2 Hz, 1H), 8.10 (d,*J* = 0.8 Hz, 1H), 4.02 (s, 3H);$^{13}$C NMR (101 MHz, DMSO-*d*$_6$) δ 159.5, 145.4, 142.1, 141.1, 139.8, 139.6, 134.0, 133.6, 108.6, 54.8;*m/z* (ESI$^+$) 262.1 ([M+H]$^+$, 100%).

**6-(4-Fluoro-2-methylphenyl)-1-(5-methoxypyrazin-2-yl)-1*H*-imidazo[4,5-*c*]pyridine (OXS007837)**

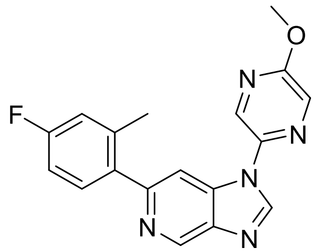

General procedure E.2 from **60** (90 mg, 0.34 mmol); yield 33% (38 mg, 0.11 mmol); light brown solid. $^1$H NMR (400 MHz, DMSO-*d*$_6$) δ 9.13 (d, *J* = 1.0 Hz, 1H), 9.02 (s, 1H), 8.88 (d, *J* = 1.4 Hz, 1H), 8.39 (d, *J* = 1.4 Hz, 1H), 8.06 (d, *J* = 1.0 Hz, 1H), 7.46 (dd, *J* = 8.5, 6.2 Hz, 1H), 7.17 (dd, *J* = 10.2, 2.7 Hz, 1H), 7.10 (td, *J* = 8.5, 2.7 Hz, 1H), 4.01 (s, 3H), 2.34 (s, 3H); $^{13}$C NMR (101 MHz, DMSO-*d*$_6$) δ 162.1 (d, *J* = 244.2 Hz), 159.4, 153.0, 144.4, 141.9, 140.1, 139.1 (d, *J* = 8.1 Hz), 138.0, 137.7 (d, *J* = 3.0 Hz), 134.0, 133.6, 132.2 (d, *J* = 8.6 Hz), 117.3 (d, *J* = 21.0 Hz), 112.9 (d, *J* = 21.0 Hz), 108.6, 54.7, 20.7 (d, *J* = 1.5 Hz);*m/z* (ESI$^+$) 336.1 ([M+H]$^+$, 100%).

**N-(2-Chloro-5-nitropyridin-4-yl)-2-methoxypyrimidin-5-amine (61)**

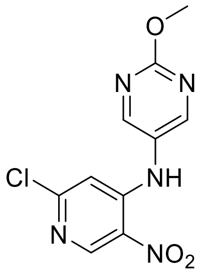

General procedure A; yield 71% (655 mg, 2.33 mmol); yellow solid. $^1$H NMR (400 MHz, $CDCl_3$) δ 9.39 (br s, 1H), 9.12 (s, 1H), 8.53 (s, 2H), 6.66 (s, 1H), 4.10 (s, 3H); $^{13}$C NMR (100 MHz, $CDCl_3$) δ 164.8, 157.9, 157.2, 149.3, 130.0, 125.6, 107.8, 55.8; *m/z* ($ESI^+$) 282.1 ($[M+H]^+$, 50%).

**6-Chloro-$N^4$-(2-methoxypyrimidin-5-yl)pyridine-3,4-diamine (62)**

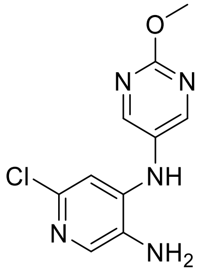

To a stirred solution of tin(II) chloride (883 mg, 4.65 mmol) in concentrated HCl (5 ml), at <10 °C was added **61** (437 mg, 1.55mmol) portionwise over 15min. The suspension was stirred at 0°C for 2 h, then basified to pH 10 with 5 N NaOH, and extracted with EtOAc. The organics were combined, dried over anhydrous $MgSO_4$ and distilled to provide **62** (141 mg, 0.560 mg, 36%) as a brown solid. $^1$H NMR (400 MHz, DMSO-$d_6$) δ 8.49 (s, 2H), 7.70 (s, 1H), 7.62 (s, 1H), 6.49 (s, 1H), 4.96 (s, 2H), 3.93 (s, 3H). *m/z* ($ESI^+$) 252.1 ($[M+H]^+$, 100%).

**6-Chloro-1-(2-methoxypyrimidin-5-yl)-1*H*-imidazo[4,5-*c*]pyridine (63)**

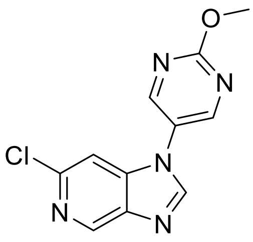

General procedure C from **62** (180 mg, 0.715mmol); yield 76%; (142 mg, 0.543 mmol); brown solid. $^1$H NMR (400 MHz, DMSO-$d_6$) δ 9.00 (s, 2H), 8.89 (s, 1H), 8.71 (s, 1H), 7.82 (s, 1H), 4.04 (s, 3H); $^{13}$C NMR (101 MHz, DMSO-$d_6$) δ 164.9 156.6, 147.2, 143.8, 141.9, 141.5, 140.8, 125.7, 106.8, 55.8; *m/z* (ESI$^+$) 262.1 ([M+H]$^+$, 100%).

**6-(4-Fluoro-2-methylphenyl)-1-(2-methoxypyrimidin-5-yl)-1*H*-imidazo[4,5-*c*]pyridine (OXS007785)**

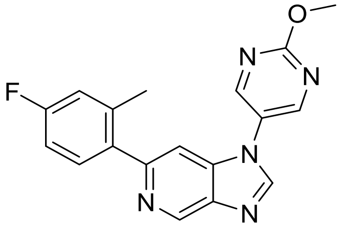

General procedure E.2 from **63** (130 mg, 0.497mmol); yield 38% (63 mg, 0.19 mmol); off-white solid. $^1$H NMR (400 MHz, DMSO-$d_6$) δ 9.13 (d, *J* = 1.0 Hz, 1H), 9.03 (s, 2H), 8.70 (s, 1H), 7.73 (d, *J* = 1.0 Hz, 1H), 7.49 (dd, *J* = 8.5, 6.2 Hz, 1H), 7.15 (dd, *J* = 10.2, 2.7 Hz, 1H), 7.12 – 7.05 (m, 1H), 4.03 (s, 3H), 2.35 (s, 3H); $^{13}$C NMR (101 MHz, DMSO-$d_6$) δ 164.8, 162.1 (d, *J* = 244.0 Hz), 156.3, 152.7, 146.0, 141.7, 139.8, 139.6, 139.2 (d, *J* = 8.1 Hz), 137.6 (d, *J* = 3.0 Hz), 132.4 (d, *J* = 8.5 Hz), 126.1, 117.3 (d, *J* = 21.1 Hz), 112.7 (d, *J* = 20.9 Hz), 106.6, 55.8, 20.8 (d, *J* = 1.5 Hz);*m/z* (ESI$^+$) 336.1 ([M+H]$^+$, 100%).

**2-Chloro-*N*-(cyclopropylmethyl)-5-nitropyridin-4-amine (64)**

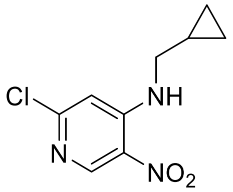

General procedure A; yield 99% (908 mg, 4.00 mmol); brown solid. $^{1}H$ NMR (400 MHz, $CDCl_3$) δ 7.64 (s, 1H), 6.42 (s, 1H), 2.98 (dd, $J$ = 7.0, 5.1 Hz, 2H), 1.21 – 1.02 (m, 1H), 0.73 – 0.51 (m, 2H), 0.28 (dt, $J$ = 5.9, 4.6 Hz, 2H); $m/z$ ($ESI^+$) 228.0 ($[M+H]^+$, 100%).

**6-Chloro-*N*$^4$-(cyclopropylmethyl)pyridine-3,4-diamine (65)**

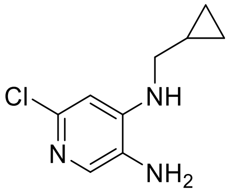

General procedure B.1 from **64** (830 mg, 3.64 mmol); yield 84% (610 mg, 3.10 mmol); grey solid. $^{1}H$ NMR (400 MHz, $CDCl_3$) δ 7.64 (s, 1H), 6.42 (s, 1H), 3.22 – 2.80 (m, 2H), 1.22 – 0.96 (m, 1H), 0.77 – 0.46 (m, 2H), 0.37 – 0.11 (m, 2H); $m/z$ ($ESI^+$) 198.1 ($[M+H]^+$, 100%).

**6-Chloro-1-(cyclopropylmethyl)-1*H*-imidazo[4,5-*c*]pyridine (66)**

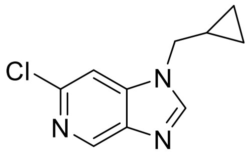

General procedure C from **65** (500 mg, 2.52 mmol); yield 78% (410 mg, 1.98 mmol); grey solid. $^{1}H$ NMR (400 MHz, $CDCl_3$) δ 8.86 (d, $J$ = 0.9 Hz, 1H), 8.07 (s, 1H), 7.40 (d, $J$ = 0.9 Hz, 1H), 4.00 (d, $J$ = 7.1 Hz, 2H), 1.35 – 1.26 (m, 1H), 0.85 – 0.66 (m, 2H), 0.45 (dt, $J$ = 6.2, 5.0 Hz, 2H); $m/z$ ($ESI^+$) 208.0 ($[M+H]^+$, 100%).

**1-(Cyclopropylmethyl)-6-(4-fluoro-2-methylphenyl)-1*H*-imidazo[4,5-*c*]pyridine (OXS007812)**

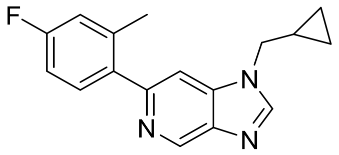

General procedure E.2 from **66** (100 mg, 0.483 mmol); yield 33% (45 mg, 0.10 mmol); grey solid. $^{1}$H NMR (500 MHz, $CDCl_3$) δ 9.26 – 9.11 (m, 1H), 8.12 (s, 1H), 7.45 – 7.38 (m, 2H), 7.06 – 6.96 (m, 2H), 4.07 (d, $J$ = 7.0 Hz, 2H), 2.39 (s, 2H), 1.35 (dqd, $J$ = 12.2, 7.5, 4.9 Hz, 1H), 0.83 – 0.68 (m, 2H), 0.47 (q, $J$ = 5.1 Hz, 2H); $^{13}$C NMR (126 MHz, CDCl3) δ 162.4 (d, $J$ = 246 Hz), 152.2, 144.3, 142.3, 139.8, 139.4, 138.6 (d, $J$ = 8.1 Hz), 137.2 (d, $J$ = 3 Hz), 131.5 (d, $J$ = 8 Hz), 117.2 (d, $J$ = 21 Hz), 112.6 (d, $J$ = 21 Hz), 105.3, 49.9, 20.6, 20.6, 10.9, 4.6; *m/z* ($ESI^+$) 282.1 ($[M+H]^+$, 100%); HRMS ($ESI^+$) $C_{17}H_{16}N_3F^+$ ($[M+H]^+$) requires 282.1328; found 282.1401.

***N*-(2-Chloro-5-nitropyridin-4-yl)-1-methyl-1*H*-indazol-5-amine (67)**

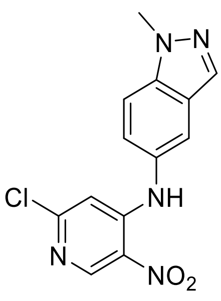

General procedure A; yield 90% (1.42 g, 4.69 mmol); brown solid. $^{1}$H NMR (400 MHz, $CDCl_3$) δ 9.68 (s, 1H), 9.10 (s, 1H), 8.04 (d, $J$ = 1.0 Hz, 1H), 7.72 – 7.46 (m, 2H), 7.28 (d, $J$ = 2.0 Hz, 1H), 6.75 (s, 1H), 4.14 (s, 3H); *m/z* ($ESI^+$) 304.0 ($[M+H]^+$, 100%).

**6-Chloro-*N*$^4$-(1-methyl-1*H*-indazol-5-yl)pyridine-3,4-diamine (68)**

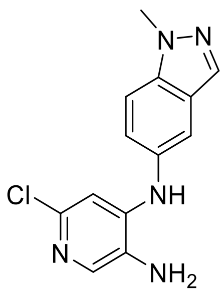

General procedure B.1 from **67** (1.40 g, 4.62 mmol); yield 78% (986 mg, 3.60 mmol); brown solid. $^1$H NMR (400 MHz, $CDCl_3$) δ 7.96 (d, *J* = 1.0 Hz, 1H), 7.79 (s, 1H), 7.54 – 7.38 (m, 2H), 7.23 (dd, *J* = 8.9, 1.9 Hz, 1H), 6.70 (s, 1H), 6.07 (s, 1H), 4.11 (s, 3H); *m/z* ($ESI^+$) 274.1 ($[M+H]^+$, 100%).

**6-Chloro-1-(1-methyl-1*H*-indazol-5-yl)-1*H*-imidazo[4,5-*c*]pyridine (69)**

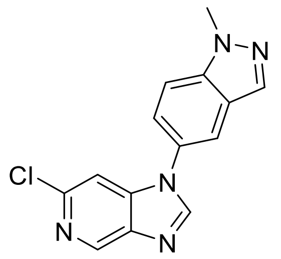

General procedure C from **68** (968 mg, 3.55 mmol); yield 84% (845 mg, 2.98 mmol); brown solid. $^1$H NMR (400 MHz, $CDCl_3$) δ 8.96 (d, *J* = 1.0 Hz, 1H), 8.17 (s, 1H), 8.12 (d, *J* = 1.0 Hz, 1H), 7.82 (dd, *J* = 2.0, 0.7 Hz, 1H), 7.63 (dt, *J* = 8.8, 0.9 Hz, 1H), 7.48 – 7.38 (m, 2H), 4.19 (s, 3H); *m/z* ($ESI^+$) 284.0 ($[M+H]^+$, 100%).

**6-(4-Fluoro-2-methylphenyl)-1-(1-methyl-1*H*-indazol-5-yl)-1*H*-imidazo[4,5-*c*]pyridine (OXS007570)**

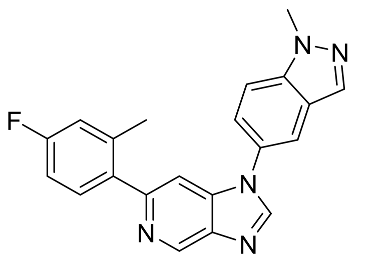

General procedure E.2 from **69** (100 mg, 0.350 mmol); yield 11% (14 mg, 0.039 mmol); beige solid. $^1$H NMR (500 MHz, $CDCl_3$) δ 9.27 (s, 1H), 8.25 (s, 1H), 8.11 (d, *J* = 1.0 Hz, 1H), 7.94 – 7.82 (m, 1H), 7.61 (d, *J* = 8.7 Hz, 1H), 7.51 – 7.41 (m, 2H), 7.36 (dd, *J* = 8.4, 6.0 Hz, 1H), 7.03 – 6.85 (m, 2H), 4.17 (s, 3H), 2.36 (s, 3H); $^{13}$C NMR (126 MHz, $CDCl_3$) δ 162.6 (d, *J* = 246 Hz), 153.3, 144.6, 142.8, 140.0, 139.9, 139.4, 138.8 (d, *J* = 8 Hz), 137.1, 133.4, 131.6 (d, *J* = 8 Hz), 128.5, 124.4, 123.2, 117.4 (d, *J* = 21 Hz), 117.2, 112.7 (d, *J* = 21 Hz), 111.0, 105.9, 36.1, 20.7; *m/z* (ESI$^+$) 358.1 ([M+H]$^+$, 100%); HRMS (ESI$^+$) $C_{21}H_{17}N_5F$ ([M+H]$^+$) requires 358.14600; found 358.14625.

**1-(4-(Difluoromethoxy)phenyl)-6-(4-fluorophenyl)-1*H*-imidazo[4,5-*c*]pyridine (OXS008278)**

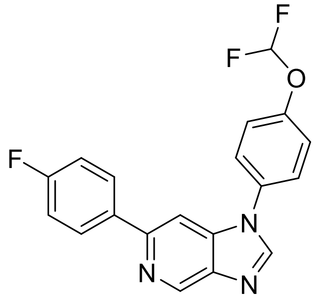

General procedure E.2 from **33** (150 mg, 0.507 mmol); yield 81% (146 mg, 0.411 mmol); pale brown solid. $^1$H NMR (400 MHz, $CDCl_3$) δ 9.54 (d, *J* = 0.8 Hz, 1H), 8.13 (s, 1H), 8.01 – 7.97 (m, 2H), 7.72 (d, *J* = 0.8 Hz, 1H), 7.56 – 7.53 (m, 2H), 7.42 (d, *J* = 1.6 Hz, 2H), 7.17 – 7.12 (m, 2H), 6.63 (t, *J* = 72.8 Hz, 1H); $^{13}$C NMR (100 MHz, $CDCl_3$) δ 163.4 (d, *J* = 248 Hz), 151.5, 151.0 (d, *J* = 3 Hz), 144.0, 143.3, 140.1 (d, *J* = 45 Hz), 136.1 (d, *J* = 3 Hz), 132.6, 129.0 (d, *J* = 8 Hz), 125.9, 121.9, 115.8 (d, *J* = 22 Hz), 115.6 (t, *J* = 263 Hz), 101.8; *m/z* (ESI$^+$) 356.0 ([M+H]$^+$, 100%).

**1-[4-(Difluoromethoxy)phenyl]-6-(4-fluoro-2-methoxyphenyl)-1H-imidazo[4,5-c]pyridine (OXS007739)**

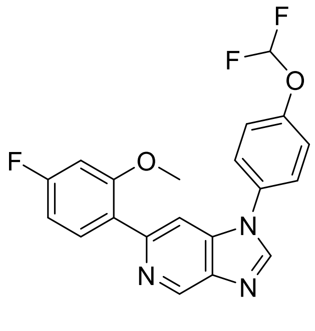

General procedure E.2 from **33** (300 mg, 1.01 mmol); yield 42% (163 mg, 0.420 mmol); pale brown solid. $^1$H NMR (400 MHz, $CDCl_3$) δ 9.25 (d, *J* = 1.2 Hz, 1H), 8.14 (s, 1H), 7.89 (d, *J* = 1.2 Hz, 1H), 7.82 (dd, *J* = 6.8,8.4 Hz, 1H), 7.54 (dd, *J* = 2, 6.8 Hz, 2H), 7.38 (dd, *J* = 2, 6.8 Hz, 2H), 6.82 – 6.77 (m, 1H), 6.73 (dd, *J* = 2.4, 10.8Hz, 1H), 6.62 (t, *J* = 72.8 Hz, 1H), 3.82 (s, 3H); $^{13}$C NMR (101 MHz, $CDCl_3$) δ 163.8 (d, *J* = 248 Hz), 158.0 (d, *J* = 10 Hz), 150.8 (t, *J* = 3 Hz), 149.4, 143.8, 143.1, 140.1, 139.0, 132.9, 132.8 (d, *J* = 10 Hz), 125.6 (d, *J* = 4 Hz), 125.6, 121.7, 115.6 (t, *J* = 263 Hz), 107.8 (d, *J* = 21 Hz), 106.5, 99.7 (d, *J* = 26 Hz), 56.1; *m/z* (ESI$^+$) 386.1 ([M+H]$^+$, 100%).

**1-[4-(Difluoromethoxy)phenyl]-6-(2,6-dimethylphenyl)imidazo[4,5-c]pyridine (OXS007924)**

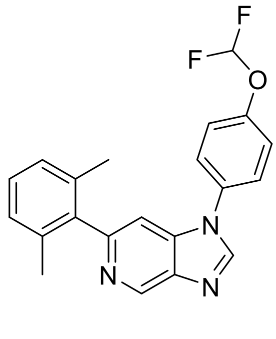

General procedure E.1 from **33** (100 mg, 0.340 mmol); yield 46% (58 mg, 0.16 mmol); white solid. $^1$H NMR (500 MHz, $CDCl_3$) δ 9.31 (d, *J* = 1.0 Hz, 1H), 8.21 (s, 1H), 7.57 – 7.51 (m, 2H), 7.42 –

7.34 (m, 3H), 7.21 (dd, $J$ = 8.3, 6.8 Hz, 1H), 7.12 (d, $J$ = 7.6 Hz, 2H), 6.61 (t, $J$ = 72.9 Hz, 1H), 2.06 (s, 6H). $^{13}$C NMR (126 MHz, $CDCl_3$) δ 153.8, 150.6 (t, $J$ = 3 Hz), 143.6, 143.4, 140.7, 139.7, 139.1, 136.1, 132.6, 127.9, 127.5, 125.4, 121.7, 115.4 (t, $J$ = 263 Hz), 105.8, 20.4; *m/z* (ESI$^+$) 366.1 ([M+H]$^+$, 100%); HRMS (ESI$^+$) $C_{21}H_{18}N_3OF_2^+$ ([M+H]$^+$) requires 362.14992; found 362.14938.

**6-(2-Chloro-4-fluorophenyl)-1-(4-(difluoromethoxy)phenyl)-1*H*-imidazo[4,5-*c*]pyridine (OXS008296)**

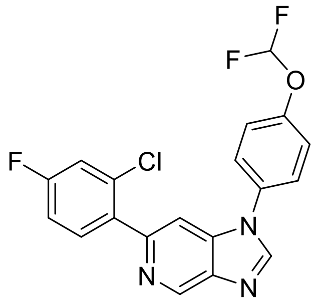

General procedure E.2 from **33** (150 mg, 0.507 mmol); yield 27% (54 mg, 0.14 mmol); white solid. $^1$H NMR (400 MHz, $CDCl_3$) δ 9.27 (d, $J$ = 1.2 Hz, 1H), 8.19 (s, 1H), 7.74 (d, $J$ = 1.2 Hz, 1H), 7.68 – 7.64 (m, 1H), 7.54 (dd, $J$ = 2.0, 6.8 Hz, 2H), 7.38 (d, $J$ = 8.8 Hz, 2H), 7.23 (dd, $J$ = 2.4, 8.4 Hz, 1H), 7.12 – 7.07 (m, 1H), 6.61 (t, $J$ = 72.8 Hz, 1H); $^{13}$C NMR (100 MHz, $CDCl_3$) δ 162.4 (d, $J$ = 251 Hz), 151.0 (t, $J$ = 3 Hz), 150.1, 144.2, 143.4, 140.5, 138.8, 136.0 (d, $J$ = 4 Hz), 133.3 (d, J = 9 Hz), 133.2 (d, $J$ = 10 Hz), 132.6, 125.7, 121.9, 117.4 (d, $J$ = 25 Hz), 115.6 (t, $J$ = 263 Hz), 114.5 (d, $J$ = 21 Hz), 106.8; *m/z* (ESI$^+$) 390.0 ([M+H]$^+$, 100%).

**1-(4-(Difluoromethoxy)phenyl)-6-(4-fluoro-2-(trifluoromethyl)phenyl)-1*H*-imidazo[4,5-*c*]pyridine (OXS008316)**

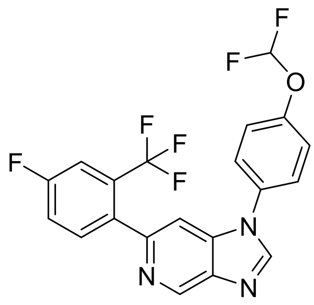

General procedure E.2 from **33** (150 mg, 0.507 mmol); yield 40% (86 mg, 0.20 mmol); brown solid. $^{1}$H NMR (400 MHz, $CDCl_3$) δ 9.24 (d, *J* = 1.2 Hz, 1H), 8.20 (s, 1H), 7.58 – 7.47 (m, 5H), 7.39 – 7.29 (m, 3H), 6.59 (t, *J* = 72.8 Hz, 1H); $^{13}$C NMR (100 MHz, $CDCl_3$) δ 162.1 (d, *J* = 250.0 Hz), 151.0 (t, *J* = 3 Hz), 150.7, 144.2, 143.1, 140.5, 138.8, 136.8 (q, *J* = 2 Hz), 134.3 (d, *J* = 8 Hz), 132.5, 130.8 – 130.2 (m), 125.6, 121.8, 118.7 (d, *J* = 21 Hz), 115.5 (t, *J* = 263 Hz), 114.0 (dq, J = 25, 5 Hz), 106.2 (d, *J* = 3 Hz).; *m/z* (ESI$^{+}$) 424.0 ([M+H]$^{+}$, 100%).

**1-(4-(Difluoromethoxy)phenyl)-6-(2-(trifluoromethoxy)phenyl)-1*H*-imidazo[4,5-*c*]pyridine (OXS008297)**

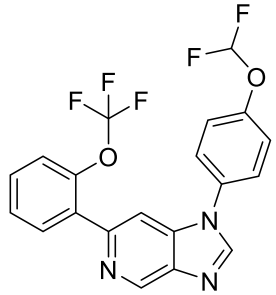

General procedure E.2 from **33** (150 mg, 0.507 mmol); yield 87% (186 mg, 0.441 mmol); pale brown solid. $^{1}$H NMR (400 MHz, $CDCl_3$) δ 9.29 (d, *J* = 0.8 Hz, 1H), 8.18 (s, 1H), 7.97 – 7.95 (m, 1H), 7.84 (d, *J* = 1.2 Hz, 1H), 7.55 – 7.52 (m, 2H), 7.45 – 7.34 (m, 5H), 6.61 (t, *J*= 72.8 Hz, 1H); $^{13}$C NMR (100 MHz, $CDCl_3$) δ 151.0 (t, *J* = 3 Hz), 148.3, 146.4 (q, *J* = 2 Hz), 144.2, 143.5, 140.5, 139.0, 134.0, 132.6, 132.3, 129.7, 127.4, 125.6, 121.8, 121.4 (q, *J* = 2 Hz), 119.3, 115.6 (t, *J* = 262 Hz), 106.6; *m/z* (ESI$^{+}$) 422.0 ([M+H]$^{+}$, 100%).

**1-(4-(Difluoromethoxy)phenyl)-6-(2-ethylphenyl)-1*H*-imidazo[4,5-*c*]pyridine (OXS007928)**

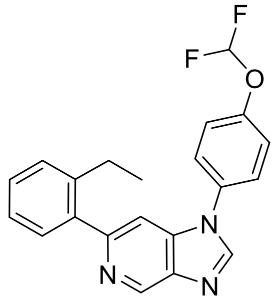

General procedure E.1 from **33** (150 mg, 0.507 mmol); yield 78% (144 mg, 0.394 mmol). $^1$H NMR (400 MHz, $CDCl_3$) δ 9.25 (d, *J* = 0.8 Hz, 1H), 8.17 (s, 1H), 7.55 – 7.49 (m, 3H), 7.37 – 7.32 (m, 5H), 7.27 – 7.23 (m, 1H), 6.59 (t, *J* = 72.8 Hz, 1H), 2.73 (q, *J* = 7.6 Hz, 2H), 1.10 (t, *J* = 7.6 Hz, 3H). $^{13}$C NMR (100 MHz, $CDCl_3$) δ 154.7, 150.7 (t, *J* = 3 Hz), 143.5, 142.6, 142.3, 140.5, 139.8, 139.0, 132.6, 130.0, 129.0, 128.4, 125.7, 125.5, 121.7, 115.4 (t, *J* = 263 Hz), 105.5, 26.2, 15.5; *m/z* (ESI$^+$) 366.1 ($[M+H]^+$, 100%).

**2-(1-(4-(Difluoromethoxy)phenyl)-1*H*-imidazo[4,5-*c*]pyridin-6-yl)-5-fluorophenol (70)**

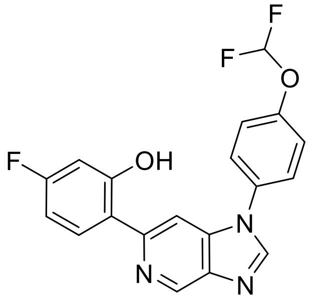

General procedure E.2 from **33** (150 mg, 0.507 mmol); yield 28% (53 mg, 0.14 mmol); light pink solid. $^1$H NMR (400 MHz, DMSO-$d_6$) 14.8 (s, 1H), 9.15 (s, 1H), 8.79 (s, 1H), 8.22 (s, 1H), 8.15 – 8.11 (m, 1H), 7.86 (dd, *J* = 2, 8.8 Hz, 2H), 7.48 (d, *J* = 8.8 Hz, 2H), 7.18 (t, *J* = 73.6 Hz, 1H), 6.74 – 6.68 (m, 2H); $^{13}$C NMR (100 MHz, DMSO-$d_6$) δ 163.2 (d, *J* = 246 Hz), 160.3 (d, *J* = 13 Hz), 150.4 (t, *J* = 3 Hz), 150.0, 146.7, 139.9, 139.2, 138.7, 131.7, 128.9 (d, *J* = 11 Hz), 126.1, 120.2,

116.5 (d, $J$ = 3 Hz), 116.1 (t, $J$ = 259 Hz), 105.7 (d, $J$ = 22 Hz), 104.0 (d, $J$ = 23 Hz), 101.6; *m/z* ($ESI^+$) 372.1 ($[M+H]^+$, 100%).

**1-(4-(Difluoromethoxy)phenyl)-6-(4-fluoro-2-propoxyphenyl)-1*H*-imidazo[4,5-*c*]pyridine (OXS008318)**

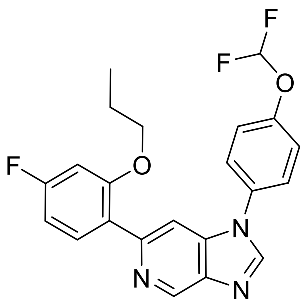

To phenol **70** (100 mg, 0.269 mmol) in acetonitrile (2.0 mL) was added $K_2CO_3$ (112 mg, 0.808 mmol) and 1-bromopropane (99 mg, 0.81 mmol). The reaction mixture was heated at 60 °C overnight, then cooled, quenched with water and extracted with EtOAc. The organics were dried with $MgSO_4$ and evaporated to dryness. The crude residue was purified by column chromatography (0-80% EtOAc in hexane) to give propoxy **OXS008318** (36 mg, 0.087 mmol, 32%) as an off-white solid. $^1H$ NMR (400 MHz, $CDCl_3$) δ 9.24 (d, $J$ = 0.8 Hz, 1H), 8.13 (s, 1H), 8.11 (d, $J$ = 1.2 Hz, 1H), 8.04 – 8.00 (m, 1H), 7.55 – 7.52 (m, 2H), 7.37 (d, $J$ = 8.8 Hz, 2H), 8.61 – 6.77 (m, 1H), 6.70 (dd, $J$ = 2.4, 10.8 Hz, 1H), 6.61 (t, $J$ = 72.8 Hz, 1H), 3.94 (t, $J$ = 6.3 Hz, 1H), 1.73 (sext, $J$ = 7.3 Hz, 1H), 0.88 (t, $J$ = 7.4 Hz, 1H).; $^{13}C$ NMR (100 MHz, $CDCl_3$) δ 163.8 (d, J = 247 Hz), 157.6 (d, $J$ = 10 Hz), 150.8 (t, $J$ = 3 Hz), 149.1, 143.7, 142.9, 139.6 (d, $J$ = 81 Hz), 132.9 (d, $J$ = 3 Hz), 132.8, 125.9, 125.1 (d, $J$ = 3 Hz), 121.8, 115.6 (t, $J$ = 263 Hz), 107.7 (d, $J$ = 21 Hz), 106.4, 100.3 (d, $J$ = 26 Hz), 70.4, 22.7, 10.8.; *m/z* ($ESI^+$) 414.1 ($[M+H]^+$, 100%).

**1-(4-(Difluoromethoxy)phenyl)-6-(4-fluoro-2-isopropoxyphenyl)-1*H*-imidazo[4,5-*c*]pyridine (OXS008317)**

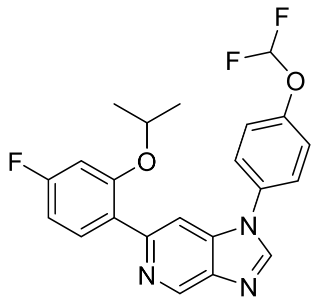

General procedure E.2 from **33** (150 mg, 0.507 mmol); yield 91% (190 mg, 0.460 mmol); brown oil. $^{1}$H NMR (400 MHz, $CDCl_3$) δ 9.24 (d, *J* = 1.2 Hz, 1H), 8.14 (s, 1H), 8.12 (d, *J* = 1.2 Hz, 1H), 8.02 (dd, *J* = 8.2, 8.4 Hz, 1H), 7.57 – 7.53 (m, 2H), 7.37 (d, J = 8.8 Hz, 2H), 6.78 (td, *J* = 2.4, 8.8 Hz, 1H), 6.71 (dd, *J* = 2.4, 10.8 Hz, 1H), 6.60 (t, *J* = 72.8 Hz, 1H), 4.56 (hept, *J* = 6.0 Hz, 1H), 1.29 (d, *J* = 6.0 Hz, 6H); $^{13}$C NMR (100 MHz, $CDCl_3$) δ 163.6 (d, *J* = 247 Hz), 156.2 (d, *J* = 10 Hz), 150.7 (t, *J* = 3 Hz), 149.2, 143.6, 142.9, 140.0, 139, 133.1, 132.4 (d, *J* = 5 Hz), 126.0 (d, *J* = 3 Hz), 125.6, 121.7, 115.6 (t, *J* = 263 Hz), 107.9 (d, *J* = 21 Hz), 106.5, 101.9 (d, *J* = 25 Hz), 71.2, 22.1; *m/z* ($ESI^+$) 414.1 ($[M+H]^+$, 100%).

**1-(4-(Difluoromethoxy)phenyl)-6-(2,4-dimethylphenyl)-1*H*-imidazo[4,5-*c*]pyridine (OXS007929)**

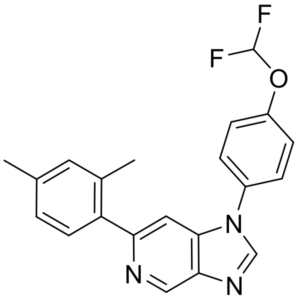

General procedure E.1 from **33** (150 mg, 0.507 mmol), yield 59% (109 mg, 0.299 mmol); off-white solid. $^{1}$H NMR (400 MHz, $CDCl_3$) δ 9.25 (d, *J* = 0.8 Hz, 1H), 8.15 (s, 1H), 7.55 – 7.51 (m, 2H), 7.48 (d, *J* = 1.2 Hz, 1H), 7.38 – 7.35 (m, 2H), 7.30 (d, *J* = 7.6 Hz, 1H), 7.11 – 7.06 (m, 2H), 6.59 (t, *J* = 72.8 Hz, 1H), 2.36 (s, 3H), 2.34 (s, 3H). $^{13}$C NMR (100 MHz, $CDCl_3$) δ 154.8, 150.8

(t, $J$ = 3 Hz), 143.6, 142.8, 139.8, 139.2, 138.2, 138.0, 136.0, 132.8, 131.7, 130.0, 126.7, 125.6, 121.8, 115.6 (t, $J$ = 263 Hz), 105.6, 21.3, 20.5; *m/z* (ESI$^+$) 366.1 ([M+H]$^+$, 100%).

**4-(1-(4-(Difluoromethoxy)phenyl)-1*H*-imidazo[4,5-*c*]pyridin-6-yl)-3-methylbenzonitrile (OXS007556)**

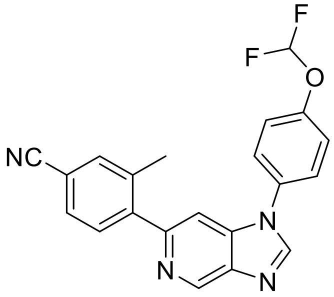

General procedure E.2 from **33** (92 mg, 0.31 mmol); yield 66% (77 mg, 0.20 mmol); pale brown foam. $^1$H NMR (500 MHz, $CDCl_3$) δ 9.27 (d, $J$ = 1.1 Hz, 1H), 8.21 (s, 1H), 7.61 – 7.45 (m, 6H), 7.38 (d, $J$ = 2.2 Hz, 1H), 6.61 (t, $J$ = 72.7 Hz, 1H), 2.40 (s, 3H); $^{13}$C NMR (126 MHz, $CDCl_3$) δ 152.4, 151.0 (t, $J$ = 3 Hz), 145.3, 144.3, 143.2, 140.3, 139.2, 137.9, 134.4, 132.4, 130.8, 129.7, 125.8, 122.0, 119.0, 115.5 (t, $J$ = 263 Hz), 112.0, 105.9, 20.5; *m/z* (ESI$^+$) 377.1 ([M+H]$^+$, 100%); HRMS (ESI) $C_{21}H_{15}ON_4F$([M+H]$^+$) requires 377.12121; found 377.12084.

**6-(4-Chloro-2-methylphenyl)-1-(4-(difluoromethoxy)phenyl)-1*H*-imidazo[4,5-*c*]pyridine (OXS007930)**

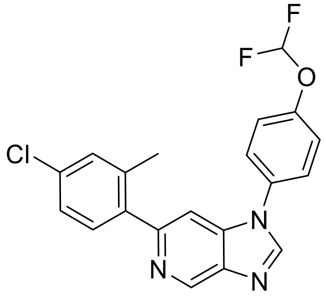

General procedure E.1 from **33** (150 mg, 0.507 mmol); yield 39%, (76 mg, 0.20 mmol); off-white solid. $^1$H NMR (400 MHz, $CDCl_3$) δ 9.25 (d, *J* = 0.8 Hz, 1H), 8.17 (s, 1H), 7.54 – 7.51 (m, 2H), 7.46 (d, *J* = 0.8 Hz, 1H), 7.40 – 7.22 (m, 5H), 6.60 (t, *J* = 72.8 Hz, 1H), 2.35 (s, 3H).$^{13}$C NMR (100 MHz, $CDCl_3$) δ 153.5, 151.0 (t, *J* = 3 Hz), 143.9, 143.0, 140.0, 139.4, 139.3, 138.2, 134.0, 132.7, 131.3, 130.7, 126.1, 125.7, 121.9, 115.5 (t, *J* = 263.0 Hz), 105.7, 20.5; *m/z* (ESI$^+$) 386.0 ([M+H]$^+$, 100%).

**1-(4-(Difluoromethoxy)phenyl)-6-(2-methyl-4-(trifluoromethyl)phenyl)-1*H*-imidazo[4,5-*c*]pyridine (OXS008161)**

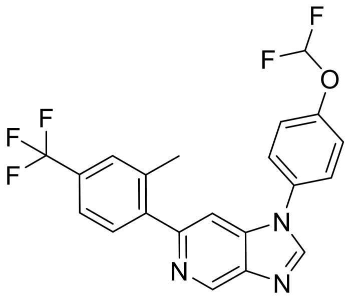

General procedure E.2 from **33** (150 mg, 0.507 mmol); yield 72%, (153 mg, 0.364 mmol); beige solid. $^1$H NMR (400 MHz, $CDCl_3$) δ 9.27 (d, *J* = 1.2 Hz, 1H), 8.20 (s, 1H), 7.55 – 7.50 (m, 6H), 7.39 – 7.37 (m, 2H), 6.60 (t, *J* = 72.8 Hz, 1H), 2.42 (s, 3H); $^{13}$C NMR ($CDCl_3$, 100 MHz) δ 153.2, 151.0 (t, *J* = 3 Hz), 144.3, 144.1, 143.2, 140.2, 139.3, 137.3, 132.6, 130.4, 127.6 (q, *J* = 4 Hz), 125.7, 122.8 (q, *J* = 4 Hz), 121.9, 115.5 (t, *J* = 263 Hz), 105.8, 20.6; *m/z* (ESI$^+$) 420.1 ([M+H]$^+$, 100%).

**4-{1-[4-(Difluoromethoxy)phenyl]-1*H*-imidazo[4,5-*c*]pyridin-6-yl}-3-methylphenol (OXS008187)**

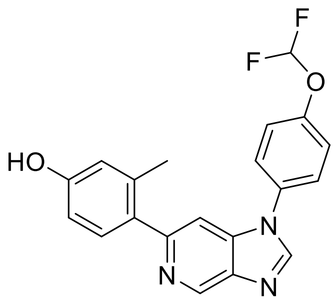

General procedure E.1 from **33** (150 mg, 0.507 mmol); yield 16% (30 mg, 0.082 mmol); beige solid. $^{1}$H NMR (400 MHz, DMSO-$d_6$) δ 9.32 (s, 1H), 9.08 (d, *J* = 0.8 Hz, 1H), 8.68 (s, 1H), 7.83 – 7.80 (m, 2H), 7.54 (d, *J* = 1.2 Hz, 1H), 7.45 – 7.42 (m, 2H), 7.32 (t, *J* = 74 Hz, 1H), 7.26 (d, *J* = 8.4 Hz, 1H), 6.69 (d, *J* = 2.0, 1H), 6.66 (dd, *J* = 8.4, 2.4 Hz, 1H), 2.28 (s, 3H); $^{13}$C NMR (100 MHz, DMSO-$d_6$) δ 156.9, 153.4, 150.2 (t, *J* = 3 Hz), 145.0, 141.1, 139.2, 138.4, 137.0, 132.2, 131.8, 131.2, 125.6, 120.3, 117.1, 116.2 (t. *J* = 259 Hz), 112.6, 105.3, 20.5; *m/z* (ESI$^{+}$) 368.1 ([M+H]$^{+}$, 100%).

**6-(4,5-Difluoro-2-methylphenyl)-1-(4-(difluoromethoxy)phenyl)-1*H*-imidazo[4,5-*c*]pyridine (OXS007733)**

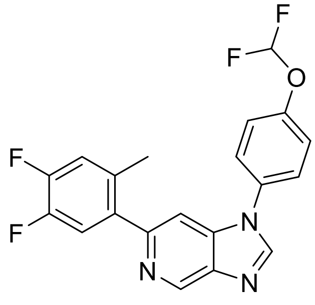

General procedure E.2 from **33** (300 mg, 1.06 mmol); yield 15% (58 mg, 0.15 mmol); grey solid. $^{1}$H NMR (400 MHz, $CDCl_3$) δ 9.25 (d, *J* = 1.0 Hz, 1H), 8.18 (s, 1H), 7.56 – 7.50 (m, 2H), 7.45 (d, *J* = 1.0 Hz, 1H), 7.43 – 7.34 (m, 2H), 7.24 (dd, *J* = 11.0, 8.1 Hz, 1H), 7.08 (dd, *J* = 11.3, 7.9 Hz, 1H), 6.61 (t, *J* = 72.8 Hz, 1H), 2.31 (s, 3H); $^{13}$C NMR (101 MHz, $CDCl_3$) δ 152.4 – 152.2 (m), 150.9 (t, *J* = 2.9 Hz), 150.2 (dd, *J* = 154.9, 12.7 Hz), 148.0 (dd, *J* = 153.1, 12.4 Hz), 144.0, 143.0, 140.0, 139.1, 137.2 (dd, *J* = 5.4, 3.7 Hz), 133.0 (dd, *J* = 6.0, 3.8 Hz), 132.4, 125.6, 121.8, 119.2

(d, $J$ = 16.9 Hz), 118.7 (dd, $J$ = 17.5, 1.0 Hz), 115.4 (t, $J$ = 263.0 Hz), 105.6, 19.9 (d, $J$ = 1.1 Hz).*m/z* ($ESI^+$) 388.0 ($[M+H]^+$, 100%).

**1-(4-(Difluoromethoxy)phenyl)-6-(6-fluoro-2-methylpyridin-3-yl)-1*H*-imidazo[4,5-*c*]pyridine (OXS007865)**

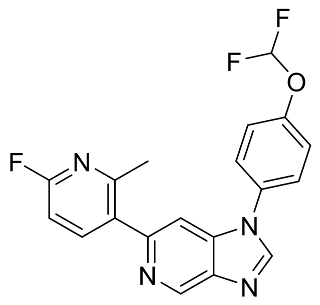

General procedure E.2 from **33** (150 mg, 0.507 mmol); yield 26% (49 mg, 0.13 mmol); brown solid. $^1$H NMR (400 MHz, DMSO-$d_6$) δ 9.16 (d, $J$ = 1.0 Hz, 1H), 8.78 (s, 1H), 8.06 (t, $J$ = 8.3 Hz, 1H), 7.89 – 7.81 (m, 2H), 7.78 (d, $J$ = 1.0 Hz, 1H), 7.47 – 7.41 (m, 3H), 7.34 (t, $J$ = 73.7 Hz, 1H), 7.10 – 7.05 (m, 2H), 2.49 (s, 3H).$^{13}$C NMR (101 MHz, DMSO-$d_6$) δ 161.9 (d, $J$ = 235.2 Hz), 155.5 (d, $J$ = 14.7 Hz), 150.8 (t, $J$ = 3.4 Hz), 150.5 (d, $J$ = 1.5 Hz), 146.2, 144.0 (d, $J$ = 8.0 Hz), 142.2, 140.4, 139.0, 134.5 (d, $J$ = 4.7 Hz), 132.5, 126.2, 120.8, 116.7 (t, $J$ = 258.8 Hz), 107.1, 106.7 (d, $J$ = 37.7 Hz), 23.3 (d, $J$ = 1.0 Hz); *m/z* ($ESI^+$) 371.1 ($[M+H]^+$, 100%).

**1-(4-(Difluoromethoxy)phenyl)-6-(6-fluoro-4-methylpyridin-3-yl)-1*H*-imidazo[4,5-*c*]pyridine (OXS007864)**

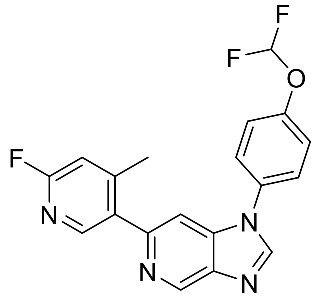

General procedure E.2 from **33** (150 mg, 0.507 mmol); yield 47% (87 mg, 0.24 mmol); off-white solid. $^{1}$H NMR (400 MHz, DMSO-$d_6$) δ 9.17 (d, *J* = 1.0 Hz, 1H), 8.78 (s, 1H), 8.29 (s, 1H), 7.89 – 7.84 (m, 2H), 7.81 (d, *J* = 1.0 Hz, 1H), 7.46 – 7.42 (m, 2H), 7.34, (t, *J* = 73.8 Hz, 1H), 7.16 (s, 1H), 2.42 (s, 3H);$^{13}$C NMR (101 MHz, DMSO-$d_6$) δ 163.1 (d, *J* = 235.6 Hz), 152.5 (d, *J* = 8.5 Hz), 150.8 (t, *J* = 3.3 Hz), 149.2 (d, *J* = 1.2 Hz), 148.1 (d, *J* = 16.0 Hz), 146.1, 142.3, 140.4, 139.0, 135.7 (d, *J* = 4.3 Hz), 132.6, 126.2, 120.8, 116.7 (t, *J* = 258.6 Hz), 110.6 (d, *J* = 37.1 Hz), 107.2, 20.4 (d, *J* = 2.7 Hz); *m/z* (ESI$^{+}$) 371.1 ([M+H]$^{+}$, 100%).

**6-Chloro-1-(4-(difluoromethoxy)phenyl)-2-methyl-1*H*-imidazo[4,5-*c*]pyridine (71)**

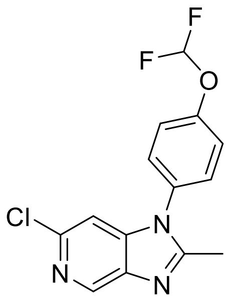

General procedure C from **32** (502 mg, 1.76 mmol), using triethylorthoacetate instead of triethylorthoformate; yield 87% (473 mg, 1.53 mmol); pale brown solid. $^{1}$H NMR (400 MHz, $CDCl_3$) δ 8.76 (d, *J* = 0.9 Hz, 1H), 7.37 (d, *J* = 0.9 Hz, 4H), 7.06 (d, *J* = 0.9 Hz, 1H), 6.85 – 6.39 (m, 1H), 2.51 (s, 3H); *m/z* (ESI$^{+}$) 310.0 ([M+H]$^{+}$, 100%).

**1-(4-(Difluoromethoxy)phenyl)-6-(4-fluoro-2-methylphenyl)-2-methyl-1*H*-imidazo[4,5-*c*]pyridine (OXS007752)**

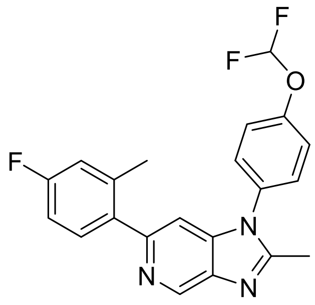

General procedure E.2 from **71** (96 mg, 0.31 mmol); yield 46% (55 mg, 0.14 mmol); brown solid. $^{1}$H NMR (600 MHz, $CDCl_3$) δ 9.09 (d, *J* = 1.0 Hz, 1H), 7.43 – 7.35 (m, 4H), 7.31 (dd, *J* = 8.5, 5.9 Hz, 1H), 7.09 (d, *J* = 1.0 Hz, 1H), 6.96 (dd, *J* = 9.8, 2.7 Hz, 1H), 6.91 (td, *J* = 8.4, 2.7 Hz, 1H), 6.62 (t, *J* = 72.8 Hz, 1H), 2.56 (s, 3H), 2.32 (s, 3H); $^{13}$C NMR (151 MHz, $CDCl_3$) δ 162.5 (d, *J* = 246 Hz), 153.2 (d, *J* = 135 Hz), 151.5, 142.0, 141.0, 138.8, 138.7, 137.2, 132.2, 131.5 (d, *J* = 8 Hz),128.5, 121.5, 117.3 (d, *J* = 21 Hz), 115.5, 113.8, 112.6 (d, *J* = 21 Hz), 105.4, 20.7, 14.6; *m/z* (ESI$^{+}$) 384.1 ([M+H]$^{+}$, 100%); HRMS (ESI$^{+}$) $C_{21}H_{17}ON_3F_3$ ([M+H]$^{+}$) requires 384.1318; found 384.1819.

**1-[4-(Difluoromethoxy)phenyl]-6-(4-fluoro-2-methyl-phenyl)-2-phenyl-imidazo[4,5-*c*]pyridine (OXS007768)**

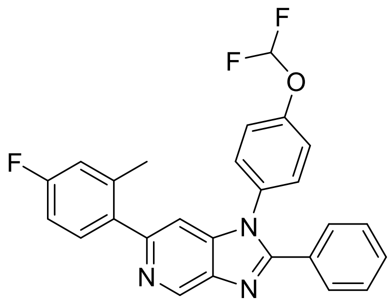

**OXS007417** (50 mg, 0.14 mmol), iodobenzene (16 µL, 0.14 mmol), $K_2CO_3$ (39 mg, 0.28 mmol), $PPh_3$ (18 mg, 0.069 mmol), copper acetate (5 mg, 0.03 mmol) and $Pd(PPh_3)_4$ (16 mg, 0.014 mmol) were dry mixed. Toluene (3 mL) was added and the reaction was heated under microwave irradiation to 100 °C for 48 h. Following this time, the mixture was diluted with $H_2O$ and extracted with $Et_2O$. The organics were combined, dried over $Na_2SO_4$ and concentrated *in vacuo*. The crude residue was purified by flash column chromatography (0 – 60% EtOAc in petroleum ether) to yield the desired product (10 mg, 0.022 mmol, 16%) as a white solid. $^{1}$H NMR (500 MHz, $CDCl_3$) δ 9.27 (s, 1H), 7.63 – 7.59 (m, 2H), 7.49 – 7.44 (m, 1H), 7.44 – 7.39 (m, 2H), 7.39 – 7.34 (m, 2H), 7.32 – 7.29 (m, 2H), 7.23 (d, *J* = 1.1 Hz, 1H), 7.01 (dd, *J* = 9.8, 2.7 Hz, 1H), 6.96 (td, *J* = 8.4, 2.7 Hz, 1H), 6.63 (t, *J* = 72.8 Hz, 2H), 2.38 (s, 3H). $^{13}$C NMR (126 MHz, $CDCl_3$) δ 162.5 (d, *J* = 248 Hz), 154.2, 153.1, 151.1 (t, *J* = 3 Hz), 142.4, 142.0, 138.9, 138.6 (d, *J* = 8 Hz), 137.1, 133.1, 131.4 (d, *J* = 3 Hz), 130.4, 129.5, 128.8, 128.7, 128.6, 121.1, 117.2 (d, *J* = 21 Hz), 115.4 (t, *J* = 262 Hz), 112.5 (d, *J* = 21 Hz), 105.6, 20.6. *m/z* ($ESI^+$) 446.1 ($[M+H]^+$, 100%); HRMS ($ESI^+$) $C_{26}H_{19}N_3OF^+$ ($[M+H]^+$) requires 446.1475; found 446.1478.

#### $^{1}H$ and $^{13}C$ NMR spectra of tested compounds

**_N_-(3-((1-(3,4-Dimethoxybenzyl)-1_H_-imidazo[4,5-_c_]pyridin-6-yl)amino)phenyl)acetamide (OXS000675)**

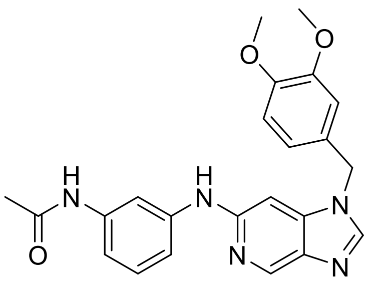

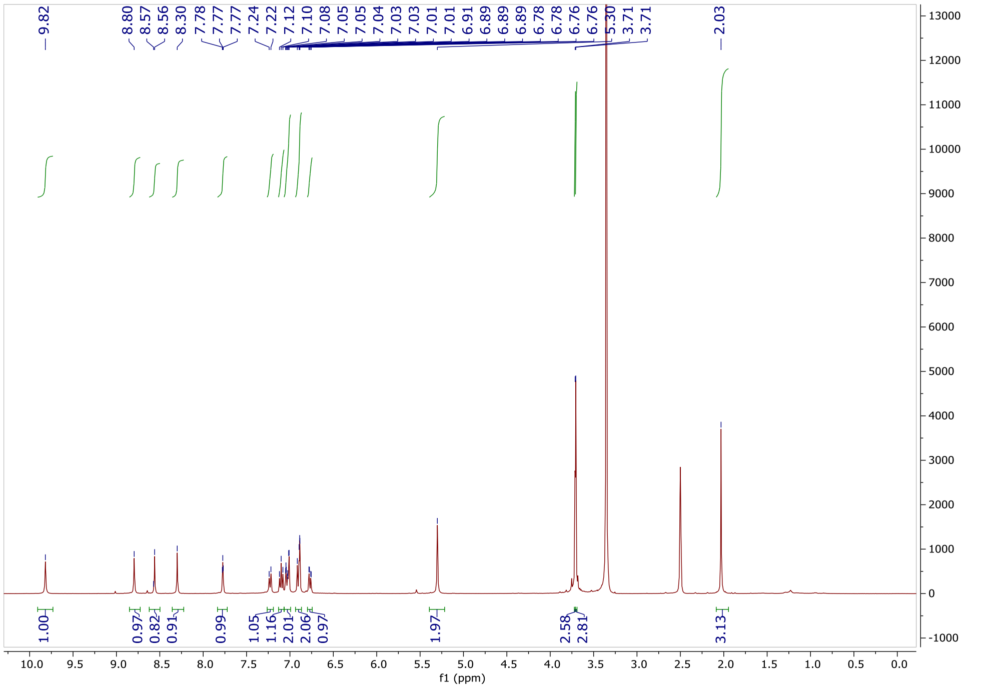
9.82
8.80
8.57
8.56
8.30
7.78
7.77
7.77
7.24
7.22
7.12
7.10
7.08
7.05
7.05
7.04
7.03
7.03
7.01
7.01
6.91
6.89
6.89
6.89
6.78
6.76
6.76
5.30
3.71
3.71
2.03
1.00
0.97
0.82
0.91
0.99
1.05
1.16
2.01
2.06
0.97
1.97
2.58
2.81
3.13
f1 (ppm)

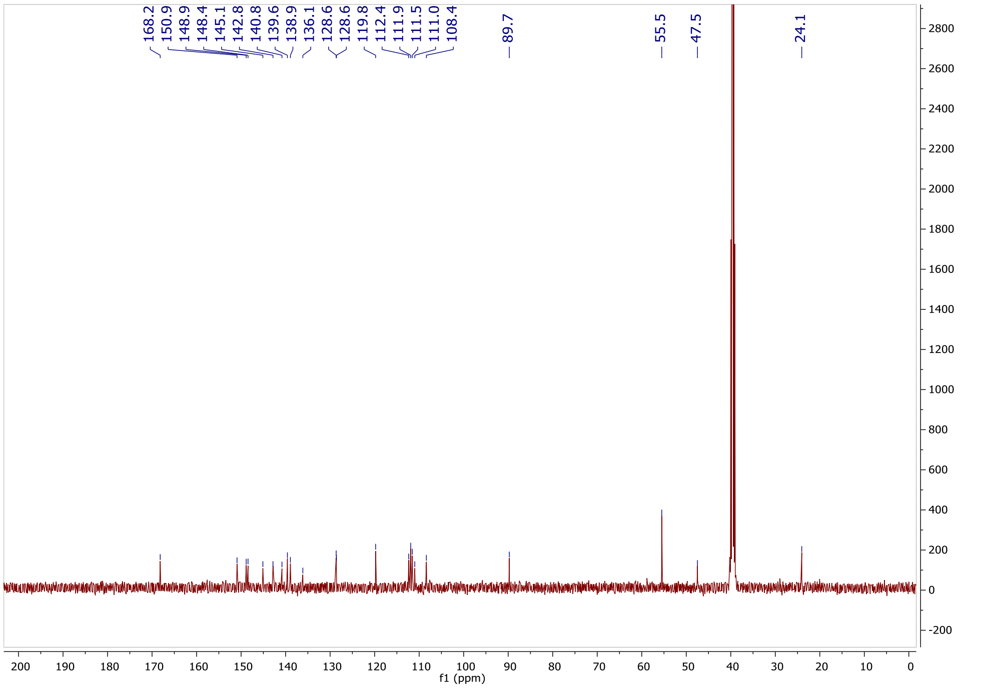
168.2
150.9
148.9
148.4
145.1
142.8
140.8
139.6
138.9
136.1
128.6
128.6
119.8
112.4
111.9
111.5
111.0
108.4
89.7
55.5
47.5
24.1
f1 (ppm)

***N*-(3-((1-(3,4-Dimethoxyphenyl)-1*H*-imidazo[4,5-*c*]pyridin-6-yl)amino)phenyl)acetamide (OXS007014)**

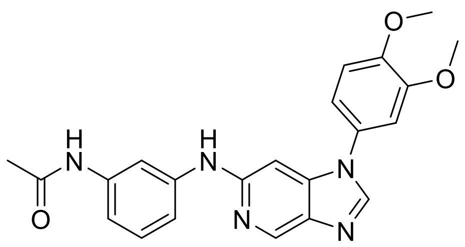

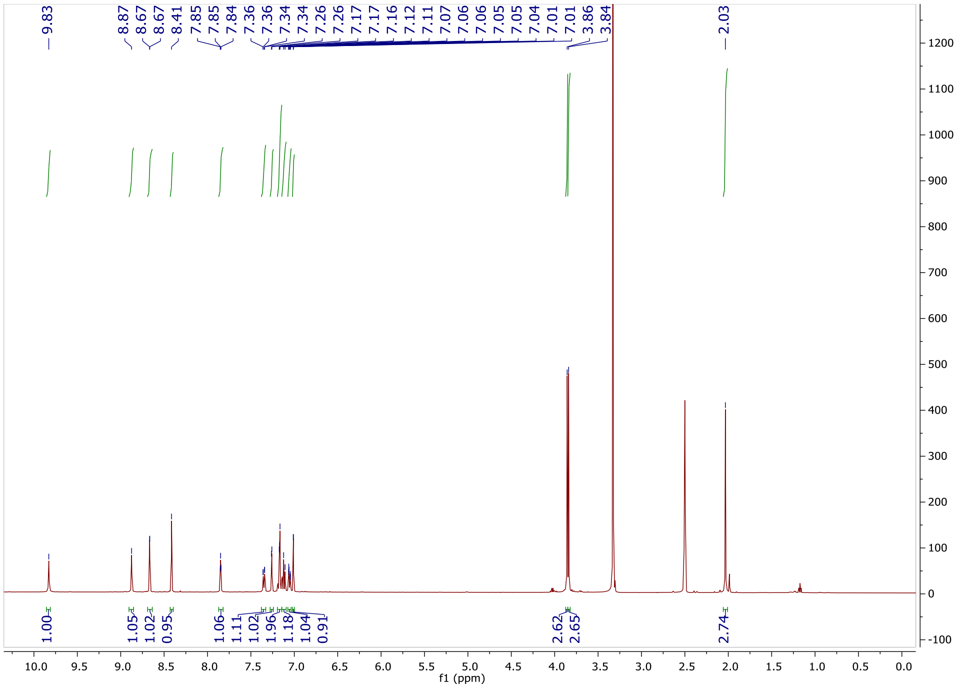
9.83
8.87
8.67
8.67
8.41
7.85
7.85
7.84
7.36
7.34
7.34
7.26
7.26
7.17
7.17
7.16
7.12
7.11
7.07
7.06
7.05
7.04
7.01
7.01
3.86
3.84
2.03
1.00
1.05
1.02
0.95
1.06
1.11
1.02
1.96
1.18
1.04
0.91
2.62
2.65
2.74
f1 (ppm)

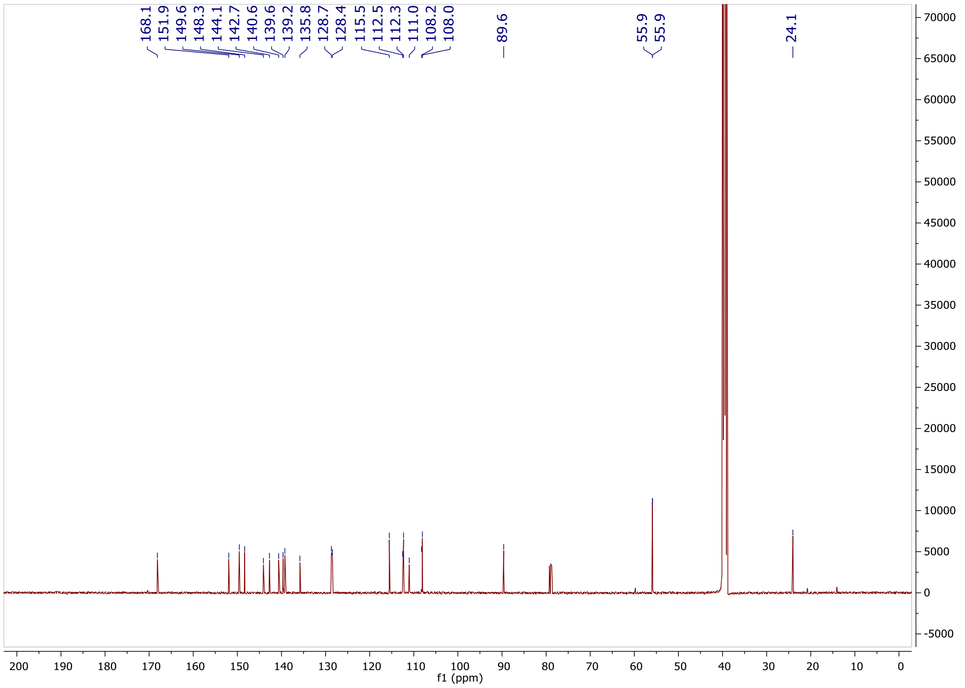
168.1
151.9
149.6
148.3
144.1
142.7
140.6
139.6
139.2
135.8
128.7
128.4
115.5
112.5
112.3
111.0
108.2
108.0
89.6
55.9
55.9
24.1
f1 (ppm)

***N*-[3-[[1-(3,4-Dimethoxyphenyl)imidazo[4,5-c]pyridin-6-yl]amino]phenyl]methanesulfonamide (OXS007746)**

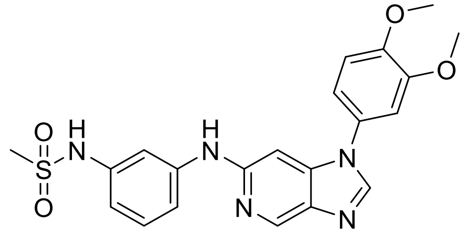

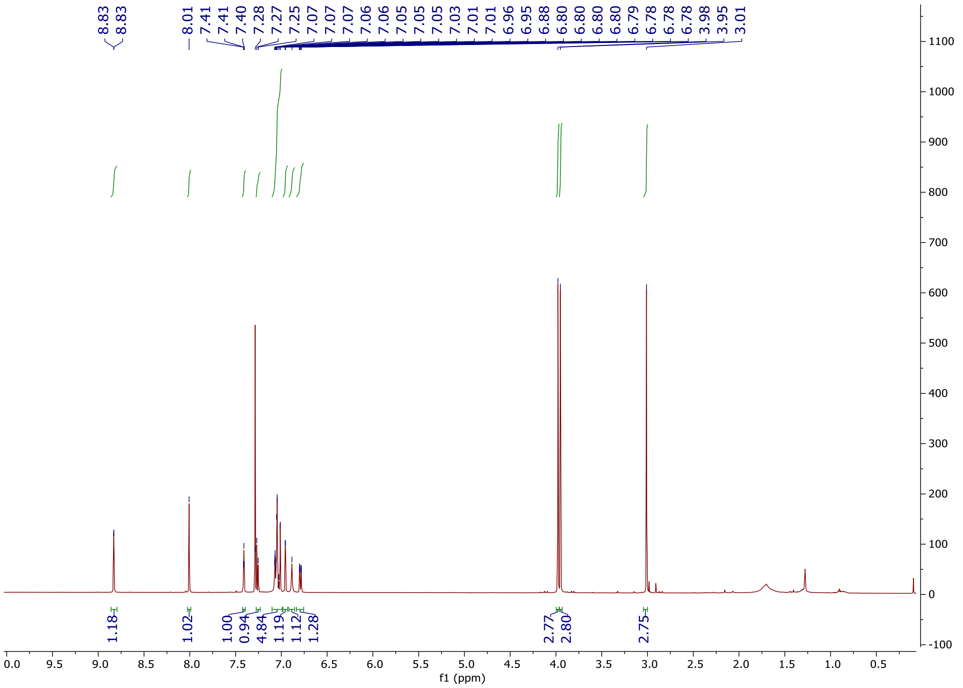

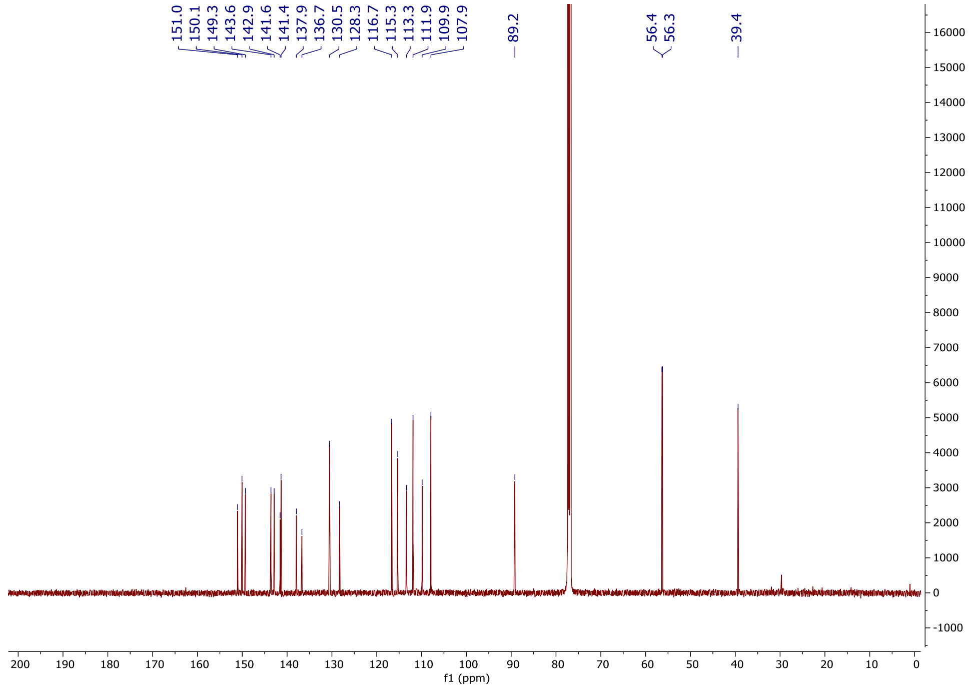

151.0
150.1
149.3
143.6
142.9
141.6
141.4
137.9
136.7
130.5
128.3
116.7
115.3
113.3
111.9
109.9
107.9
89.2
56.4
56.3
39.4
f1 (ppm)

#### *N*-(3-((1-(3,4-Dimethoxyphenyl)-1*H*-imidazo[4,5-*c*]pyridin-6-yl)oxy)phenyl)acetamide (OXS007266)

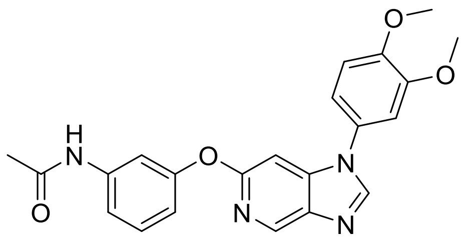

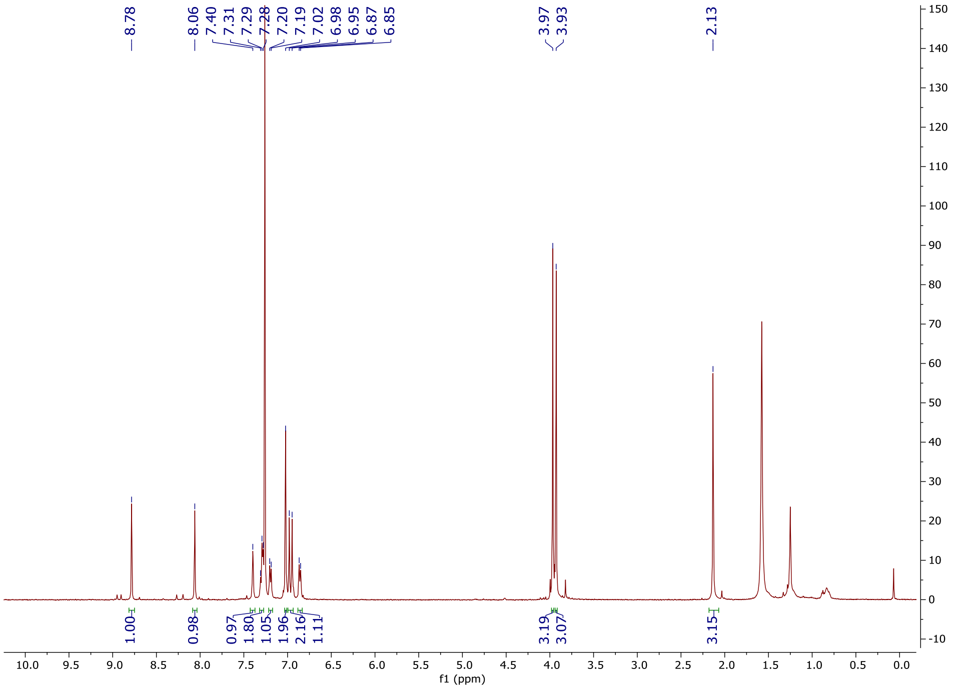

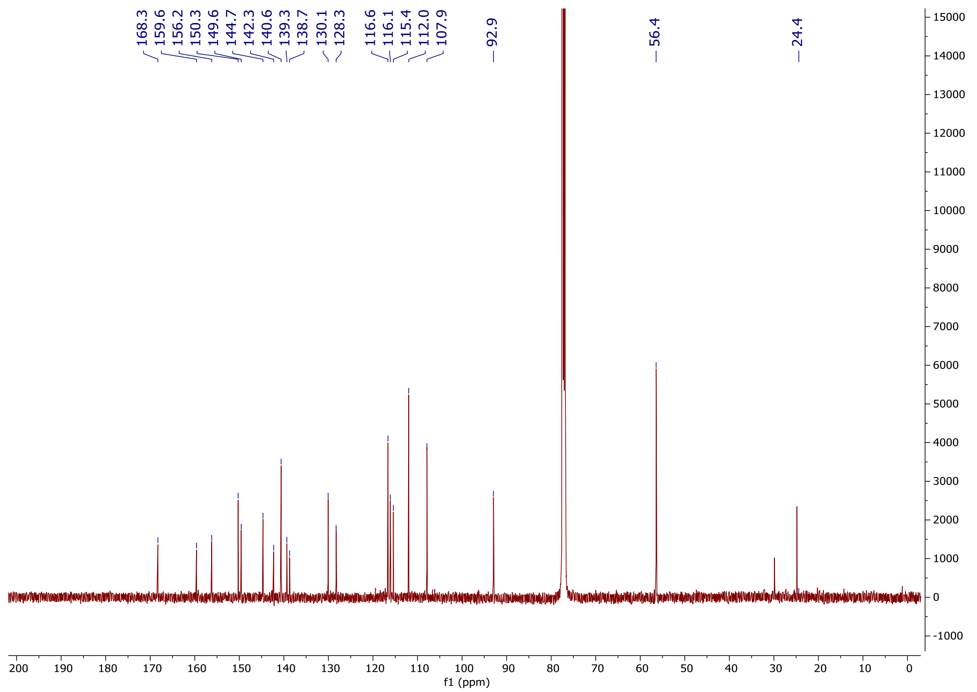

**N-(3-(1-(3,4-Dimethoxyphenyl)-1H-imidazo[4,5-c]pyridin-6-yl)phenyl)acetamide (OXS007034)**

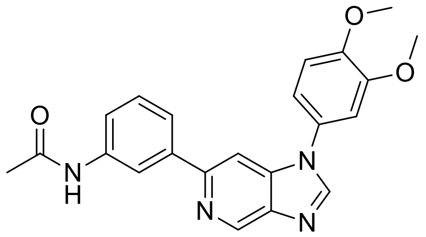

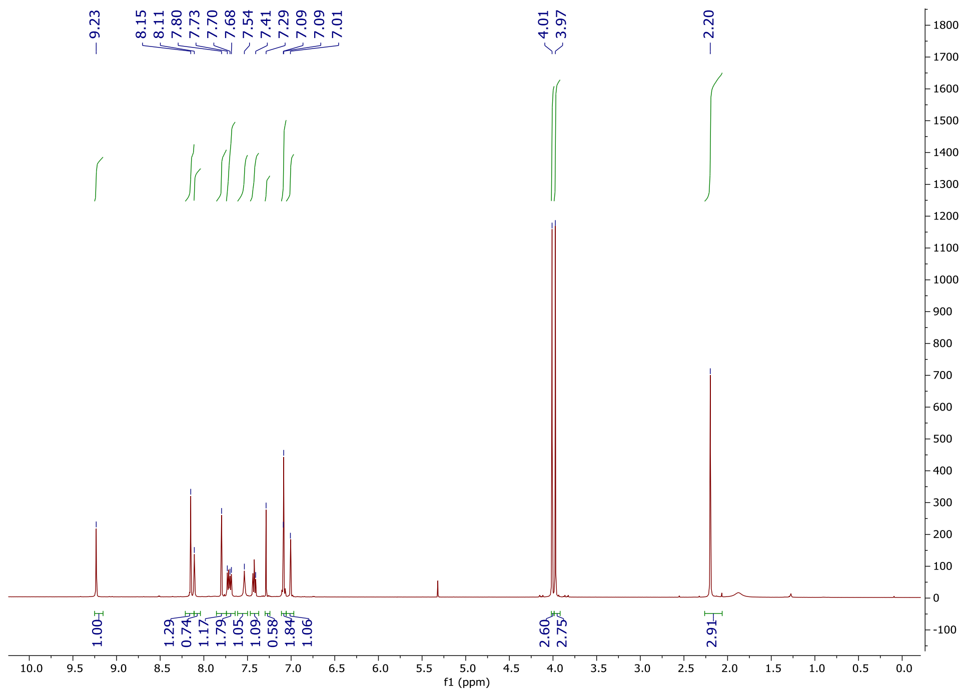
9.23
8.15
8.11
7.80
7.73
7.70
7.68
7.54
7.41
7.29
7.09
7.09
7.01
4.01
3.97
2.20
1.00
1.29
0.74
1.17
1.79
1.05
1.09
0.58
1.84
1.06
2.60
2.75
2.91
f1 (ppm)

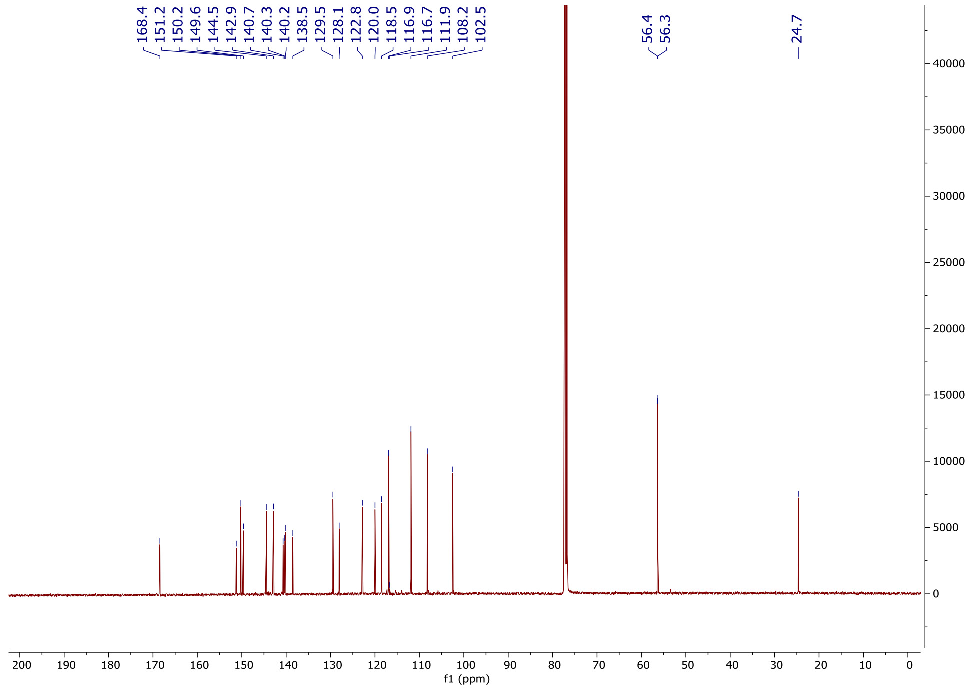
168.4
151.2
150.2
149.6
144.5
142.9
140.7
140.3
140.2
138.5
129.5
128.1
122.8
120.0
118.5
116.9
116.7
111.9
108.2
102.5
56.4
56.3
24.7
f1 (ppm)

### 1-(3,4-Dimethoxyphenyl)-6-phenyl-1*H*-imidazo[4,5-*c*]pyridine (OXS007136)

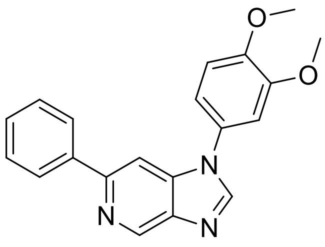

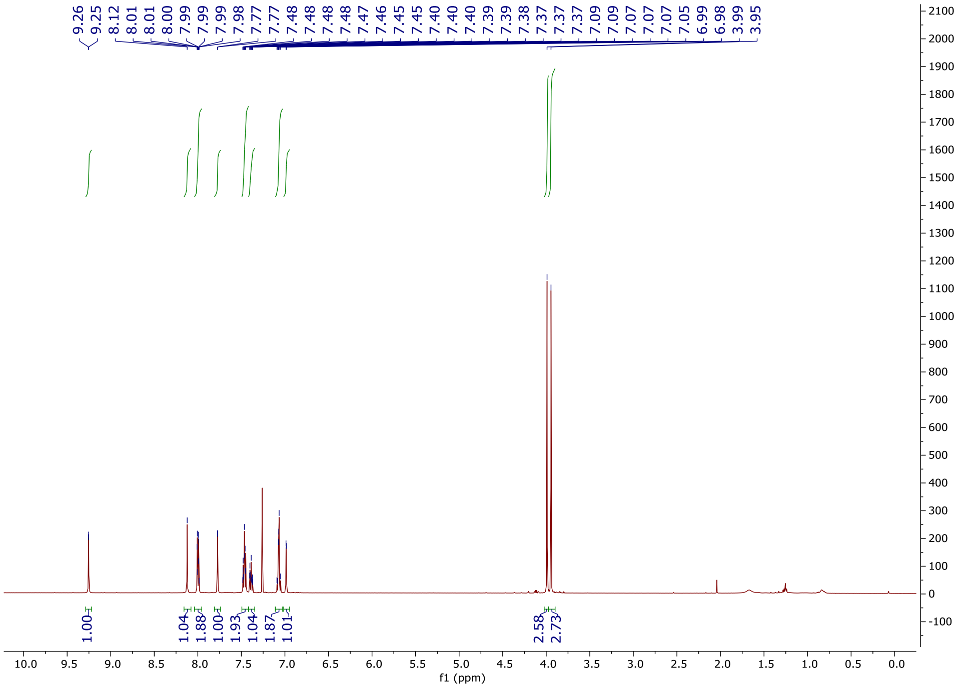

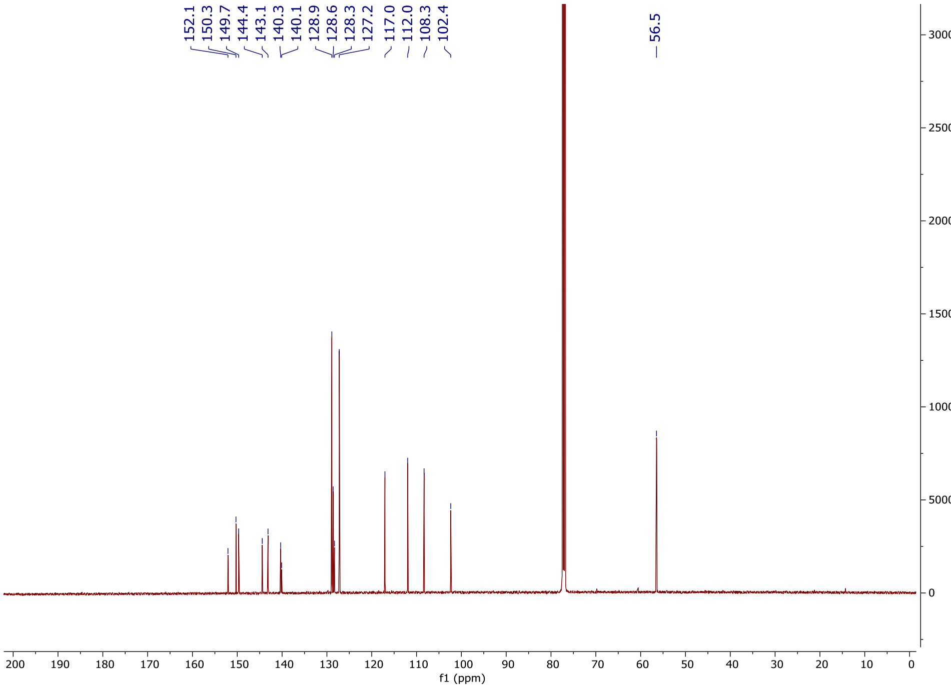
152.1
150.3
149.7
144.4
143.1
140.3
140.1
128.9
128.6
128.3
127.2
117.0
112.0
108.3
102.4
56.5
30000
25000
20000
15000
10000
5000
0
200 190 180 170 160 150 140 130 120 110 100 90 80 70 60 50 40 30 20 10 0
f1 (ppm)

### 1-(3,4-Dimethoxyphenyl)-6-(*o*-tolyl)-1*H*-imidazo[4,5-*c*]pyridine (OXS007142)

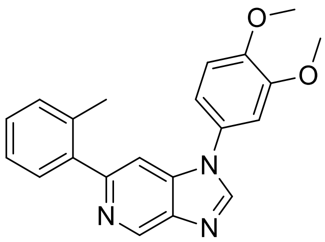

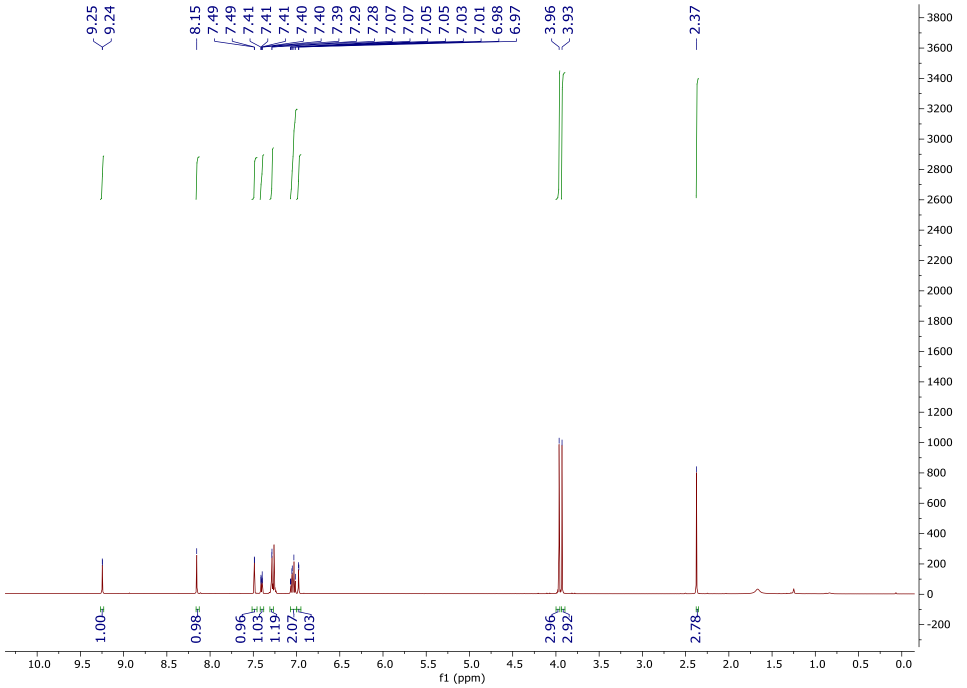

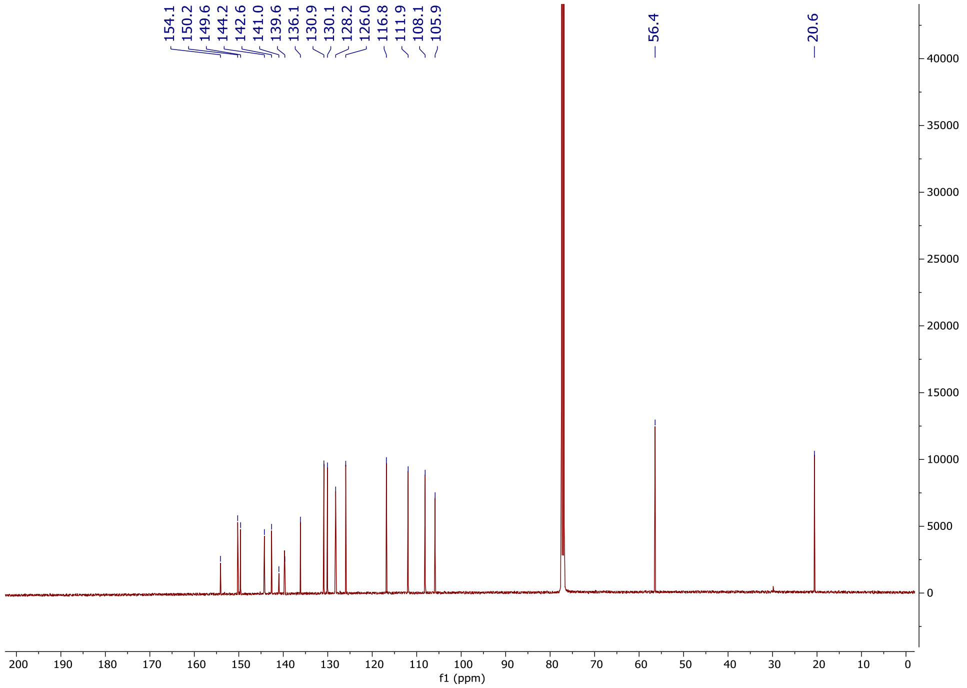

154.1
150.2
149.6
144.2
142.6
141.0
139.6
136.1
130.9
130.1
128.2
126.0
116.8
111.9
108.1
105.9
56.4
20.6
40000
35000
30000
25000
20000
15000
10000
5000
0
200 190 180 170 160 150 140 130 120 110 100 90 80 70 60 50 40 30 20 10 0
f1 (ppm)

**1-(3,4-Dimethoxyphenyl)-6-(2-methoxyphenyl)-1*H*-imidazo[4,5-*c*]pyridine (OXS006988)**

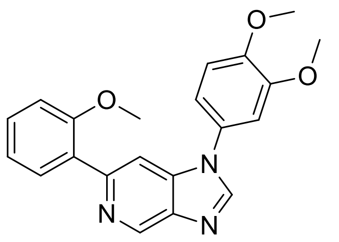

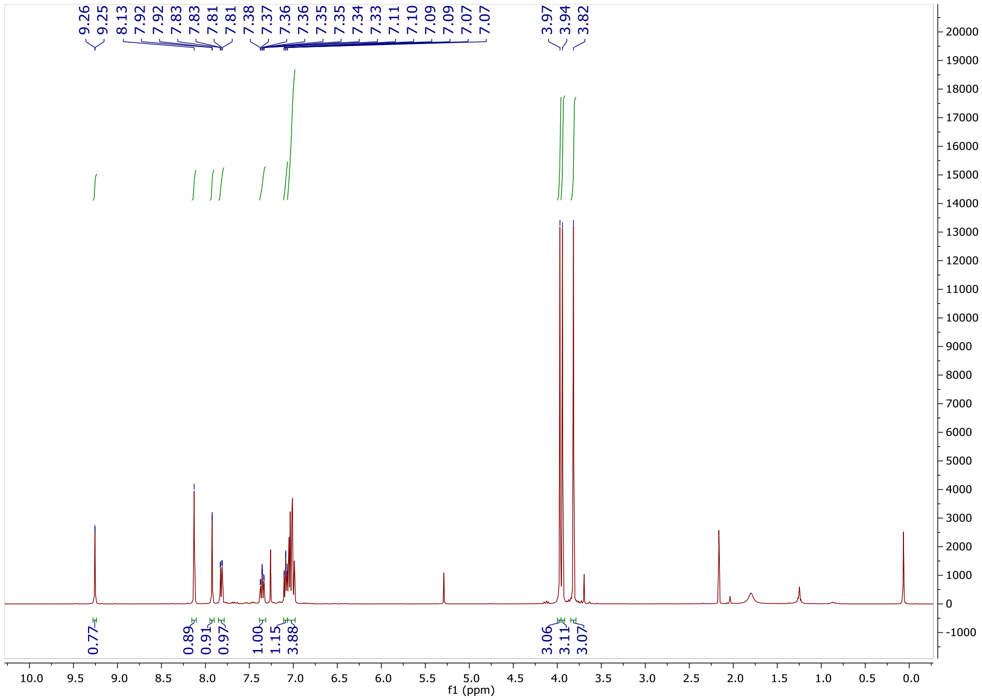
9.26
9.25
8.13
7.92
7.92
7.83
7.83
7.81
7.81
7.38
7.37
7.36
7.36
7.35
7.35
7.34
7.33
7.11
7.10
7.09
7.09
7.07
7.07
3.97
3.94
3.82
0.77
0.89
0.91
0.97
1.00
1.15
3.88
3.06
3.11
3.07
f1 (ppm)

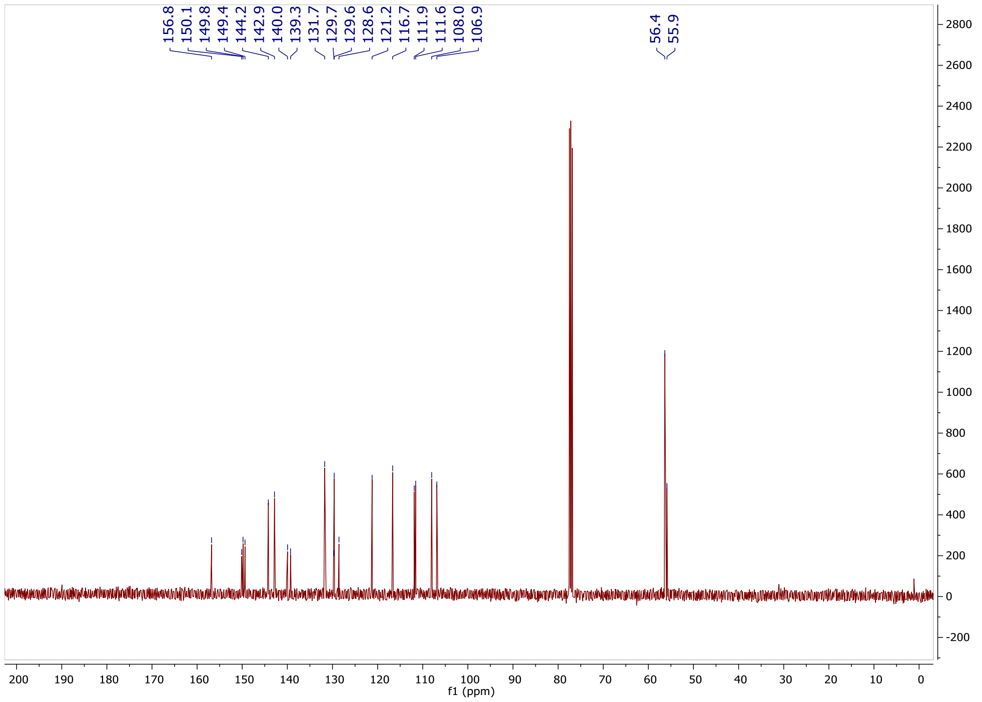
156.8
150.1
149.8
149.4
144.2
142.9
140.0
139.3
131.7
129.7
129.6
128.6
121.2
116.7
111.9
111.6
108.0
106.9
56.4
55.9
f1 (ppm)

#### 1-(3,4-Dimethoxyphenyl)-6-(2-(trifluoromethoxy)phenyl)-1*H*-imidazo[4,5-*c*]pyridine (OXS007316)

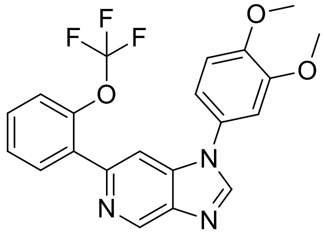

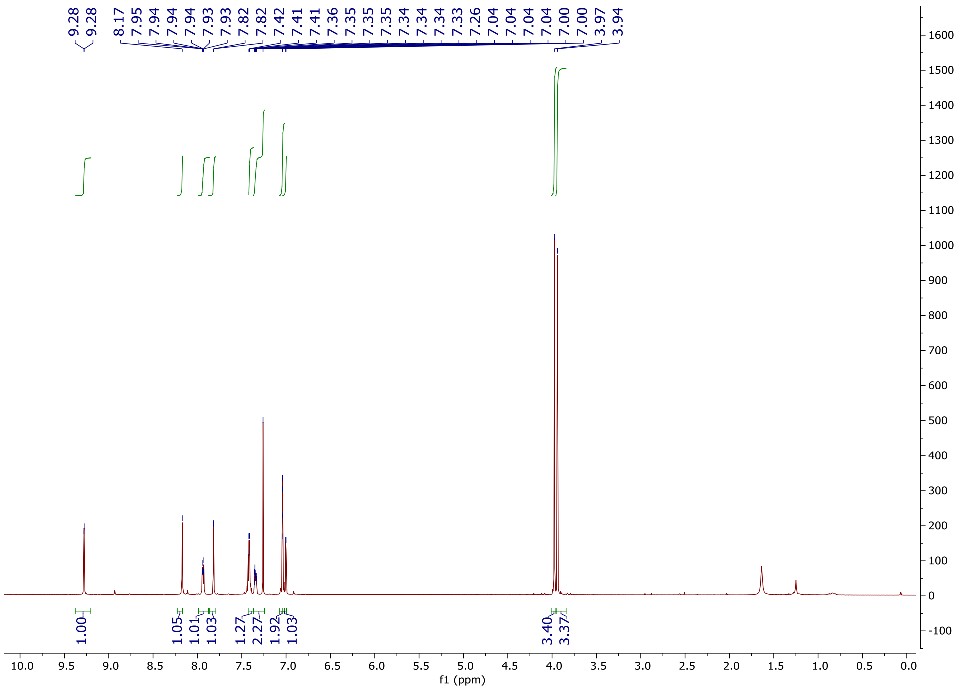

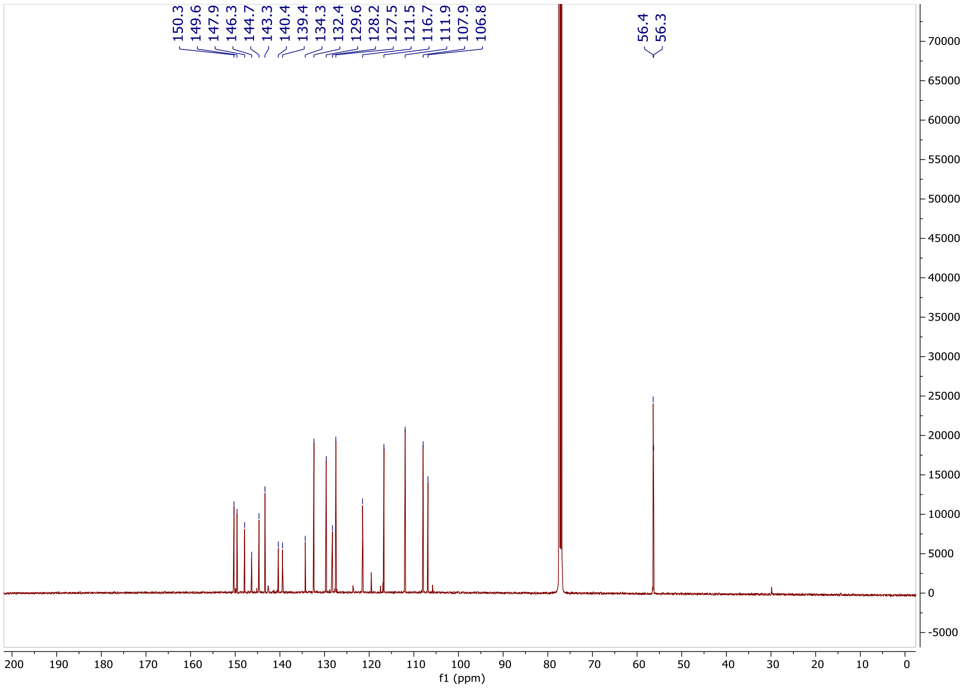

150.3
149.6
147.9
146.3
144.7
143.3
140.4
139.4
134.3
132.4
129.6
128.2
127.5
121.5
116.7
111.9
107.9
106.8
56.4
56.3
f1 (ppm)

### 1-(3,4-Dimethoxyphenyl)-6-(4-fluoro-2-methylphenyl)-1*H*-imidazo[4,5-*c*]pyridine(OXS007287)

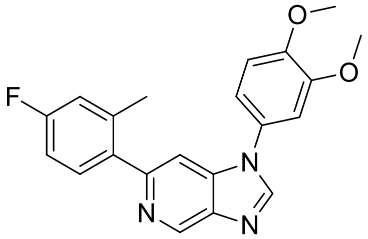

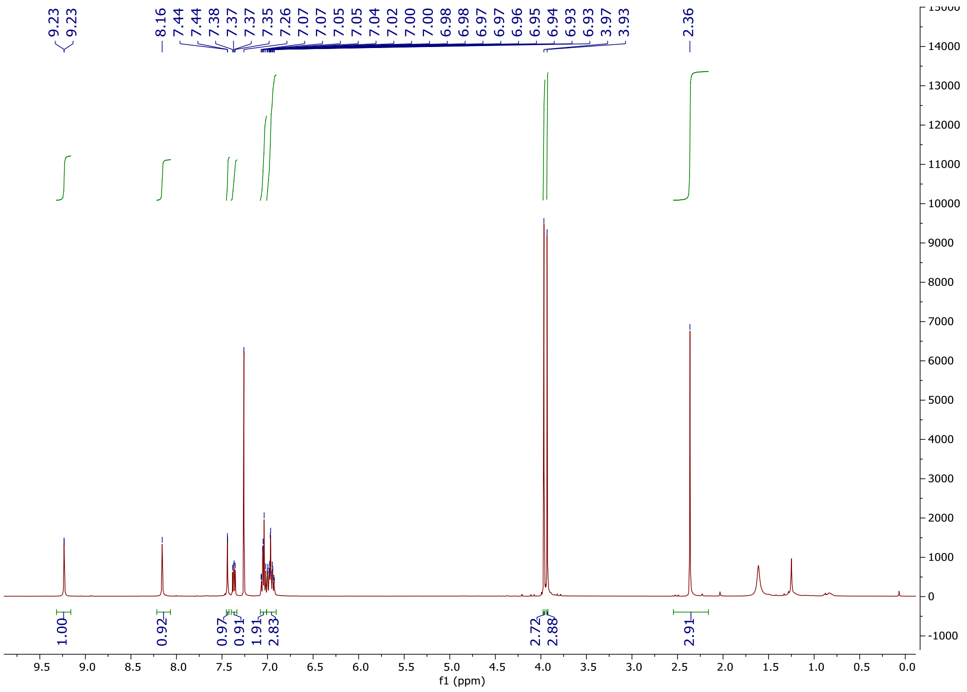

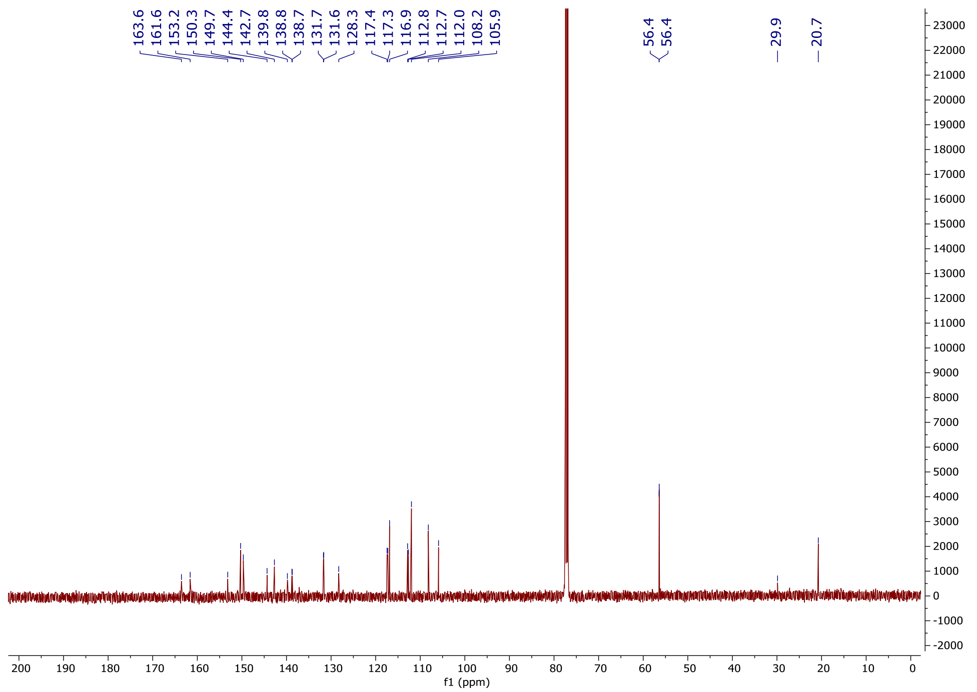

163.6
161.6
153.2
150.3
149.7
144.4
142.7
139.8
138.8
138.7
131.7
131.6
128.3
117.4
117.3
116.9
112.8
112.7
112.0
108.2
105.9
56.4
56.4
29.9
20.7
f1 (ppm)

**1-(3,4-Dimethoxyphenyl)-6-(4-fluoro-2-methoxyphenyl)-1*H*-imidazo[4,5-*c*]pyridine (OXS007184)**

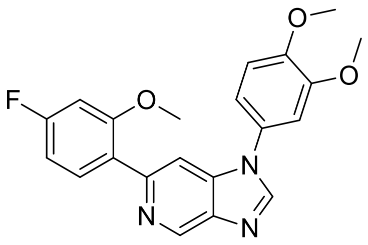

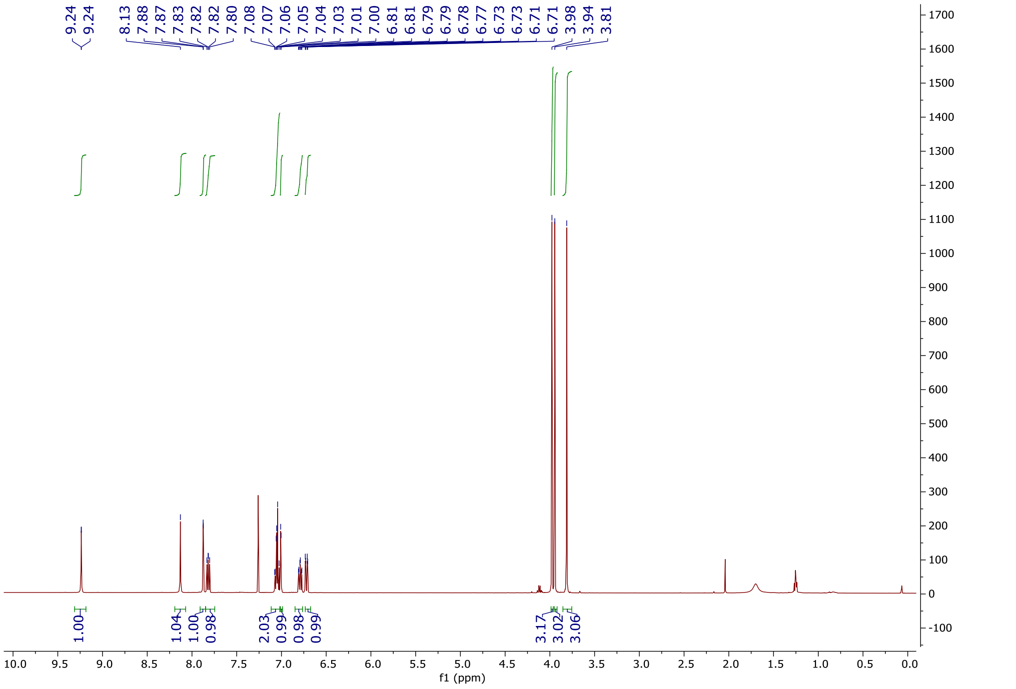

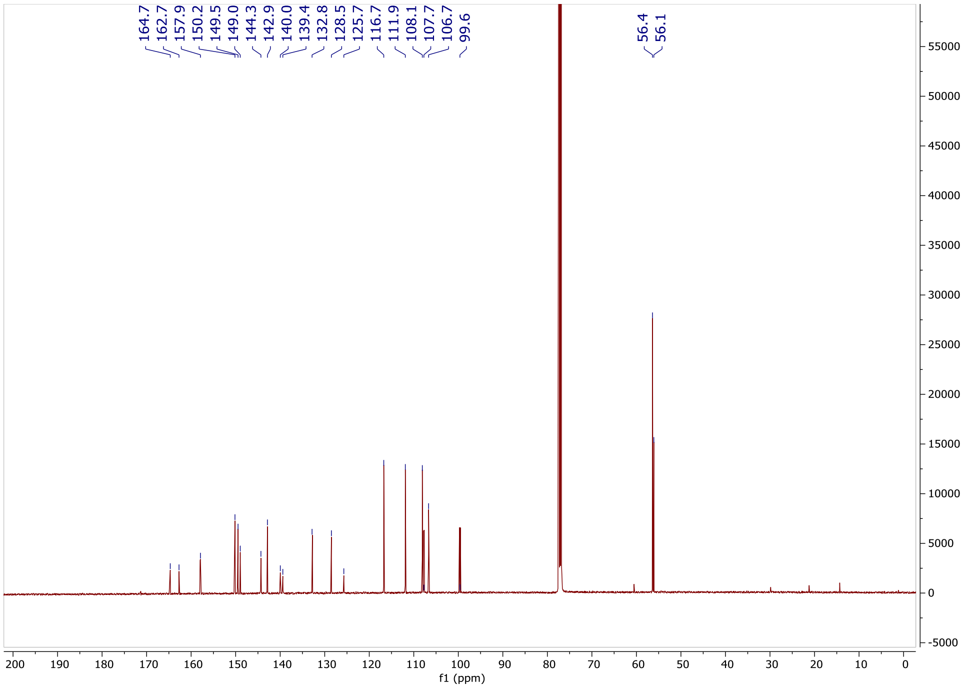
164.7
162.7
157.9
150.2
149.5
149.0
144.3
142.9
140.0
139.4
132.8
128.5
125.7
116.7
111.9
108.1
107.7
106.7
99.6
56.4
56.1
f1 (ppm)

**1-(3,4-Dimethoxyphenyl)-6-(5-fluoro-2-methylphenyl)-1*H*-imidazo[4,5-*c*]pyridine (OXS007317)**

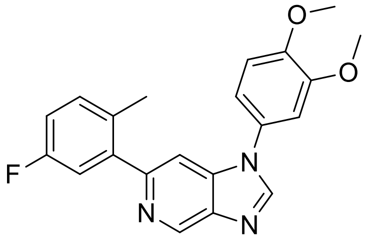

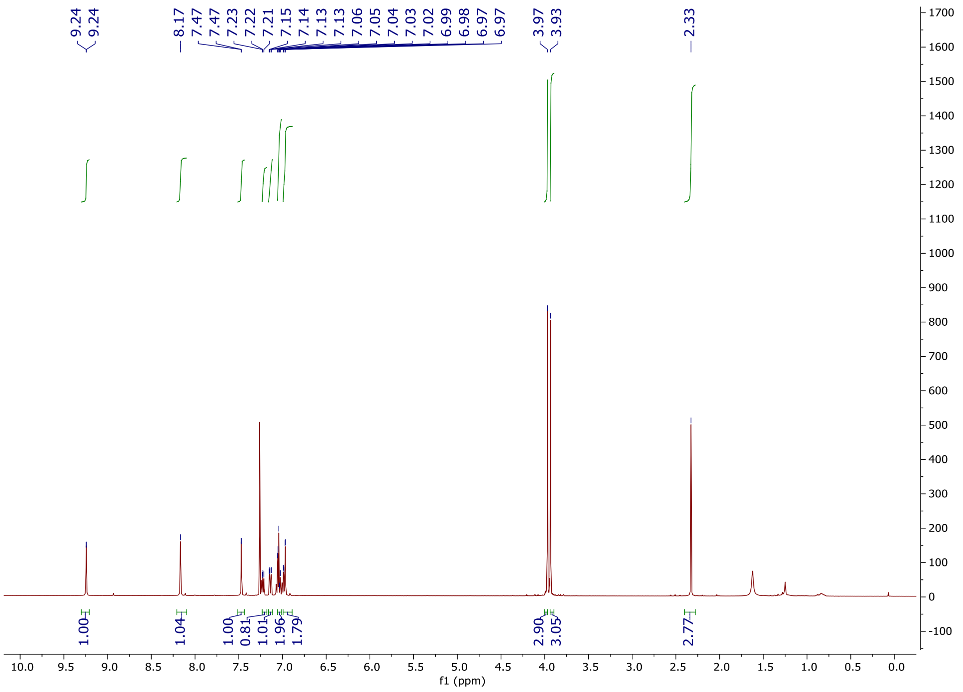

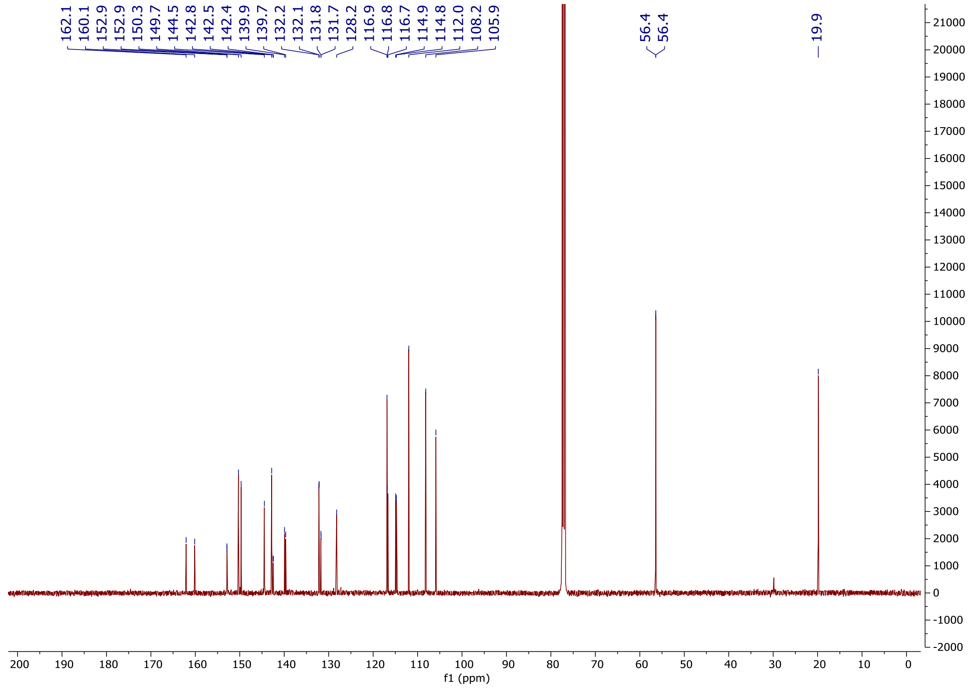
162.1
160.1
152.9
152.9
150.3
149.7
144.5
142.8
142.5
142.4
139.9
139.7
132.2
132.1
131.8
131.7
128.2
116.9
116.8
116.7
114.9
114.8
112.0
108.2
105.9
56.4
56.4
19.9
f1 (ppm)

### 1-(3,4-Dimethoxyphenyl)-6-(5-fluoro-2-methoxyphenyl)-1*H*-imidazo[4,5-*c*]pyridine (OXS007268)

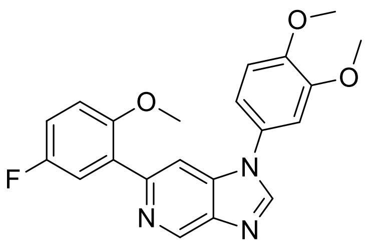

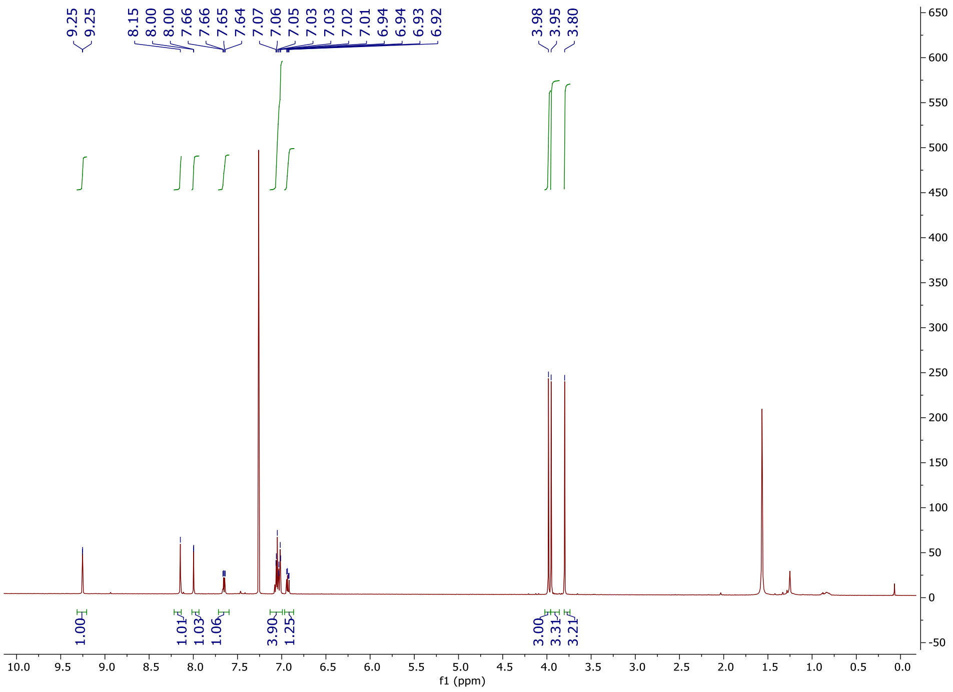

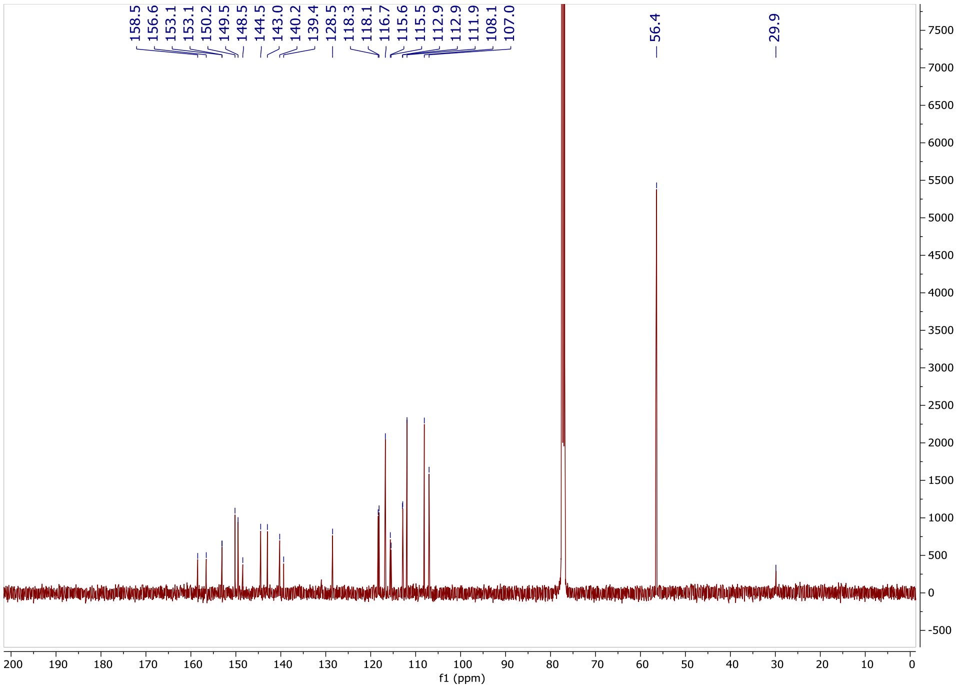
158.5
156.6
153.1
153.1
150.2
149.5
148.5
144.5
143.0
140.2
139.4
128.5
118.3
118.1
116.7
115.6
115.5
112.9
112.9
111.9
108.1
107.0
56.4
29.9
f1 (ppm)

**1-(3,4-Dimethoxyphenyl)-6-(4-fluorophenyl)-1*H*-imidazo[4,5-*c*]pyridine (OXS007823)**

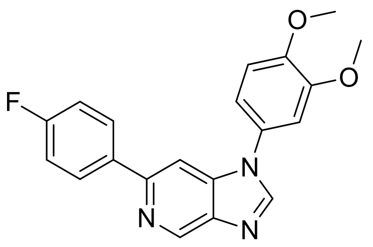

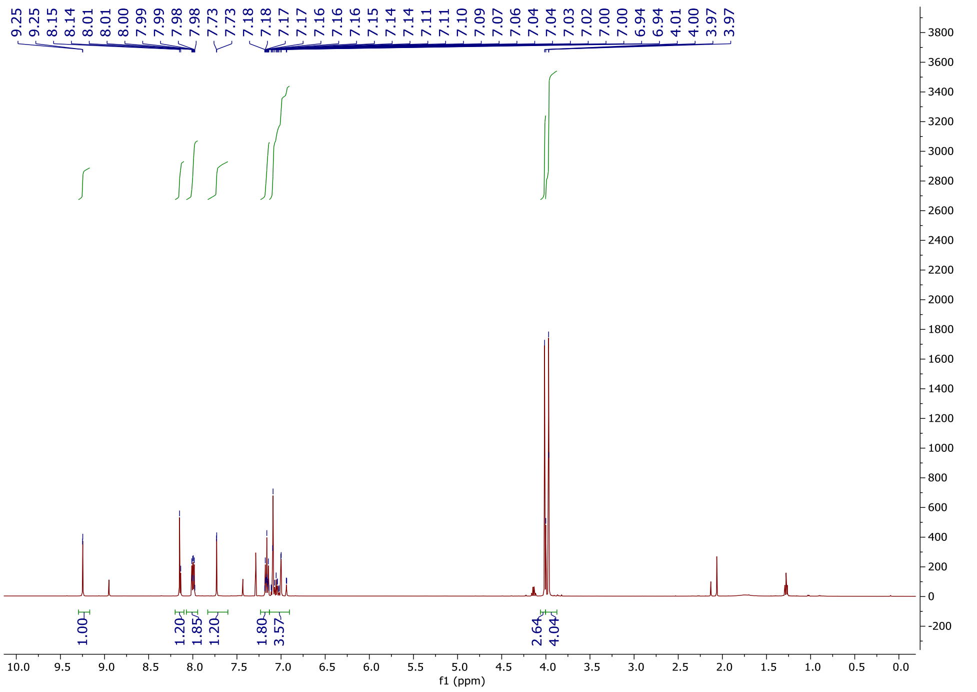

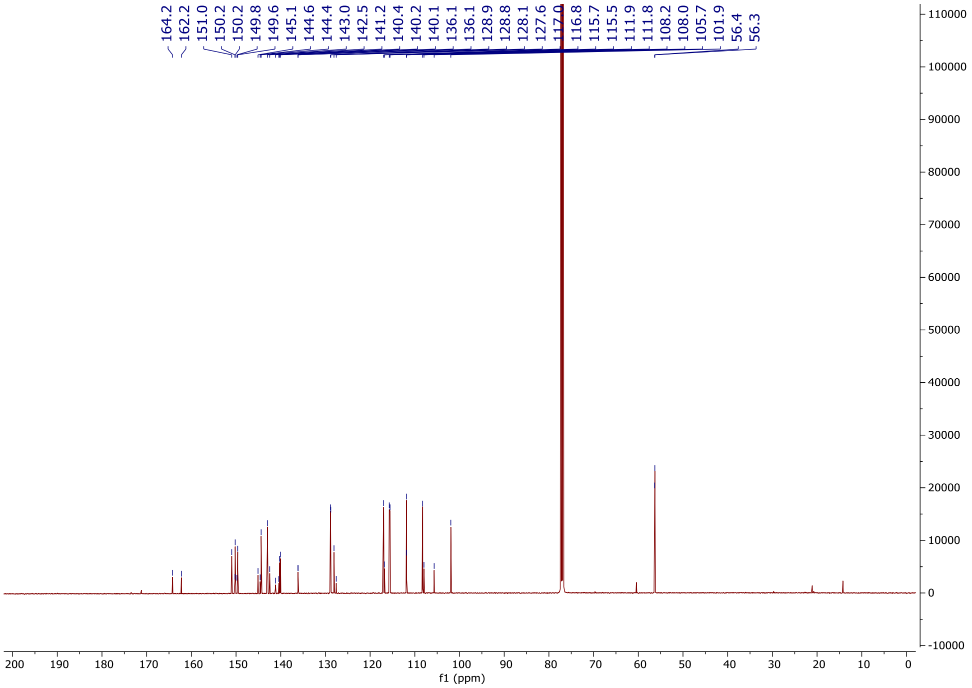
164.2
162.2
151.0
150.2
150.2
149.8
149.6
145.1
144.6
144.4
143.0
142.5
141.2
140.4
140.2
140.1
136.1
136.1
128.9
128.8
128.1
127.6
117.0
116.8
115.7
115.5
111.9
111.8
108.2
108.0
105.7
101.9
56.4
56.3
f1 (ppm)

**4-(1-(3,4-Dimethoxyphenyl)-1*H*-imidazo[4,5-*c*]pyridin-6-yl)morpholine (OXS007267)**

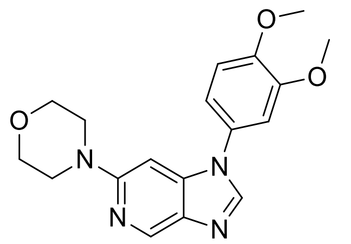

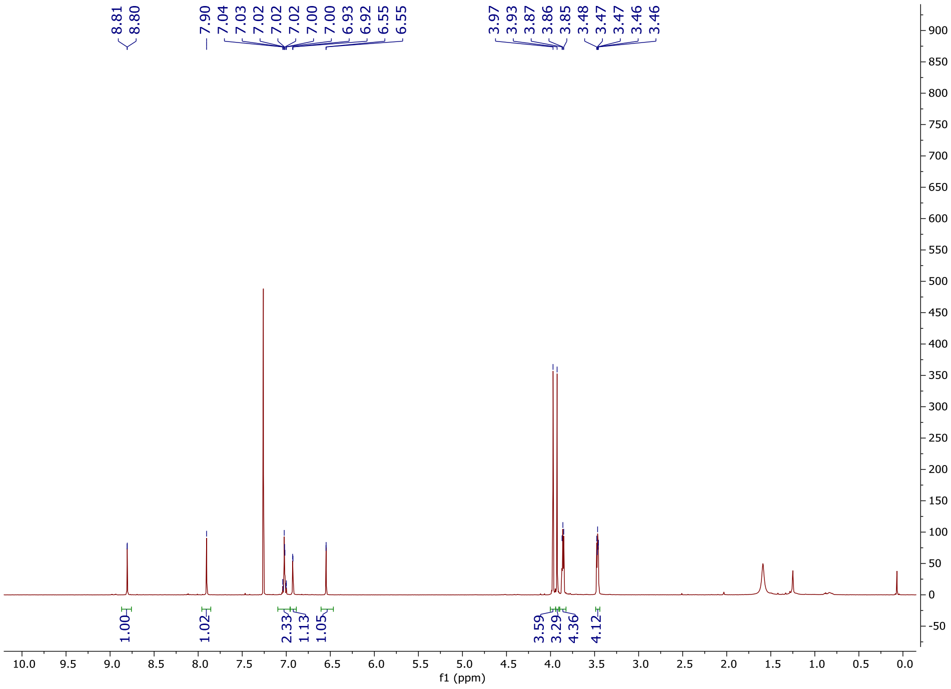

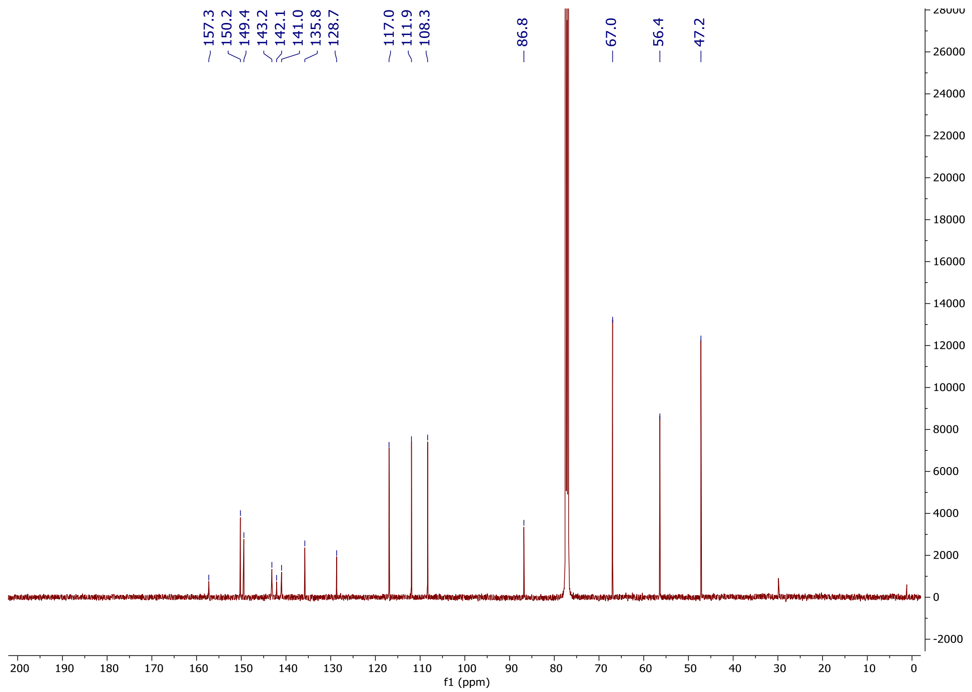

#### 6-(4-Fluoro-2-methylphenyl)-1-(4-methoxyphenyl)-1*H*-imidazo[4,5-*c*]pyridine (OXS007372)

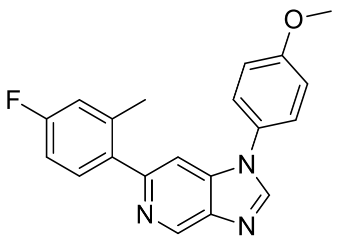

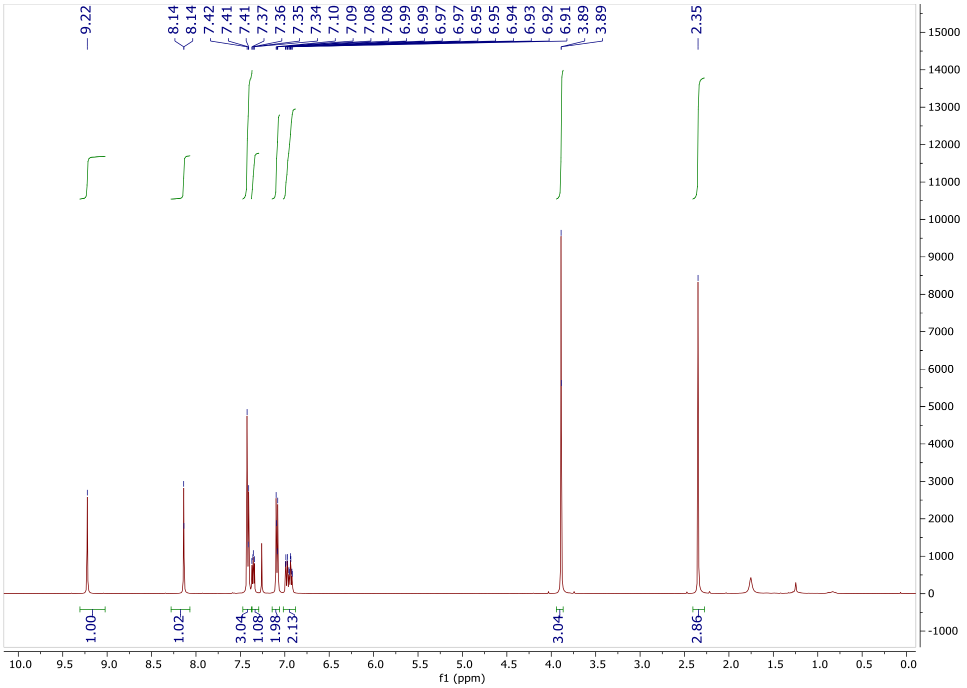
9.22
8.14
8.14
7.42
7.41
7.41
7.37
7.36
7.35
7.34
7.10
7.09
7.08
7.08
6.99
6.97
6.97
6.95
6.95
6.94
6.93
6.92
6.91
3.89
3.89
2.35
1.00
1.02
3.04
1.08
1.98
2.13
3.04
2.86
f1 (ppm)

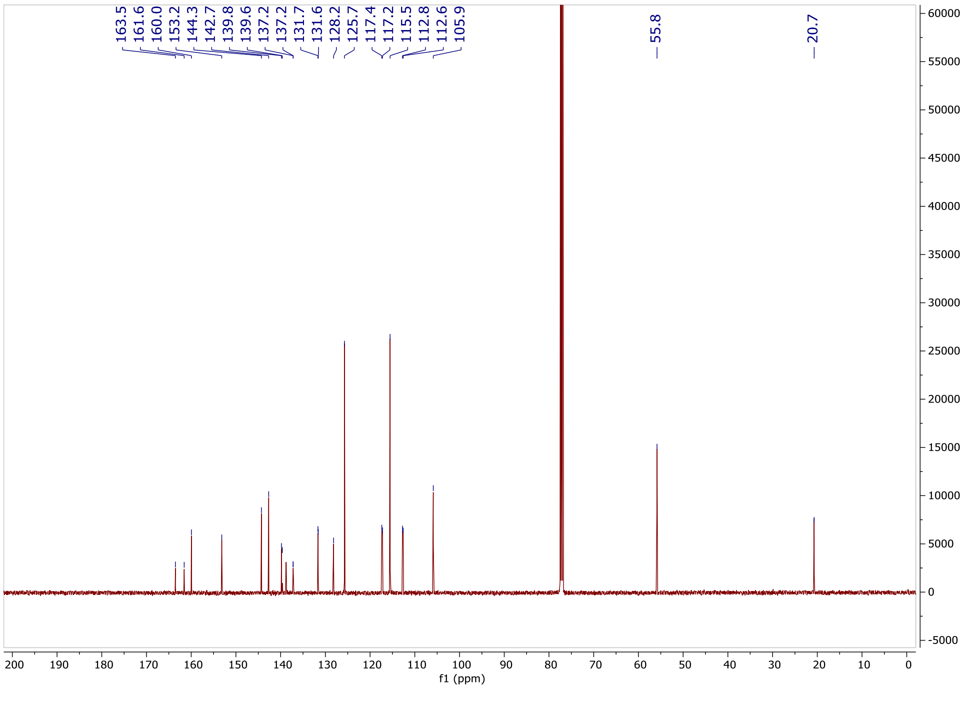
163.5
161.6
160.0
153.2
144.3
142.7
139.8
139.6
137.2
137.2
131.7
131.6
128.2
125.7
117.4
117.2
115.5
112.8
112.6
105.9
55.8
20.7
f1 (ppm)

**1-(4-Ethoxyphenyl)-6-(4-fluoro-2-methylphenyl)-1*H*-imidazo[4,5-*c*]pyridine (OXS007370)**

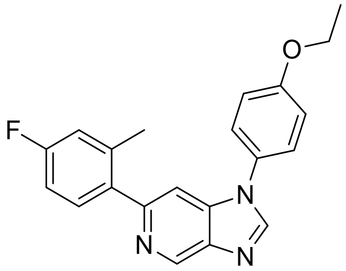
F
N
N
N
O

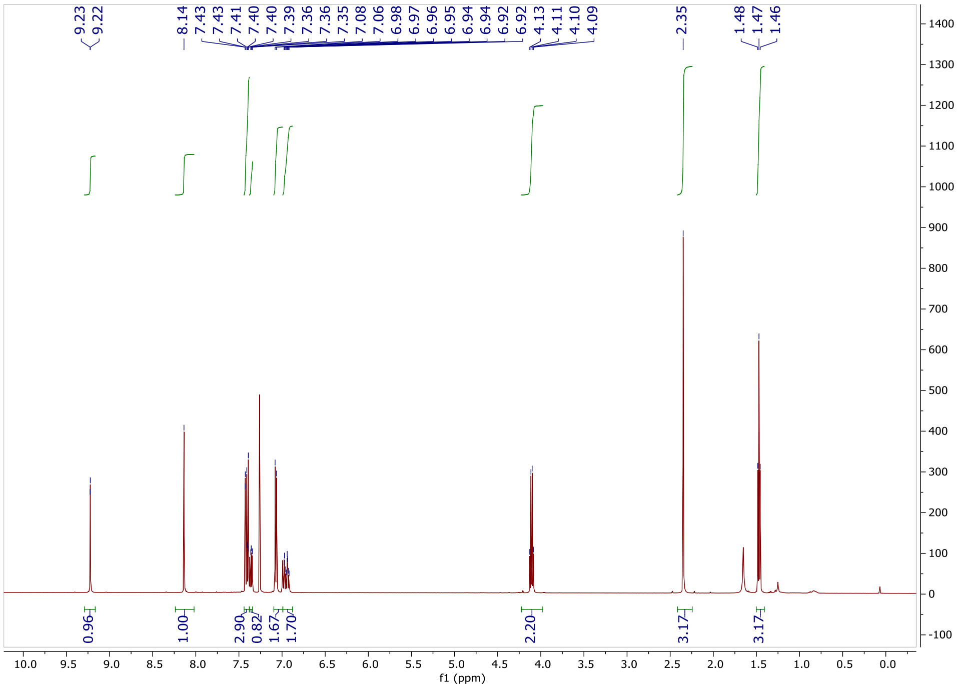
9.23
9.22
8.14
7.43
7.43
7.41
7.40
7.40
7.39
7.36
7.36
7.35
7.08
7.06
6.98
6.97
6.96
6.95
6.94
6.94
6.92
6.92
4.13
4.11
4.10
4.09
2.35
1.48
1.47
1.46
0.96
1.00
2.90
0.82
1.67
1.70
2.20
3.17
3.17
f1 (ppm)

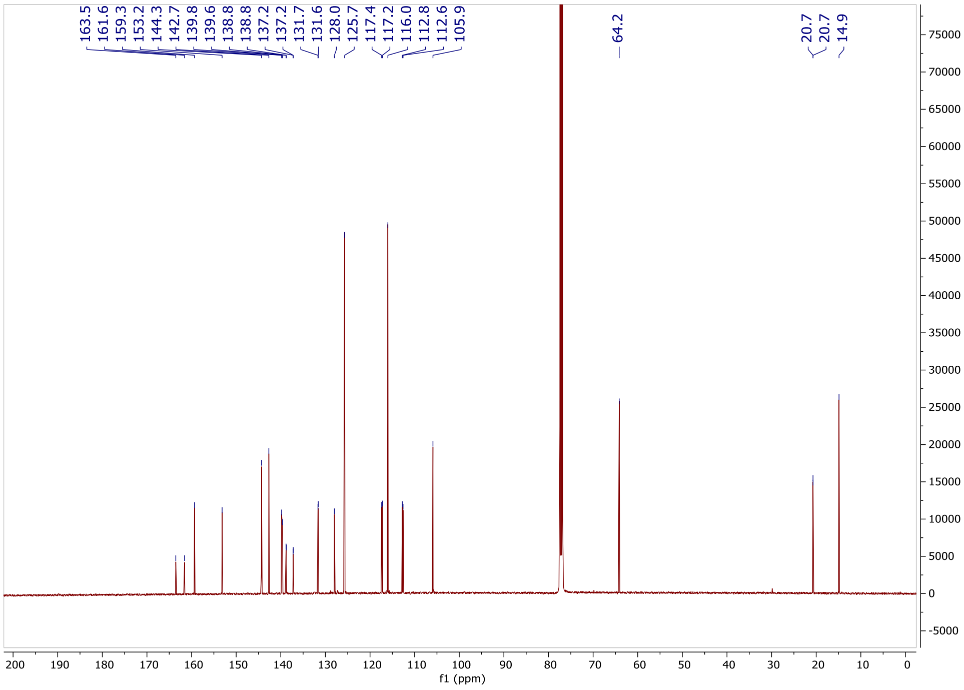
163.5
161.6
159.3
153.2
144.3
142.7
139.8
139.6
138.8
138.8
137.2
137.2
131.7
131.6
128.0
125.7
117.4
117.2
116.0
112.8
112.6
105.9
64.2
20.7
20.7
14.9
f1 (ppm)

**1-(Benzo[*d*][1,3]dioxol-5-yl)-6-(4-fluoro-2-methylphenyl)-1*H*-imidazo[4,5-*c*]pyridine (OXS007392)**

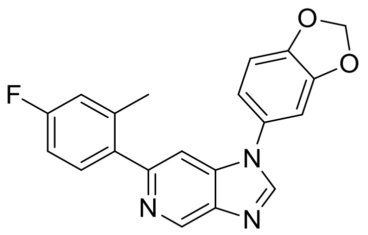

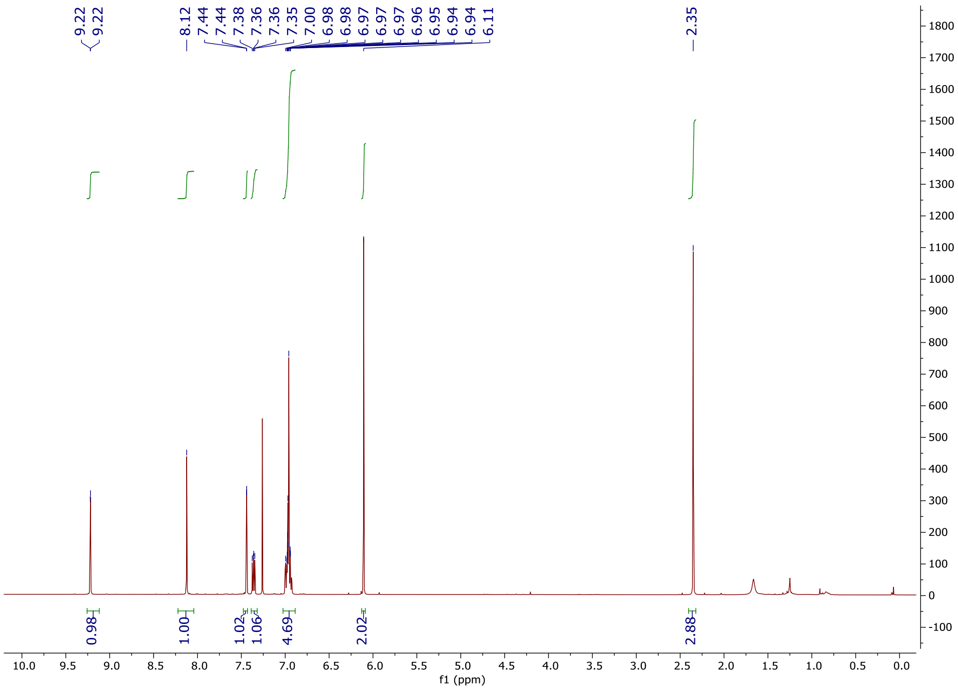

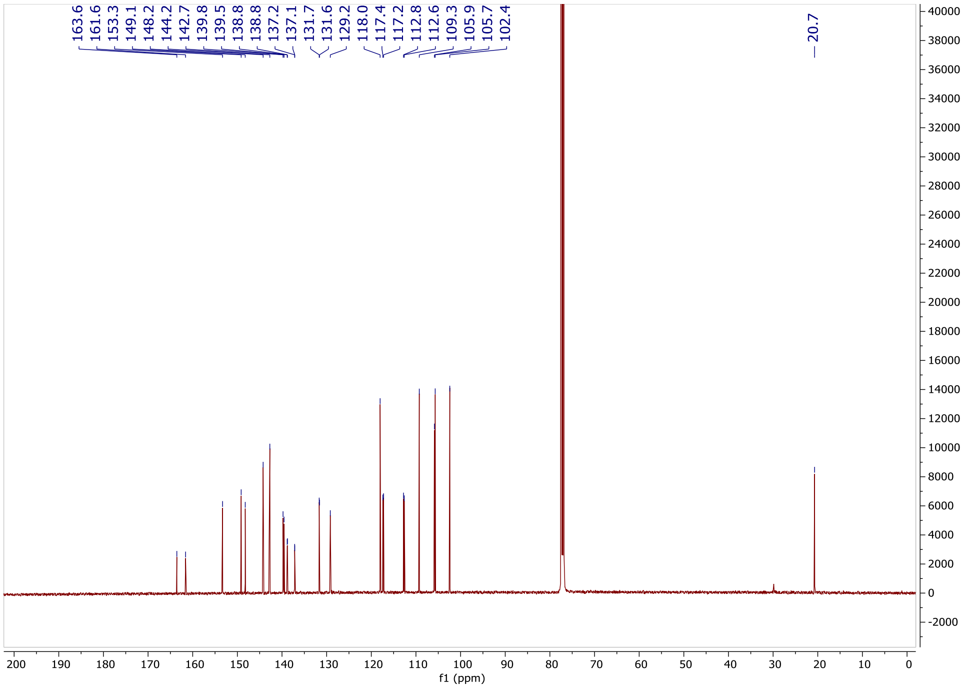

163.6
161.6
153.3
149.1
148.2
144.2
142.7
139.8
139.5
138.8
138.8
137.2
137.1
131.7
131.6
129.2
118.0
117.4
117.2
112.8
112.6
109.3
105.9
105.7
102.4
20.7
f1 (ppm)

### 1-(2,3-Dihydrobenzofuran-5-yl)-6-(4-fluoro-2-methylphenyl)-1*H*-imidazo[4,5-*c*]pyridine (OXS007540)

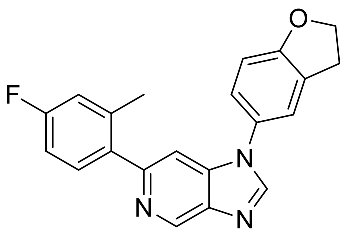

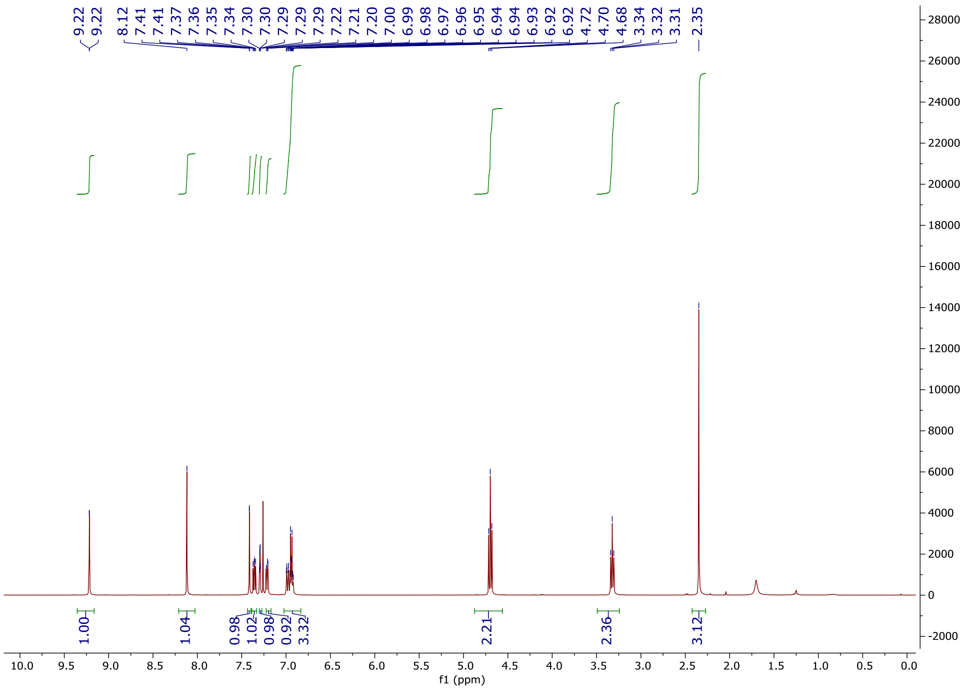

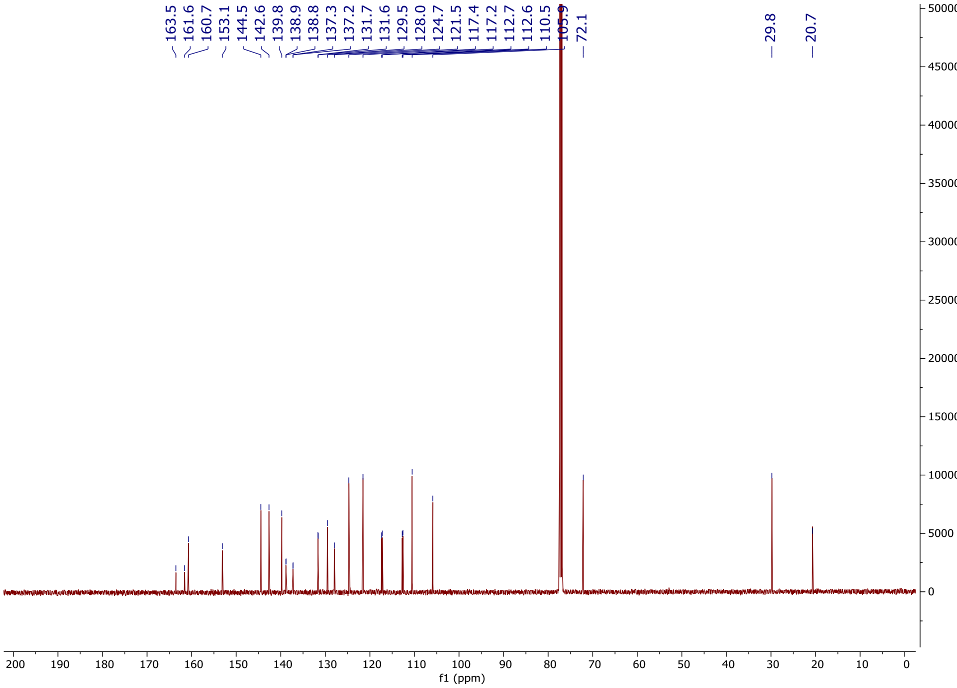

163.5
161.6
160.7
153.1
144.5
142.6
139.8
138.9
138.8
137.3
137.2
131.7
131.6
129.5
128.0
124.7
121.5
117.4
117.2
112.7
112.6
110.5
105.9
72.1
29.8
20.7
50000
45000
40000
35000
30000
25000
20000
15000
10000
5000
0
200 190 180 170 160 150 140 130 120 110 100 90 80 70 60 50 40 30 20 10 0
f1 (ppm)

**6-(4-Fluoro-2-methylphenyl)-1-(4-methylphenyl)-1*H*-imidazo[4,5-*c*]pyridine (OXS008411)**

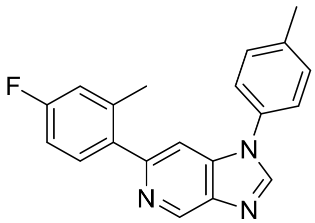

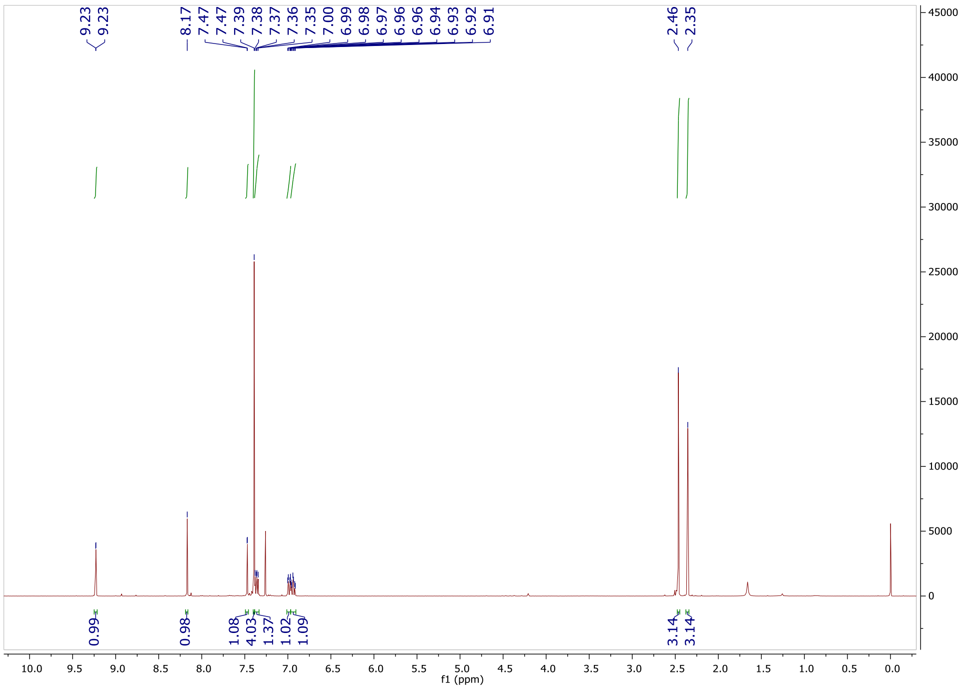

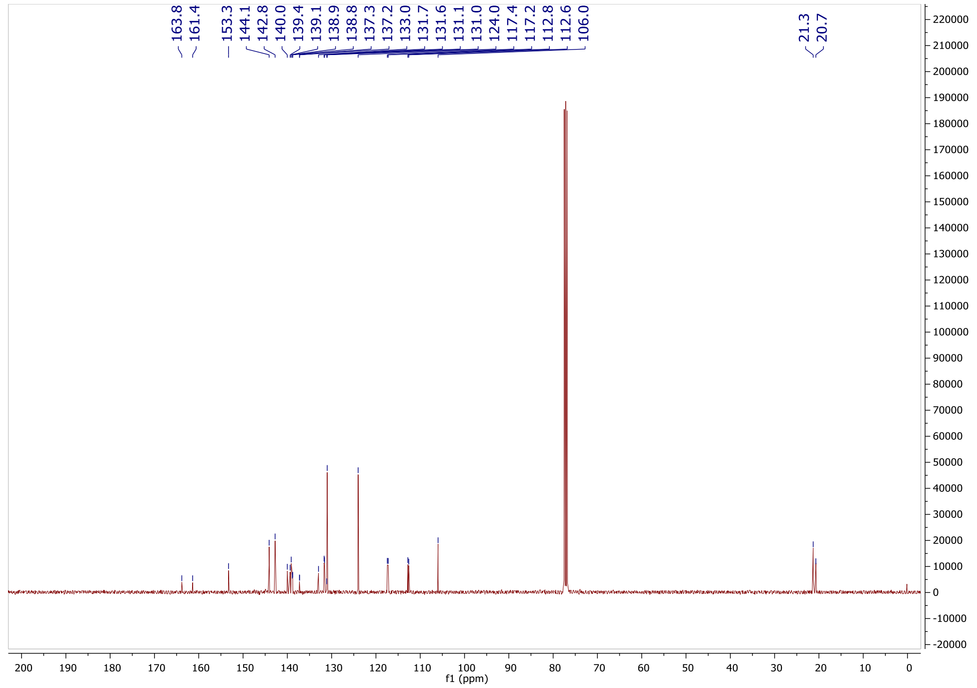
163.8
161.4
153.3
144.1
142.8
140.0
139.4
139.1
138.9
138.8
137.3
137.2
133.0
131.7
131.6
131.1
131.0
124.0
117.4
117.2
112.8
112.6
106.0
21.3
20.7
f1 (ppm)

**6-(4-Fluoro-2-methylphenyl)-1-(4-(methylsulfonyl)phenyl)-1*H*-imidazo[4,5-*c*]pyridine (OXS007839)**

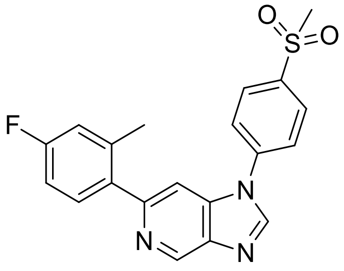

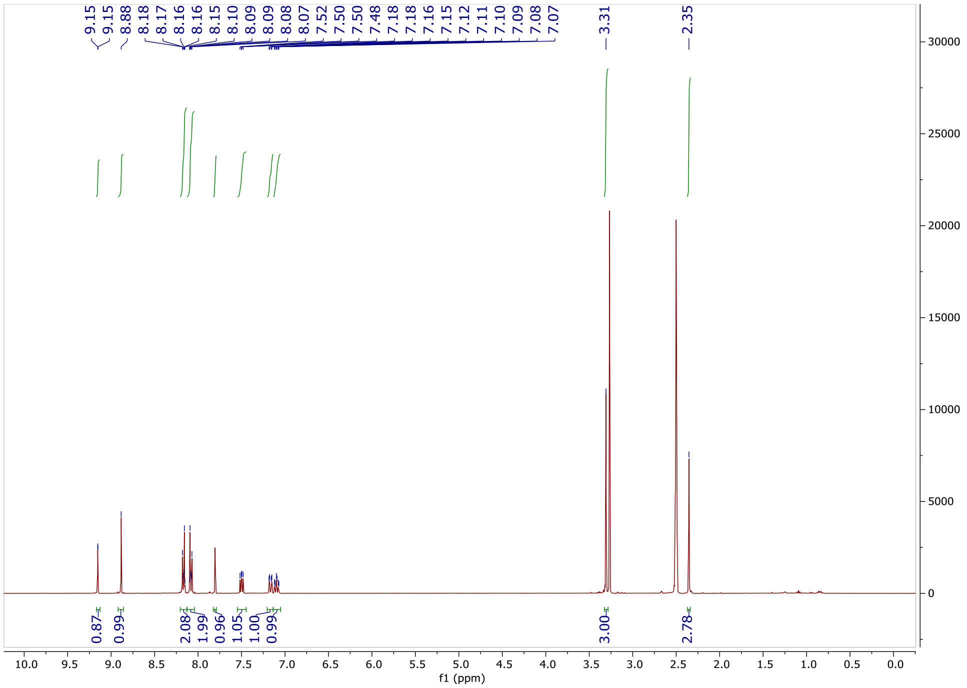

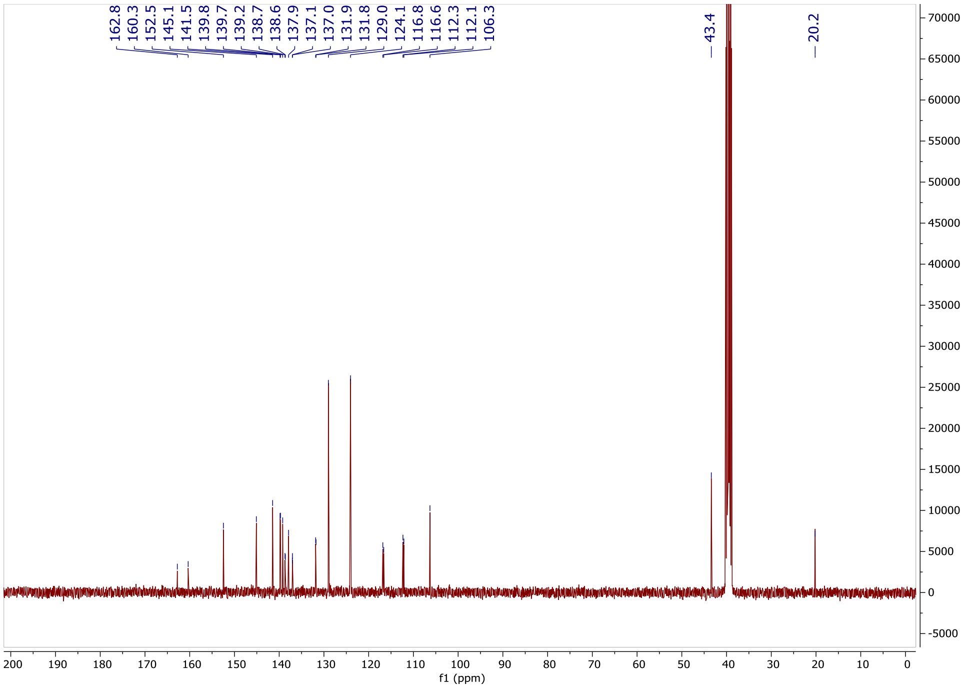

162.8
160.3
152.5
145.1
141.5
139.8
139.7
139.2
138.7
138.6
137.9
137.1
137.0
131.9
131.8
129.0
124.1
116.8
116.6
112.3
112.1
106.3
43.4
20.2
f1 (ppm)

#### 1-(4-(Difluoromethoxy)phenyl)-6-(4-fluoro-2-methylphenyl)-1*H*-imidazo[4,5-*c*]pyridine (OXS007417)

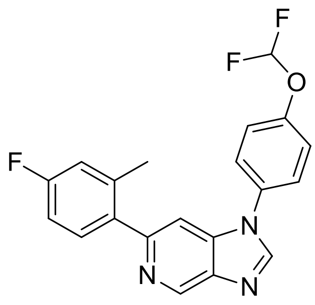

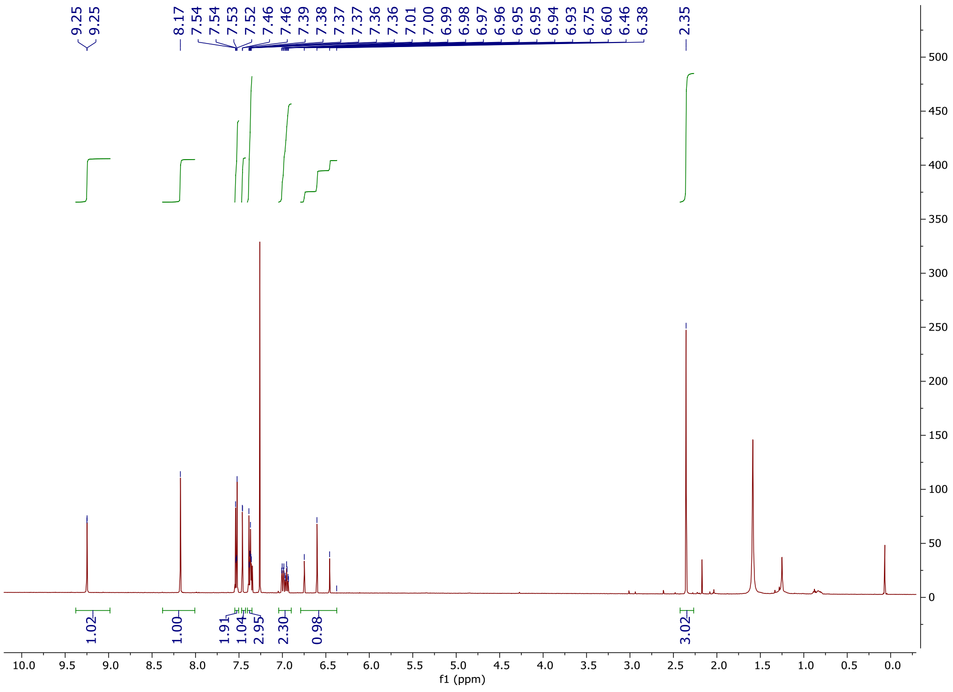

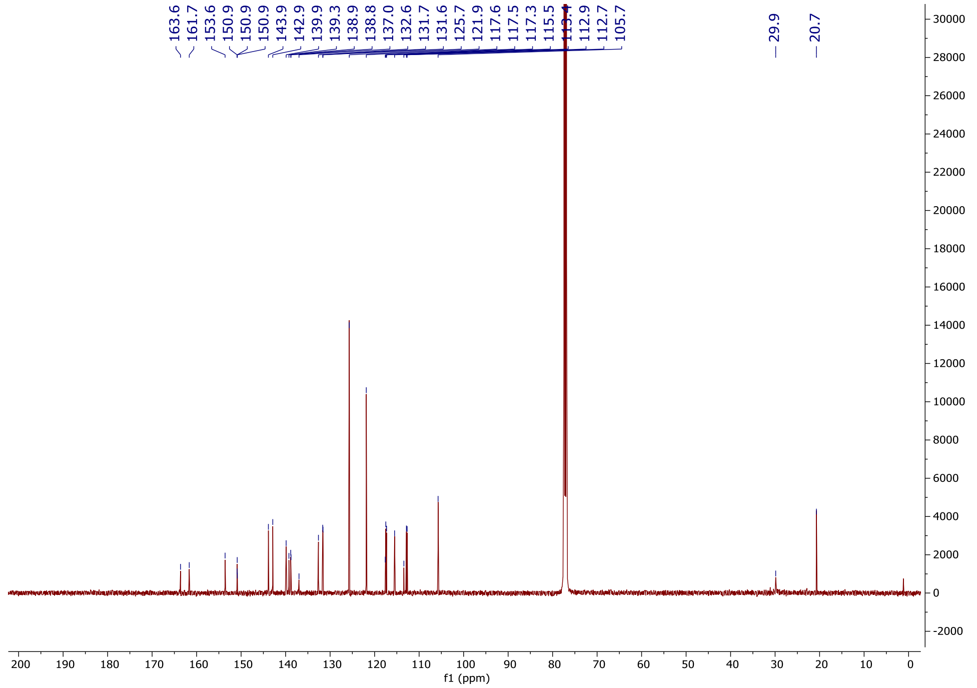

163.6
161.7
153.6
150.9
150.9
150.9
143.9
142.9
139.9
139.3
138.9
138.8
137.0
132.6
131.7
131.6
125.7
121.9
117.6
117.5
117.3
115.5
113.4
112.9
112.7
105.7
29.9
20.7
f1 (ppm)

**6-(4-Fluoro-2-methylphenyl)-1-(4-(trifluoromethoxy)phenyl)-1*H*-imidazo[4,5-*c*]pyridine (OXS007951)**

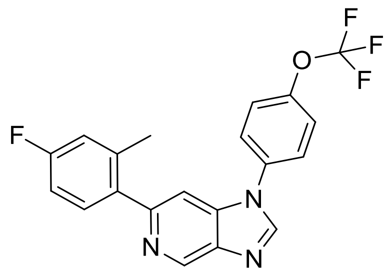

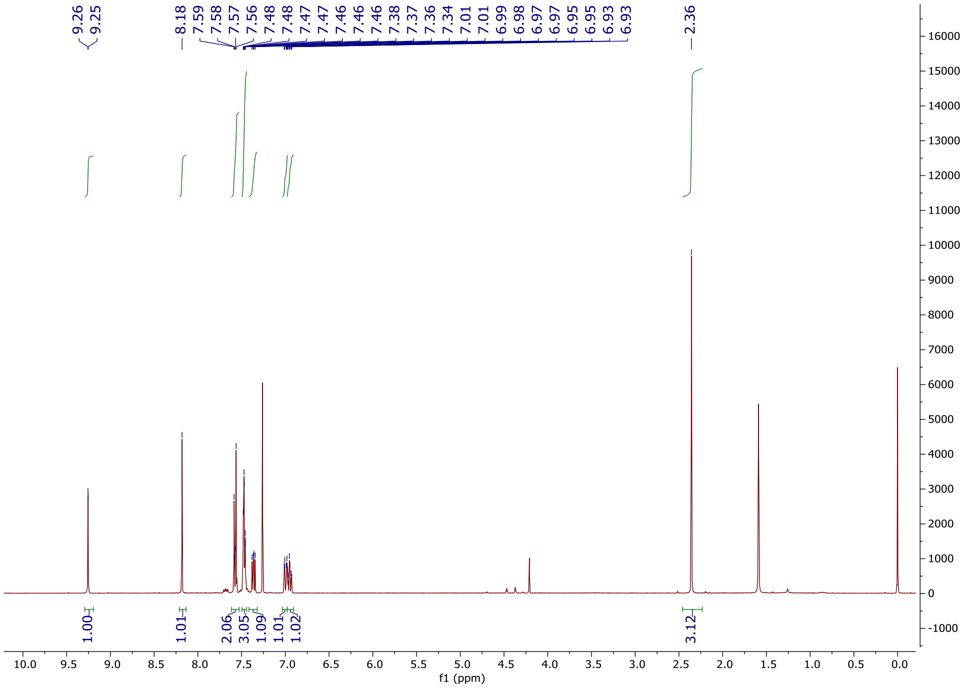

9.26
9.25
8.18
7.59
7.58
7.57
7.56
7.48
7.47
7.46
7.46
7.38
7.37
7.36
7.34
7.01
6.99
6.98
6.97
6.95
6.95
6.93
6.93
2.36
1.00
1.01
2.06
3.05
1.09
1.01
1.02
3.12
f1 (ppm)

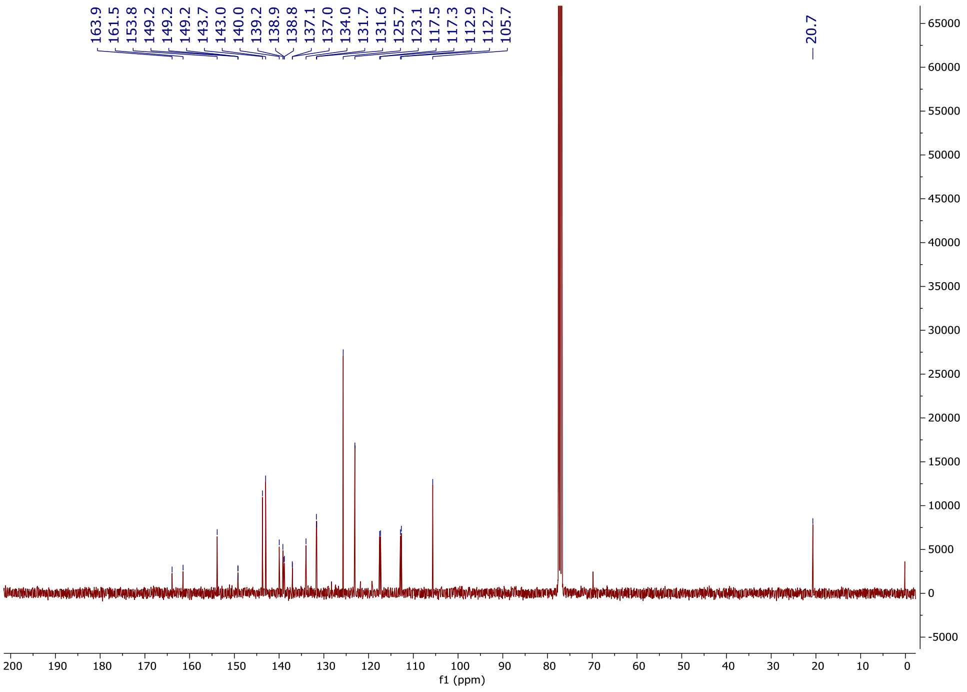

163.9
161.5
153.8
149.2
149.2
143.7
143.0
140.0
139.2
138.9
138.8
137.1
137.0
134.0
131.7
131.6
125.7
123.1
117.5
117.3
112.9
112.7
105.7
20.7
f1 (ppm)

**6-(4-Fluoro-2-methylphenyl)-1-(3-fluoro-4-methoxyphenyl)-1*H*-imidazo[4,5-*c*]pyridine (OXS007373)**

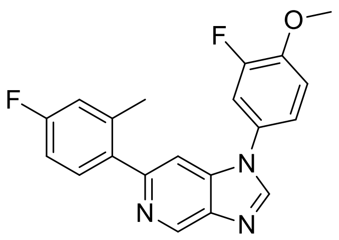

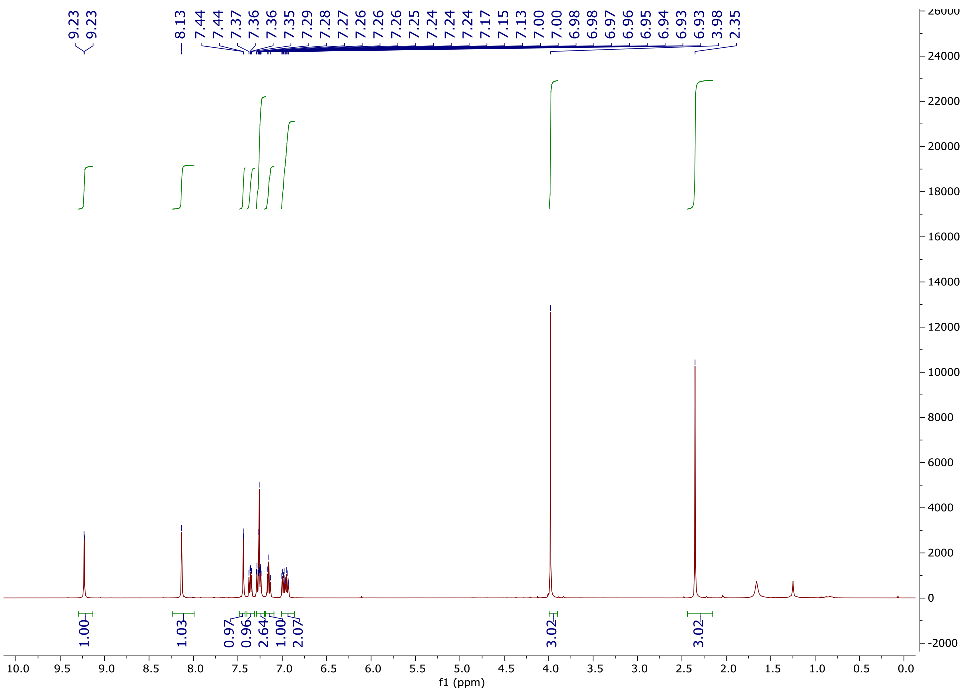

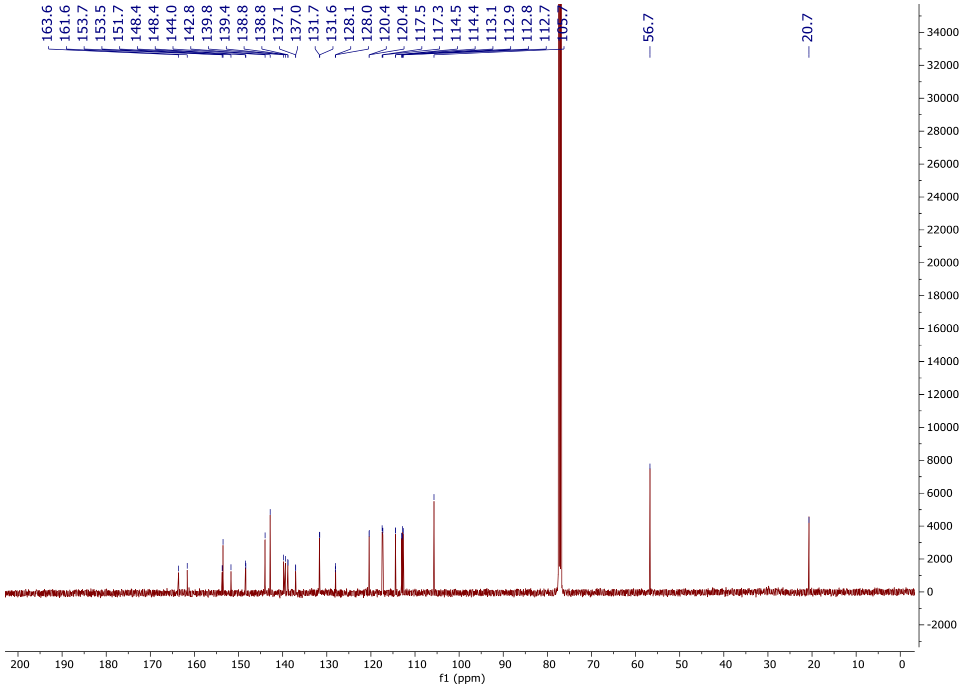

163.6
161.6
153.7
153.5
151.7
148.4
148.4
144.0
142.8
139.8
139.4
138.8
138.8
137.1
137.0
131.7
131.6
128.1
128.0
120.4
120.4
117.5
117.3
114.5
114.4
113.1
112.9
112.8
112.7
105.7
56.7
20.7
f1 (ppm)

**1-(3-Chloro-4-methoxyphenyl)-6-(4-fluoro-2-methylphenyl)-1*H*-imidazo[4,5-c]pyridine (OXS008397)**

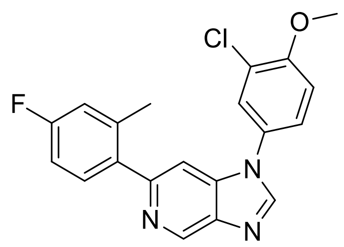

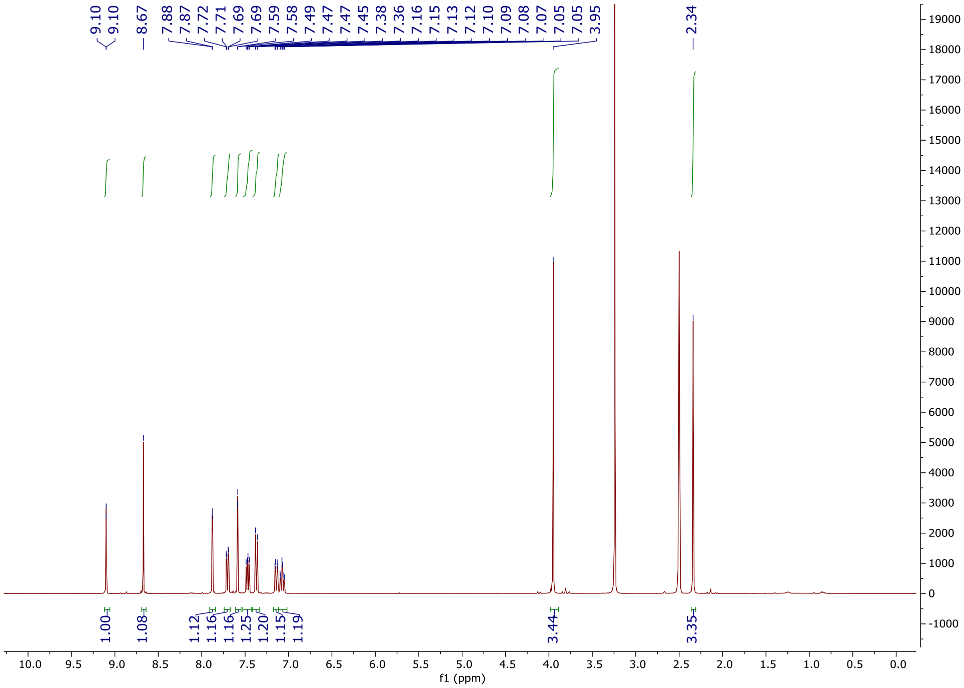

9.10
9.10
8.67
7.88
7.87
7.72
7.71
7.69
7.69
7.59
7.58
7.49
7.47
7.47
7.45
7.38
7.36
7.16
7.15
7.13
7.12
7.10
7.09
7.08
7.07
7.05
7.05
3.95
2.34
1.00
1.08
1.12
1.16
1.16
1.25
1.20
1.15
1.19
3.44
3.35
f1 (ppm)

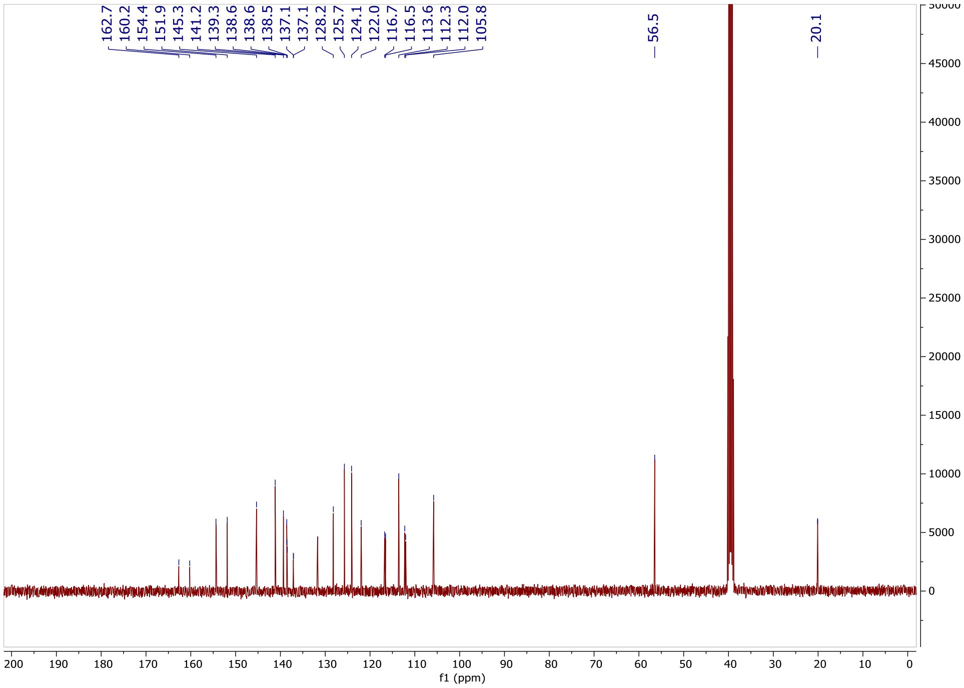

162.7
160.2
154.4
151.9
145.3
141.2
139.3
138.6
138.6
138.5
137.1
137.1
128.2
125.7
124.1
122.0
116.7
116.5
113.6
112.3
112.0
105.8
56.5
20.1
f1 (ppm)

**6-(4-Fluoro-2-methylphenyl)-1-(4-methoxy-3-methylphenyl)-1*H*-imidazo[4,5-*c*]pyridine (OXS008444)**

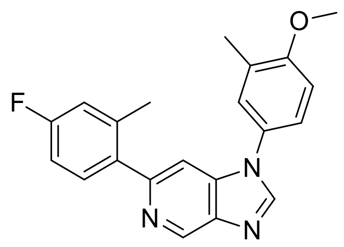

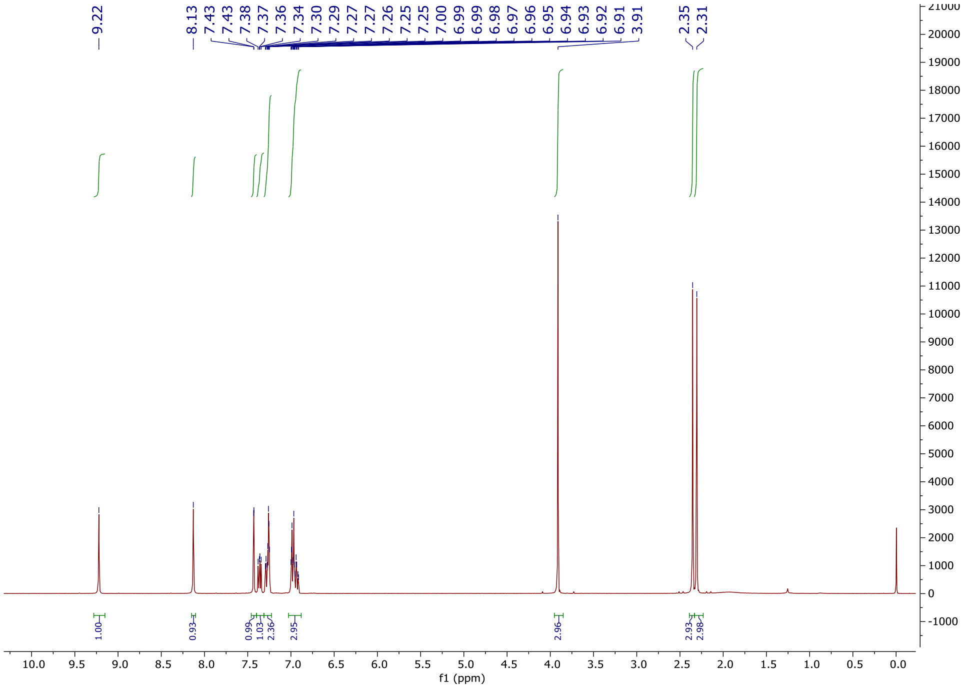
9.22
8.13
7.43
7.43
7.38
7.37
7.36
7.34
7.30
7.29
7.27
7.27
7.26
7.25
7.25
7.00
6.99
6.99
6.98
6.97
6.96
6.95
6.94
6.93
6.92
6.91
3.91
2.35
2.31
1.00
0.93
0.99
1.03
2.36
2.95
2.96
2.93
2.98
f1 (ppm)

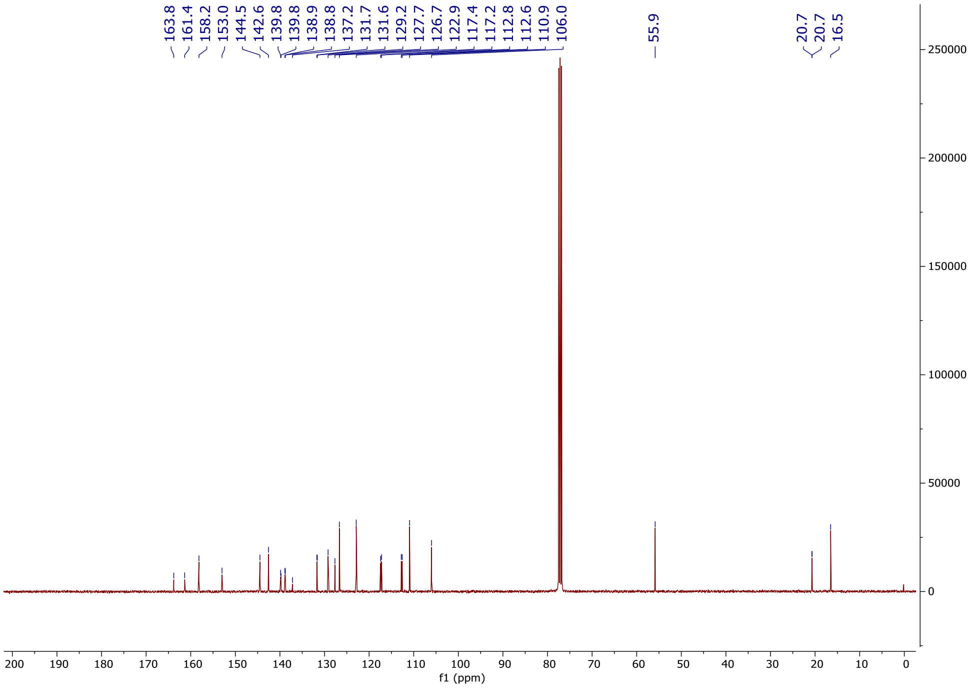
163.8
161.4
158.2
153.0
144.5
142.6
139.8
138.9
138.8
137.2
131.7
131.6
129.2
127.7
126.7
122.9
117.4
117.2
112.8
112.6
110.9
106.0
55.9
20.7
20.7
16.5
f1 (ppm)

**6-(4-Fluoro-2-methylphenyl)-1-(3-isopropyl-4-methoxyphenyl)-1*H*-imidazo[4,5-*c*]pyridine (OXS007787)**

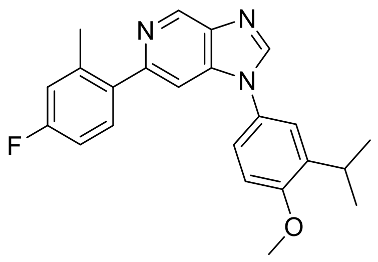

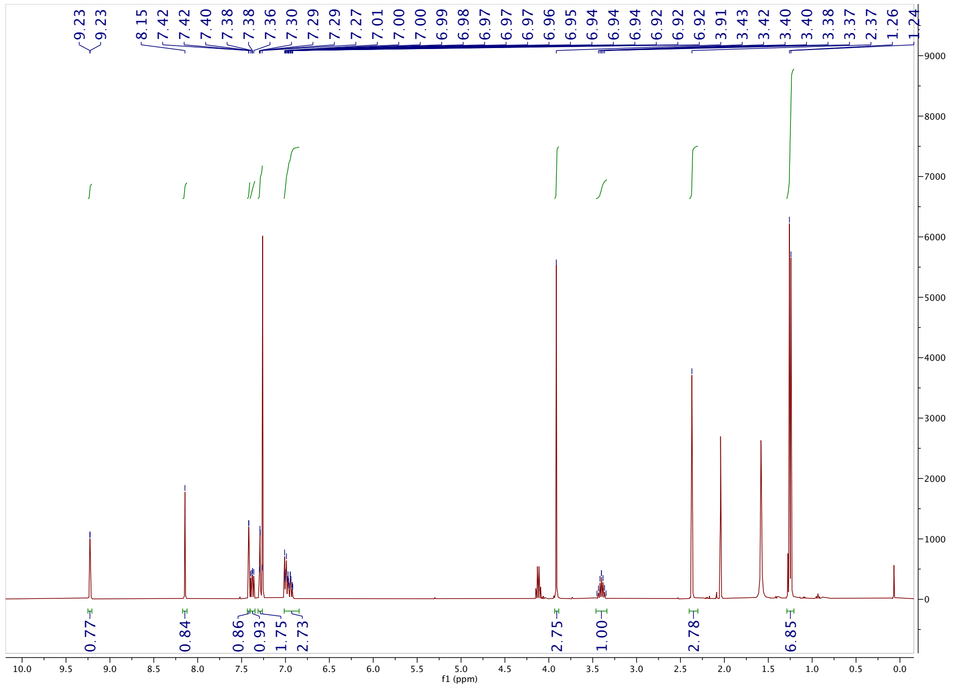
9.23
9.23
8.15
7.42
7.42
7.40
7.38
7.38
7.36
7.30
7.29
7.29
7.27
7.01
7.00
7.00
6.99
6.98
6.97
6.97
6.97
6.96
6.95
6.94
6.94
6.94
6.92
6.92
6.92
3.91
3.43
3.42
3.40
3.40
3.38
3.37
2.37
1.26
1.24
0.77
0.84
0.86
0.93
1.75
2.73
2.75
1.00
2.78
6.85
f1 (ppm)

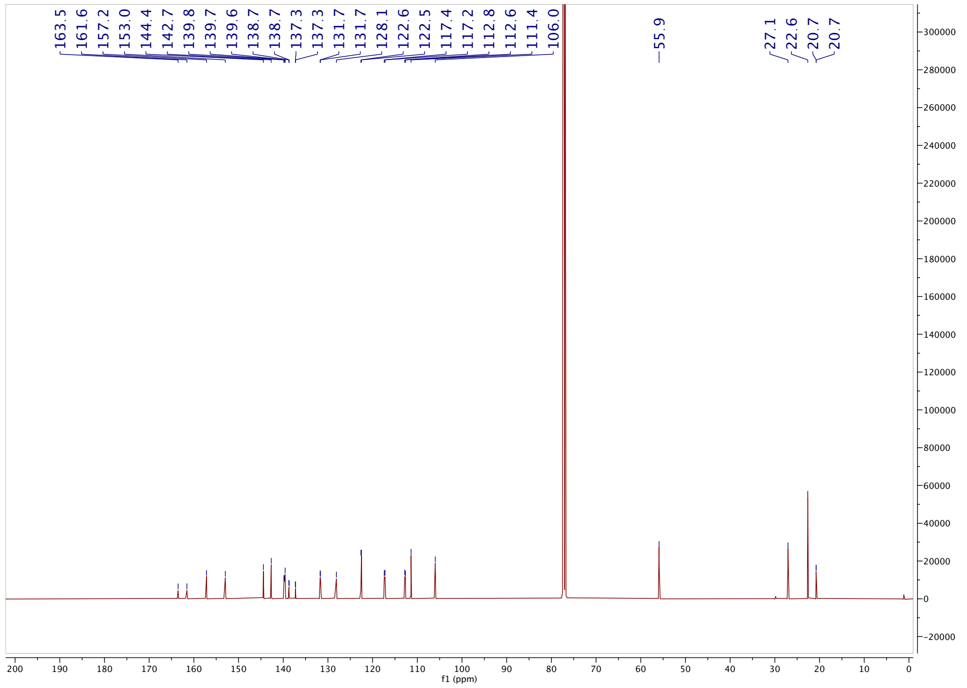
163.5
161.6
157.2
153.0
144.4
142.7
139.8
139.7
139.6
138.7
138.7
137.3
137.3
131.7
131.7
128.1
122.6
122.5
117.4
117.2
112.8
112.6
111.4
106.0
55.9
27.1
22.6
20.7
20.7
f1 (ppm)

**6-(4-Fluoro-2-methylphenyl)-1-(6-methoxypyridin-3-yl)-1*H*-imidazo[4,5-*c*]pyridine (OXS007749)**

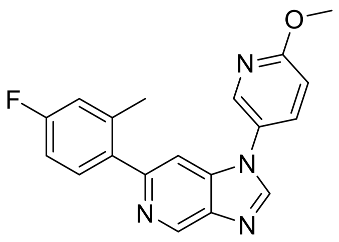

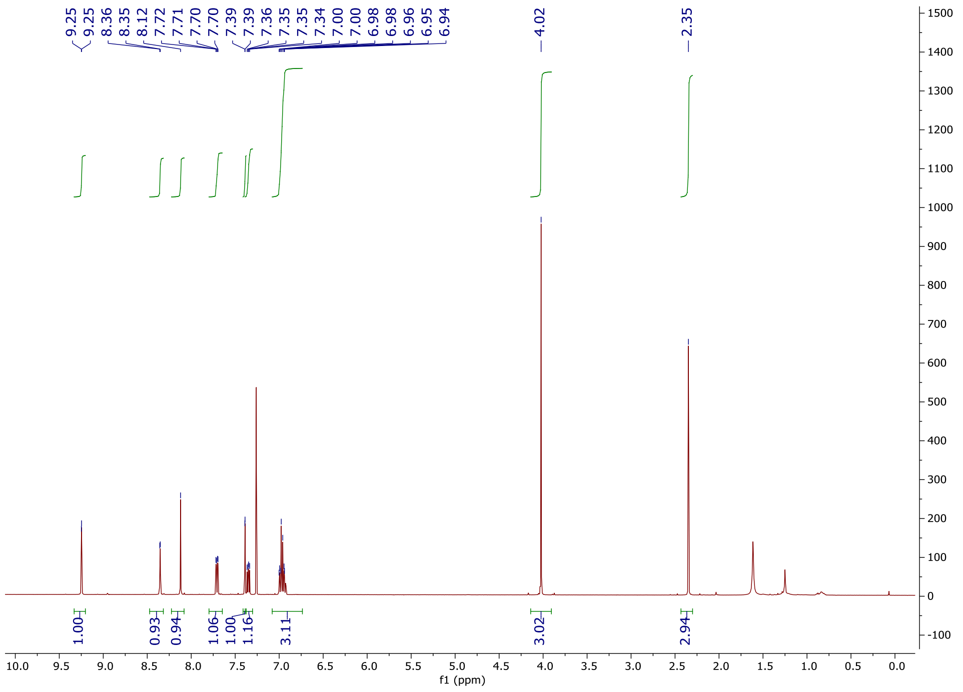

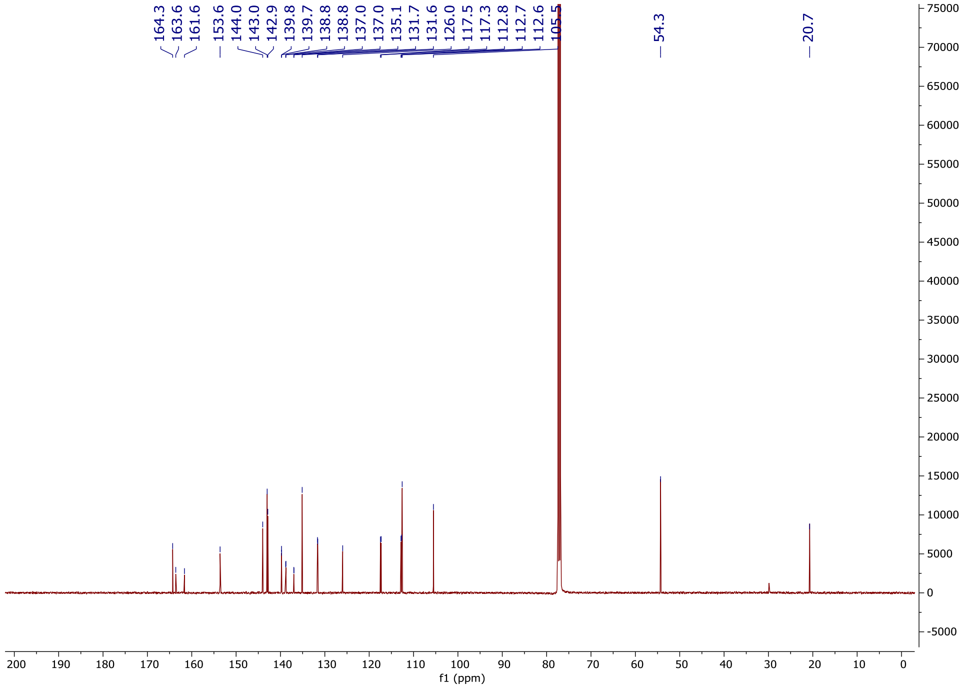
164.3
163.6
161.6
153.6
144.0
143.0
142.9
139.8
139.7
138.8
138.8
137.0
137.0
135.1
131.7
131.6
126.0
117.5
117.3
112.8
112.7
112.6
105.5
54.3
20.7
f1 (ppm)

**5-(6-(4-fluoro-2-methylphenyl)-1H-imidazo[4,5-c]pyridin-1-yl)pyridin-2-ol (OXS007801)**

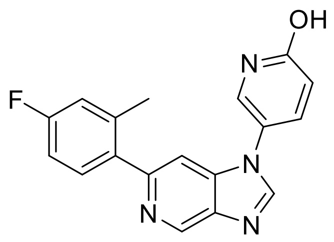

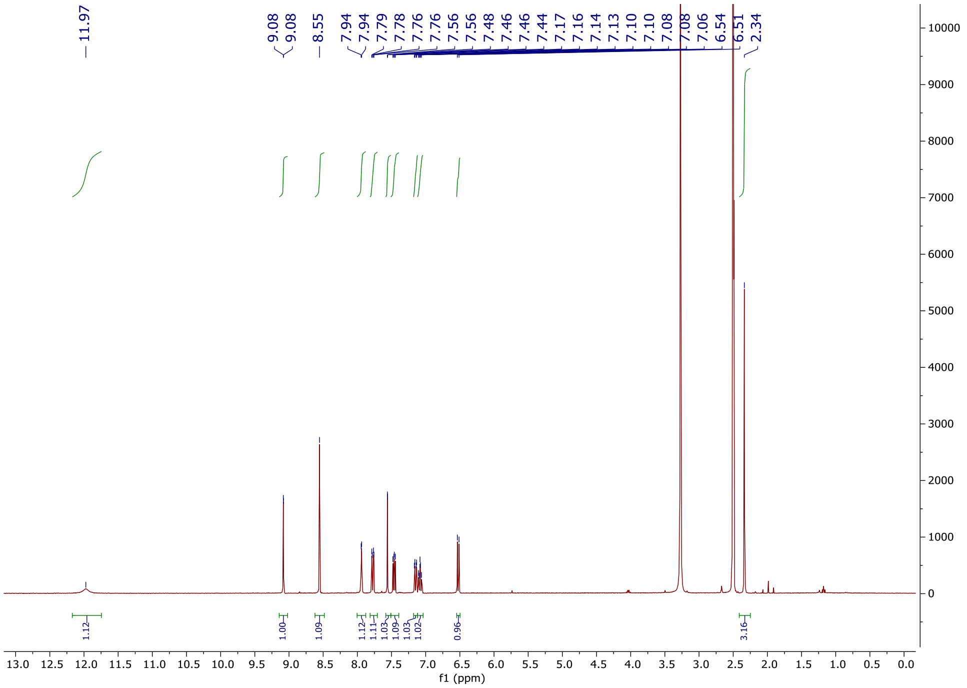
11.97
9.08
9.08
8.55
7.94
7.94
7.79
7.78
7.76
7.76
7.56
7.56
7.48
7.46
7.46
7.44
7.17
7.16
7.14
7.13
7.10
7.10
7.08
7.08
7.06
6.54
6.51
2.34
1.12
1.00
1.09
1.12
1.11
1.03
1.09
1.03
1.02
0.96
3.16
f1 (ppm)

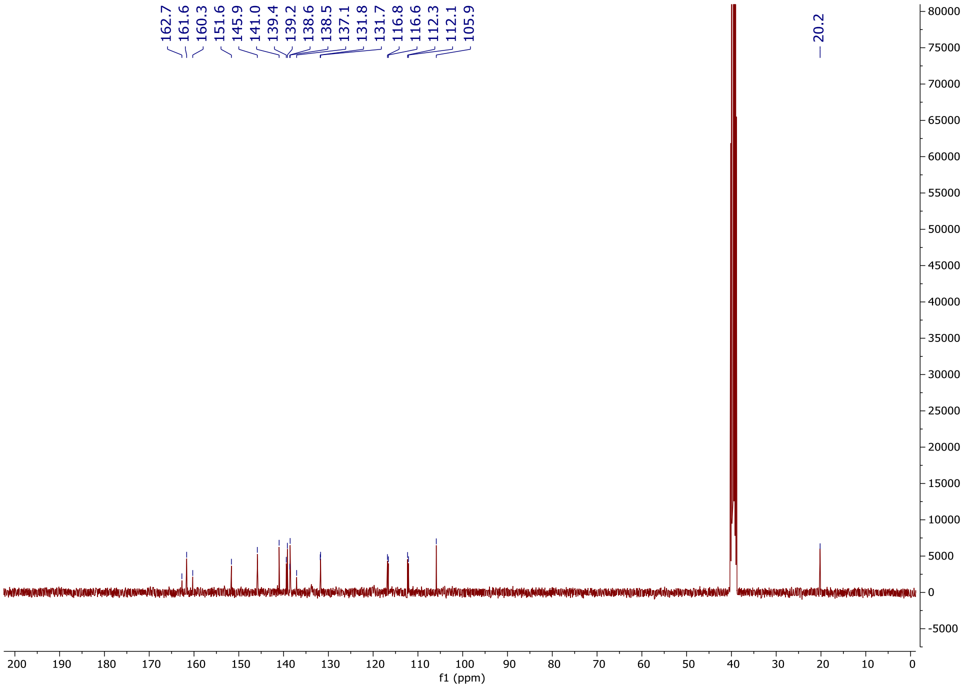
162.7
161.6
160.3
151.6
145.9
141.0
139.4
139.2
138.6
138.5
137.1
131.8
131.7
116.8
116.6
112.3
112.1
105.9
20.2
f1 (ppm)

### 1-(6-(Difluoromethoxy)pyridin-3-yl)-6-(4-fluoro-2-methylphenyl)-1*H*-imidazo[4,5-*c*]pyridine (OXS007571)

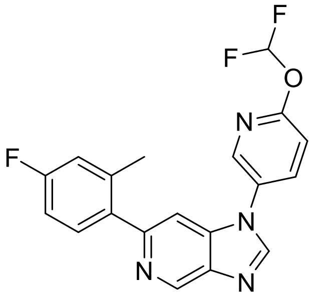

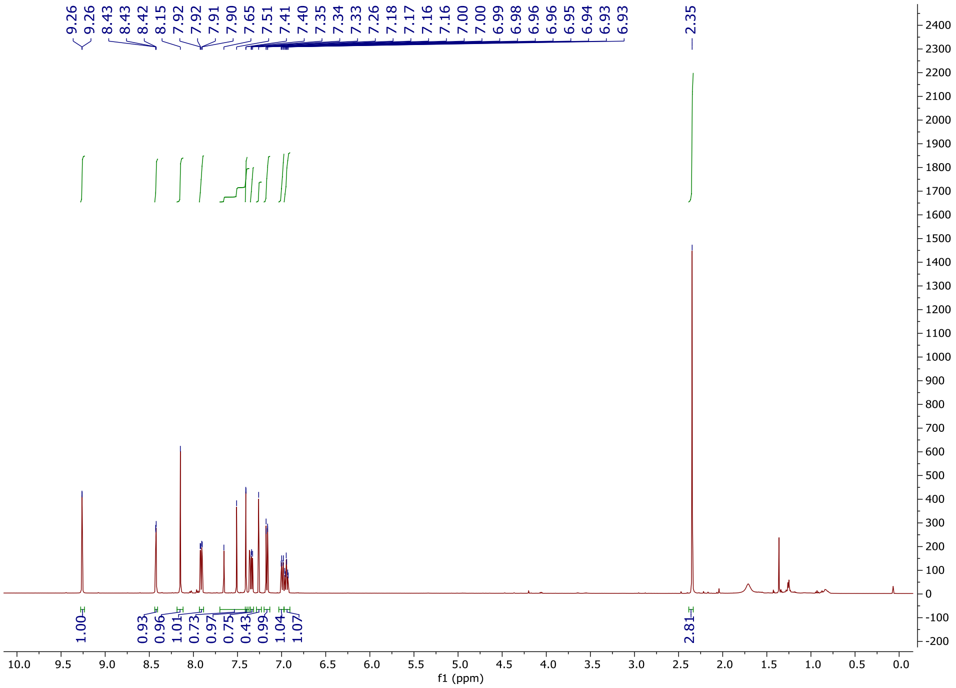

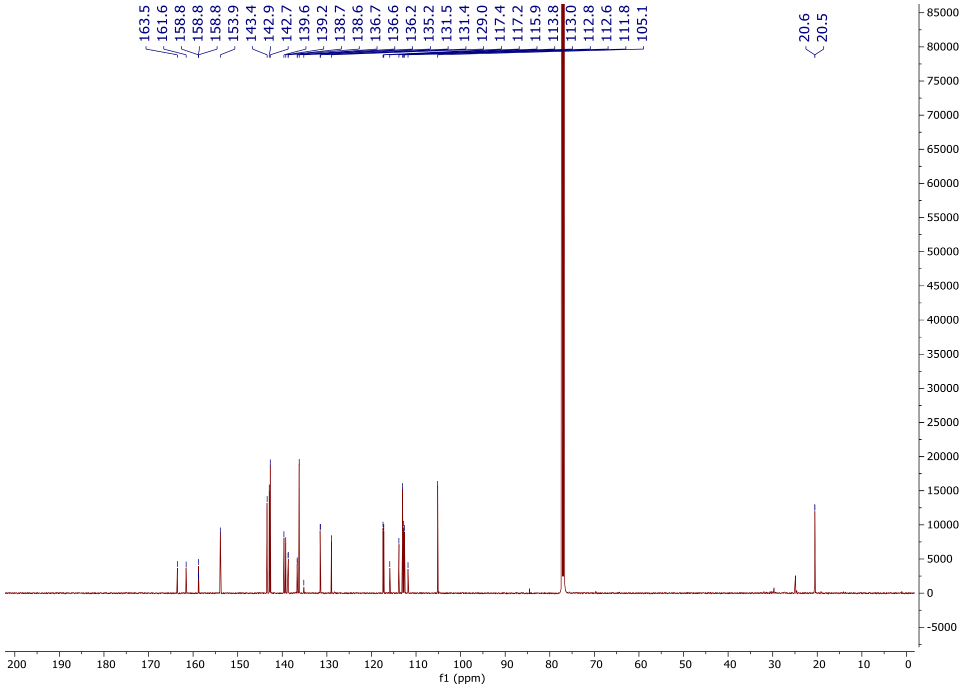
163.5
161.6
158.8
158.8
153.9
143.4
142.9
142.7
139.6
139.2
138.7
138.6
136.7
136.6
136.2
135.2
131.5
131.4
129.0
117.4
117.2
115.9
113.8
113.0
112.8
112.6
111.8
105.1
20.6
20.5
f1 (ppm)

**6-(4-Fluoro-2-methylphenyl)-1-(5-methoxypyridin-2-yl)-1*H*-imidazo[4,5-*c*]pyridine (OXS008495)**

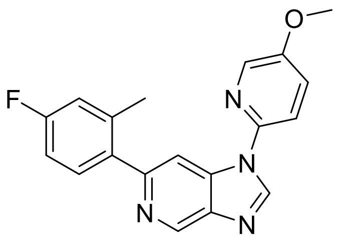

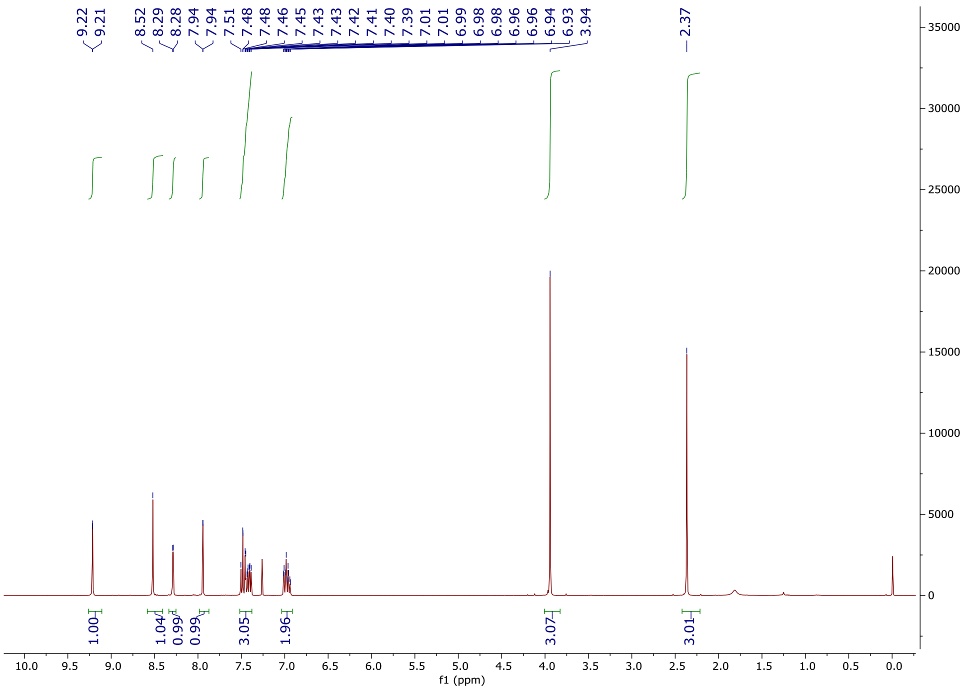

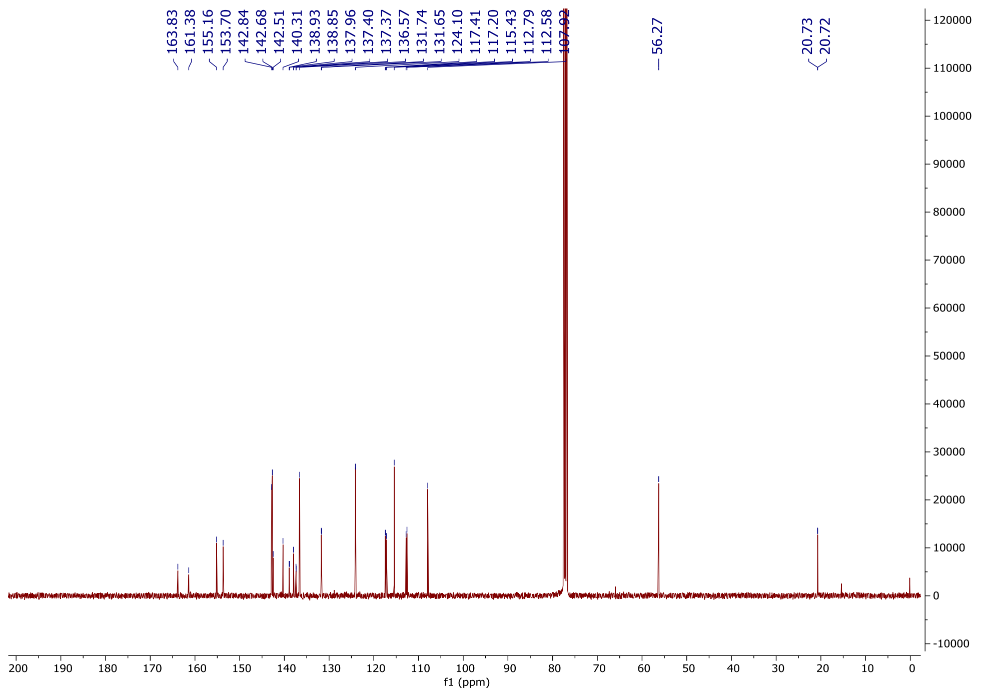
163.83
161.38
155.16
153.70
142.84
142.68
142.51
140.31
138.93
138.85
137.96
137.40
137.37
136.57
131.74
131.65
124.10
117.41
117.20
115.43
112.79
112.58
107.92
56.27
20.73
20.72
f1 (ppm)

**6-(4-Fluoro-2-methylphenyl)-1-(5-methoxypyrazin-2-yl)-1*H*-imidazo[4,5-*c*]pyridine (OXS007837)**

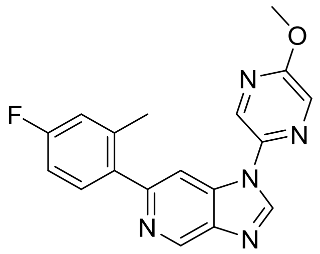

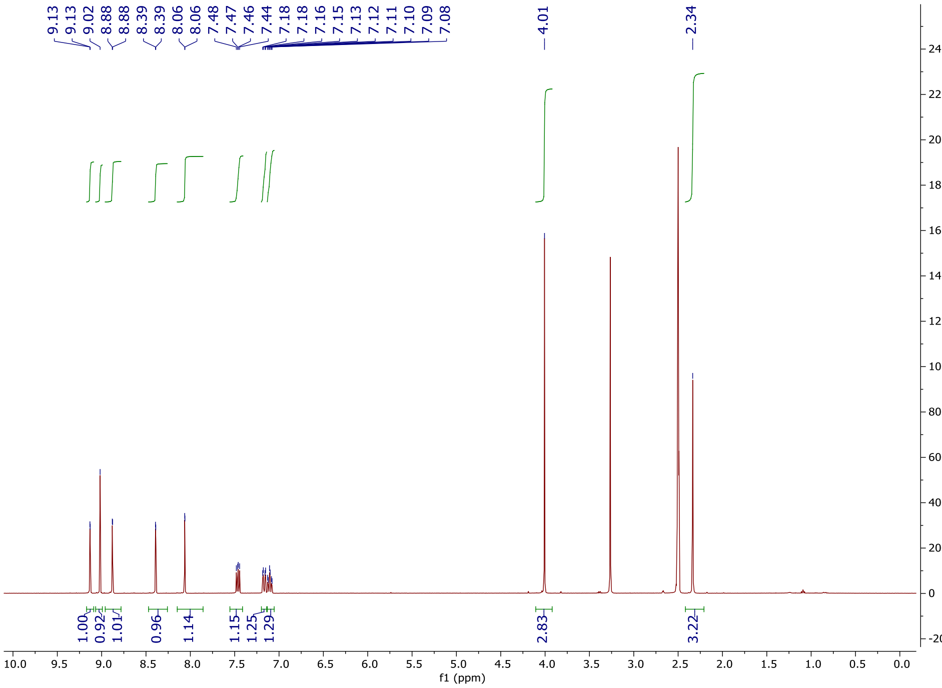

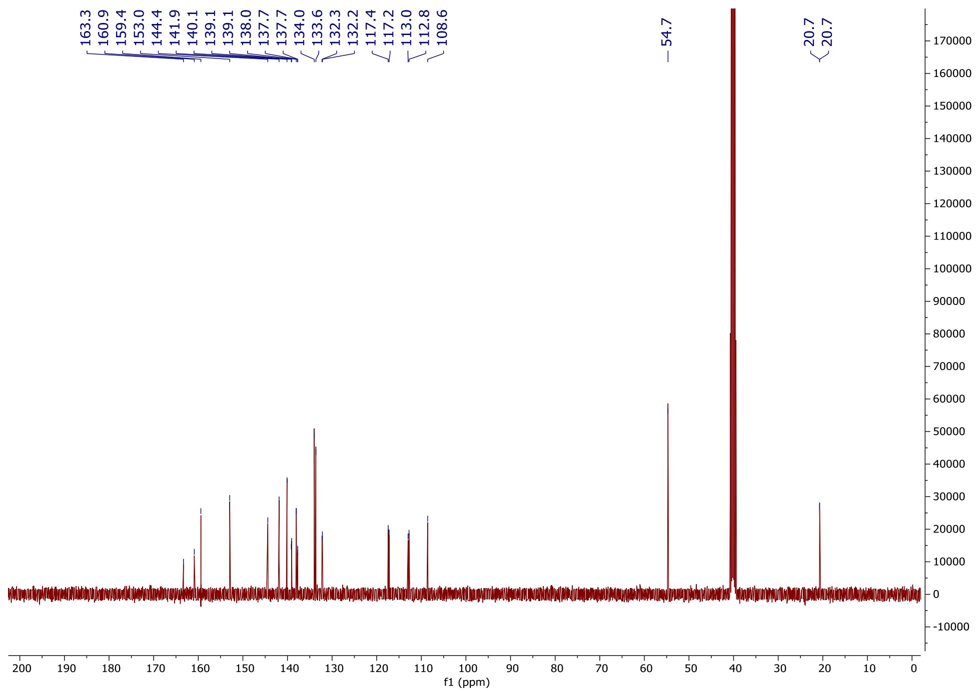

163.3
160.9
159.4
153.0
144.4
141.9
140.1
139.1
139.1
138.0
137.7
137.7
134.0
133.6
132.3
132.2
117.4
117.2
113.0
112.8
108.6
54.7
20.7
20.7
f1 (ppm)

**6-(4-Fluoro-2-methylphenyl)-1-(2-methoxypyrimidin-5-yl)-1*H*-imidazo[4,5-*c*]pyridine (OXS007785)**

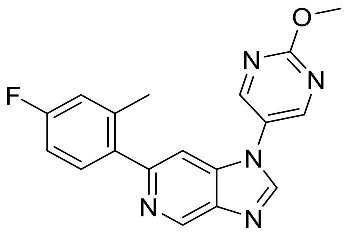

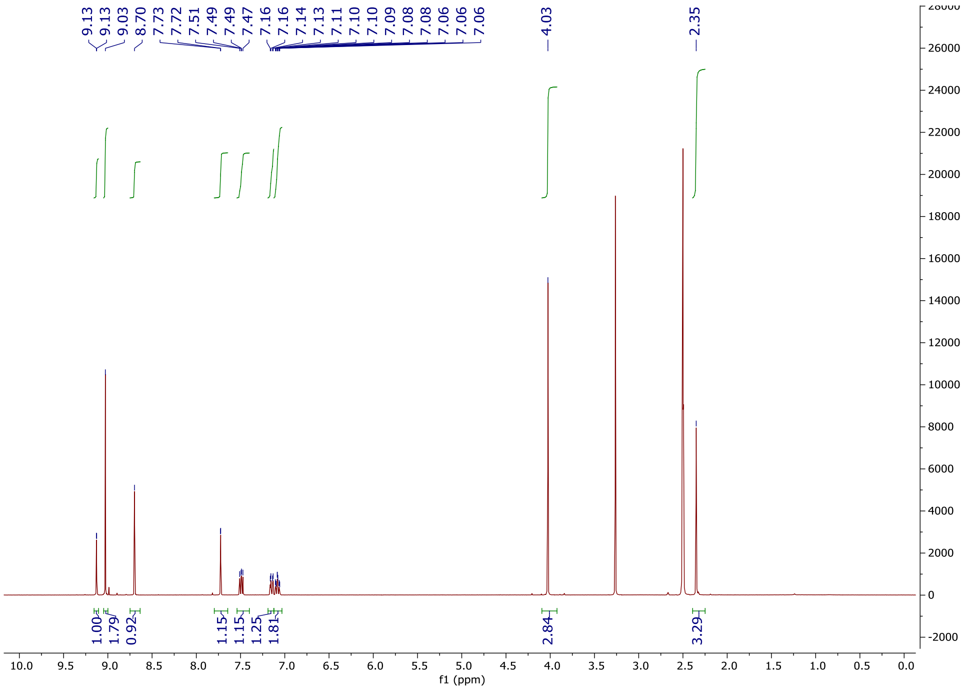

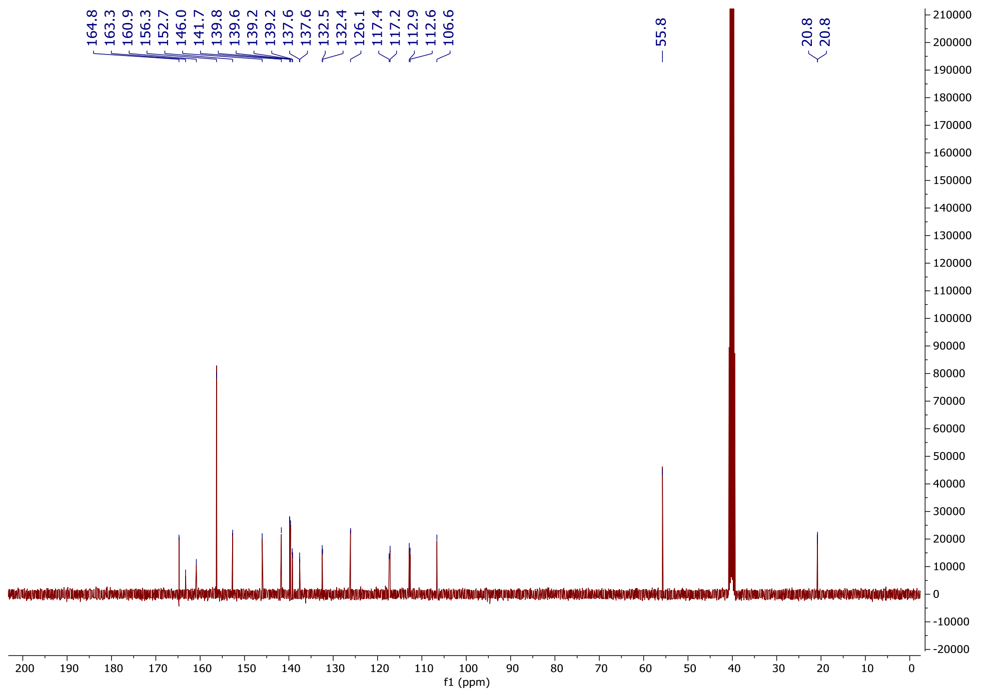

164.8
163.3
160.9
156.3
152.7
146.0
141.7
139.8
139.6
139.2
139.2
137.6
137.6
132.5
132.4
126.1
117.4
117.2
112.9
112.6
106.6
55.8
20.8
20.8
f1 (ppm)

**1-(Cyclopropylmethyl)-6-(4-fluoro-2-methylphenyl)-1*H*-imidazo[4,5-*c*]pyridine (OXS007812)**

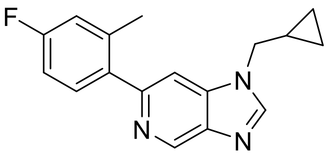

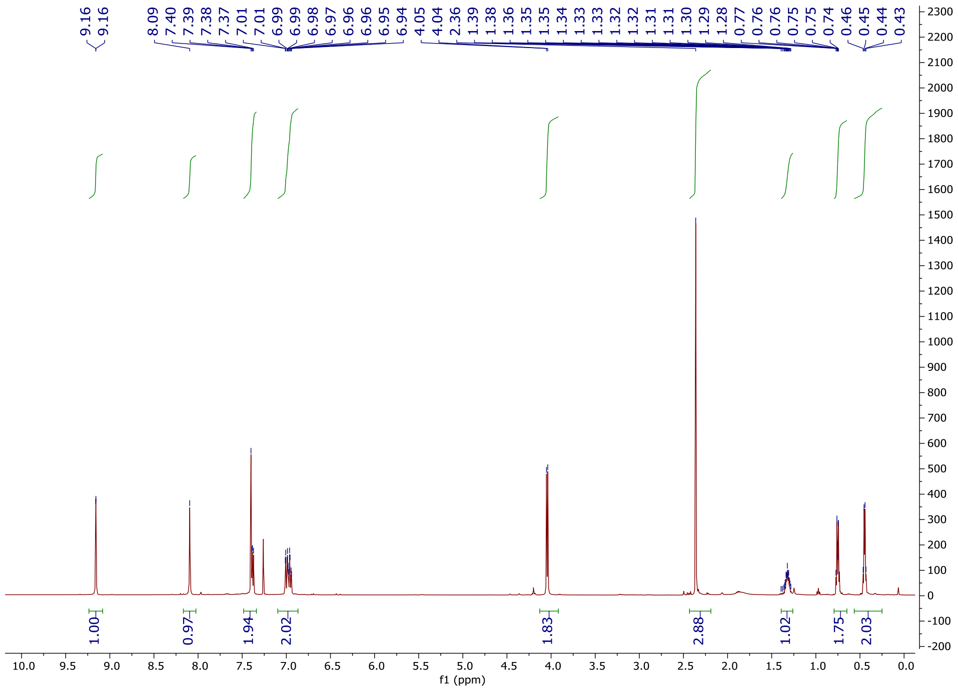

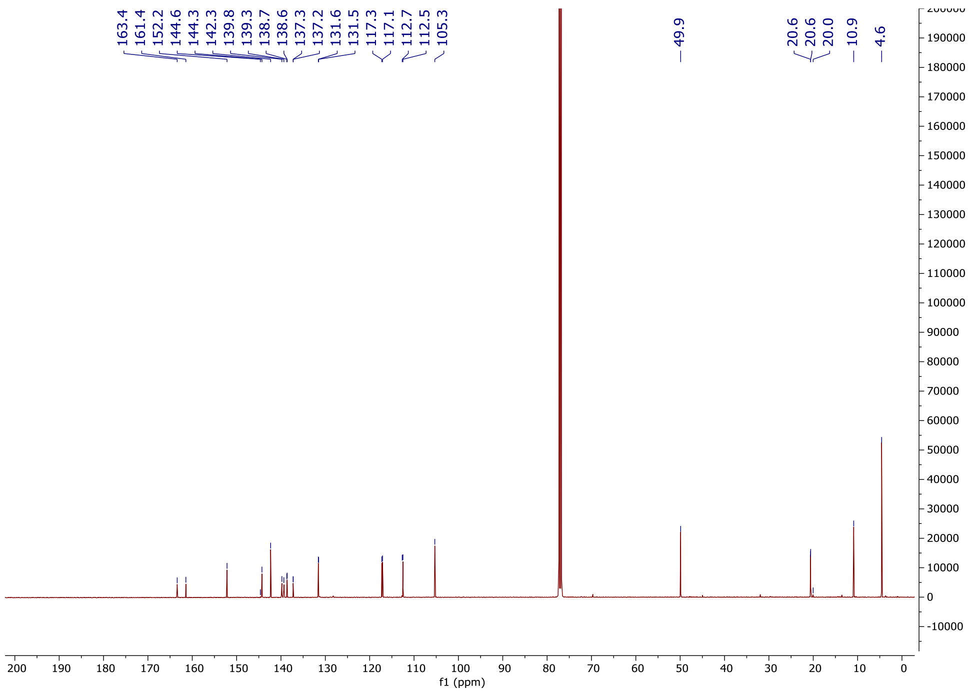

163.4
161.4
152.2
144.6
144.3
142.3
139.8
139.3
138.7
138.6
137.3
137.2
131.6
131.5
117.3
117.1
112.7
112.5
105.3
49.9
20.6
20.6
20.0
10.9
4.6
f1 (ppm)

**6-(4-Fluoro-2-methylphenyl)-1-(1-methyl-1*H*-indazol-5-yl)-1*H*-imidazo[4,5-*c*]pyridine (OXS007570)**

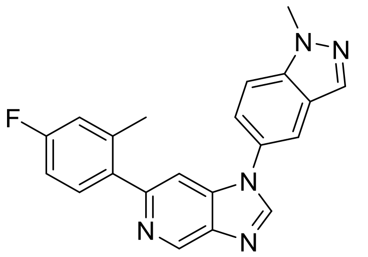

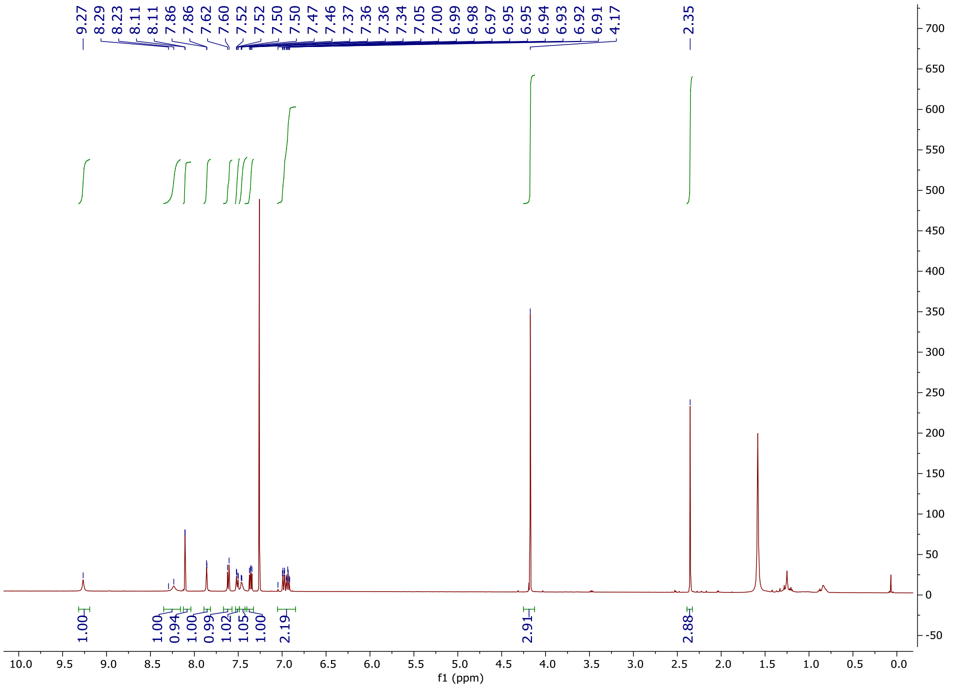

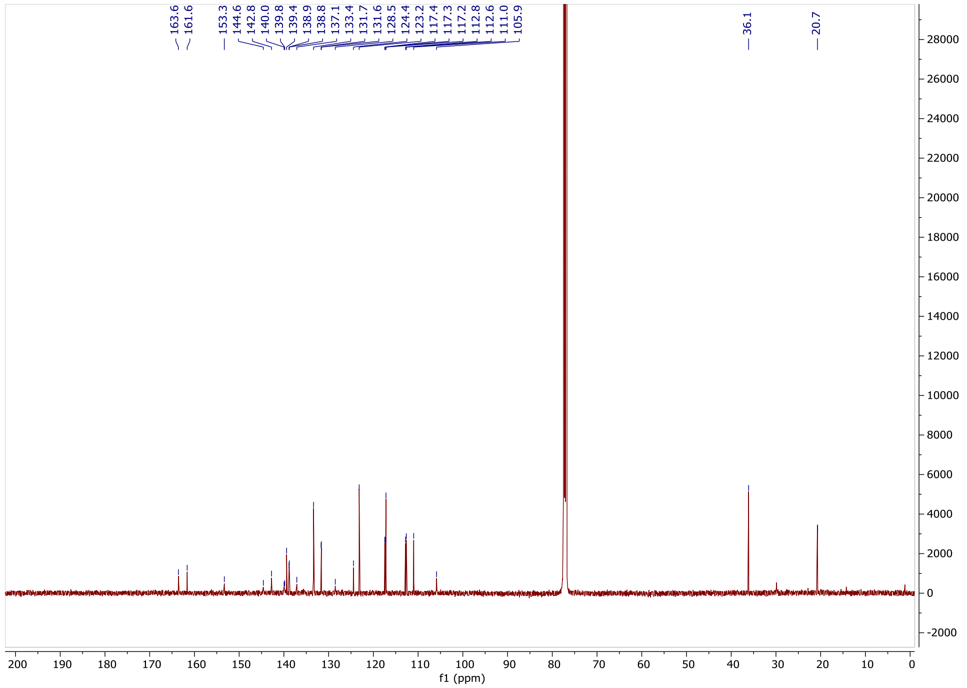

163.6
161.6
153.3
144.6
142.8
140.0
139.8
139.4
138.9
138.8
137.1
133.4
131.7
131.6
128.5
124.4
123.2
117.4
117.3
117.2
112.8
112.6
111.0
105.9
36.1
20.7
f1 (ppm)

**1-(4-(Difluoromethoxy)phenyl)-6-(4-fluorophenyl)-1*H*-imidazo[4,5-*c*]pyridine (OXS008278)**

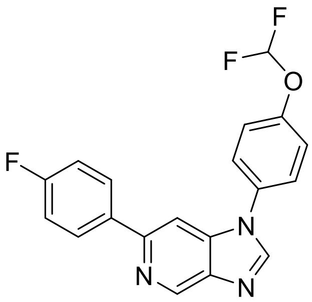
F
F
O
F
N
N
N

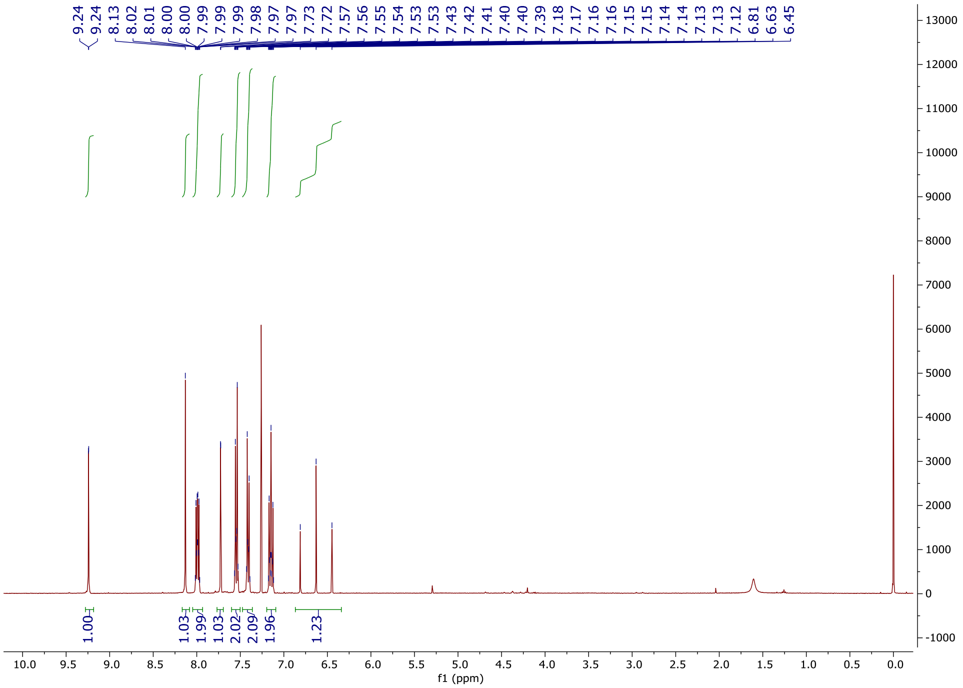
9.24
9.24
8.13
8.02
8.01
8.00
7.99
7.99
7.99
7.98
7.97
7.97
7.73
7.72
7.57
7.56
7.55
7.54
7.53
7.53
7.43
7.42
7.41
7.40
7.39
7.18
7.17
7.16
7.16
7.15
7.15
7.14
7.14
7.13
7.13
7.12
6.81
6.63
6.45
1.00
1.03
1.99
1.03
2.02
2.09
1.96
1.23
f1 (ppm)

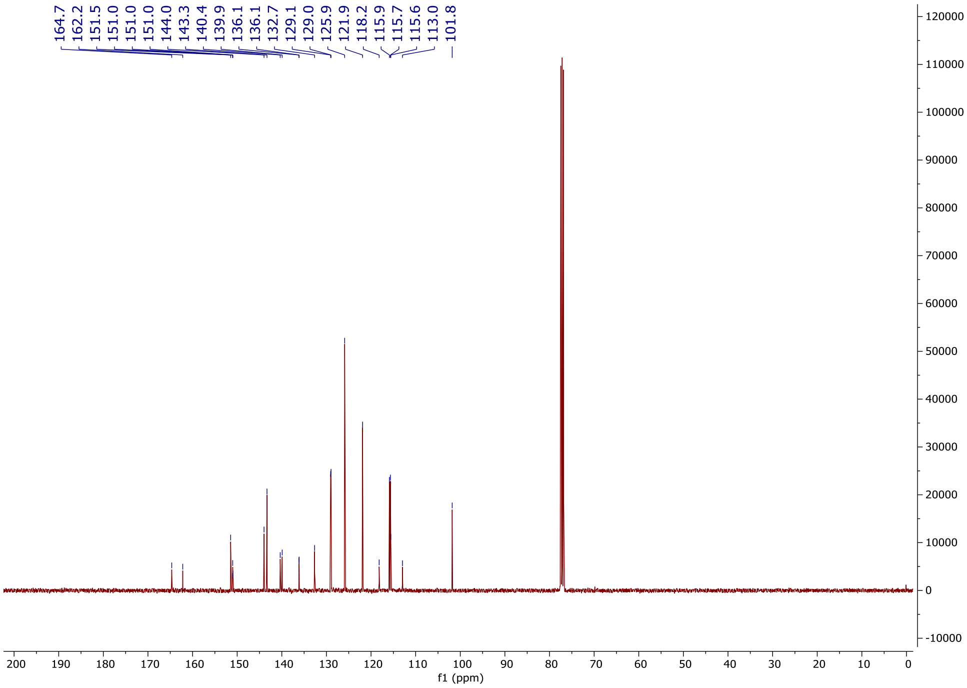
164.7
162.2
151.5
151.0
151.0
151.0
144.0
143.3
140.4
139.9
136.1
136.1
132.7
129.1
129.0
125.9
121.9
118.2
115.9
115.7
115.6
113.0
101.8
f1 (ppm)

**1-[4-(Difluoromethoxy)phenyl]-6-(4-fluoro-2-methoxyphenyl)-1H-imidazo[4,5-c]pyridine (OXS007739)**

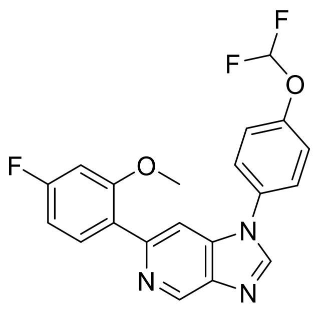

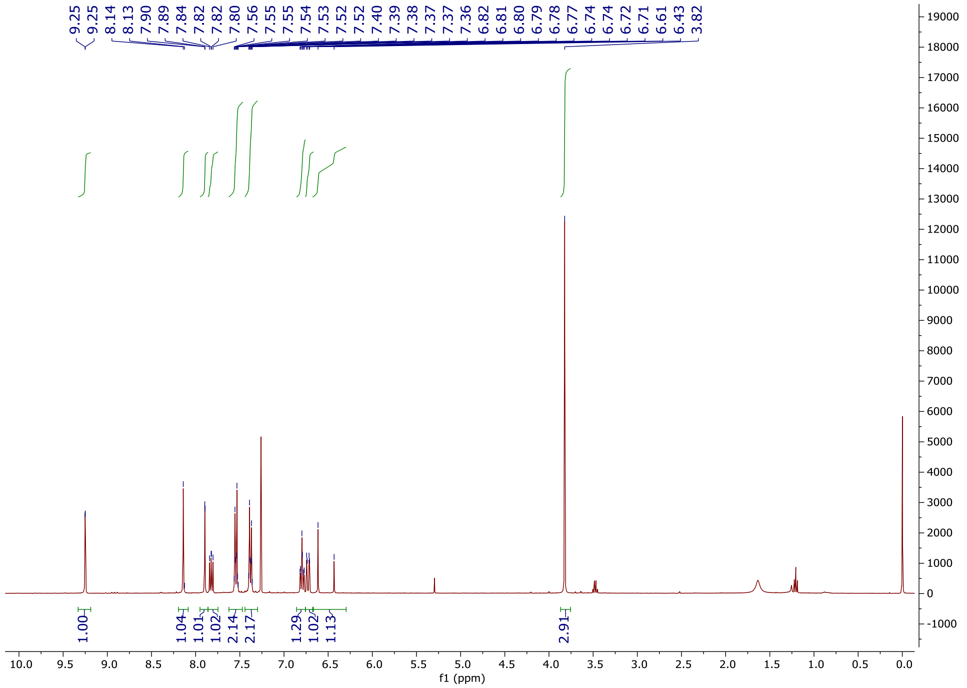
9.25
9.25
8.14
8.13
7.90
7.89
7.84
7.82
7.82
7.80
7.56
7.55
7.55
7.54
7.53
7.52
7.52
7.40
7.39
7.38
7.37
7.37
7.36
6.82
6.81
6.80
6.79
6.78
6.77
6.74
6.74
6.72
6.71
6.61
6.43
3.82
1.00
1.04
1.01
1.02
2.14
2.17
1.29
1.02
1.13
2.91
f1 (ppm)

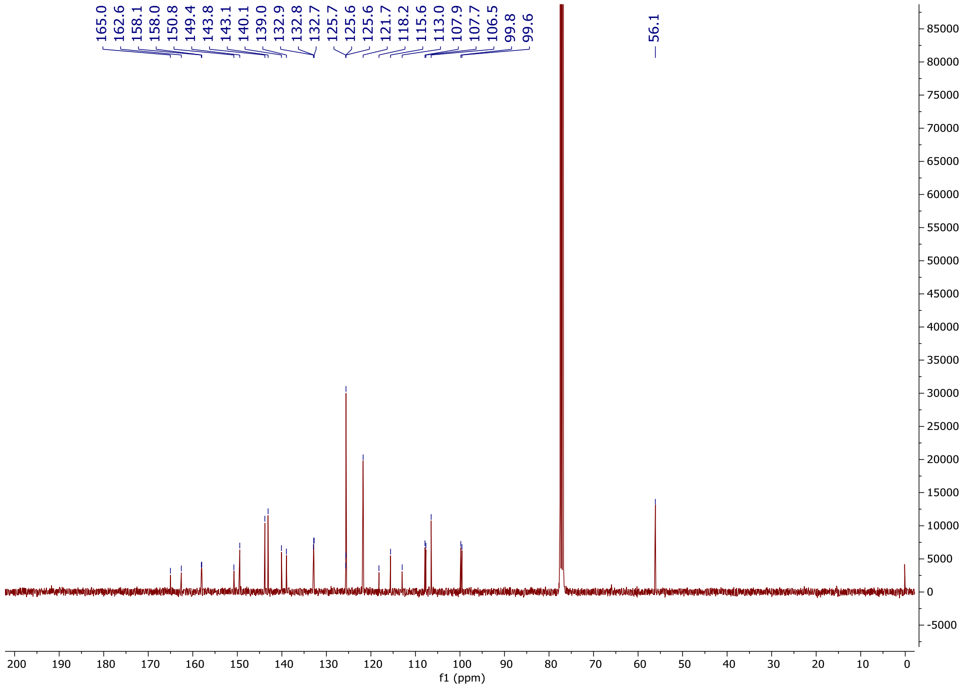
165.0
162.6
158.1
158.0
150.8
149.4
143.8
143.1
140.1
139.0
132.9
132.8
132.7
125.7
125.6
125.6
121.7
118.2
115.6
113.0
107.9
107.7
106.5
99.8
99.6
56.1
f1 (ppm)

**1-[4-(Difluoromethoxy)phenyl]-6-(2,6-dimethylphenyl)imidazo[4,5-c]pyridine (OXS007924)**

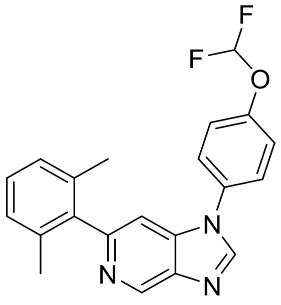

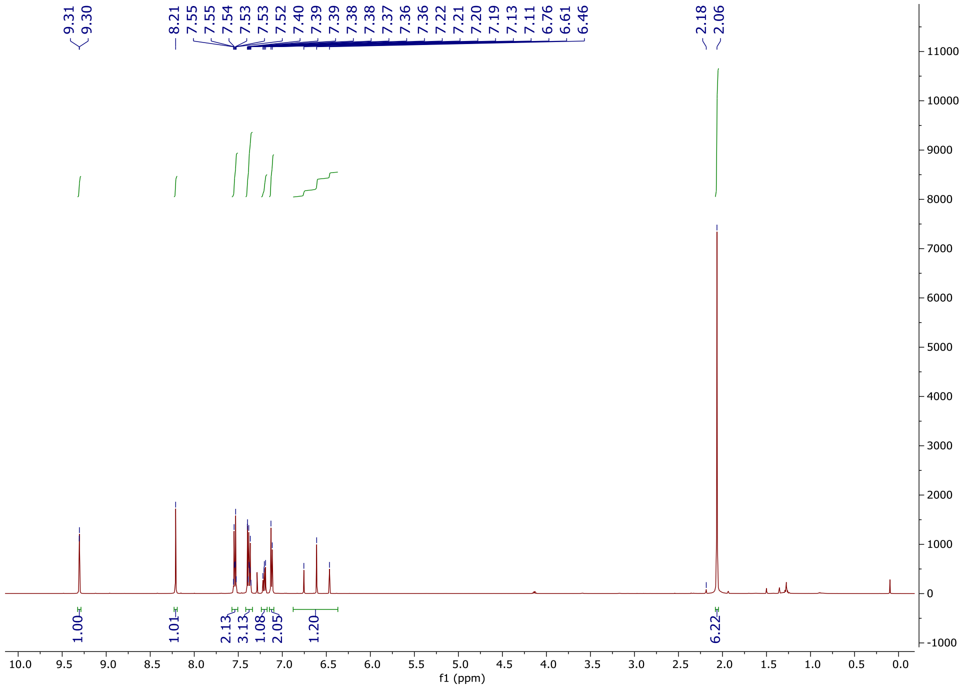
9.31
9.30
8.21
7.55
7.55
7.54
7.53
7.53
7.52
7.40
7.39
7.39
7.38
7.38
7.37
7.36
7.36
7.22
7.21
7.20
7.19
7.13
7.11
6.76
6.61
6.46
2.18
2.06
1.00
1.01
2.13
3.13
1.08
2.05
1.20
6.22
f1 (ppm)

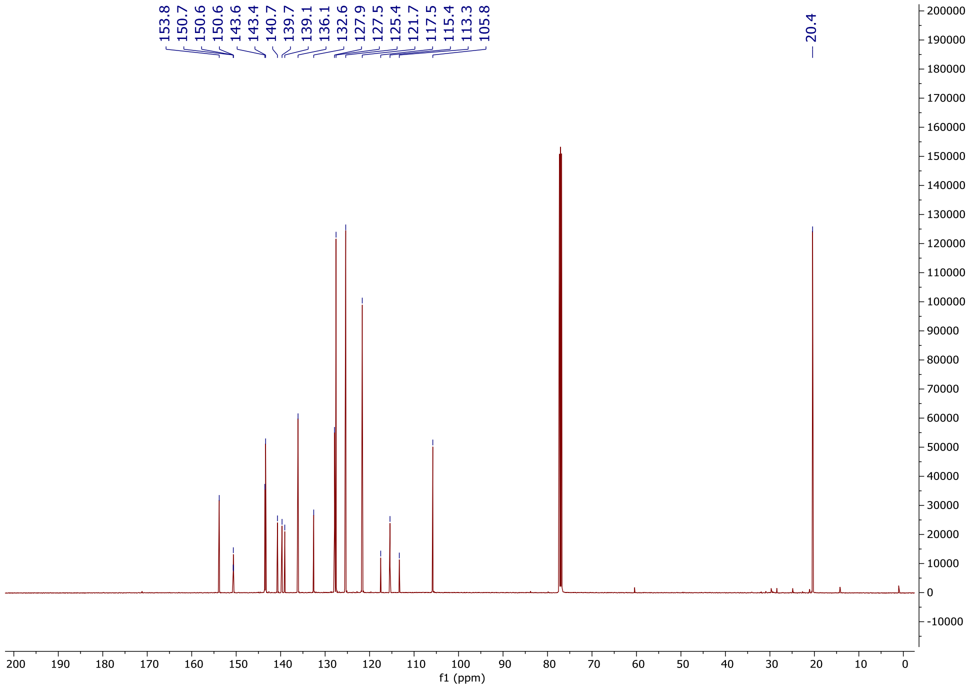
153.8
150.7
150.6
150.6
143.6
143.4
140.7
139.7
139.1
136.1
132.6
127.9
127.5
125.4
121.7
117.5
115.4
113.3
105.8
20.4
f1 (ppm)

**6-(2-Chloro-4-fluorophenyl)-1-(4-(difluoromethoxy)phenyl)-1*H*-imidazo[4,5-*c*]pyridine (OXS008296)**

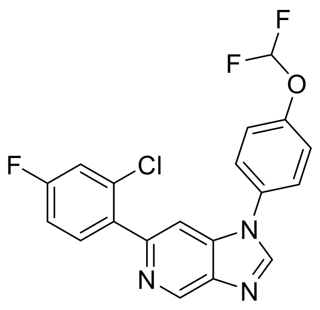

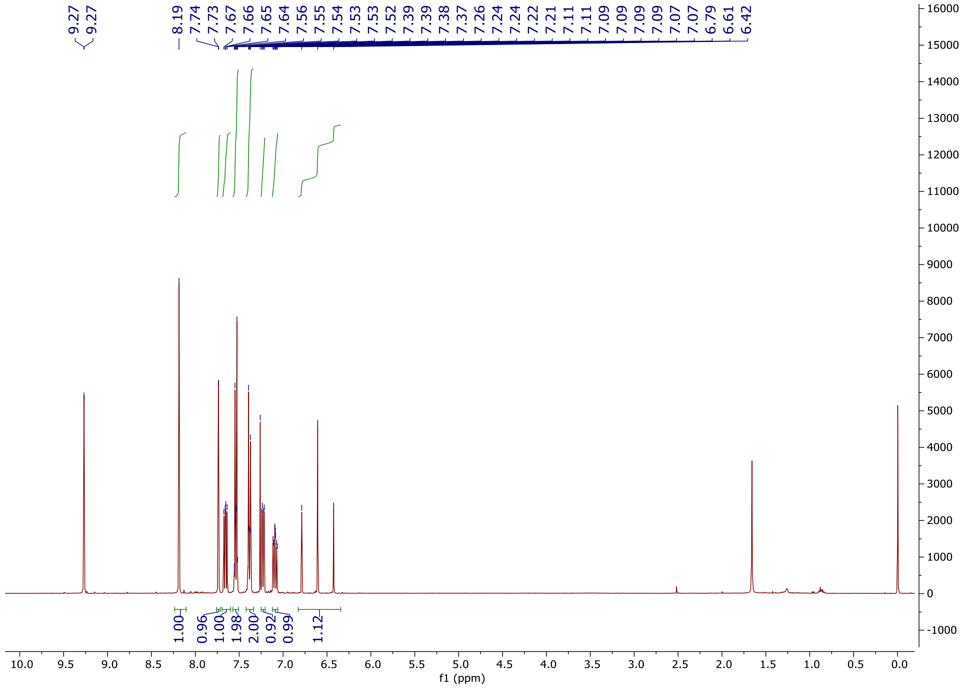
9.27
9.27
8.19
7.74
7.73
7.67
7.66
7.65
7.64
7.56
7.55
7.54
7.53
7.52
7.39
7.39
7.38
7.37
7.26
7.24
7.24
7.22
7.21
7.11
7.11
7.09
7.09
7.09
7.07
7.07
6.79
6.61
6.42
1.00
0.96
1.00
1.98
2.00
0.92
0.99
1.12
f1 (ppm)

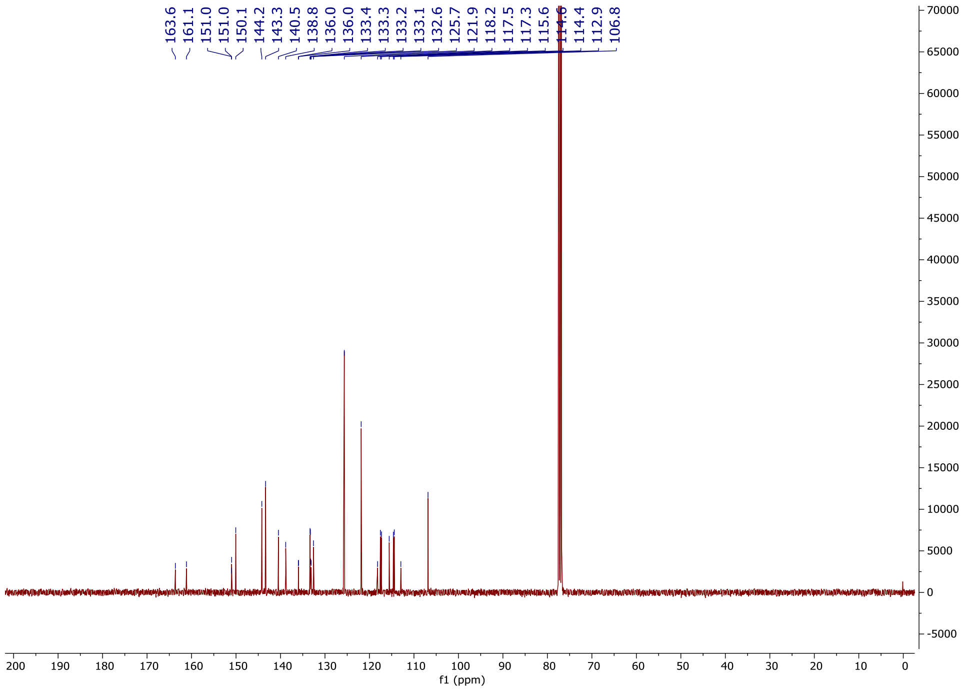
163.6
161.1
151.0
151.0
150.1
144.2
143.3
140.5
138.8
136.0
136.0
133.4
133.3
133.2
133.1
132.6
125.7
121.9
118.2
117.5
117.3
115.6
114.6
114.4
112.9
106.8
f1 (ppm)

**1-(4-(Difluoromethoxy)phenyl)-6-(4-fluoro-2-(trifluoromethyl)phenyl)-1*H*-imidazo[4,5-*c*]pyridine (OXS008316)**

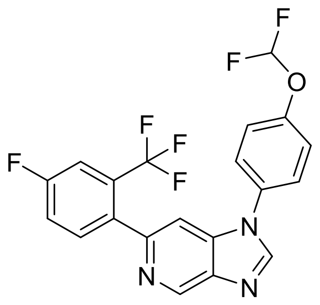
F
F
O
F
F
F
F
N
N
N

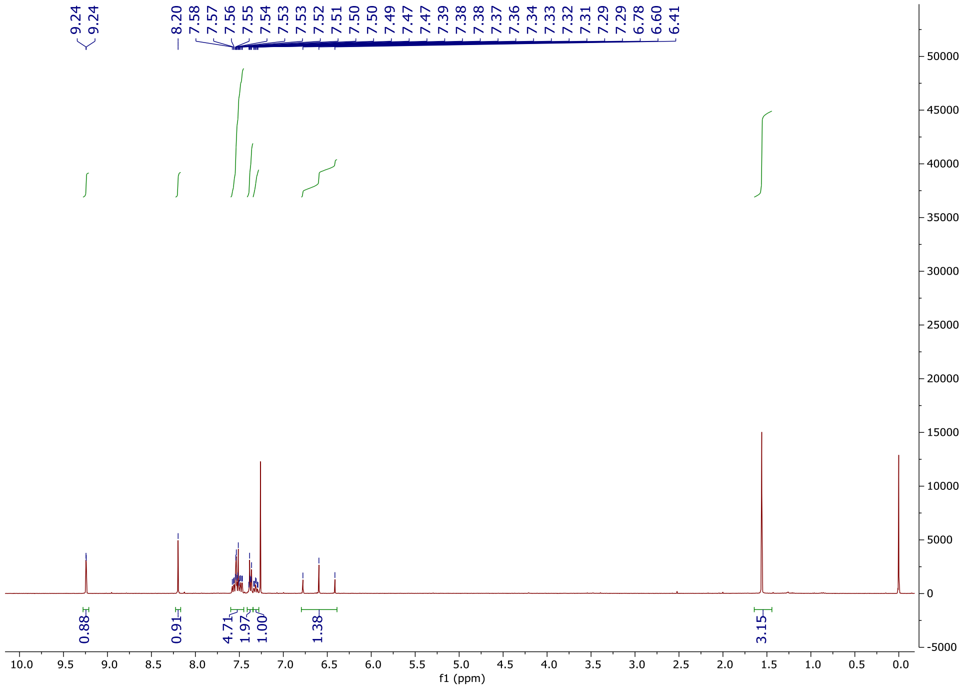
9.24
9.24
8.20
7.58
7.57
7.56
7.55
7.54
7.53
7.53
7.52
7.51
7.50
7.50
7.49
7.47
7.47
7.39
7.38
7.38
7.37
7.36
7.34
7.33
7.32
7.31
7.29
7.29
6.78
6.60
6.41
0.88
0.91
4.71
1.97
1.00
1.38
3.15
f1 (ppm)

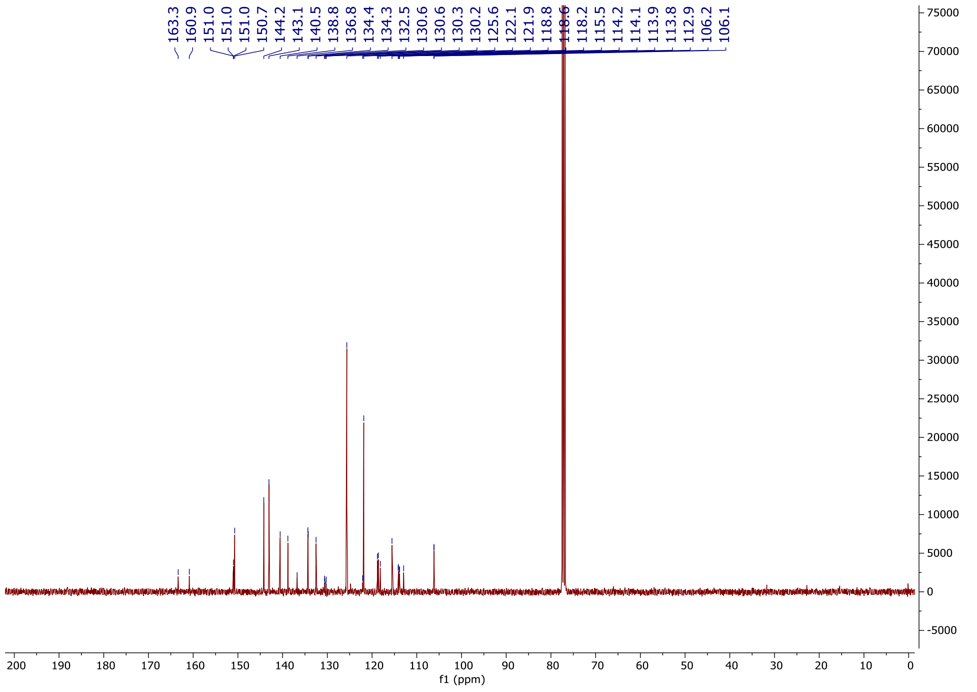
163.3
160.9
151.0
151.0
150.7
144.2
143.1
140.5
138.8
136.8
134.4
134.3
132.5
130.6
130.3
130.2
125.6
122.1
121.9
118.8
118.6
118.2
115.5
114.2
114.1
113.9
113.8
112.9
106.2
106.1
f1 (ppm)

### 1-(4-(Difluoromethoxy)phenyl)-6-(2-(trifluoromethoxy)phenyl)-1*H*-imidazo[4,5-*c*]pyridine (OXS008297)

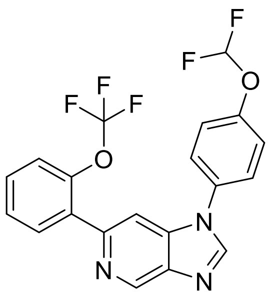

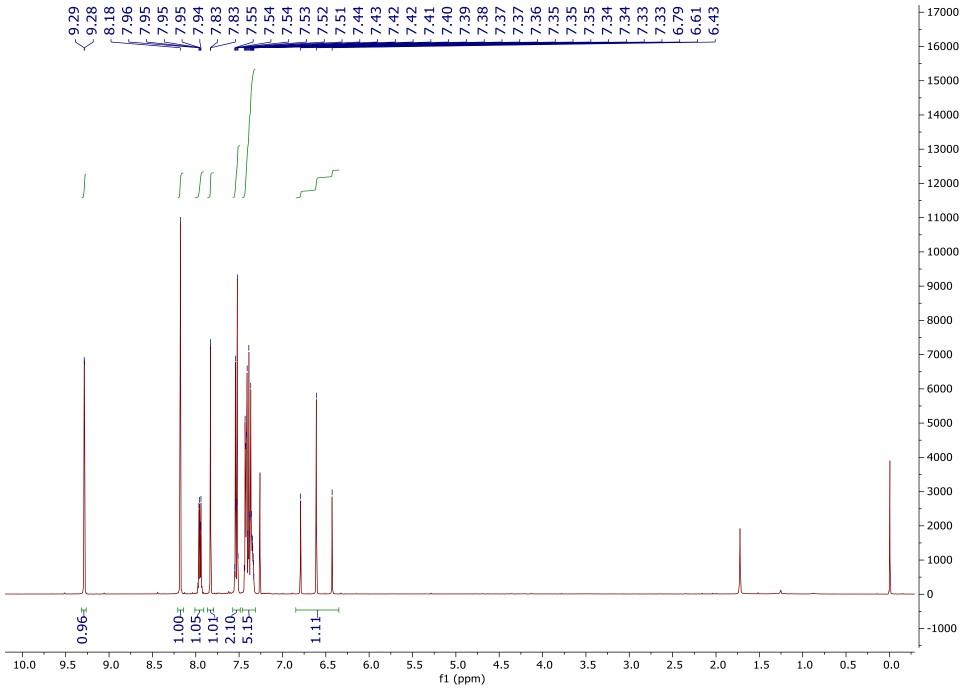

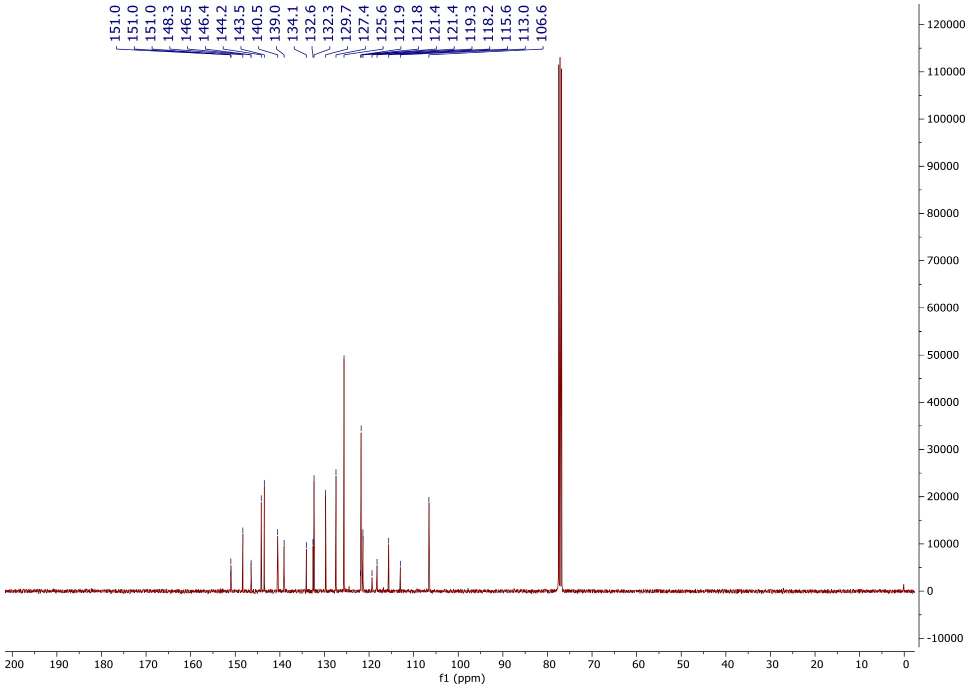
151.0
151.0
151.0
148.3
146.5
146.4
144.2
143.5
140.5
139.0
134.1
132.6
132.3
129.7
127.4
125.6
121.9
121.8
121.4
121.4
119.3
118.2
115.6
113.0
106.6
f1 (ppm)

### 1-(4-(Difluoromethoxy)phenyl)-6-(2-ethylphenyl)-1*H*-imidazo[4,5-*c*]pyridine (OXS007928)

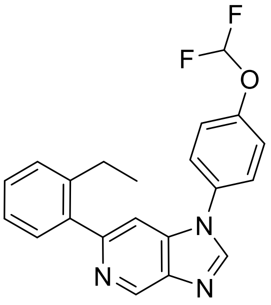

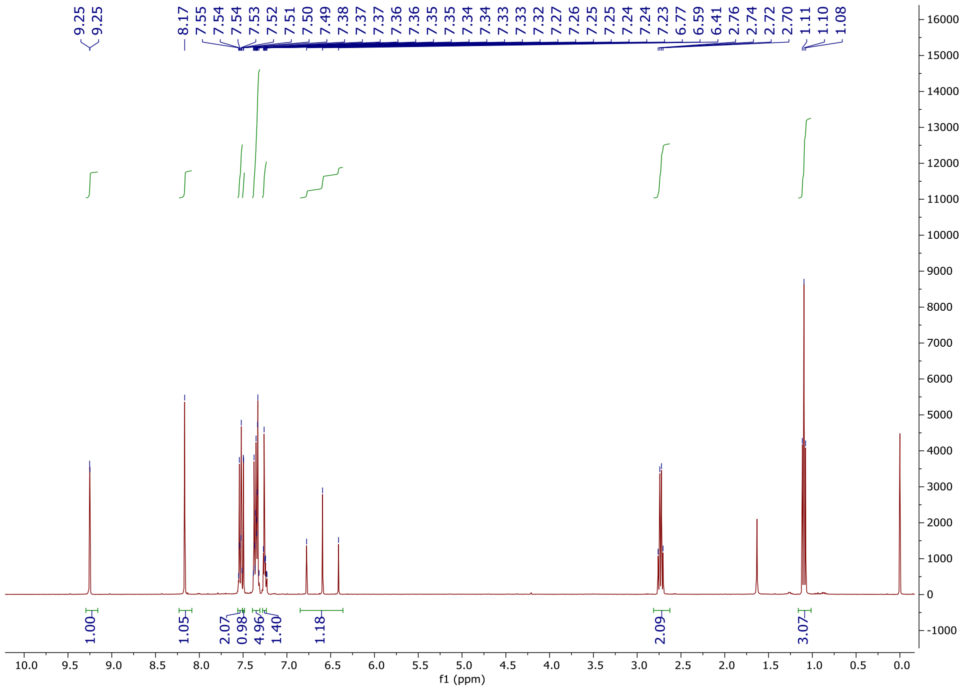

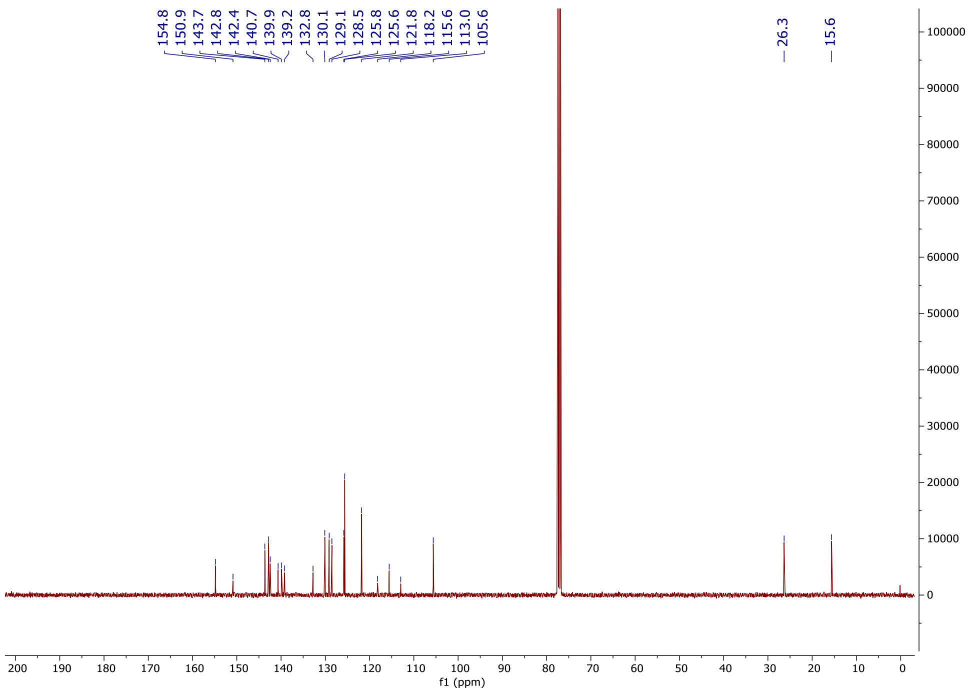
154.8
150.9
143.7
142.8
142.4
140.7
139.9
139.2
132.8
130.1
129.1
128.5
125.8
125.6
121.8
118.2
115.6
113.0
105.6
26.3
15.6
100000
90000
80000
70000
60000
50000
40000
30000
20000
10000
0
200 190 180 170 160 150 140 130 120 110 100 90 80 70 60 50 40 30 20 10 0
f1 (ppm)

**1-(4-(Difluoromethoxy)phenyl)-6-(4-fluoro-2-propoxyphenyl)-1*H*-imidazo[4,5-*c*]pyridine (OXS008318)**

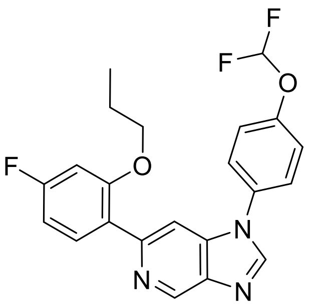
F
F
O
F
O
N
N
N

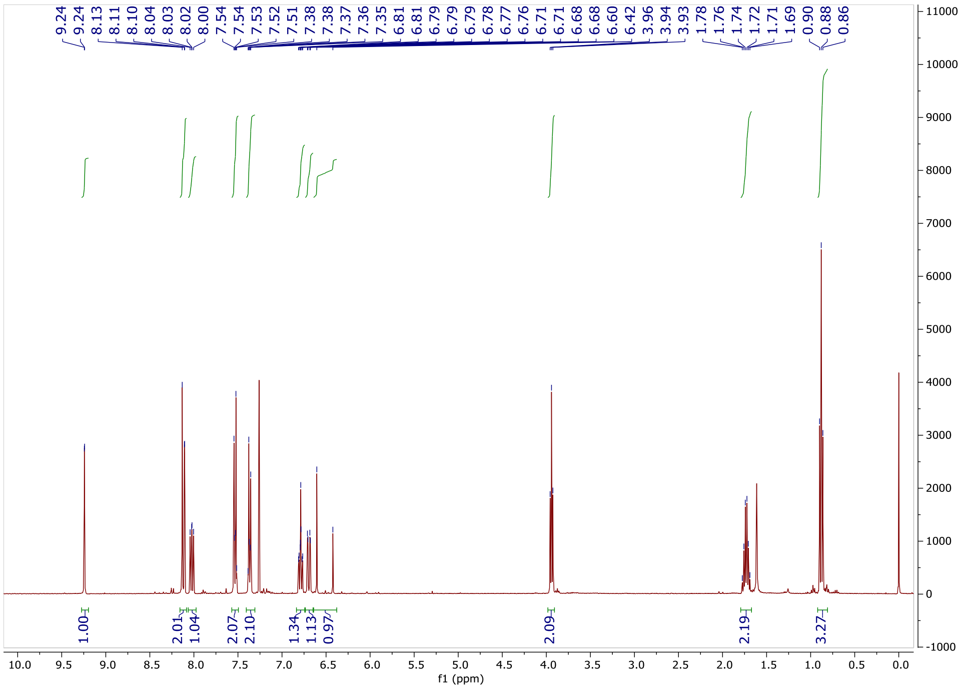
9.24 9.24 8.13 8.11 8.10 8.04 8.03 8.02 8.00 7.54 7.54 7.53 7.52 7.51 7.38 7.38 7.37 7.36 7.35 6.81 6.81 6.79 6.79 6.79 6.78 6.77 6.76 6.71 6.71 6.68 6.68 6.60 6.42 3.96 3.94 3.93 1.78 1.76 1.74 1.72 1.71 1.69 0.90 0.88 0.86
1.00 2.01 1.04 2.07 2.10 1.34 1.13 0.97 2.09 2.19 3.27
f1 (ppm)

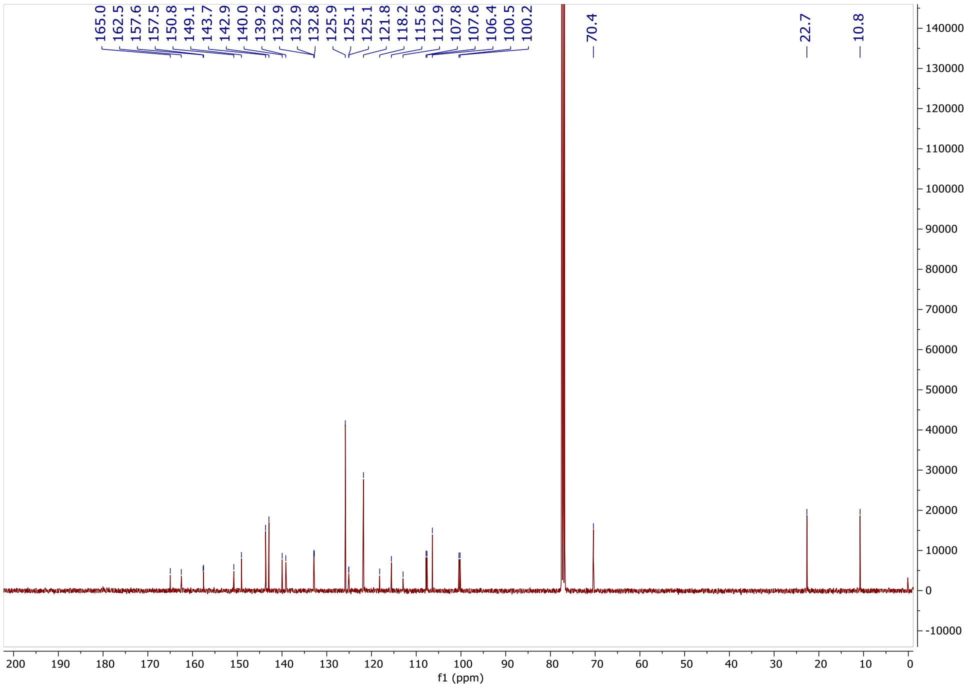
165.0 162.5 157.6 157.5 150.8 149.1 143.7 142.9 140.0 139.2 132.9 132.9 132.8 125.9 125.1 125.1 121.8 118.2 115.6 112.9 107.8 107.6 106.4 100.5 100.2
70.4
22.7
10.8
f1 (ppm)

**1-(4-(Difluoromethoxy)phenyl)-6-(4-fluoro-2-isopropoxyphenyl)-1*H*-imidazo[4,5-*c*]pyridine (OXS008317)**

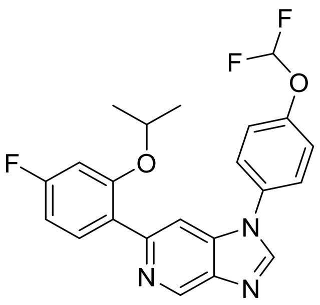

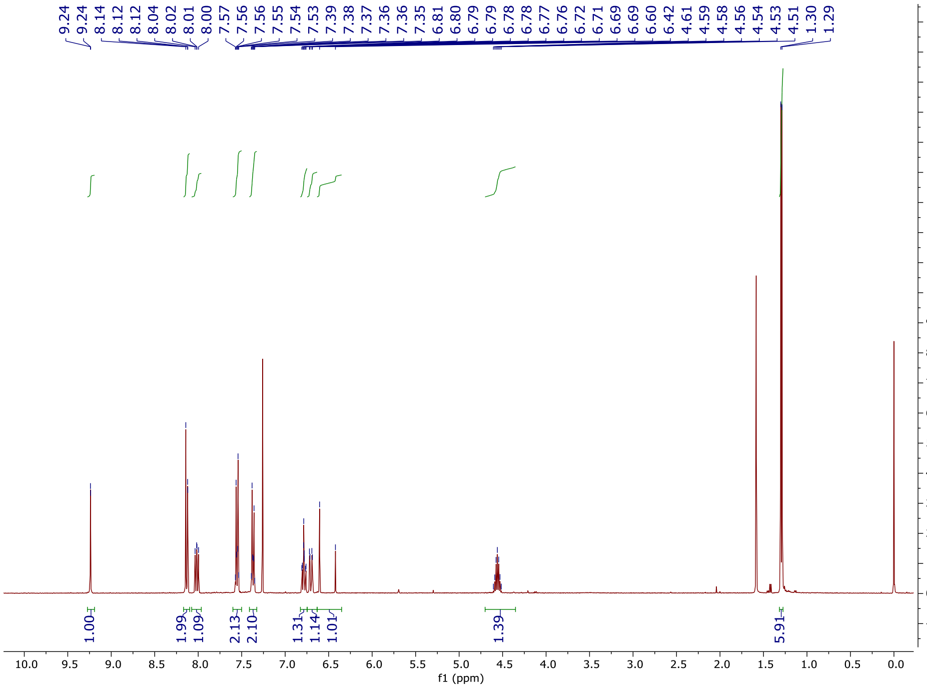
9.24
9.24
8.14
8.12
8.04
8.02
8.01
8.00
7.57
7.56
7.56
7.55
7.54
7.53
7.39
7.38
7.37
7.36
7.36
7.35
6.81
6.80
6.79
6.79
6.78
6.78
6.77
6.76
6.72
6.71
6.69
6.69
6.60
6.42
4.61
4.59
4.58
4.56
4.54
4.53
4.51
1.30
1.29
1.00
1.99
1.09
2.13
2.10
1.31
1.14
1.01
1.39
5.91
f1 (ppm)

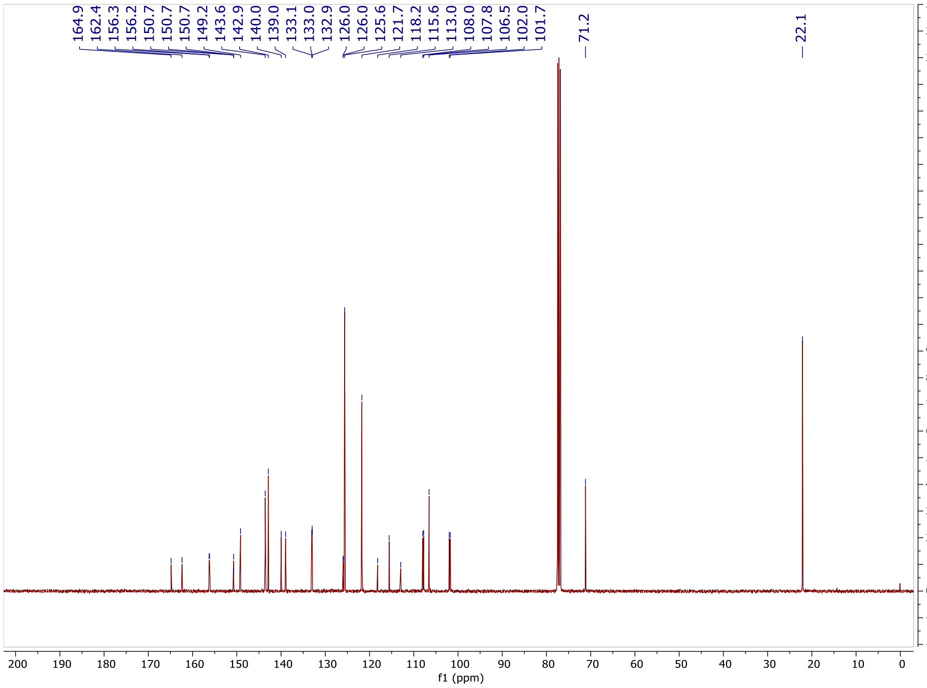
164.9
162.4
156.3
156.2
150.7
150.7
149.2
143.6
142.9
140.0
139.0
133.1
133.0
132.9
126.0
126.0
125.6
121.7
118.2
115.6
113.0
108.0
107.8
106.5
102.0
101.7
71.2
22.1
f1 (ppm)

**1-(4-(Difluoromethoxy)phenyl)-6-(2,4-dimethylphenyl)-1*H*-imidazo[4,5-*c*]pyridine (OXS007929)**

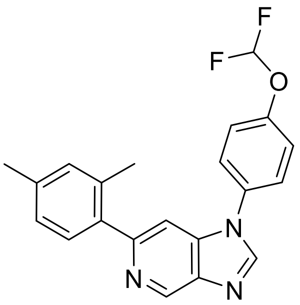

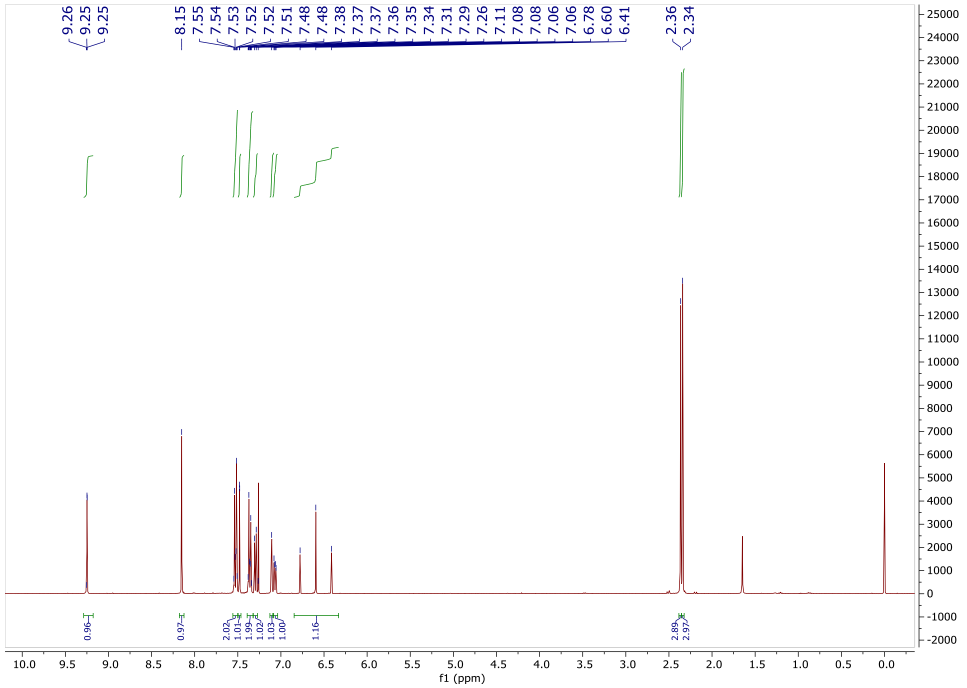

9.26
9.25
9.25
8.15
7.55
7.54
7.53
7.52
7.52
7.51
7.48
7.48
7.38
7.37
7.37
7.36
7.35
7.34
7.31
7.29
7.26
7.11
7.08
7.08
7.06
7.06
6.78
6.60
6.41
2.36
2.34
0.96
0.97
2.02
1.01
1.99
1.02
1.03
1.00
1.16
2.89
2.97
f1 (ppm)

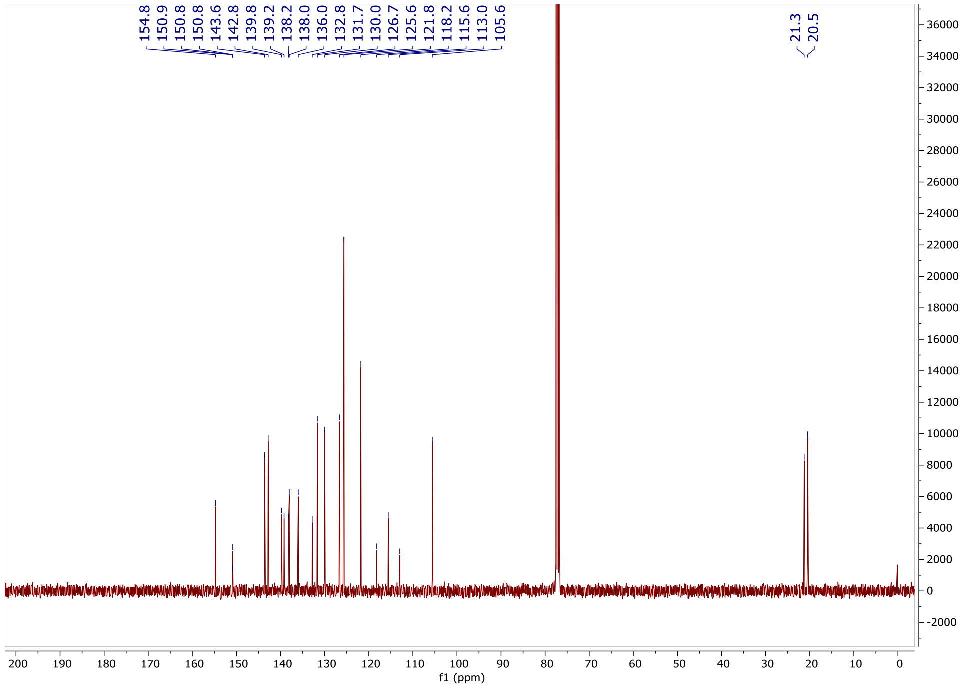

154.8
150.9
150.8
143.6
142.8
139.8
139.2
138.2
138.0
136.0
132.8
131.7
130.0
126.7
125.6
121.8
118.2
115.6
113.0
105.6
21.3
20.5
f1 (ppm)

#### 4-(1-(4-(Difluoromethoxy)phenyl)-1*H*-imidazo[4,5-*c*]pyridin-6-yl)-3-methylbenzonitrile (OXS007556)

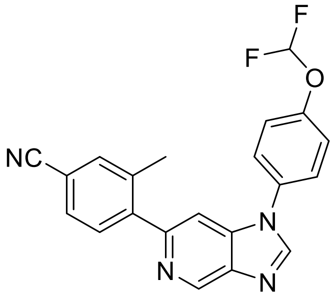

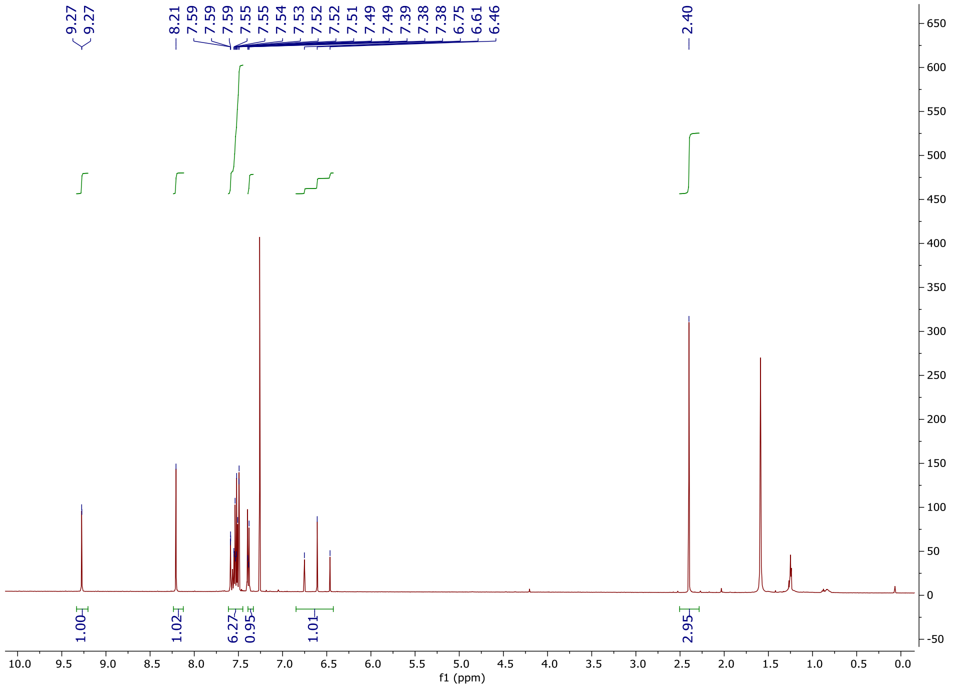

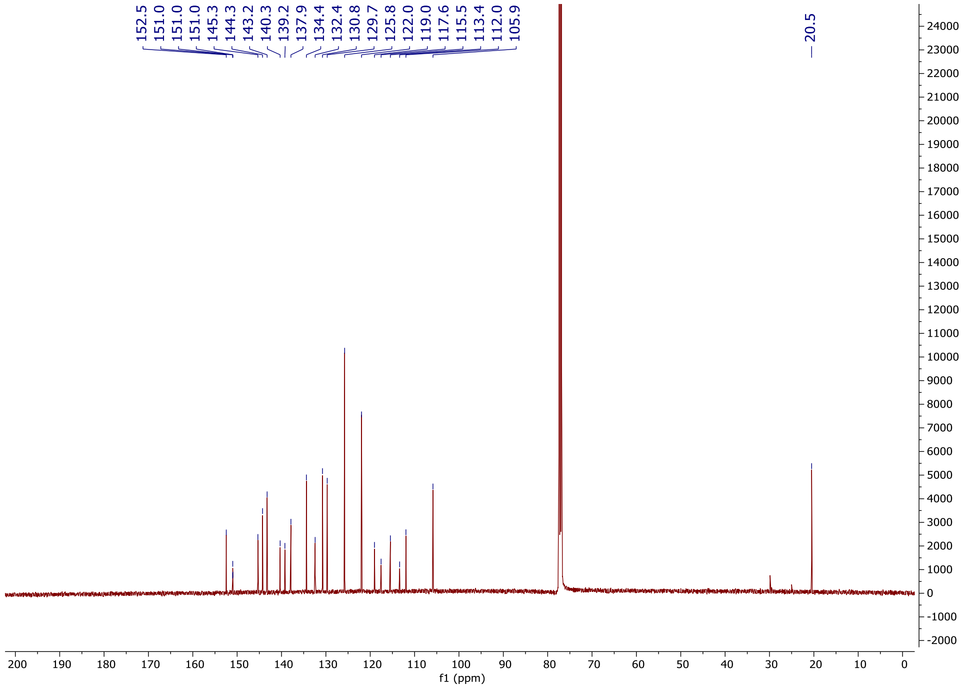

152.5
151.0
151.0
145.3
144.3
143.2
140.3
139.2
137.9
134.4
132.4
130.8
129.7
125.8
122.0
119.0
117.6
115.5
113.4
112.0
105.9
20.5
f1 (ppm)

**6-(4-Chloro-2-methylphenyl)-1-(4-(difluoromethoxy)phenyl)-1*H*-imidazo[4,5-*c*]pyridine (OXS007930)**

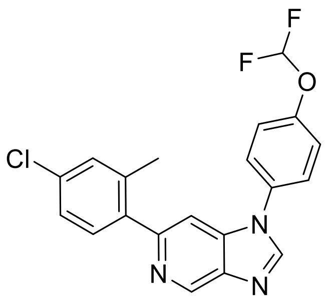

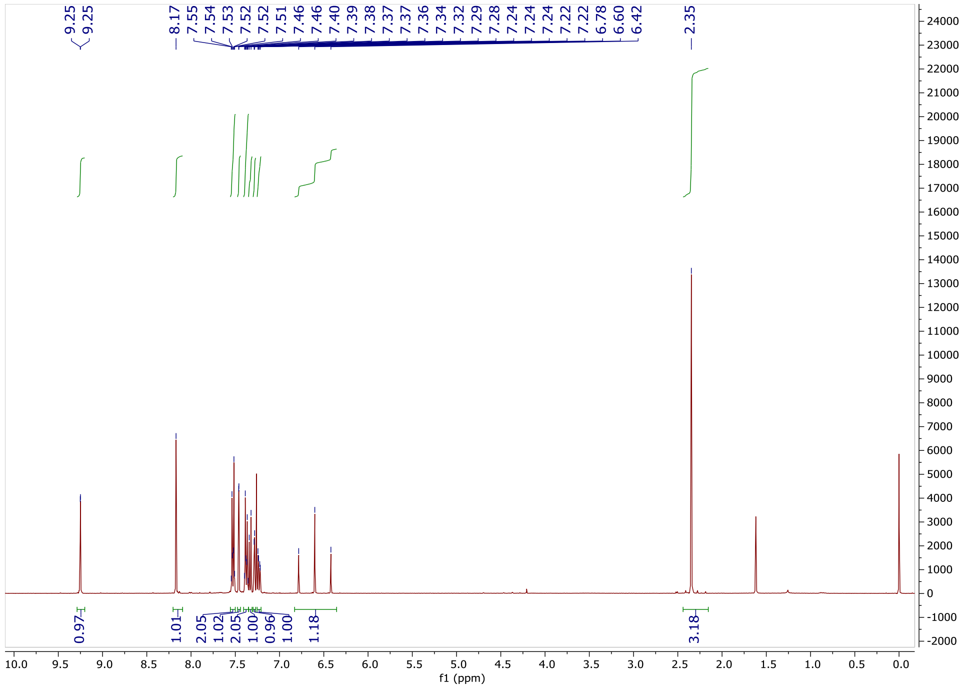
9.25
9.25
8.17
7.55
7.54
7.53
7.52
7.52
7.51
7.46
7.46
7.40
7.39
7.38
7.37
7.37
7.36
7.34
7.32
7.29
7.28
7.24
7.24
7.24
7.22
7.22
6.78
6.60
6.42
2.35
0.97
1.01
2.05
1.02
2.05
1.00
0.96
1.00
1.18
3.18
f1 (ppm)

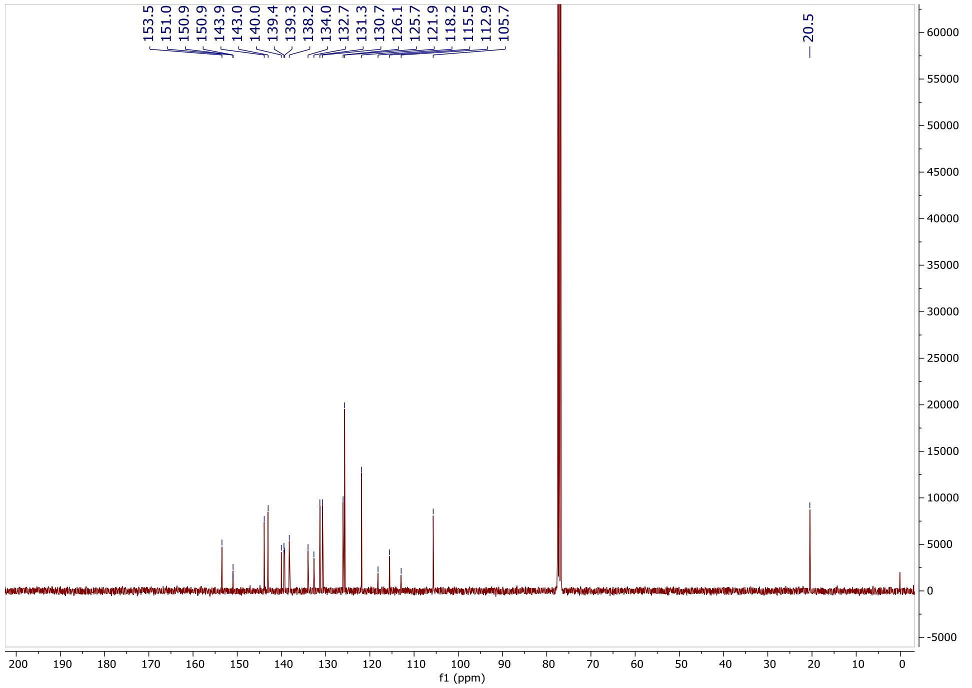
153.5
151.0
150.9
143.9
143.0
140.0
139.4
139.3
138.2
134.0
132.7
131.3
130.7
126.1
125.7
121.9
118.2
115.5
112.9
105.7
20.5
f1 (ppm)

**1-(4-(Difluoromethoxy)phenyl)-6-(2-methyl-4-(trifluoromethyl)phenyl)-1*H*-imidazo[4,5-*c*]pyridine (OXS008161)**

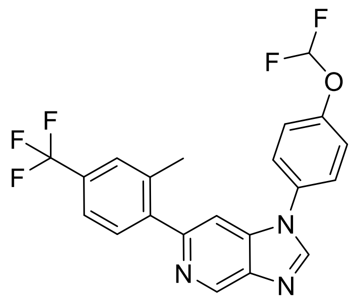

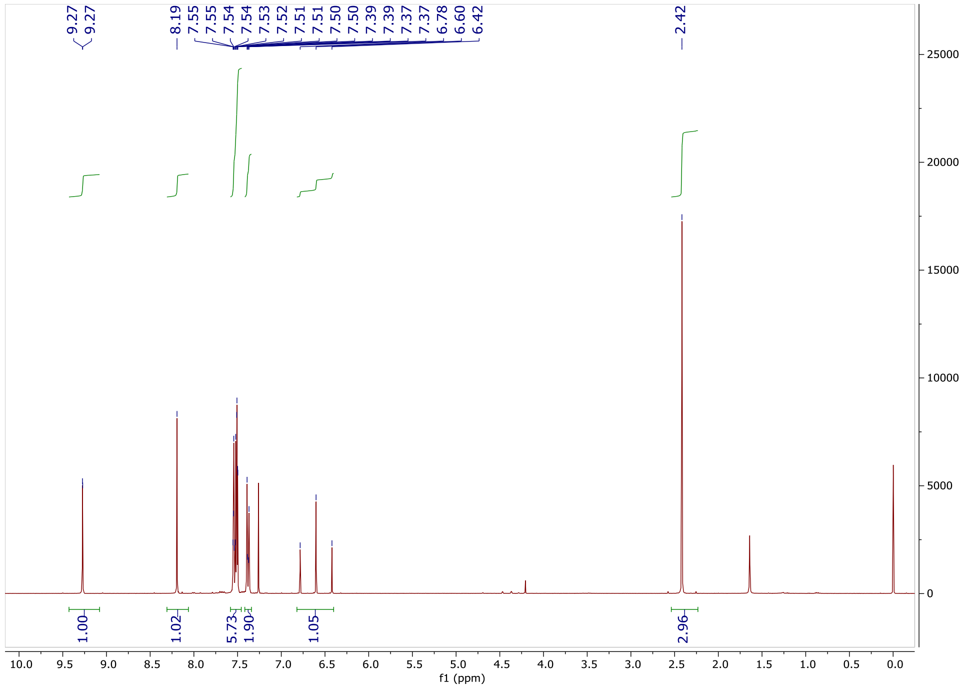

9.27
9.27
8.19
7.55
7.55
7.54
7.54
7.53
7.52
7.51
7.51
7.50
7.50
7.39
7.39
7.37
7.37
6.78
6.60
6.42
2.42
1.00
1.02
5.73
1.90
1.05
2.96
f1 (ppm)

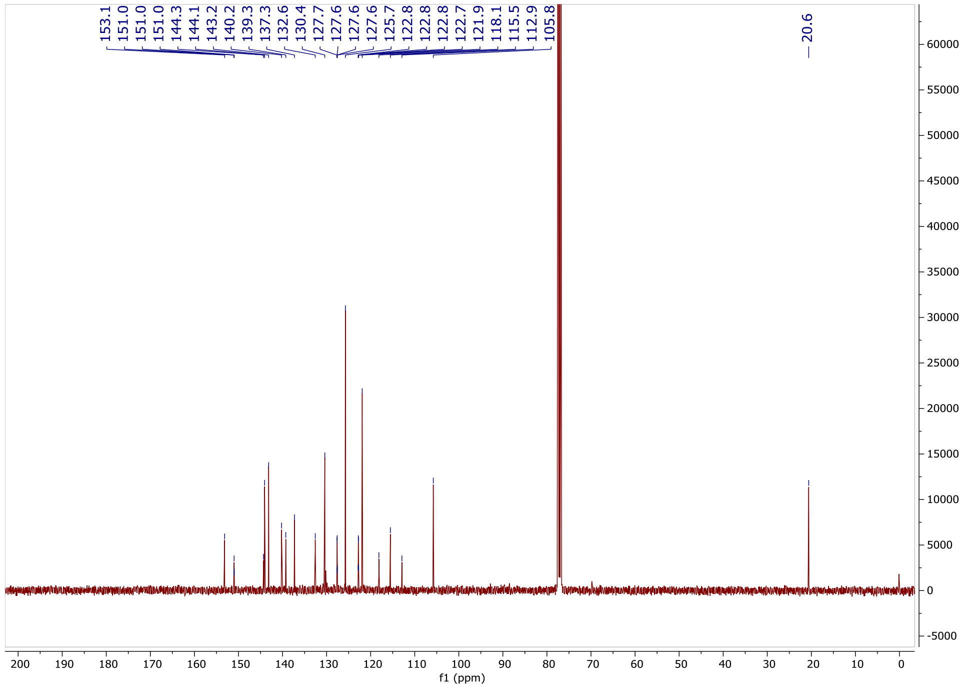

153.1
151.0
151.0
144.3
144.1
143.2
140.2
139.3
137.3
132.6
130.4
127.7
127.6
127.6
125.7
122.8
122.8
122.8
122.7
121.9
118.1
115.5
112.9
105.8
20.6
f1 (ppm)

**4-{1-[4-(Difluoromethoxy)phenyl]-1*H*-imidazo[4,5-*c*]pyridin-6-yl}-3-methylphenol (OXS008187)**

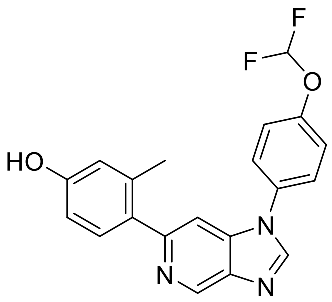
F
F
O
HO
N
N
N

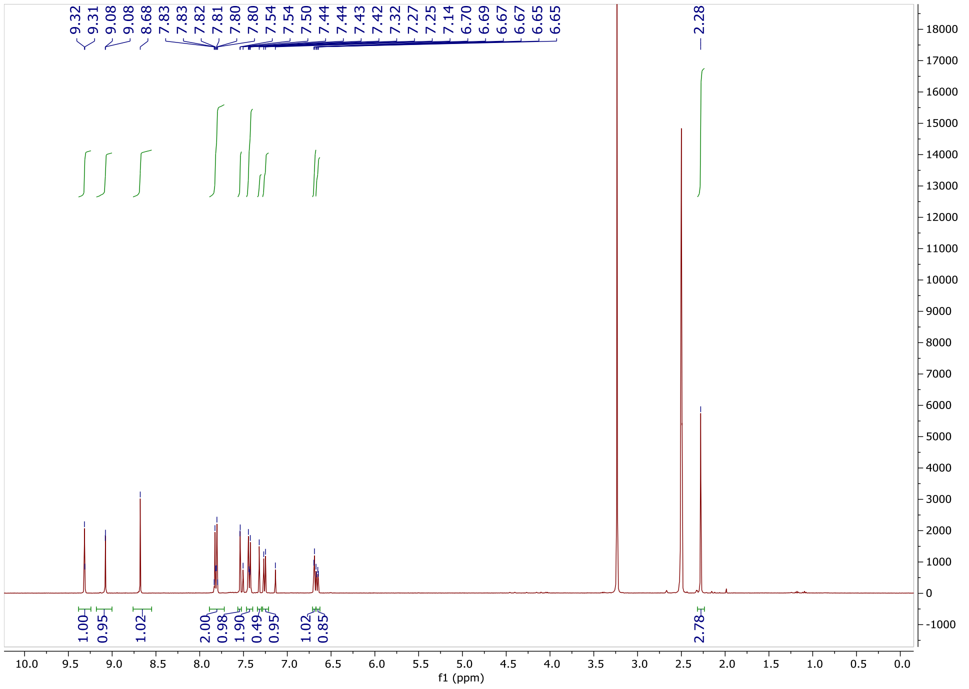
9.32
9.31
9.08
9.08
8.68
7.83
7.83
7.82
7.81
7.80
7.80
7.54
7.54
7.50
7.44
7.44
7.43
7.42
7.32
7.27
7.25
7.14
6.70
6.69
6.67
6.67
6.65
6.65
2.28
1.00
0.95
1.02
2.00
0.98
1.90
0.49
0.95
1.02
0.85
2.78
f1 (ppm)

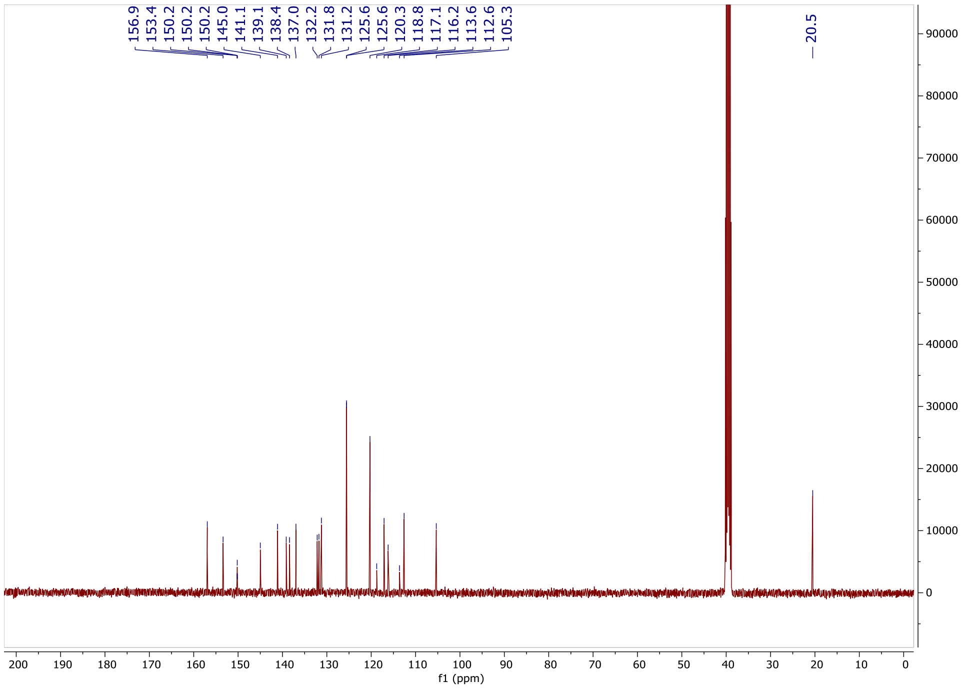
156.9
153.4
150.2
150.2
150.2
145.0
141.1
139.1
138.4
137.0
132.2
131.8
131.2
125.6
125.6
120.3
118.8
117.1
116.2
113.6
112.6
105.3
20.5
f1 (ppm)

**6-(4,5-Difluoro-2-methylphenyl)-1-(4-(difluoromethoxy)phenyl)-1*H*-imidazo[4,5-*c*]pyridine (OXS007733)**

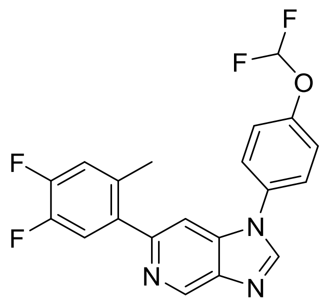

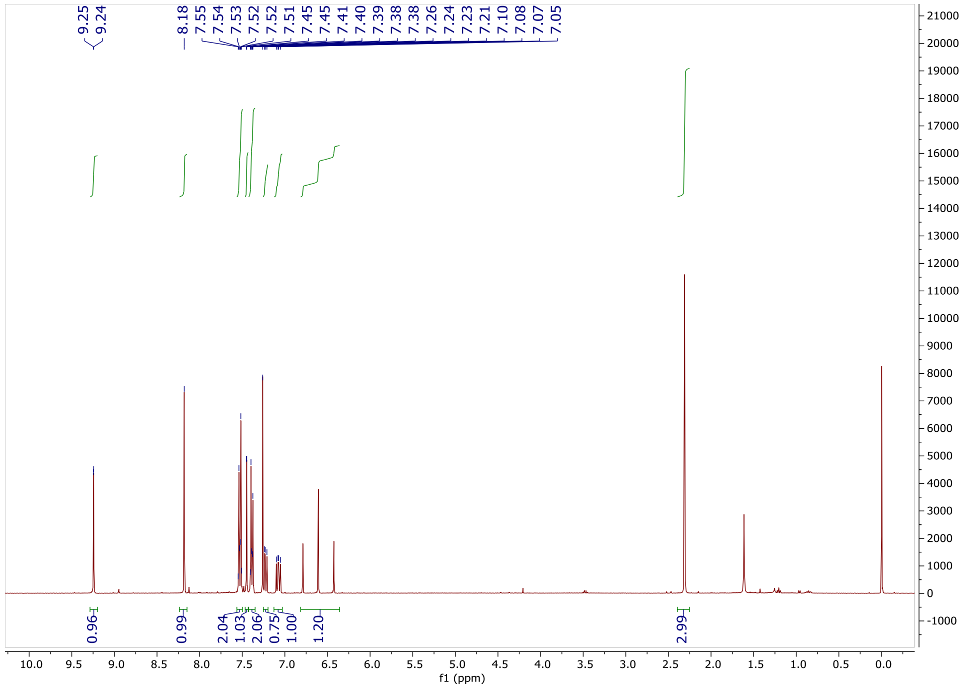

9.25
9.24
8.18
7.55
7.54
7.53
7.52
7.52
7.51
7.45
7.45
7.41
7.40
7.39
7.38
7.38
7.26
7.24
7.23
7.21
7.10
7.08
7.07
7.05
0.96
0.99
2.04
1.03
2.06
0.75
1.00
1.20
2.99
f1 (ppm)

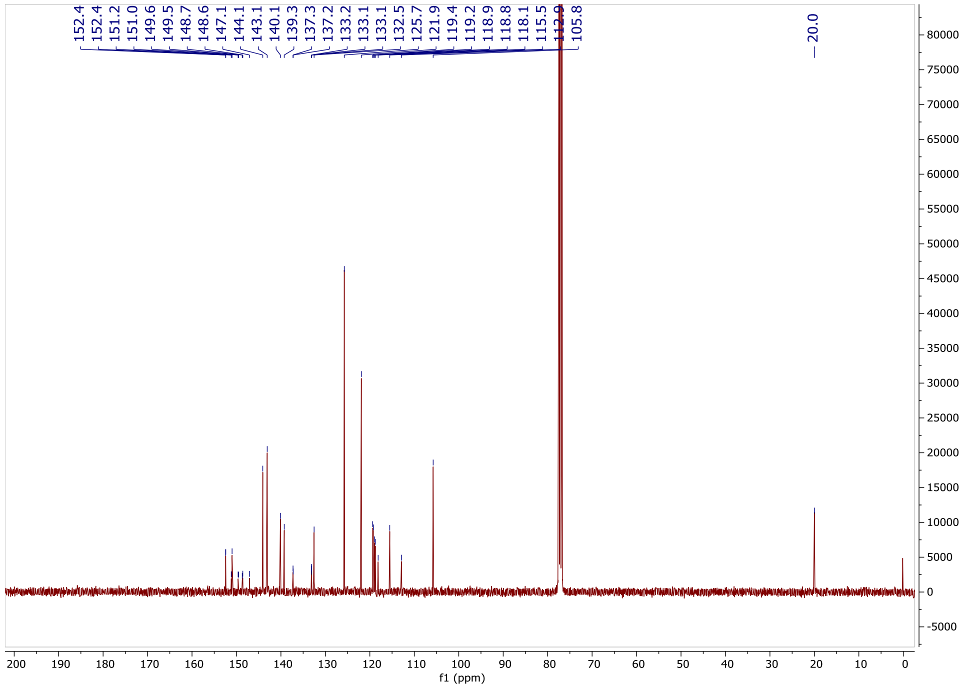

152.4
152.4
151.2
151.0
149.6
149.5
148.7
148.6
147.1
144.1
143.1
140.1
139.3
137.3
137.2
133.2
133.1
133.1
132.5
125.7
121.9
119.4
119.2
118.9
118.8
118.1
115.5
105.8
20.0
f1 (ppm)

**1-(4-(Difluoromethoxy)phenyl)-6-(6-fluoro-2-methylpyridin-3-yl)-1*H*-imidazo[4,5-*c*]pyridine (OXS007865)**

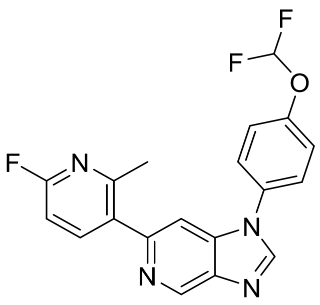

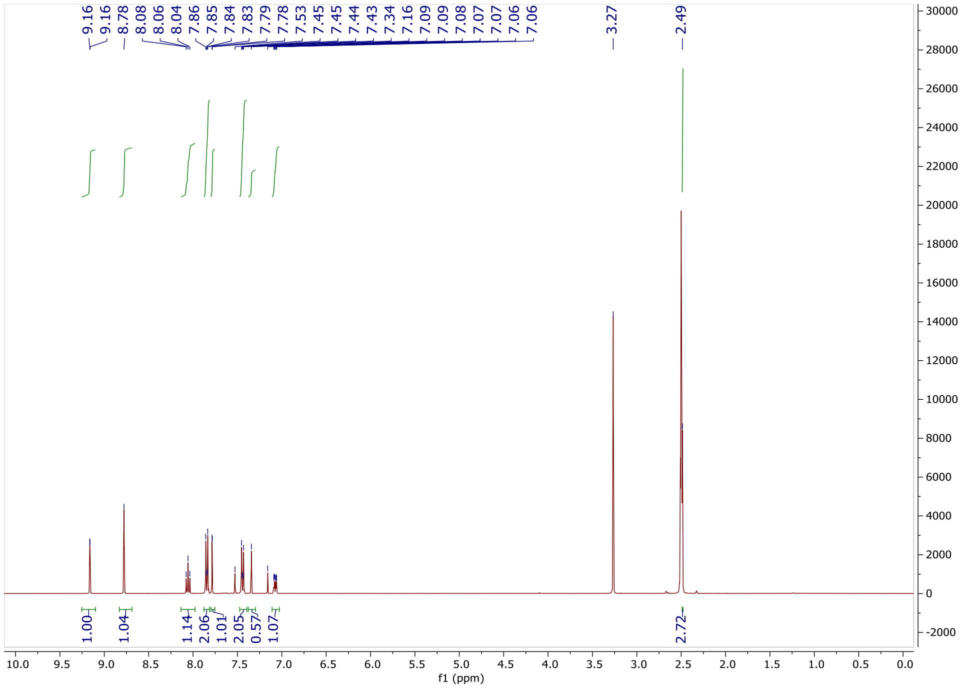
9.16
9.16
8.78
8.08
8.06
8.04
7.86
7.85
7.84
7.83
7.79
7.78
7.53
7.45
7.45
7.44
7.43
7.34
7.16
7.09
7.09
7.08
7.07
7.07
7.06
7.06
3.27
2.49
1.00
1.04
1.14
2.06
1.01
2.05
0.57
1.07
2.72
f1 (ppm)

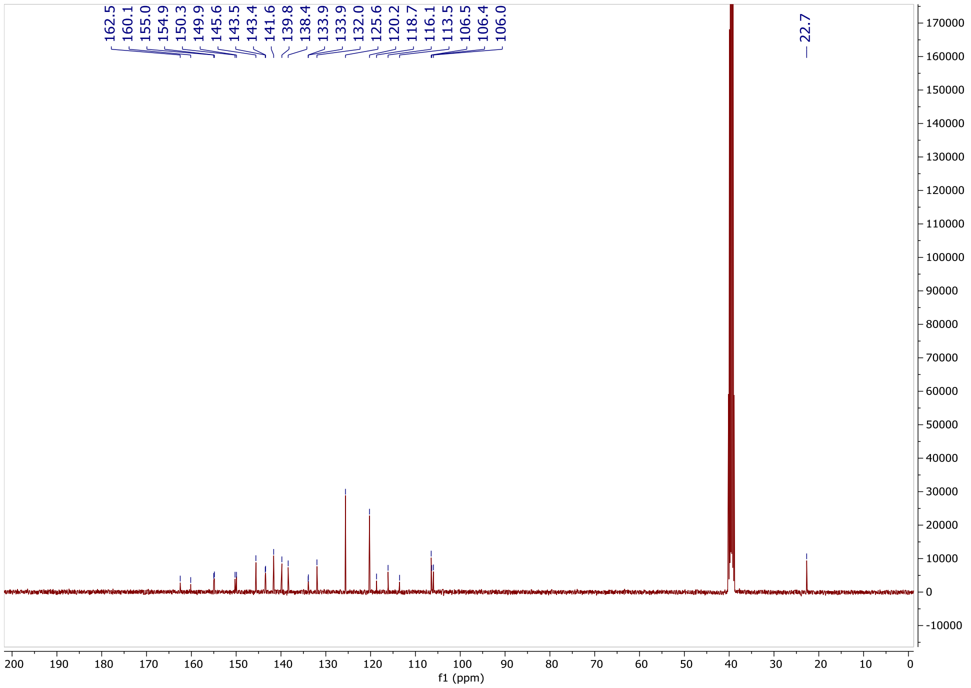
162.5
160.1
155.0
154.9
150.3
149.9
145.6
143.5
143.4
141.6
139.8
138.4
133.9
133.9
132.0
125.6
120.2
118.7
116.1
113.5
106.5
106.4
106.0
22.7
f1 (ppm)

**1-(4-(Difluoromethoxy)phenyl)-6-(6-fluoro-4-methylpyridin-3-yl)-1*H*-imidazo[4,5-*c*]pyridine (OXS007864)**

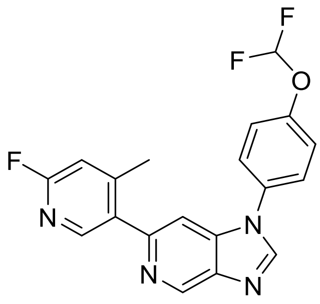

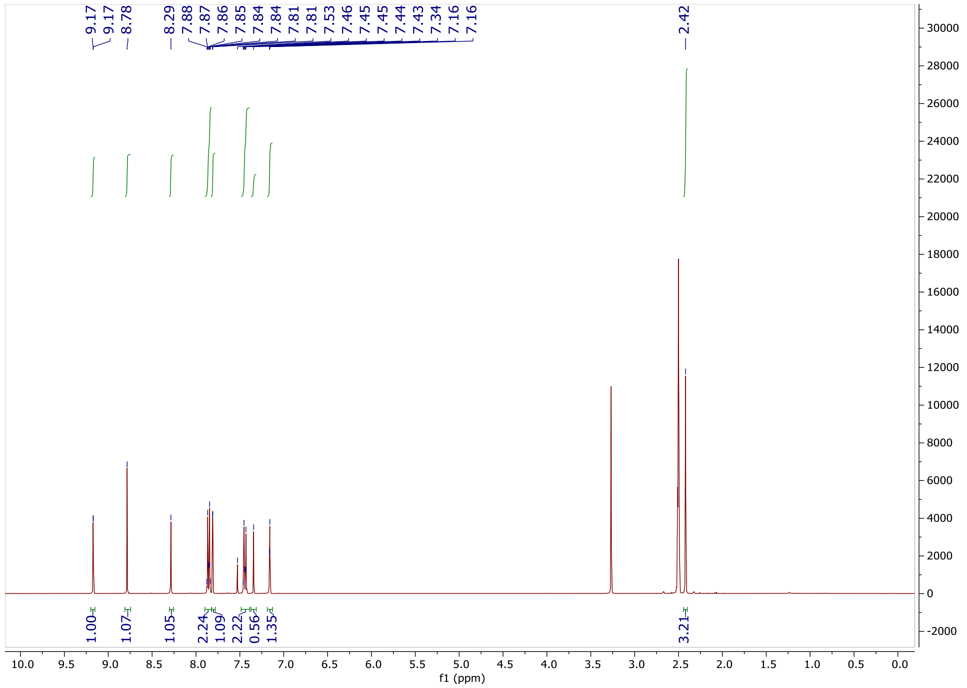
9.17
9.17
8.78
8.29
7.88
7.87
7.86
7.85
7.84
7.84
7.81
7.53
7.46
7.45
7.45
7.44
7.43
7.34
7.16
7.16
2.42
1.00
1.07
1.05
2.24
1.09
2.22
0.56
1.35
3.21
f1 (ppm)

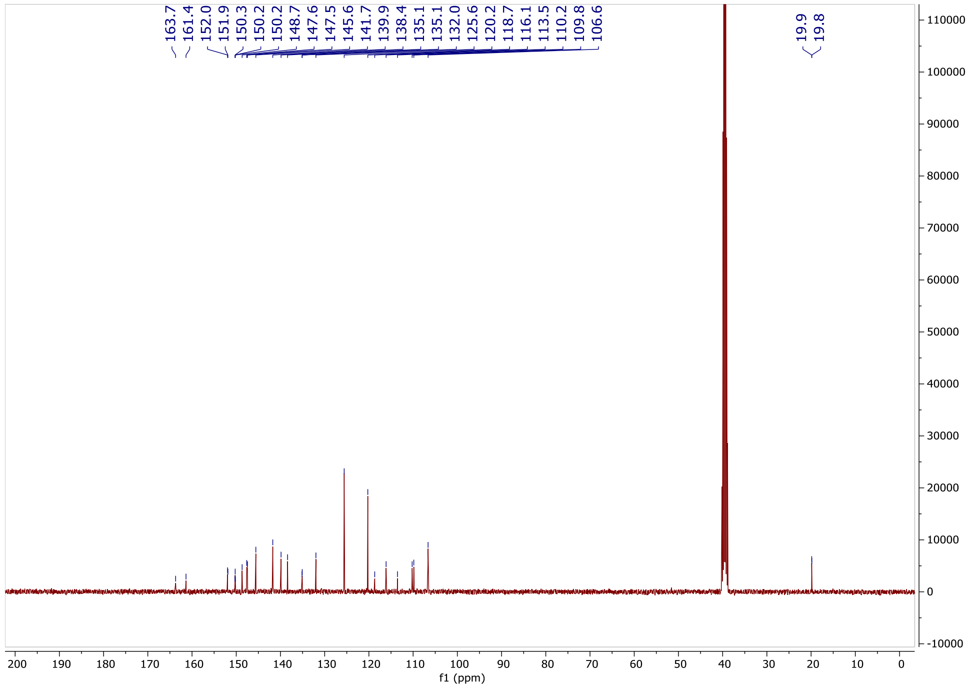
163.7
161.4
152.0
151.9
150.3
150.2
148.7
147.6
147.5
145.6
141.7
139.9
138.4
135.1
135.1
132.0
125.6
120.2
118.7
116.1
113.5
110.2
109.8
106.6
19.9
19.8
f1 (ppm)

### 1-(4-(Difluoromethoxy)phenyl)-6-(4-fluoro-2-methylphenyl)-2-methyl-1*H*-imidazo[4,5-*c*]pyridine (OXS007752)

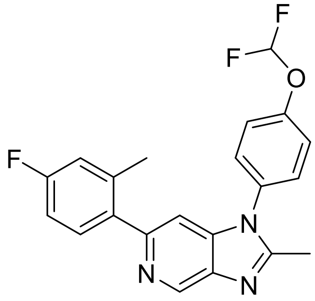

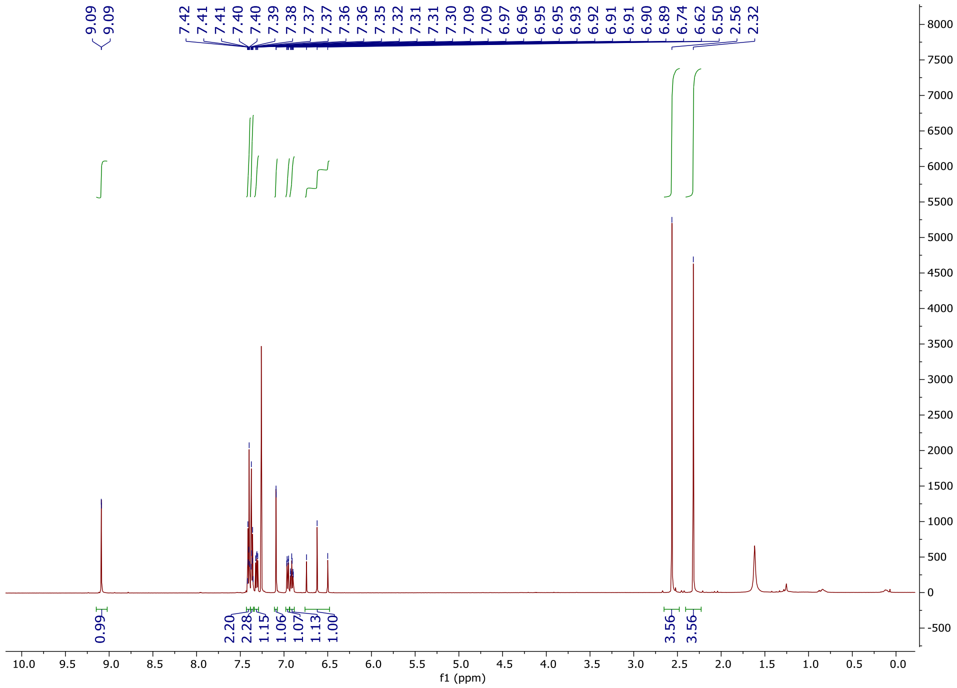

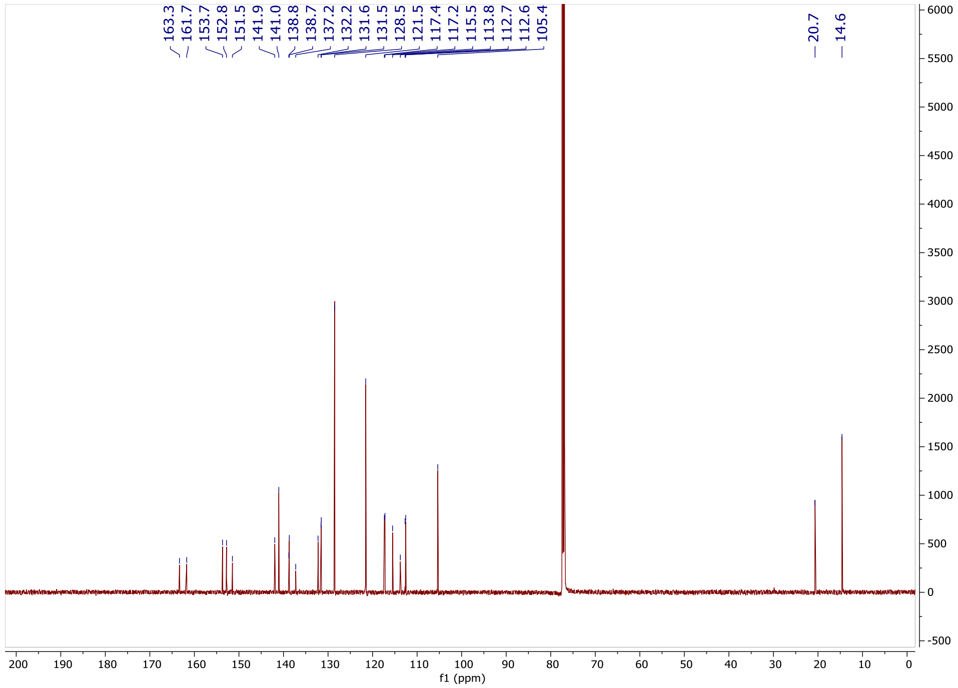

163.3
161.7
153.7
152.8
151.5
141.9
141.0
138.8
138.7
137.2
132.2
131.6
131.5
128.5
121.5
117.4
117.2
115.5
113.8
112.7
112.6
105.4
20.7
14.6
f1 (ppm)

**1-[4-(Difluoromethoxy)phenyl]-6-(4-fluoro-2-methyl-phenyl)-2-phenyl-imidazo[4,5-*c*]pyridine (OXS007768)**

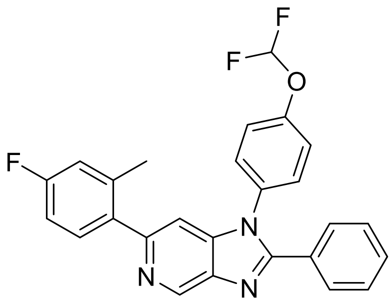

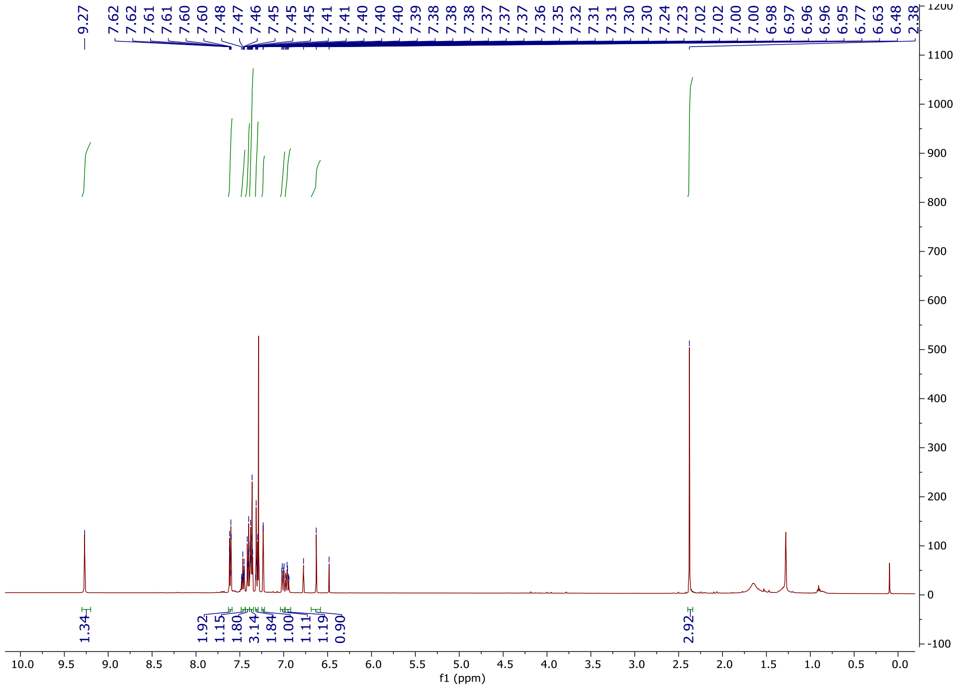
9.27
7.62 7.62 7.61 7.61 7.60 7.60 7.48 7.47 7.46 7.45 7.45 7.45 7.41 7.41 7.40 7.40 7.40 7.39 7.38 7.38 7.38 7.37 7.37 7.37 7.36 7.35 7.32 7.31 7.31 7.30 7.30 7.24 7.23 7.02 7.02 7.00 7.00 6.98 6.97 6.96 6.96 6.95 6.77 6.63 6.48 2.38
1.34 1.92 1.15 1.80 3.14 1.84 1.00 1.11 1.19 0.90 2.92
f1 (ppm)

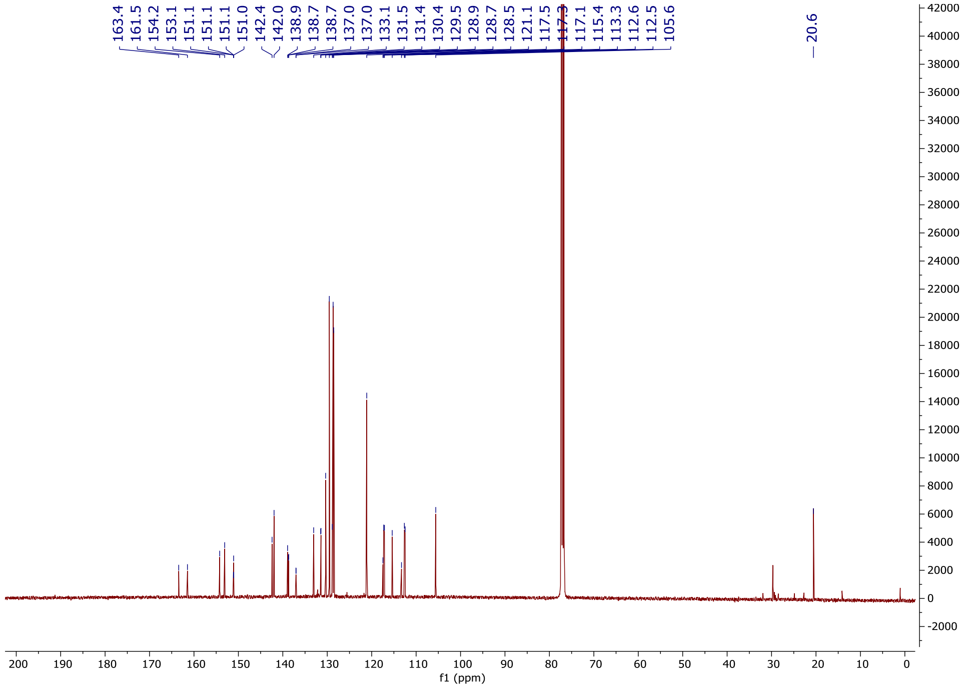
163.4 161.5 154.2 153.1 151.1 151.1 151.1 151.0 142.4 142.0 138.9 138.7 138.7 137.0 137.0 133.1 131.5 131.4 130.4 129.5 128.9 128.7 128.5 121.1 117.5 117.3 117.1 115.4 113.3 112.6 112.5 105.6
20.6
f1 (ppm)
